# The FBA Solution Space Kernel – Introduction and Illustrative Examples

**DOI:** 10.1101/2024.06.10.598372

**Authors:** Wynand S. Verwoerd, Longfei Mao

## Abstract

**Background:** **T**he solution space of an FBA-based model of cellular metabolism, can be characterised by extraction of a bounded, low dimensional kernel (the SSK) that facilitates perceiving it as a geometric object in multidimensional flux space. The aim is to produce an amenable description, intermediate between the single feasible extreme flux of FBA, and the intractable proliferation of extreme modes in conventional solution space descriptions. Fluxes that remain fixed are separated off while the focus of interest is put on the subset of variable fluxes that have a nonzero but finite range of values. For unbounded fluxes, a finite subrange that geometrically corresponds to the variable flux range is determined and is supplemented by a limited set of rays that encapsulates their unbounded aspects. In this way the kernel emphasises the realistic range of flux variation allowed in the interconnected biochemical network by e.g. limited nutrient uptake, an optimised objective and other model constraints. This work builds on the full presentation of the SSK approach in a research monograph.

**Methods:** Calculations are performed with the publicly available software package SSKernel, the source code and user manual of which is included as a supplementary file.

**Results:** It is demonstrated how knowledge of the SSK and accompanying rays can be exploited to explore representative flux states of the metabolic network. Noting that bioengineering interventions such as gene knockouts modify the solution space, new tools based on the SSK analysis are presented here that predict the effects of such interventions on a target flux constructed to represent a desired metabolic output. A simple metabolic model is used first to demonstrate the special concepts and constructions needed to define and compute the SSK. The demonstration model is tweaked to produce typical behaviours of larger models, but with kernels in 1,2 or 3 dimensions that are explicitly displayed to visualise the concepts. General applicability to models where visualisation is inaccessible, is illustrated by showing evaluation of potential bioengineering strategies for a genome scale model.

**Conclusions:** SSKernel is a flexible interactive tool that facilitates an overview of the FBA solution space as a multidimensional geometric object, in terms of a manageable number of parameters. It allows exploration of effects on this solution space from metabolic interventions, and can be used to investigate bioengineering strategies to manipulate cellular metabolism.

## 1 Background

A fundamental premise of the constraint-based modelling of cellular metabolism is that stoichiometric, thermodynamic, and capacity constraints limit the vector of flux values for biochemical reactions in a cell to a feasible region (the solution space or SS) within the flux space created by taking each reaction flux as an independent axis.

Flux balance analysis (FBA) [1], [2] takes this one step further, assuming that the metabolic state of a cell is a steady state that is well represented by the flux vector that optimises an appropriately chosen objective such as the biomass growth rate. With linear constraints and objectives, solving for an optimal flux value is a standard linear programming (LP) [3] problem.

However, even though realistic models remain extensively underdetermined even after an objective is fixed, LP only delivers a single optimal flux value. In contrast, extreme pathway or elementary mode analysis [4], [5], [6] gives an exact mathematical description of all feasible flux values by finding a vector basis that completely spans the solution space. In particular, the FBA solution space is a convex polytope in the flux space, and this basis consists of the vectors that define the positions of the vertices of the polytope.

Neither of these approaches provide a readily utilizable description of the flux or range of fluxes available to a cell. In the case of FBA, this is because the single flux it yields cannot be taken as typical; for the simplex LP algorithm, it is, in fact by construction located at a vertex, i.e., at the very extreme of the solution space. For extreme pathways, the problem is that the sheer number of basis vectors becomes overwhelmingly large for realistic metabolic models.

A popular way to address this difficulty is an elaboration of FBA called flux variability analysis (FVA) [7]. This consists of using LP to find the minimum and maximum value that each reaction flux can achieve within the model constraints. This fixes a cuboid bounding box in the multidimensional flux space, within which the solution space lies. Numerical difficulties are often encountered in performing FVA [8], but a more fundamental limitation is that in high-dimensional spaces, the solution space polytope occupies a negligible fraction of the bounding box, rendering the FVA box uninformative. This point is further discussed below.

The solution space kernel (SSK) approach [9] steers midway between the extremes and pitfalls described. It expands on FBA by specifying the range of feasible flux values rather than a single value. It characterises this region in a way that is far more specific and informative than FVA is, but in terms of far fewer parameters than the extreme pathway/elementary mode approach. In direct contrast to the FVA bounding box, every flux value that falls in the SSK is an actual feasible flux. Also, the SSK approach specifically provides for describing unbounded solutions spaces that are commonly observed in metabolic models. In such a SS there is one or more directions in flux space in which a flux vector of arbitrary length remains feasible; a unit vector in such a direction is defined as a *ray vector*.

This goal is achieved by identifying a compact, low-dimensional subset of the SS (a polytope) from which most, if not all, feasible fluxes can be reached mathematically by the addition of a linear combination of a limited number of ray vectors that are also identified. This polytope is termed the *kernel* of the SS. Even when the full SS is unbounded in some directions, the kernel remains bounded. It is specified by a small number of constraints in far fewer dimensions than the full SS.

To enhance comprehension, the SSK is treated as a geometrical object in multidimensional space with a shape described by a set of mutually orthogonal chords of maximal length, and the aspect ratios given by their quotients. Even more concretely, a list of explicit flux vectors that specify the centre and chosen periphery points of the SSK, allows the expected values and ranges for flux-dependent attributes of cell metabolism to be evaluated.

This paper clarifies these concepts by detailing their application to a simple metabolic network. No attempt is made here to explain the extensive computation that is needed to construct an SSK; more details are available in the work referenced in the next section.

However, in broad outline, the major stages are as follows:

1. All reaction fluxes that remain fixed over the entire SS are separated off.
2. Identify directions in flux space for which the SS is unbounded and find corresponding ray vectors.
3. Identify bounded faces of the SS polytope, then introduce additional *capping constraints* that cap ray vectors without truncating the bounded faces. This partitions the SS into a bounded kernel (the SSK) and an unbounded conical section.
4. The extent and shape of the SSK are delineated by finding a set of mutually orthogonal, maximal chords spanning the SSK.
5. The center and periphery of the compact kernel are determined, which further characterizes the SSK location, size and shape in a reduced-dimensional space.

Stage 5 allows an even more amenable approximation of the SSK for cases where it still has a sizable dimensionality *N*. A collection of representative periphery points is explicitly determined in this stage, and their convex hull delineates a central region that typically covers 80% or more of the full extent of the SSK. This is termed the peripheral point polytope (PPP) that by construction has only in the order of 4N to 6*N* explicitly listed vertices that can be used as a “poor man’s version” of extreme pathways, to specify the most physically significant central part of the SSK. The central part is considered most significant, as it is least affected by slight variations or uncertainties of the metabolic conditions that give rise to the polytope boundaries.

The solution space (SS) concept needs further discussion. In constraint-based analysis, this is generally considered to be the flux region that satisfies stoichiometry and other external constraints. For FVA, the optimised objective is added as a further constraint, further reducing the region to what is termed the optimal solution space (OS). The distinction is ignored in most of what follows since kernel analysis can be applied to either the SS or OS. However, for most models, the upper limits for individual fluxes are unknown, and the SS is a pointed polyhedral cone whose apex at the flux space coordinate origin represents the zero-flux solution. In this case, the apex point by itself serves as the kernel since all other feasible fluxes can be reached from it by adding a suitable ray vector combination.

For this reason, the SSKernel analysis mostly comes into its own for the OS, which has a more complex shape. A linear objective defines a hyperplane, and the OS is the intersection between this hyperplane and the conical SS. This intersection generally retains some unbounded directions but also acquires bounded faces. These bounded faces embody the biologically significant flux constraints. Emphasizing both sets of conditions that a bounded face satisfies, it is referred to as a feasible, bounded face (FBF). Determining all FBF’s is one of the major computational challenges of the SSK calculation. Once found, the kernel is constructed as the hypervolume bordered by all FBF’s, together with any additional capping constraints needed to partition it from the unbounded section of the OS.

This computational framework can also be exploited to characterise the dimensionality and shape of a pointed cone SS more fully. To do this, its minimal single-point kernel is extended by capping the cone at a chosen radius. This allows the “cross-sectional” shape of the pointed cone to be characterised by chords and aspect ratios in a similar way as is done for the OS.

In this sense, the SS kernel approach can be usefully applied to both the SS and OS, and to simplify terminology, both concepts are henceforth referred to as the solution space.

A more formal treatment of metabolic solution spaces was presented by Klamt et al [10]. The SSK concept may be further clarified by relating it to the terminology used there. In the presence of non-homogenous constraints such as flux bounds or an optimised objective, ref [10] designates the SS as the *flux polyhedron*. If sufficiently constrained, this polyhedron may be bounded and is termed the *flux polytope*.

Associated with the flux polyhedron or polytope, is its *recession cone* which is the polyhedral cone obtained by taking only its homogenous constraints. Considering the flux polyhedron as composed of a flux polytope plus its recession cone, an arbitrary feasible flux can be written a suitable linear combination of a finite set of generators, consisting of all *extreme rays* of the recession cone and all vertex vectors of the polyhedron. Extreme rays, e.g for a 3D cone, would point along each of its edges, and any other ray can be constructed as a convex combination of them.

Constructing the solution space kernel (SSK) is somewhat similar to reversing the recession cone construction. However, it generally starts from the flux polyhedron rather than a flux cone, and constructs a flux polytope that shares all the bounded polyhedron faces (and hence vertices), but the SSK features additional faces and vertices derived from the capping constraints. These serve to separate the unbounded region of the flux polyhedron from the bounded convex polytope, namely the SSK. The key feature of this construction is that capping boundaries are required only to intersect bounded faces of the flux polytope tangentially, i.e. at a vertex. That is what guarantees that all vertices of the flux polyhedron remain vertices of the SSK.

The recession cone decomposition yields a mathematically complete vector basis, but in each of its two main components (the vertices and recession cone extreme rays) the number of members proliferate exponentially with increasing dimensions. In contrast, the SSK uses a halfspace representation of the flux polytope so that it can be fully specified by a small number of constraints, generally only a fraction of those in the original metabolic model. Also, for the unbounded part of the flux polyhedron, it does not use extreme rays, but uses special procedures to find a ray basis with its membership count a low multiple of the dimensions. Unlike extreme rays the resulting ray set is not a complete basis, but the extensive convergence of the iterative ray searching procedure in all genome scale models investigated, builds confidence that the limited basis is adequate to reproduce most rays through convex superposition.

The brief introduction presented above merely touches upon the main SSK concepts, and the main purpose of this article is to further elucidate them by explicit, concrete examples in low dimensions that can be readily visualised. A more leisurely introduction to the solution space kernel approach may be found in the software manual included in the supplementary information attached to this article [See Additional File 3]. Also, a more extensive exposition of how this work relates to existing FBA approaches and literature, is provided in the first chapter of the monograph [9] in which the Solution Space Kernel (SSK) was first introduced.

## 2 Implementation

The computational processing is performed via the software package SSKernel [11], which is readily available as a GitHub release, and its Wolfram Language source code is also provided in the Supplementary Data [See Additional File 3]. This implements an interactive interface that allows the user to perform the major stages as outlined in the previous section, adjusting relevant parameters for each subsequent stage as needed. The software is accompanied by a user manual that describes the concepts and calculations in more detail than is possible here.

To process a metabolic model, it is specified in the format defined by the COBRA toolbox [12] as an *.mat file or alternatively in SBML format.

An interactive run of the SSKernel software produces two sets of outputs. First, there is a 2-page PDF file that summarises the main outcomes of the calculation, such as stage-by-stage dimension reduction and SSK shape information. Second, the full numerical results, including a constraint-based specification of the SSK, explicit ray vectors and peripheral flux points, are exported to an output file in the Data Interchange Format (*.dif). Examples of the input files and two output files for the toy model discussed in section 3 are provided in the supplementary data [See Additional File 2].

Also included in the SSK software are tools for exploring the solution space and how it is changed by bioengineering interventions. These are used independently of the interactive interface for postprocessing and take the abovementioned DIF file as input. The realistic network example discussed in section 3.2 is based on processing the DIF output file of an SSK run on the red blood cell model iAB_RBC_283 [13].

The calculation of the SSK involves many special concepts and algorithms for dealing with polytopes in high-dimensional spaces, and a full exposition has been presented in a recent monograph [9]. However, the exploratory tools are a subsequent development, and the formal details of the new algorithms they use are presented in the Supplementary Information [See Additional File 1].

## 3 Results and discussion

The results are presented in two main parts.

First, the main features of the SSK are illustrated on the basis of a simple model system. This is a toy model formulated in a 10-dimensional flux space and allows SSK reductions to 2 or 3 dimensions so that they can be graphically displayed. The emphasis here is on transparency. Such a simple model does not demonstrate the full power of simplification that SSK analysis is capable of. For that, the reader is referred to the dedicated monograph [9], which discusses applications to state-of-the-art models involving thousands of flux space dimensions.

Second, it is shown how the SSK and rays can be used to explore the solution space and its response to flux restrictions. The toy model illustrates the concepts, and an application to a realistic metabolic model is also shown to demonstrate its practical usefulness.

### 3.1 Elucidating SSK Concepts

The toy network used here originates from a textbook demonstration of extreme pathways [14] and is shown in Figure 1. Various slight elaborations of the toy model are made below to reproduce several scenarios commonly encountered in models of real cellular metabolism. Explicit input data sets to reproduce the results for all scenarios below are supplied in the Supplementary Information [See Additional File 2].

**Figure 1.**
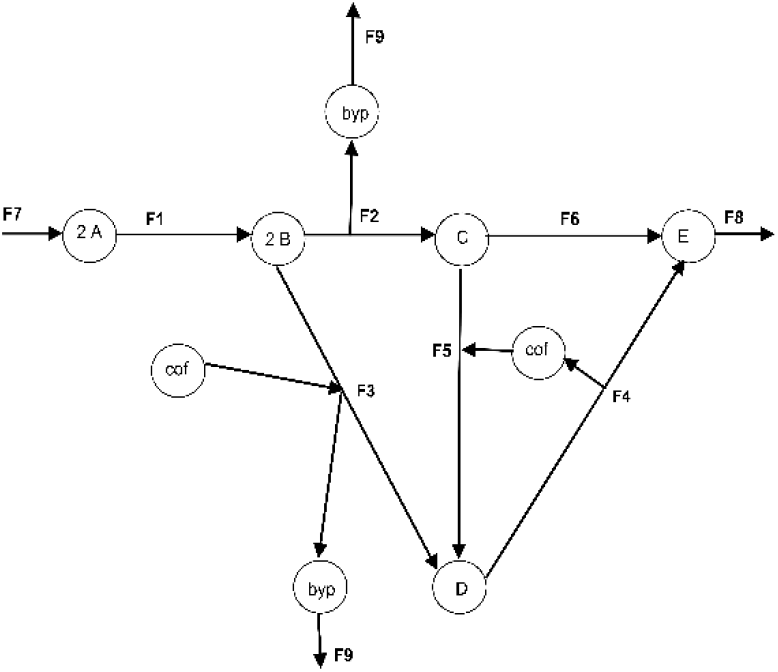
Toy network that produces a product *E* and byproduct *byp* from six internal reactions with fluxes F1 to F6. F7 is an external input flux and F8 and F9 are external output fluxes.

Each of the five SSK calculation stages mentioned in the introduction requires multiple elaborate numerical algorithms, specially designed for the purpose. It would not be practical to describe them here, but instead the 2D and 3D plots of the SS presented below for each scenario are sufficient to visually trace those steps and confirm that the computation yields the required results. For the first stage in particular, finding fixed fluxes, FVA calculation is quick and sufficient for the simple toy model, so is used as a familiar point of departure although a more efficient special algorithm is used in the SSKernel implementation.

#### 3.1.1 Scenario 1: A bounded SS, with its SSK and FVA

To set the scene, the toy model is first specialized to a case where only 3 fluxes have variable values over the extent of the SS. This is the result of introducing and optimizing a further biomass flux F10 (discussed further for Scenario 4 below) and also introducing a fixed value for F2 and an upper limit for the input flux F7. Then fluxes F1,F2, F3, F7, F8, F9 and F10 all become fixed to specific values, and feasible flux points hence fall in a 3D hyperplane within the 10D flux space, spanned by the flux axes F4, F5 and F6 and which is graphically displayed in Figure 2.

**Figure 2.**
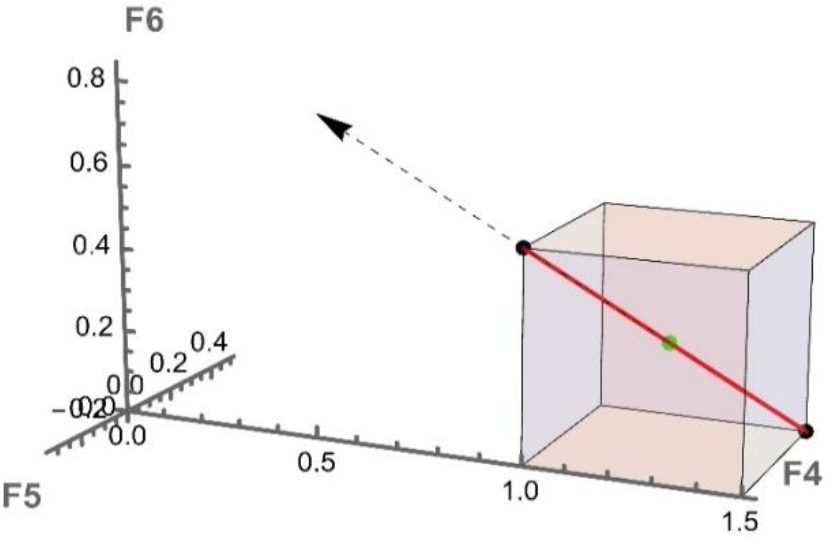
The 1D linear SSK (red line) and 3D FVA bounding box (in colour) for the toy model, with an optimised objective but constrained to leave only F4, F5 and F6 variable, while other fluxes acquire fixed nonzero values.

**Figure 3.**
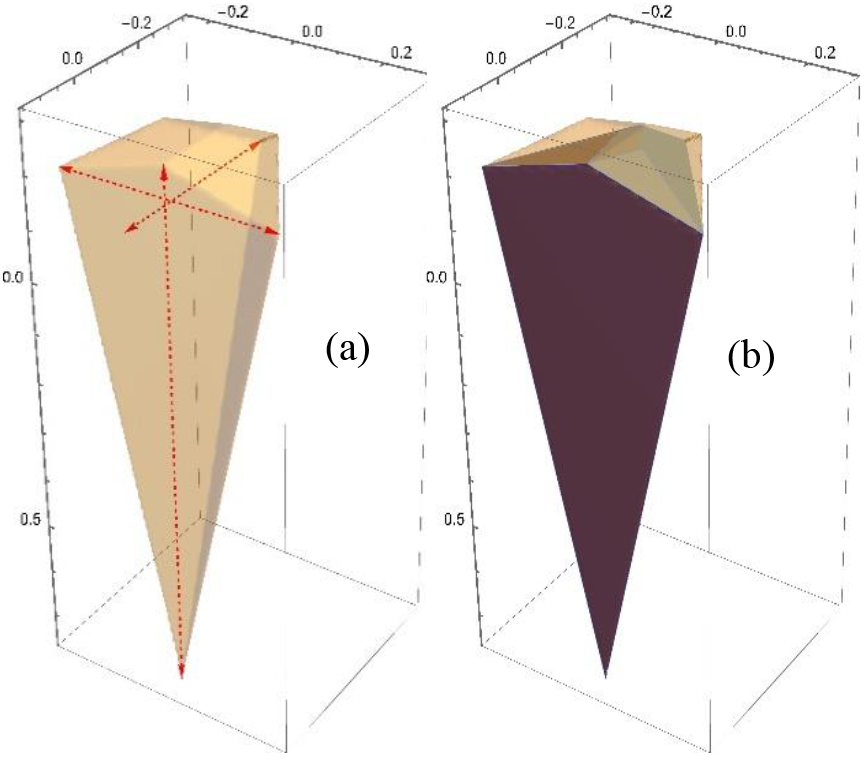
(a) The capped SSK for the toy model without objective. The main chords are shown as red arrows. (b) The periphery point polytope (PPP) superimposed on the SSK (darker colour) shows coverage of most but not all of its volume.

In this case, it turns out that the solution space is bounded, so the calculated SSK that is shown in the figure as a red line section, coincides with the SS. Also shown on the figure is the cuboid bounding box that represents the FVA limits on individual flux ranges. Clearly this bounding box provides a very poor rendering of the actual extent of feasible flux solutions. By contrast the SSK shows this exactly, namely, as just a single line, the diagonal of the cuboid box. This only occupies a negligible fraction of the FVA box volume; thus the box overestimates the flux variability, since e.g. random sampling from the cuboid is unlikely to reveal a single feasible flux combination.

Such dimensional discrepancy between the bounding cuboid and the SSK occurs also in the scenarios below and is a universal feature of realistic metabolic models. For those the FVA overestimation is far worse, because even in cases where the bounding box occupies the same dimensions as the SSK, the fraction of the box that the SSK occupies is still negligible. This is because of a typical high aspect ratio of the SSK and the scaling effects in high dimensions mentioned in the introduction and discussed in more detail in [9].

The FVA limits do provide a familiar indication of flux ranges that is helpful to discover fixed fluxes, but we note that the Hop, Skip and Jump algorithm incorporated in the SSK calculation [9] performs this task faster by orders of magnitude and also more reliably. These advantages are not manifested in the small toy model

#### 3.1.2 Scenario 2: A simple unbounded cone solution space

While graphically convenient, the coincidence of the SS and SSK in the first scenario might leave the relevance of the concept in doubt. To explore how SSK features are distinct, this and subsequent scenarios presents systematic variations on the model from the simplest to those chosen to demonstrate a specific feature. Starting with the bare toy model, it is representative of realistic models that contain no objective. In the conventional description, the metabolic states of this network are superpositions of its three distinct extreme pathways [14], i.e., its SS is a 3-sided infinite cone in 9 flux dimensions.

The SSK analysis uses a default capping radius, as mentioned above, to reduce this to a finite kernel, which is explicitly specified by 6 constraints on 3 variables. The reduction from the 16 constraints on 9 variables of the original model merely hints at the large reductions in realistic models.

**Error! Reference source not found**. shows a 3D plot of the SSK. The SSK algorithm finds 3 peripheral rays aligned along the 3 long edges of the cone, each yielding one capping plane visible at the top in the figure. In this particular case these ray directions are identical to the extreme pathways listed in Figure 13.1 of [14], but that is not guaranteed in more complex models.

The coordinate axes in **Error! Reference source not found**. represent the particular 3D subspace within the full 9D flux space, in which the model allows feasible metabolic fluxes. Specifically, they are by construction the directions of the main orthogonal chords of the SSK, shown as red arrows in the figure. As such, the axes point in special directions in the 9D space, as determined by the SSK shape, and this is the reason why axes are not explicitly labeled in **Error! Reference source not found**. and subsequent figures.

For the simple unbounded cone of this model, the arbitrary capping radius means that the actual chord lengths are not meaningful. However, their ratios are independent of the radius and characterise the shape of the cone. For this model, these aspect ratios are 1.2, 2.1 and 2.5, respectively. The lowest value reflects a fairly regular cross section, and the other values reflect a quite sharp cone, as shown in **Error! Reference source not found**..

#### 3.1.3. Scenario 3: A fully bounded solution space

Next, the model is made slightly more realistic by introducing an upper limit for the input flux F7. FVA shows that this constrains all nine fluxes to finite, nonzero ranges. There are no fixed fluxes, and the FVA bounding box is a cuboid in 9 dimensions.

With this SS already bounded, no rays or capping hyperplanes need to be calculated, and the SSK is in fact identical to the SS for this case, though specified in only 3 dimensions. The SSK shape analysis remains applicable, and the result is very similar to that shown in **Error! Reference source not found**.; only the cone is now truncated by a single plane fixed by the upper limit on F7. Therefore, the SSK is, in this case, given by just 4 constraints on 3 variables.

Provided that the F7 cut-off value has been set at a physically meaningful value, the 3 main chords indicate the range of metabolic flux variation allowed by the model along the 3 available orthogonal directions. Note how this is far more specific than the 9D FVA cuboid within which this truncated cone is contained.

The aspect ratios derived from the chords are 1.5, 1.8 and 2.8, indicating a similarly sharp cone but with a slightly flattened cross section compared with **Error! Reference source not found**..

#### 3.1.4. Scenario 4: Optimised objective in a bounded SS

Moving on to a case more appropriate for the purpose for which the SSK analysis is designed, we introduce an optimised objective into the model while keeping the input flux limitation. A biomass composition of 80% of metabolite *E* and 20% of the byproduct *byp* is assumed and incorporated into the model by adding a further output flux F10 that exports these metabolites in a 4:1 ratio. The objective is taken as maximising F10.

An FVA calculation to characterise the OS with F10 kept fixed to its optimised value, shows that a total of 5 fluxes become fixed, leaving a 5D cuboid as the bounding box for feasible metabolic states.

The OS is now bounded, so no rays or capping is needed, and the SSK is identical to the OS. The calculation algorithm shows that the SSK/OS is only 2-dimensional and is plotted in Figure 4. Clearly, this SSK is simply the intersection of the objective plane with the triangular cone of Figure 4, and as expected, the aspect ratio of 1.4 is similar to that of the cone cross section.

**Figure 4.**
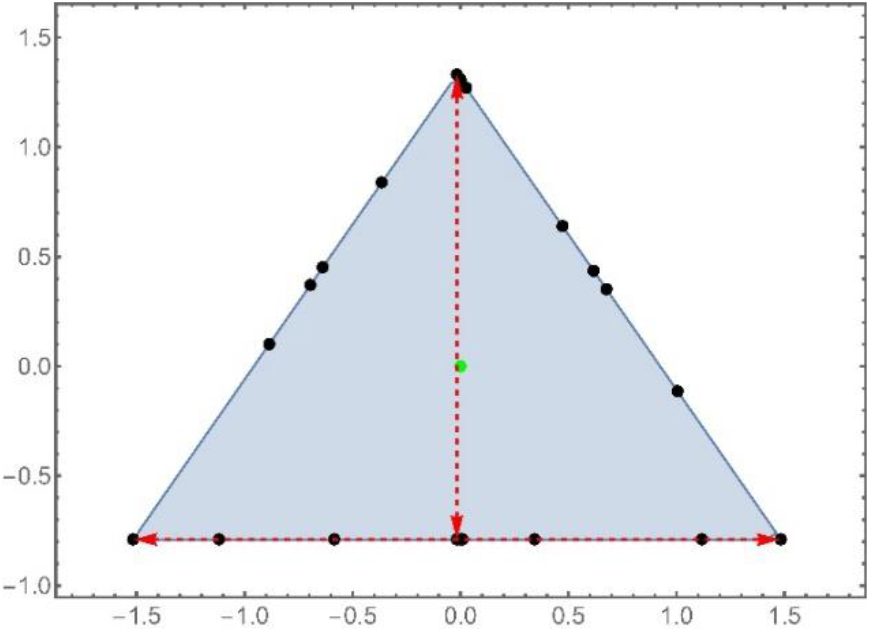
The OS and SSK for the toy model with a restricted input flux and a fixed objective. A centering algorithm determines the green center point and a semirandom selection of periphery points shown in black. The main chords are shown as red arrows.

The figure also illustrates the outcome of the centering procedure incorporated in the SSK calculation. The center is the green dot, which also serves as the coordinate origin in the SSK specification of 3 constraints on 2 variables. Centering also finds a series of periphery points shown as black dots. Visually it is clear that these are not very uniformly distributed, as there are random aspects in the underlying algorithm. Nevertheless, these points form a plausible sample of explicit feasible fluxes that encapsulates the range of variation of metabolic states allowed by the model. In this case their convex hull (the PPP) covers 100% of the SSK, but as shown in **Error! Reference source not found**. (b), this is not generally the case.

#### 3.1.5 Scenario 5: SS partially bounded; tangent capping

The next scenario is chosen to resemble a realistic solution space as closely as possible within the size limitations of the toy model.

This is achieved by putting a lower limit on F2 and allowing F3 to be reversible. The same objective and input flux limitations as in scenario 4 are used. There are again 5 fixed fluxes and 5 variable fluxes, and the feasible solutions are restricted to a 2D plane within this 5D subspace. Unlike the previous scenarios, this *reduced solution space* or RSS (reduced by the exclusion of fixed fluxes) is no longer bounded. Therefore, there are ray directions in the solution space—in fact, only a single ray—the green arrow shown in Figure 5.

**Figure 5.**
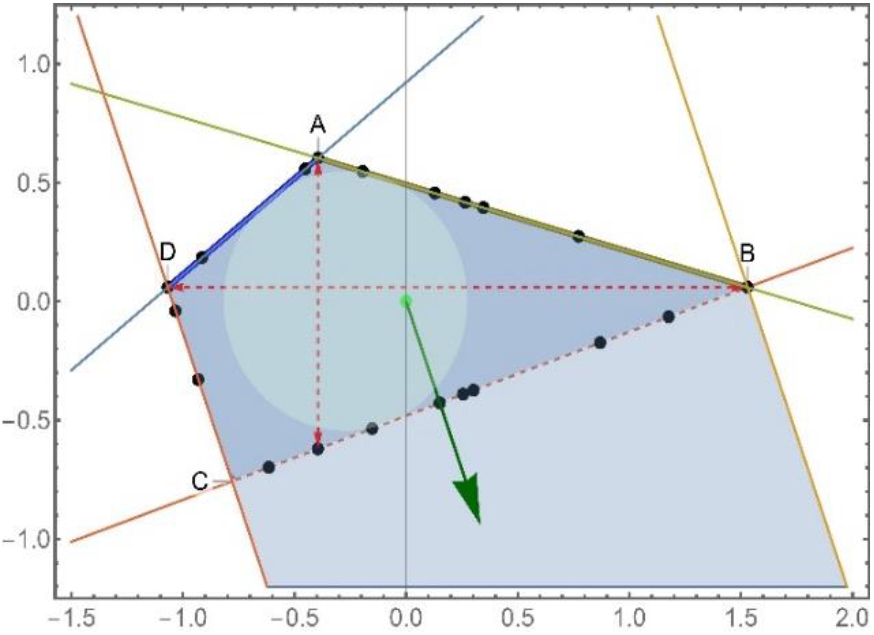
The RSS (light blue shading) and SSK (darker shading) for the toy model with a single reversible reaction. The RSS has two bounded faces, AB and AD. The other two parallel faces remain unbounded and leaves a single ray direction (green arrow). The SSK is bounded by the introduced capping line BC, perpendicular to the ray and only intersecting the bounded faces tangentially at the single point B. The inscribed circle is shown in a lighter shade, but the (green) centroid of the chosen periphery points (black dots) is taken as the coordinate origin and the axes are aligned with the main chords shown as dashed red arrows.

Despite the RSS being unbounded, some of its faces remain bounded, emphasised as thick lines in the figure. In addition to the 2 bounded faces (FBF’s), there are also 2 unbounded faces. To achieve a bounded kernel, the SSK procedure introduces a single *capping hyperplane*, the dotted line BC, that partitions the RSS.

Geometrically, any flux value in the unbounded section can be reached from at least one flux point in the bounded section (the SSK) by adding a multiple of the ray vector.

This is guaranteed by having chosen the capping line such that it does not bisect any of the bounded faces. The capping radius (the perpendicular distance from the origin to the capping line) is chosen to allow the capping line to intercept any bounded face at most at a single point. This procedure is denoted as *tangential capping*.

To grasp the biochemical meaning of the capping, note that the ray vector in fact represents the single cycle created in the network of Figure 1 when reaction 3 was made reversible. This is shown by the calculated 10D ray vector {0, 0.57735, −0.57735, 0, 0.57735, 0, 0, 0, 0, 0}, i.e., only fluxes F2, F3 and F5 participate equally.

This cycle has no net consumption/production of metabolites, so it can support arbitrarily large flux values without violating limits such as the maximum uptake imposed in the scenario under discussion. This means that partitioning off the unbounded section of the RSS by capping simply excludes nonphysical cyclical fluxes. The remaining solutions that define the SSK embody the physically meaningful flux variation as limited by the model constraints.

Note that this partitioning is not unique. The line BD would be an alternative (and minimal) capping line. The key feature is that a viable SSK needs to include all bounded faces (FBF’s), which is why

FBF’s are considered to encompass the physically meaningful constraints that result from the way that individual flux constraints combine across a network. In higher dimensions, minimal tangent capping is hard to compute, so merely capping all ray directions orthogonally (as line BC does in this example) is considered a sufficient strategy.

#### 3.1.6 Scenario 6: Ray space and coincidence capping

This case is similar to scenario 5, but reaction 6 is now also allowed to be reversed. The result, shown in Figure 6, is that only the single bounded face AB of scenario 5 survives. The RSS remains 2D, but there are now two peripheral ray directions aligned with the two unbounded faces. Intermediate rays can be formed by a linear combination of these two rays, i.e., they span the *ray space*.

**Figure 6.**
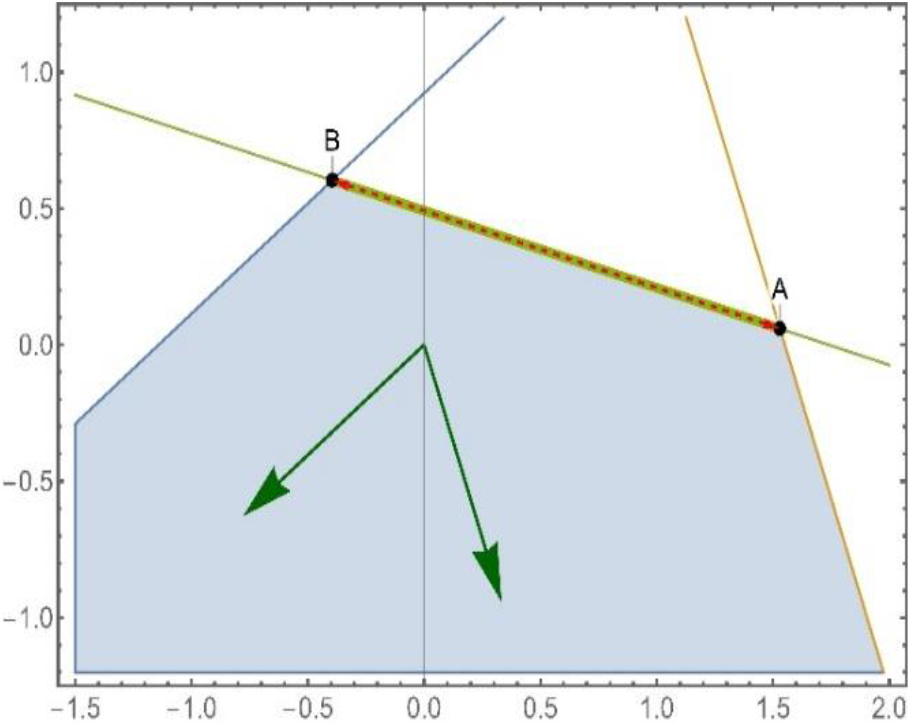
The toy model RSS (unbounded 2D shaded region) and 1D SSK (the line AB) for scenario 6. There are two peripheral ray directions shown as green arrows. The coordinate system of scenario 5 is chosen to facilitate comparison.

It would be possible to define a 2D SSK for this case by taking a capping line perpendicular to each peripheral ray to form a triangle with AB on one side. However, a better alternative is to note that there exists a ray that is perpendicular to line AB. Suppose that this ray is used to define a capping constraint, resulting in a capping line parallel to AB. The capping radius is adjusted until the capping line first intersects the FBF (i.e., line AB itself), which occurs only when the two lines coincide. At that point, the entire shaded region of the RSS is separated off, and only the single, one-dimensional SSK defined by line AB remains.

This is an example of *coincidence capping*. In contrast to tangent capping, it gives an SSK of lower dimensions than the RSS.

With respect to the biochemical interpretation, inspection of the two 10D numerical ray vectors reveals that they represent fluxes that involve (F2, F3, F5) and (F4, F5, F6) respectively.. These paths are easily recognizable from Figure 1 as overlapping but independent cyclical paths formed when F3 and F6 are both reversed.

Rays are, however, not necessarily associated with cycles as happens for the examples shown; e.g., if the toy network contained a linear network path that does not involve the constrained uptake flux F2 or the metabolites that make up the objective, this might also have no upper limit on its flux and thus constitute a ray.

#### 3.1.7 Scenario 7: Coincidence capping of prismatic rays

The RSSs in scenarios 5 and 6 are examples of faceted cones, a term coined to emphasize the presence of bounded faces in addition to the unbounded faces of a simple pointed cone. The rays in Figure 5 and 6 are described as *conical rays*. However, for the full SSK calculation, several additional types of rays are distinguished [9], and the most common type, typically approximately half of all rays for realistic metabolic models, are *prismatic rays*. This type can be induced into the toy model by inserting an additional unidirectional reaction, which involves only some newly added metabolites, into the network. This would add an additional flux axis F11 to the flux space. If the same constraints are used as in scenario 4, the OS shown in Figure 4 is modified by the addition of a vertical axis perpendicular to the plane of the paper. The assumed direction of the reaction limits its feasible fluxes to positive values. Therefore, instead of the 2D triangle shown in the figure, the OS is now a semi-infinite triangular prism that stretches vertically to positive infinity and is bordered by one horizontal constraint plane (the plane of the paper, which constrains F11 to positive values) and three vertical constraint planes.

This modification adds a single ray direction to the OS, which is clearly described as a *prismatic ray*. The distinguishing feature of a prismatic ray is that it is antiparallel to one constraint vector (the one defining the horizontal plane in this example) and orthogonal to all others. Prismatic rays are particularly easy to deal with, as they can be coincidence capped. In this case, that reduces the 3D OS back to the 2D triangular SSK shown in Figure 4.

Similarly, if the constraints of scenarios 5 or 6 are applied to the 11D extension of the toy model, the RSSs in both cases become threedimensional semi-infinite prisms, which are partially unbounded in both the horizontal and vertical directions. They each acquire an additional prismatic ray as a result. Once more, this can be coincidence capped, and then tangent capping applied in the horizontal plane. This produces the same 2D or 1D SSK’s as illustrated in Figures 5 and 6, respectively, despite the extra dimension in the extended model.

#### 3.1.8 Discussion

Table 1 gives a summary of the model modifications and SSK results for the various scenarios, and explicit SSKernel input data for each scenario are provided in Additional File 2.

**Table 1.** Scenario summary, showing modifications of bare toy model to produce various SSK outcomes. Arrow indicates non-zero stoichiometry entries for added reactions, and flux ranges are shown when different from toy model.

| Scenario | Features | Dimension count |  |  | Modifications |
| --- | --- | --- | --- | --- | --- |
|  |  | FS | RSS | SSK |  |
| 1 | SS bounded; 0 rays<br>SSK = SS | 10 | 3 | 1 | $F2 = 0.5, F7 \leq 3$ |
| 2 | SS 3-sided unbounded cone<br>3 conical rays<br>SSK 6-sided polyhedron | 9 | 3 | 3 | Capping radius = 1.0 |
| 3 | SS bounded cone; 0 rays<br>SSK = SS | 9 | 3 | 3 | $F7 \leq 3,$ |
| 4 | OS bounded 2D triangle; 0 rays<br>SSK = OS | 10 | 5 | 2 | $R10 \rightarrow 0.8E + 0.2cof$<br>$F7 \leq 3, F10$ optimized to 1.875 |
| 5 | RSS faceted cone; 1 conical ray<br>SSK 2D polygon | 10 | 5 | 2 | $R10 \rightarrow 0.8E + 0.2cof$<br>$F2 \leq 0.5, -\infty \leq F3 \leq \infty, F7 \leq 3, F10 = 1.875$ |
| 6 | RSS faceted cone; 2 conical rays<br>SSK line section | 10 | 5 | 1 | $R10 \rightarrow 0.8E + 0.2cof$<br>$F2 \leq 0.5, -\infty \leq F3 \leq \infty, -\infty \leq F6 \leq \infty, F7 \leq 3, F10 = 1.875$ |
| 7 | RSS 3D prism<br>2 conical and 1 prismatic ray<br>SSK 2d polygon | 11 | 6 | 2 | $R10 \rightarrow 0.8E + 0.2cof, R11 \rightarrow X,$<br>$F2 \leq 0.5, -\infty \leq F3 \leq \infty, F7 \leq 3, F10 = 1.875, 0 \leq F11 \leq \infty$ |
| cof target | RSS faceted cone; 1 conical ray<br>SSK 2D polygon | 10 | 5 | 2 | $R3 \rightarrow -2B + D + byp -2cof, R4 \rightarrow -D + E + 2cof$<br>$R10 \rightarrow 0.8E + 0.2cof$<br>$F2 \leq 0.5, -\infty \leq F3 \leq \infty, F7 \leq 3, F10 = 1.875$ |

The toy model scenarios that were sketched reproduce in isolation some of the features encountered in full-scale metabolic models. However, in a realistic model, multiple instances are usually combined, such as the simultaneous occurrence of multiple ray types and capping methods along different dimensions.

From a computational point of view, realistic higher-dimensional solution spaces present many complications not encountered in the simple toy model.

An example is that polytope faces can be classified into multiple levels, similar to the boundary planes, edges and vertices of a 3D polytope. In fact, for realistic models, the boundaries formed by the original model constraint hyperplanes are invariably unbounded, and bounded faces occur only at much higher levels. This requires an extensive computational framework for their detection and utilisation to construct appropriate tangent capping. A combinatorial proliferation of faces at higher levels makes this challenging, but it turns out that with appropriate algorithms, it is possible to reduce most models to a fairly low-dimensional SSK with a moderately sized accompanying set of ray vectors that suffice to specify the FBA solution space. That is, for example, the case for the red blood cell model discussed in the next section.

The leap in complexity is illustrated by the seemingly simple task of finding a centre point of a multidimensional polytope. Even in 2D, thousands of different definitions have been proposed just for a triangle and several are in common use. Trials show that these mostly do not generalize well to higher dimensions or to awkward shapes, e.g. with the high aspect ratios commonly found for metabolic solution spaces. A recurring problem is that the calculated centre can fall outside the polytope, unless explicitly subjected to feasibility constraints in which case it shifts to lie on a boundary face. The SSK calculation employs multiple different centring algorithms, specially designed to counteract this tendency. Three of these are used in the early stages because they are applicable to unbounded polytopes, but allow different trade-offs between efficiency and scaling with the dimension count.

Once all capping constraints have been applied to produce a bounded SSK, new options appear. The final procedure arrived at from extensive trials, uses a Chebyshev centre, namely the centre of the maximal inscribed hypersphere, as a starting point. Calculating this is done by formulating it as minimax LP problem. While readily soluble, this centre point is usually not unique and the LP solution tends to place it far closer to some faces than to others. So, an iterative refinement is carried out in which a sample of directions in flux space are chosen. For each direction, the pair of offcut radii to the boundary intercepts, both forwards and backwards from the current centre, are found. The goal of the refinement is to reduce the difference between these pair radii, summed over all sample directions, to a minimum. To achieve that, the next iteration of the centre is chosen as the centroid of all the intercepts of all sample directions. Iteration is continued until satisfactory convergence is achieved. The final centre point is at the centroid of the final set of intercepts, which are the periphery points shown for example in Figure 4, Figure 5 and used in the resampling analysis discussed below. Figure 5 also shows an example of how the refinement shifts the centre point away from the inscribed circle centre.

For the refinement to succeed, the sample directions need to be carefully chosen. In the SSKernel implementation, 3 distinct sets are combined. The first of these are the endpoints of the main chords of the polytope. This set represents the orthogonal directions in which the range of variation of feasible fluxes are maximised, and includes at least one SSK vertex in each pair. The second is the directions of the orthogonal projection from the centre onto each polytope face. These directions take into account the actual shape of the polytope, and helps to place a periphery point on each face. The third set is intended to ensure that all directions are covered as evenly as possible. For an SSK in a *n*-dimensional space, the chosen directions are the (*n*+1) directions of the vertices of a regular simplex centred on the current origin. By construction, regular simplex vertices have equal spacing in angular space, i.e. the dihedral angle between all vertex pairs are the same. The simplex as a whole is randomly oriented in each iteration of the centre refinement cycle. Hence a large variety of equally spaced directions are sampled over the course of the iterations. It is plausible that the periphery points set, composed of a pair of points for each direction of each of the 3 sets, give a reasonable account of the flux variation allowed by the SSK constraints.

The centre point calculated in this fashion is a typical flux value, in a similar sense as the median of a 1-dimensional statistical distribution is regarded as typical. The centre point is as close to a median simultaneously along all directions, as is allowed by the geometrical shape of the SSK. The iterative refinement amounts to a sequence of trials of centre points and their accompanying peripherals, pursued until no further improvement can be obtained, so can reasonably be regarded as the best available single representative of the SSK flux values.

For metabolic fluxes there is another reason to consider the centre point as representative. The SSK polytope boundaries are ultimately determined by capacity constraints, nutrient availability and the optimised value of an objective such as biomass growth rate. Slight variations in the actual values for these constraints are to be expected between individual cells due e.g. to differences in genetics and environment. Hence somewhat different polytopes, defined by different boundaries, would apply to different cells, and perhaps even to the same cell over the course of time. The single point most likely to survive boundary changes, is the interior point located as far as possible from the boundaries, i.e. the calculated centre. From another perspective, if the SSK polytopes of all cells in a colony are superimposed, the centre point is the one most likely to be feasible in all of them. Similarly for a single cell in an evolving environment.

This argument obviously also applies more broadly to a central region near to the centre point. The PPP encloses such a region. It is by construction a polytope which has the calculated centre as the centroid of all its vertices. Moreover, it is composed of the endpoints of a set of diameters; on each diameter, the endpoints are located at the extreme feasible flux values along one direction, but central regarding all other orthogonal directions.

Another reason to consider the PPP as representative, is that it is found to cover a large part of the SSK hypervolume. This is visually clear in the scenarios presented; in Figure 4 it is in fact 100%, but in Figure 3 and Figure 5 it is closer to 90%. For a high dimensional SSK the coverage is not easy to estimate, but a rough indication would be to compare the radii of the hyperspheres that respectively circumscribe the SSK and the PPP. This idea is further pursued in [9] and leads to a more refined estimate that is routinely calculated by the SSKernel software. The ratio of radii for the red blood cell genome scale model discussed in the next section is 81.5%, and anecdotally values between 60% and 98% were obtained for a variety of genome scale models.

### 3.2 Exploring and Manipulating the SS

Knowledge of the SSK provides insight into the location, extent and shape of the solution space and can plausibly be exploited to direct cellular metabolism along desired channels. This section introduces some new computational tools for that and demonstrates their use.

Although realistic models are not amenable to visualisation, the SSK periphery point calculation yields a representative sample of feasible metabolic states that similarly serves to display the range of variation allowed by the model.

The key idea pursued here is to resample a particular subregion of the SS that corresponds to a metabolic intervention of interest, such as manipulating nutrient availability, gene regulation, etc. Inspection of the resulting new sample yields information about the cellular responses predicted by the model.

Resampling a section of the SSK could be done by mere interpolation of the SSK periphery point set, but the *HyperSampler* function provided in the SSKernel software goes beyond that by also using the ray basis to extrapolate sample points located in SS regions outside the SSK. Execution of the combined interpolation and extrapolation is not entirely straightforward, and details of the algorithm used are provided in the Supplementary Information [See Additional File 1]. Note that this algorithm also explicitly calculates two additional sample points, namely the points where a target flux supplied to it, reaches its minimum/maximum value within the resampled section of the SS.

Given a target metabolite or flux on which some intended bioengineering effort focusses, a further function, *TargetVariation*, tabulates the mean value and range of variation of the target, observed in the new sample, for each of a chosen list of interventions such as gene knockouts. Examples are shown and interpreted below.

#### 3.2.1 Toy model with *cof* as a target metabolite

Consider the toy model modified in scenario 5 above, and for illustration, suppose that we wish to influence the production of the metabolite *cof* (see Figure 1) via an external intervention. Two tweaks of the model are needed. First, in the model as stated, the stoichiometry guarantees net zero production of *cof* (as appropriate for its intended role as a cofactor). Increasing the coefficient of the metabolite for reactions 3 and 4 from 1.0 to 2.0 results in excess production. Second, a sink reaction to absorb this excess is added to the model, and its flux is defined as the target.

The effects on the target flux of various interventions, which were chosen to reflect typical cases and include some single gene knockouts, are shown in Table 2 and illustrated in Figure 7

**Table 2.**
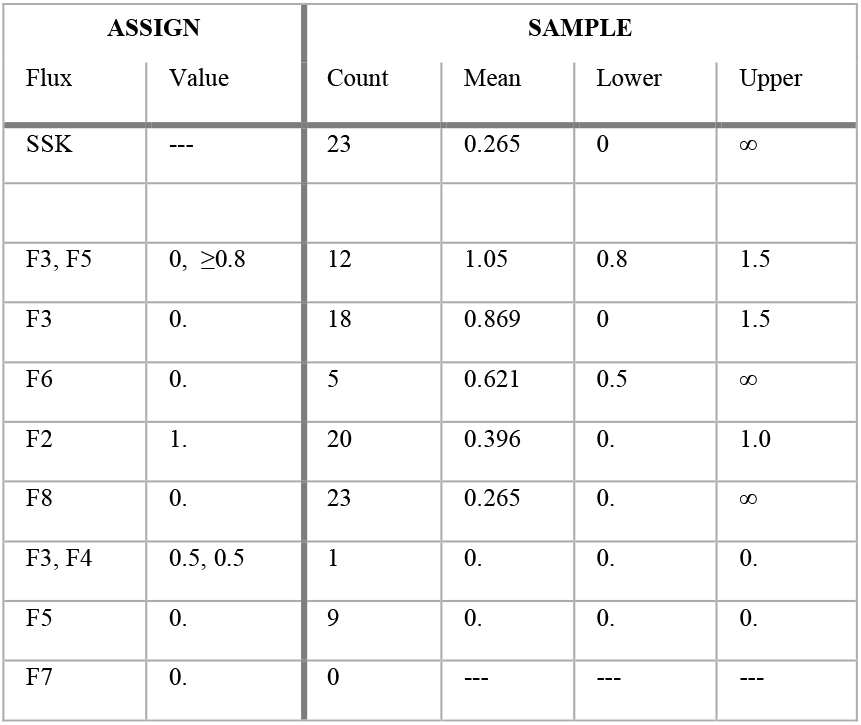
Target flux variation in response to flux assignments.

| ASSIGN |  | SAMPLE |  |  |  |
| --- | --- | --- | --- | --- | --- |
| Flux | Value | Count | Mean | Lower | Upper |
| SSK | --- | 23 | 0.265 | 0 | ∞ |
| F3, F5 | 0, $\geq 0.8$ | 12 | 1.05 | 0.8 | 1.5 |
| F3 | 0. | 18 | 0.869 | 0 | 1.5 |
| F6 | 0. | 5 | 0.621 | 0.5 | ∞ |
| F2 | 1. | 20 | 0.396 | 0. | 1.0 |
| F8 | 0. | 23 | 0.265 | 0. | ∞ |
| F3, F4 | 0.5, 0.5 | 1 | 0. | 0. | 0. |
| F5 | 0. | 9 | 0. | 0. | 0. |
| F7 | 0. | 0 | --- | --- | --- |

**Figure 7:**
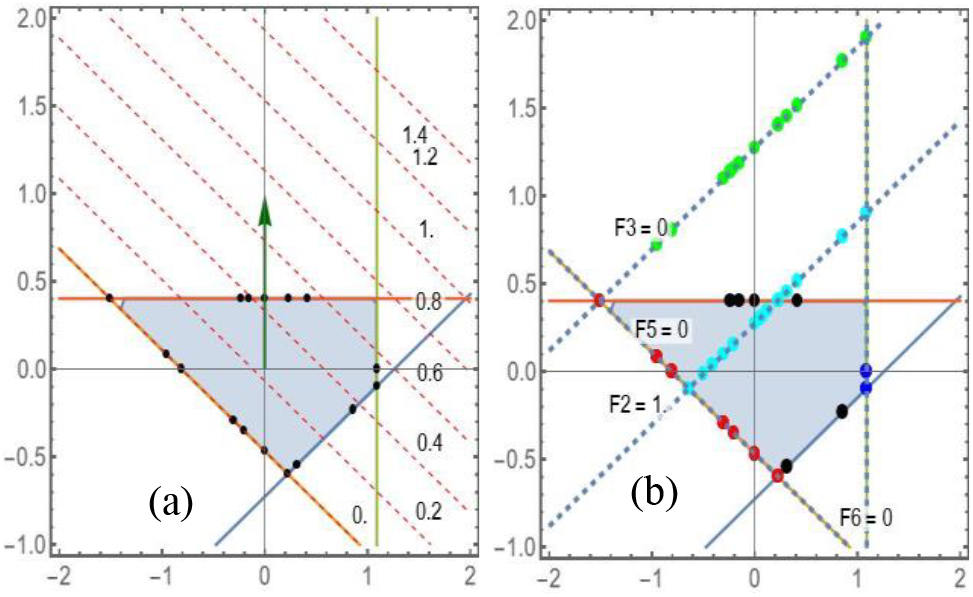
**a)** SSK with sampled periphery points shown as black dots, and contours of fixed target flux values as dashed lines. b) Fixed flux lines (dotted) and resampled points for some flux assignments: F3 = 0. (green), F2 = 1. (cyan), F5 = 0. (red) and F6 = 0 (blue).

For comparison, the first line in the table shows values for the unperturbed SSK.

Starting at the bottom of the table, blocking the input flux F7 makes the model infeasible, as indicated by 0 sample points. Blocking F5 allows a range of feasible points, but the target flux remains fixed to zero. Fixing F3 and F4 to 0.5 units each restricts feasibility to a single flux point still with a zero target flux value. In contrast, blocking F8 has no discernible effect: the sample count and target flux values remain the same as those for the SSK. Fixing F2 to 1.0 produces almost as many sample points, but now, the maximum target flux is reduced to a finite value while the mean value is increased. Blocking either F6 or F3 respectively restricts the lower and upper target flux limits, increasing the mean value in both cases. The strongest boosting effect is observed by simultaneously blocking F3 and restraining F5 to a range of values above 0.8 units.

All of these behaviours are easily interpreted by graphically displaying the SS and various flux contours, as shown in Figure 7. The SSK is the same as that in Figure 5 and is only reoriented for better display, and the unbounded SS now stretches to infinity along the vertical direction. The actual sample points generated are shown as coloured dots for a few of the single fixed flux cases included in the table. Note how these extend beyond the SSK where appropriate and cover the full range of variation fairly uniformly.

The case of assigning a value range is not explicitly shown, but for example, resampling F2 ≤ 1 produces a sample corresponding to the cyan, blue and some of the red points, in addition to some points on the lower right SSK boundary. The points represent all finite perturbed SS boundaries but may also include a few interior points inherited from the SSK sample.

Comparing a fixed flux line in Figure 7b with the target flux contours in Figure 7a, the value range shown in Table 2 for the corresponding case is easily reconciled. It also indicates how the sample set often has fewer members than the original periphery points, due to the smaller dimensionality or extent of the subsection, and sample points consequently coinciding. Such graphic interpretation is not possible for realistic models, but the table still conveys the relevant information.

#### 3.2.2 Resampling a realistic model

The red blood cell (RBC) metabolic model [13] included as a demonstration example with the SSKernel software is used to illustrate that resampling is also viable for realistic models. This model has 469 fluxes, of which 408 are found to have the same value for all feasible fluxes (i.e., they are fixed fluxes), leaving 61 fluxes that vary across the SS but are restricted to an 8-dimensional bounded SSK with 4 ray directions.

By way of comparison, an FVA calculation of this model identifies the same 408 fluxes as fixed values, but it takes 132.5 seconds of computing time compared to the 1.33 seconds for the entire SSK calculation, of which fixed flux determination is only a small part. The SSK calculation is not only around 100 times faster, but also more informative than FVA as previously discussed.

A further example of that, is that in this case the FVA result identifies 9 reaction fluxes that remain unbounded, while SSK shows that there are only 4 independent ray vectors. Figure 5 shows how such a discrepancy can arise: the single ray vector points in a direction in flux space that combines several flux axes, all of which tend to infinity if the flux increases along the ray direction. From the biochemical perspective, the single ray in this case represents a loop in the network, and involves several reactions. The presence of unbounded fluxes in the FVA shows that the SS is unbounded, but gives no information on the actual independent rays belonging to the SS. We also note that anecdotally, the SSK calculation scales better with flux count than the FVA, so for large models, even larger runtime discrepancies apply than the factor 100 found in this example.

Before discussing target metabolites, we note that simply inspecting points sampled from the SS can reveal aspects of metabolism. For example, the flux for the water transport sink reaction EX_h2O_e in the RBC model varies between −6.3 and 4.8 units, allowing both the import and export of water. What would happen if either of these is ruled out or if no net water transport is allowed?

This is investigated by resampling from the subregions where for that particular reaction, either F ≤ 0, F ≥ 0 or taking the intersection of hyperplane F = 0 with the SS. In the latter case, for example, all changes in the remaining mean fluxes from their values over the SSK are plotted in Figure 8. There is a large contrast in the observed responses. For reference, the seven reactions where the magnitude of the change exceeded 0.1 are labelled EX_ade_e, EX_adn_e, EX_bilglcur_e, EX_fum_e, EX_h2o2_e, EX_hco3_e and EX_man_e. The biochemical significance of this observation, if any, could perhaps be further explored by sampling from SS subregions that focus attention on subsets of these particular fluxes.

**Figure 8:**
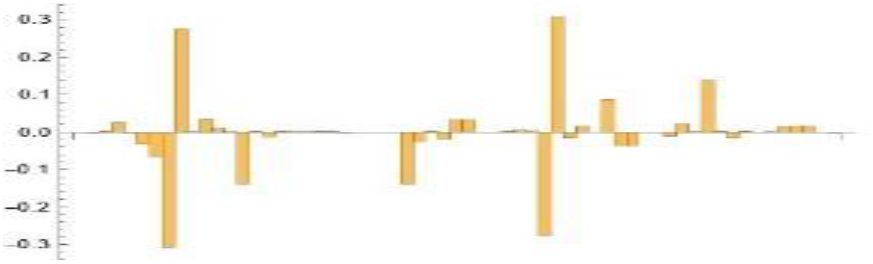
Mean value changes from SSK values for the variable fluxes, in response to requiring net zero water exchange in RBC model.

#### 3.2.3 Maximising RBC oxygen production

To show an example of how choosing a target metabolite could suggest a bioengineering target, consider that the function of the red blood cell is to transport oxygen. Is it possible to enhance this function by knocking out any single gene? To address that question, we identify the oxygen export reaction EX_O2_e in the existing model as the target flux. The broader SSK analysis discussed above has already narrowed the field from 469 fluxes to 61 variable fluxes, for which variability is compatible with the RBC model constraints.

The *TargetVariation* function illustrated above for the toy model provides a table of predicted outcomes for a systematic trial of each of the 61 knockouts. An extract of the results is shown in Table 3.

**Table 3.**
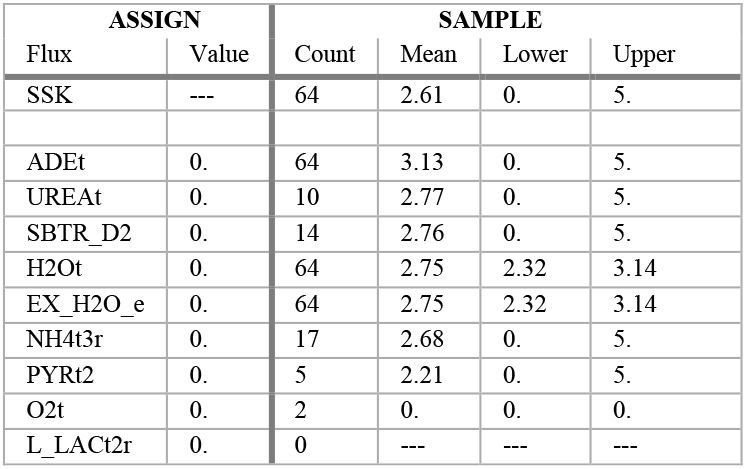
Red blood cell oxygen export flux for selected singlegene knockouts.

There are again some cases in which an infeasible model is produced, or the target flux is restricted to zero. Most knockouts leave the range of the target flux unchanged while either increasing or reducing the mean target flux relative to the SSK value. Two cases stand out, however: knocking out either the H2Ot or EX-H2o_e reaction restricts the target EX_O2 flux to a fairly narrow, positive value range. Although there are other knockouts, such as ADEt, that produce a higher average value, this is arguably a more potent intervention since it guarantees a substantial oxygen export flux.

A close link between oxygen export and water export may come as no surprise to some readers, in which case the example confirms that the resampling procedure yields reliable results. Otherwise, it is a demonstration that interrogation of the SSK using resampling tools can be useful for identifying new bioengineering interventions for further study.

For this realistic but still rather small model, it is practical to use a brute force approach of testing all available gene knockouts since only 8.5 s of computing time was required to trial all 61 cases. For larger and more complex models, this may well become intractable. However, since the method merely requires a list of candidate interventions as input, the application of biochemical knowledge to set up such a list should be able to reduce the task to manageable proportions even for large models and for simultaneous knockouts of several genes..

The utility of the resampling approach can be further assessed by comparing it with using FVA to investigate the same questions.

For the case of section 3.2.2, FVA could only show a changed flux range, not the change in mean flux displayed in Figure 8. Table 3 shows that a different size and shape of the SS can often produce a different mean value while leaving the flux range intact. Not only does the resampling give this more nuanced information, but it runs in less than 2 seconds, including the initial SSK calculation. Two FVA runs, one for the base model and one for the perturbed model are required, and would need around 250 seconds to run.

The case of section 3.2.3 seems more favourable for FVA since the main inference was based on flux ranges. However, it would offer no accuracy advantage; it gives identical values to those tabulated in Table 3 since the resampling also finds these extremes by explicit optimization rather than interpolation. But each knockout, of which there are 61 in this case, would require a separate FVA calculation because each has a different set of model constraints. Hence the computational effort would run into hours rather than seconds, and would still miss the more subtle differences in mean values shown in the table.

## 4 Conclusions

An understanding of the range of metabolic states available to the metabolism of a cell, as delineated by available FBA metabolic models, is a powerful tool to design bioengineering interventions that produce desired metabolic outcomes. Equally such understanding can contribute to explain observed effects of the cellular environment, such as nutrient availability, on the metabolism.

Computing the solution space kernel (SSK) facilitates such understanding. Careful approximation reduces the vast complexity induced by the millions of extreme pathways needed to give an exact description of the solution space of genome-scale metabolic models. At the same time, it is much faster and far more informative than the reduction to a cuboid bounding box produced by flux variability analysis (FVA).

From a solution space dimension count of thousands, the SSK reduction focusses attention on a kernel polytope with a dimension count usually well below 100 and characterized by a correspondingly small set of constraint equations. It allows geometrical perception of this kernel by explicit determination of its centre and an orthogonal set of maximal length chords. At an even more practical level, a semi-random but empirically representative sample of feasible fluxes are produced, that can be used for statistical analysis or as the basis for experimenting with the effects of imposing chosen genetic regulation or other manipulations of the metabolism.

A focus of the work is to allow description of the SSK as a geometric object in flux space, with a location, size and shape. Two straightforward explicit examples of how this view can yield practical information useful for bioengineering were shown. We believe, however, that this only scratches the surface of what might be possible. For example, it seems plausible that the shape and size of the SSK might reflect the robustness of the metabolism of a cell to external challenges, since it delineates which adaptations are feasible given a set of constraints such as nutrient availability. The SSKernel software opens the door for such conjectures to be further studied.

## Supporting information

Resampling algorithm

SSK software manual and code

Input data and workflow

## Abbreviations

DIF: Digital Interchange Format
FBA: Flux Balance Analysis
FBF: Feasible Bounded Face
FVA: Flux Variability Analysis
LP: Linear Programming
OS: Objective Space
PPP: Periphery Point Polytope
SBML: Systems Biology Markup Language
SS: Solution Space
SSK: Solution Space Kernel

## Availability and Requirements

**Project name:** SolutionSpaceKernel

**Project home page:** https://github.com/wynand-verwoerd

**Current Release:** doi:10.5281/zenodo.11483838

**Operating system(s):** Platform independent, provided Mathematica implementation available.

**Programming language:** <u>Wolfram Language</u>

**Other requirements:** Wolfram Mathematica or Wolfram Engine

**License:** GNU GPL 3.

**Any restrictions to use by non-academics:** None

### Computing environment

Full utilization of the SSKernel software including its interactive control interface, requires installation of the proprietary *Mathematica* system. Work is in progress to produce a stand-alone version of the software and make this available in due course as part of the GitHub release.

However, for reproducing the results in Section 3, a command line implementation is perfectly adequate and for that only the installation of the <u>Wolfram Engine</u> is required. This can be downloaded for free, though subject to use restrictions that do allow testing applications. It is also available for online use via a Wolfram cloud server.

## Declarations

### Ethics approval and consent to participate

Not Applicable

### Consent for publication

Not Applicable

### Availability of data and materials

Input data sets are referenced and explicitly listed in the Supplementary Information.

### Authors’ contributions

WSV: Conceptualization, Methodology, Formal Analysis, Software, Writing – original draft preparation.

LM: Conceptualization, Validation and software testing, Writing – review and editing.

## Acknowledgements

Not Applicable

## Supplementary Information

The online version contains supplementary material in the form of three additional files, as follows:

- Additional File 1.docx: The mathematical formulation and computational procedure to resample points in a subregion of the solutions space, for use in exploring metabolic interventions.
- Additional File 2.docx: Explicit input data sets for producing the various Toy Model scenarios described in Section 3 of the article.
- Additional File 3.zip: The compressed file directory SSKernel Implementation, which contains the current release of the SSKernel software. This includes a 55 page software manual, all source code and test data.

