## Supplementary material for "The FBA Solution Space Kernel – Introduction and Illustrative Examples": Resampling algorithm

### Resampling an FBA Solution Space

This work is an extension of the research monograph (Verwoerd, 2022) which describes the FBA solution space kernel (SSK) and its computation, and the terminology and formalism used here may require reference to that source.

For practical usage the example data presented in section 2 below and the workflow described in section 3 may suffice and the technical description that follows could be skipped.

#### 1.1  Formulating the problem

The metabolic network model is taken to contain *M* metabolites and *N* reaction fluxes, linked by a (*M* x *N*) stoichiometry matrix. Resampling comes into play once a full solution space kernel (SSK) calculation has been performed on this model, which generally includes the optimization of an FBA objective. The following numerical arrays produced by the SSKernel software need to be read from its XXXX.dif output file as input to the resampling calculation:

1. The reduced solution space (RSS) specification consisting of a constraint set and a vector transformation from the full flux space (FFS), as described in (Verwoerd, 2022) Eq. 1.12.
2. The SSK periphery point matrix *P*. For π points, this is a (π x *N*) matrix with each row containing the position vector of a periphery point in FFS.
3. The (ρ x N) matrix *R* containing each of the ρ ray vectors in FFS as it rows.

In addition, the user is expected to define the resampling goal by supplying:

4. A constraint or set of constraints that defines the hyperplane H or subregion to be sampled. This consists firstly of a matrix *C*_H_ with λ rows and *N* columns. Each row specifies a direction in FFS that defines a perpendicular hyperplane which partitions the FFS into half-space regions. Secondly, the corresponding values vector *V*_H_ with λ rows; each specifies the perpendicular distance value and an index that signifies whether it is an equality or inequality.
5. The target flux column *t* for which the range of its values over H are to be monitored.

The user supplied data described in point 4 above provides for consideration of two distinct cases. The simplest is where all constraints defining H are strict equalities. Then sampling is restricted to the hyperplane H itself and this forms the point of departure for the formalism introduced below.

However, if some of the constraints are inequalities, sampling is done over a subregion of the SS which is bordered by hyperplanes contained in the H specification. Each individual inequality defines a half-space, so defines part of the boundary. The entire composite boundary is obtained by changing one inequality at a time into an equality. This subregion sampling requires some generalizations of the simple H sampling procedure, as indicated where applicable below.

The overall concept of the algorithm is that new sample points are found by using linear programming (LP) to find a feasible point that is close to each periphery point, but located on H.

A key feature is that the actual calculations are performed in RSS rather than FFS. This exploits the fact that the RSS constraints matrix *C*_RSS_ has *m* constraints (rows) on *n* variables (columns) where generally *m* << *M* and *n* << *N*. Some formal manipulations are required to achieve this computational simplification and are detailed in what follows.

The feasibility requirement means that in fact it is the intersection of H with the SS that is to be resampled. The simplest case would be if H intersects the SS only within its bounded kernel. In this case just interpolating between the known periphery points would be sufficient and guaranteed to generate feasible points without needing to invoke the SS or SSK constraints explicitly.

However, to cover the general case that parts or all of the intersection lies outside the SSK, extrapolation using rays is required. This is based on the defining property of the SSK that any point in the SS can be reached from some point in the SSK by adding a multiple of a ray vector. The discrete set *P* does not guarantee that; so it is necessary to start the extrapolation from an unknown interpolated point. In other words, a combined interpolation and extrapolation procedure needs to be constructed.

One limitation must be recognized. Using the predetermined periphery points limits the interpolation/extrapolation to points in the SS that are reachable from their convex hull (the periphery point polytope or PPP). The underlying assumption is that the PPP is representative of the entire SSK, even though it is only a subset constructed to cover a major central region of the SSK. Then it should be enough to approximate the mean or median value of a flux from PPP-generated samples only. However, it is conceivable that the outlying SS regions not included in the sample may modify these values.

In the extreme, it is in principle possible that H may only intersect the SS in such outlying regions and then no sample points would be found, and this could misleadingly suggest that H is infeasible. That is avoided in what follows by performing an independent test for infeasibilty of H and raising the alarm if a sampling failure turns out to be a false negative. The only remedy would be the computationally demanding process to generate each sample point from scratch using a large LP that combines *C*^H^ with the model constraints, but as current experience has not encountered such a case it is not considered further here.

The range of the target flux is the aspect that is most susceptible to be misrepresented by limiting sample points to those that can be reached from the PPP. As a plausible goal of bioengineering is to restrict this range, it is worthwhile to extend the sampling so that it includes two points that respectively minimize and maximize target fluxes. That eliminates (at least regarding flux range) the concern about using the PPP rather than the full SSK and is incorporated in the discussion below.

Points in *P* are by construction located on the periphery of the SSK. However, when projected by the interpolation/extrapolation procedure to H, some may fall in the interior of the convex hull of resampled points. Such a point is in principle redundant, since it could be formed by a linear combination of convex hull points.

How such interior points are treated differs between the case where H is defined by pure equality constraints versus the case where there are one or more inequality constraints. In the first case H is a simple hyperplane and interior points are kept because they still contribute to giving representative target flux values.

The second case implies that the resampling is taken from a subregion of the SS which is bordered by the hyperplanes in which *C*_H_ constraints are changed to equalities. Locating sample points in the interior of this subregion would leave its new borders unrepresented. That is avoided by explicitly resampling the bordering hyperplanes, obtained by taking each inequality in turn as an equality.

For simple hyperplanes multiple points in *P* may project to the same point on H, so that the size of the new sample is often smaller than the original. However, the iterative nature of the procedure for subregions means that a single point in *P* may be projected to several borders and the new sample can be larger than the original. It is not unreasonable that a larger sample is required to be representative of a subregion with extra borders not present in the SSK, but it indicates that resampling a complicated subregion can become computationally demanding.

For a bounded H-SS intersection, an adequate representation of feasible fluxes is given by the set of resampled points. However, in many cases the intersection remains unbounded, and then it is in addition necessary to find a ray basis for the intersection. In fact, determining its ray basis to be empty, is the most straightforward way to establish that the intersection is bounded.

Merely projecting the SS ray set to the intersection is not enough, as a ray projection may not be a ray in the intersection anymore. Consider, for example, the case where the SS is unbounded but the H-SS intersection is bounded; then clearly none of the projections of SS rays remain rays in the intersection. This problem of finding intersection rays is addressed in section 1.4 below.

Two subsidiary problems raised above are whether the H-SS intersection is in fact empty, so establishing beyond doubt the feasibility status of H; and how to calculate the target flux minimum and maximum needed for its exact range specification. Each of these are addressed in a separate section below, while collection of relevant statistics is outlined in the last section.

#### 1.2 Constraints for Interpolation and Extrapolation

In the simplest case a new sample point S constructed by just interpolating the point matrix *P* and is represented by

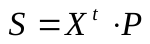

where *X* is a column vector of coefficients to be determined.

To ensure feasibility of an interpolated S, it is sufficient to impose the convex conditions
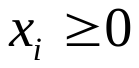
 and
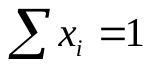
 on the components of *X*.

Extrapolation, on the other hand, starting from a specific point *p*_i_ at row *i* in *P* and using the ray basis is given by the row vector

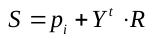

Here, the column vector *Y* contains the coefficients for superposition of the rays in matrix *R* and needs to be subjected to a constraint to ensure that the linear combination remains a ray. This constraint is the ray condition (Verwoerd, 2022) Eq. 4.1 and for a column vector *x* is given by
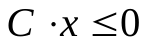
Here *C* is the (*M* x *N*) constraints matrix of the original model including stoichiometry, objective and range constraints.. Applying the ray condition to the ray superposition term in Eq. gives

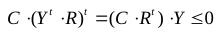

This condition on the allowable coefficients *Y* only involves the (*M* x ρ) matrix
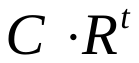
. This only needs to be computed once and applied repeatedly to all *i*, and moreover as shown next can be evaluated in the reduced solution space RSS rather than FFS, reducing matrix sizes by 2 or more orders of magnitude.

The RSS specification as read from the SSKernel output file is given as the compound list

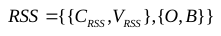

consisting of the RSS constraints in the first sublist and the position vector *O* of the RSS origin and its basis vectors *B* as the second sublist.

The nature of the RSS reduction is such that most of the constraint vectors in the FFS matrix *C*, are orthogonal to the RSS. This stems from the fact that RSS is located in the null space of the stoichiometry matrix, as well as from the elimination of fixed fluxes, linealities and prismatic rays. Express this fact by explicitly partitioning the rows of *C* into the subset *C*_r_ that have a non-zero component in the RSS, and the remainder *C*_e_ that are orthogonal to the RSS and hence project to zero and are eliminated during the reduction. The rows of *C* can without loss of generality be sorted so those that belong to *C*_r_ appear first followed by *C*_e_. Similarly, the rows of *R* can be partitioned and sorted so that all conical rays (that remain after projection to RSS) appear first, as the submatrix *R*_r_, followed by the submatrix of eliminated rays *R*_e_, namely prismatic rays and linealities. Using the partitioning, the FFS matrix product in Eq. can be expanded as

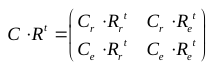

Both off-diagonal elements in the expansion reduce to zero. That is obvious for the lower left entry
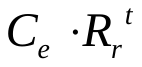
 since by definition their vectors belong to orthogonal spaces. Regarding the upper right entry
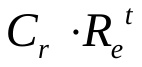
, the constraint vectors in *C*_r_ overlap with, but do not necessarily belong to, the RSS so it is less obvious that this entry reduces to zero as well. First, we note that for linealities, they by definition satisfy
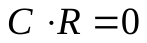
 i.e. are orthogonal to all constraint vectors and hence also to the subset *C*_r_. The second group of rays that make up *R*_e_ are prismatic rays, and by definition they are orthogonal to all rays except one, with which they have an overlap of -1. That particular constraint vector is eliminated by coincidence capping of the prismatic ray to produce the RSS, and so is excluded from the subset *C*_r_. It follows that also
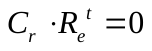

The submatrix
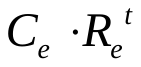
 that appears at the lower right in Eq. also simplifies substantially. Again, any column in the submatrix that derives from a lineality row in *R*_e_ will reduce to zero. The prismatic ray cases each overlaps with exactly one unique row of *C*_e_ and produces the entry value -1. So
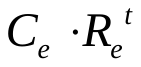
 contains mostly zero rows, except for a single negative unit vector corresponding to each prismatic ray. The zero rows identically satisfy Eq. for any *Y* and can be omitted without loss of generality. Also omitting lineality columns, the remaining columns of
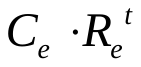
can be sorted so that it becomes a negative identity matrix.

Finally, consider the upper left submatrix
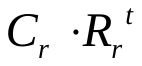
 in Eq. . When combined with Eq. this submatrix ensures that a vector formed by linear superposition of rays belonging to *R*_r_ remains a ray in the full solution space. In principle, *R*_r_ contains all rays that are not orthogonal to the RSS but they do not have to be confined to this subspace. In practice, though, the SSKernel software performs the calculation of conical rays only after completing reduction the RSS. Since the rows in *R*_r_ originate from the ray matrix obtained from the SSKernel output file, they will thus in fact belong to the RSS. Therefore they remain unaffected by matrix projection to the RSS, using the orthonormal RSS basis vectors specified by matrix *B* from Eq. . Formally, that means that

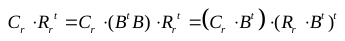

However, the matrix defining the constraints in the RSS is obtained by downcasting all constraints in C that survive the RSS reduction to vectors in RSS, i.e
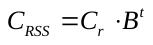
. The expression
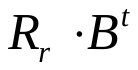
 represents the rays expressed as vectors in RSS, so the implication is that the FFS matrix product on the left in Eq. may be evaluated as an equivalent product in the much smaller reduced space. Bearing in mind that the ray matrix delivered in the SSKernel output file is specified in full flux space, it is however computationally convenient to keep the explicit form for the second factor and write

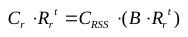

A point glossed over so far is how to practically perform the partitioning introduced conceptually in Eq. . For the constraint matrix this becomes irrelevant because only *C*^RSS^ is used. For the rays, it is simple to calculate
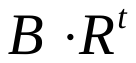
. All zero rows in this product identify the excluded rays that constitute *R*_e_ ; deleting those, what remains is the bracketed term in Eq. .

Having ensured that S is feasible, the next step is to ensure that it falls in the subregion or on the hyperplane H. This requires that

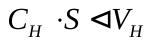

where *C*_H_ and *V*_H_ are user supplied as detailed in point (4) in section 1.1. The ◁ sign used in Eq. is meant to represent ≤ , = or ≥ as the case may be for each line individually, and as specified as the second part of the element in *V*_H_.

All the constraints may be collected into a single matrix equation formulated in the space formed by concatenating the vectors *X* and *Y* , but with the Y vector components arranged into its subsets associated with retained (conical) rays and eliminated rays respectively :

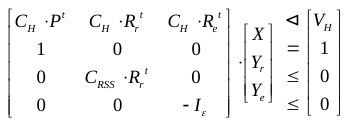

Here, *I*_ε ­_signifies the ε-dimensional unit matrix and 0 and 1 the appropriately dimensioned matrices of 0’s and 1’s respectively.

In some metabolic models, the rays are such that either the subset *R*_r_ or *R*_e_ is empty. The constraint equation accordingly simplifies in each case. If the RSS is bounded, there are no retained rays and then

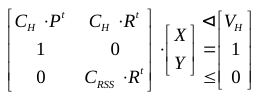

On the other hand, when no rays were eliminated to form the RSS we have

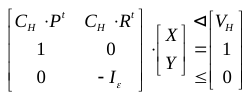

#### 1.3 The optimization procedure

We next address the procedure to find X and Y for each resampled point S. For clarity, consider first the case of a simple hyperplane H, while the subregion case will be discussed later.

As set out in section 1.1, a new sample point on H is generated by combined interpolation and extrapolation from the known sample *P* of points located in the solution space. Since H and the SS intersect, some members of *P* may already belong to H and they are merely transferred to the new sample set. The complementary subset is referred to as the focus point set. The strategy is to focus on one of its members at a time, and resample a new point located as closely as possible to the focus point but located in the H-SS intersection. The rationale for keeping the resampled point close, is to preserve as far as possible the fairly uniform spatial distribution of points that was built into the SSKernel peripheral point sampling procedure by which *P* was generated.

So the main calculation is a loop over all focus points, in which the currently selected one is identified by an index *i*. An obvious way to resample a close point is to maximize the interpolation coefficient *x*_i_ while minimizing the contribution from ray vectors. That could be accomplished by a linear programming (LP) problem, subject to the constraint set or its simplifications, and with the (π+ρ)-dimensional objective vector

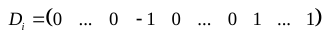

where the vector product
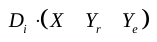
 is minimized. The range of *x*-variables is restricted to non-negative values, while *y*-variables are unrestricted.

The right hand side of is meant to convey that the *i*-th component is set to -1 to induce maximization of *x*_i_, while the last ρ entries have the value 1 implying that
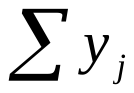
 is minimized. This will in fact only guarantee a minimal ray contribution if all *y*_j_ are positive as is the case for a convex ray basis. The ray basis as actually constructed by SSKernel does not have this property; it is merely vectorially complete so will allow negative coefficients. That could be provided for in the LP problem by introducing a set of auxiliary variables that instead minimizes
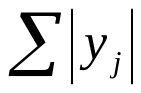
, albeit at the expense of an increased vector space dimension.

However, even with such elaboration this approach is not successful. Due to the minimizing of the ray contribution, it becomes skewed towards always using interpolation when possible, rather than extrapolation. That is OK if the H-SS intersection is either completely inside or outside the SSK; in the first case interpolation is used exclusively, and in the second, since interpolation does not work, extrapolation is used. However, in the case where the H intersection is only partially in the SSK, no sample points are ever found in the part of the intersection outside the SSK as this would require extrapolation using rays. Hence no matter which focus point is used, it always ends up with an interpolated sample point inside the SSK. Failure to also resample the SS outside the SSK misses an important goal as it undermines the representative aspect of the sample.

One remedy would be to drop the requirement to minimize the ray contribution. The only optimization included in the objective, is to maximize the contribution of one particular periphery point chosen as focus. However, entirely removing the penalty for using rays introduces a bias towards extrapolation. This can result in an extrapolated resample point located far away from the focus point, even when a nearby point could have been reached by interpolation.

This is because the focus point coefficient for extrapolation always has the value 1, while for interpolation it is typically between 0.6 and 0.8. Moreover, when there are multiple rays, any of them that produces an extrapolation point on H are completely equivalent since all give the focus point coefficient 1. The ray-related part of the LP problem is then identical for all focus points, so the same ray is usually selected for all of them (typically just the first on the arbitrarily ordered list of rays). This means that the extrapolated points are not uniformly distributed over H; it is more like the set of focus points are just projected onto a limited patch of H by a common translation along the arbitrarily selected ray direction.

The final strategy chosen to counteract both biases, is to modify the objective vector by installing a set of small, randomly selected positive weights *ω*_i j_ for the ray coefficients, and varying the random choice for each focus point:

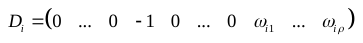

The *ω*_i j_ are scaled such that
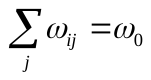
 and *ω*_0_ is an adjustable parameter currently chosen as *ω*_0_ = 0.2. The positive *ω*_i j_ values means a penalty for using rays. Bearing in mind that the convex condition guarantees that the positive coefficients of the interpolated contribution to S add up to a value of 1, the chosen scaling value is meant to be small enough to equally balance tendencies to favour interpolation and extrapolation. Otherwise, it is an arbitrary choice, but appears to work well enough in low-dimensional trials to produce resample points that are well spaced on the target hyperplane H.

In compiling the resampled point set, the subset of *P* that satisfy the *C*_H_ constraints is used as a starting set and the optimized point derived from each focus point is added to the set as the resampling progresses.

As described so far, the resampling consists of solving an LP problem for each focus point, in which the constraint set is the same for all points but the objective vector is constructed individually for each point and signified by the index *i* in Equation . As for Equation , the objective vector gives rise to an objective value that is minimized.

This single loop over the focus point set is applicable to cases where H constitutes a simple hyperplane, i.e. the relations symbolized by ◁ in Equation all reduce to equalities. For a subregion, where some of the relations are inequalities, the strategy still needs elaboration to implement the sampling of individual boundary hyperplanes outlined in section 1.1

This elaboration consists firstly of embedding the point loop in an outer loop that extends over all inequalities contained in ◁ . In each iteration, one such inequality is temporarily replaced by an equality. This defines a new boundary hyperplane of the subregion that is introduced by H. Then the inner loop generates all new sample points, located on this boundary hyperplane, originating from the set of focus points. Clearly there could be quite a proliferation of resampled points involving a substantial increase of computational effort. Depending on the inherent size of the metabolic model, this can restrict the number of inequalities contained in ◁ that can be realistically computed.

A second aspect needing elaboration is how the focus point set is constituted. For a simple H, the rationale for excluding the subset of *P* that satisfy the H constraints from the focus group, is that such points already fall on the boundary and would trivially interpolate to themselves. For a subregion, however, many points in *P* may satisfy the H inequalities but are in the interior of the subregion and so can usefully extrapolate to a new point on the boundary. So instead of partitioning *P* into complementary subsets, the first step is to take all its members that satisfy the full set of H constraint inequalities as a starting collection of resampled points. However, the points that are excluded to form the focus point set, are limited to those that satisfy a stricter requirement, namely that that it in addition satisfies at least one inequality as a strict equality. This requirement places it on one of the hyperplanes that make up the composite boundary and so will again trivially interpolate to itself were it not excluded.

Note that while some points are excluded from the focus set over which the resampling loop iterates, all members of *P* are allowed to contribute to each interpolation, as signified by the use of the full *P* matrix in Equations , and .

#### 1.4 Finding the H hyperplane rays

An unbounded solution space (the usual case) generally has both bounded and unbounded boundary facets. The resampling hyperplane H may or may not intersect its unbounded facets in such a way as to truncate SS rays. Hence it is not a priori clear whether the H-SS intersection is bounded or not. Finding a non-empty ray basis for the intersection is a direct way to establish that.

Moreover, the resampled points are by construction finite, even if they happen to fall on an unbounded boundary facet. So collectively they will always indicate a finite value range for the target metabolite flux. To establish whether the target flux is in fact bounded in the H-SS intersection, it is necessary to find out if the target flux points along a ray direction.

For both of these reasons it is of interest to calculate the ray basis of the intersection. Considering first the case of a simple hyperplane H, an obvious start is to project all SS rays to H. To that end, since all constraints are equalities, H is in effect the null space of its constraints matrix CH and so formally its orthogonal basis is given by the rows of the *Mathematica* matrix construct
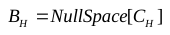
. Then the set of projected rays are defined by

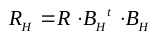

Knowing that some rays may project to the zero vector, multiple projections may be identical and some projections may even fail to remain rays, the first task is to establish that the projected rays still form a complete basis for rays in the H-SS intersection.. Let *r* be an arbitrary SS ray vector, then completeness of the SS ray basis *R* guarantees that there is a coefficient vector *X* such that
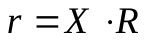
. The projection of this vector to H is given by

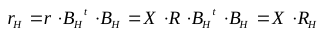

Since any ray in the intersection is also a ray in the SS, Equation applies also to this case where *r* and *r*_H_ are identical. Hence this shows that there exists a coefficient vector *X* that can be applied as coefficients to the projected ray set *R*_H_ to form such an intersection ray, namely the same set *X* that was used to form it as a superposition of the SS rays. It follows that the projected set forms a complete basis.

In the practical construction of the intersection ray basis, just as for the resampling points there may be some low hanging fruits in the form of SS rays that already satisfy the H ray condition
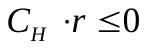
. These are directly used as a starting set of intersection rays.

The remaining SS rays are projected according to Equation . The resulting *R*_H_ matrix may contain zero rows stemming from rays orthogonal to H. Delete them and normalize the remaining rows; then delete any duplicate rows that occur. This leaves a matrix *Q* still with *N* columns, but fewer rows than *R* and each of those represent a candidate ray vector for the H-SS intersection.

Since projection can change the vector direction, there is no guarantee that a candidate will still satisfy the ray condition based on the SS constraint matrix *C* and this needs to be explicitly tested. This test in principle requires evaluating
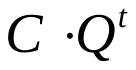
 but the arguments employed in section 1.2 applies again, except that since the eliminated rays in the *R*_e_ ray subset do not contribute to *Q* its is only the *C*_r_ submatrix that survives and evaluating this in the RSS as done in reaching Equation , we remain with the test

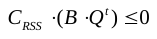

Any rows in *Q* that gives rise to a positive component value for the matrix on the left of this equation are rejected, and the rows that remain are combined with the starting set to constitute the H-SS intersection ray vector basis.

As before some elaboration is required in the case that H represents a subregion. In that case, only the strict equalities among the H constraints are included in the null space calculation of Equation to determine a vector basis for the projection, while the remaining inequality equations are appended to the RSS constraints to give an extended ray test condition in Equation .

Also, for a subregion it is desirable to include rays aligned to the newly added boundary hyperplanes, so as to maintain the criterion that rays are chosen to be as peripheral as possible, as set out in (Verwoerd, 2022) section 4.5. That is achieved by repeating the projection step during the subregion outer resampling loop, and where each inequality in turn becomes an equality and so gives a ray aligned with that boundary. Any additional rays found in this way are also added to the ray collection. This resulting ray basis may well be overcomplete, but that is of no concern as the original SS ray basis was by construction already potentially overcomplete.

Finally, note that it may happen that the starting set, the rays projected to H and those projected to boundary hyperplanes of H may all turn out to be empty. Since the projected ray basis has been shown to be complete, it means that in this case there does not exist any rays. In other words, this outcome proves that the H-SS intersection is bounded. That is an important characterization in its own right.

#### 1.5 Checking that H and SS do intersect

Failure to find any resampled points suggests that H does not intersect the SS, but as noted above could also result from inadequate representation of the intersection by the limited set of SSK periphery points that form the basis of the resampling. For a conclusive result an explicit test for a non-empty intersection is needed. In fact, performing the test first can avoid futile computation of resampled points in cases where no intersection exists. As it turns out, the test can be formulated as an LP problem in the reduced solution space or RSS and with a suitable choice of the objective will also indicate whether the intersection falls within the RSS.

Let an arbitrary point in the RSS be specified by an *n*-dimensional column vector *U*. To belong to the RSS it has to satisfy the constraints

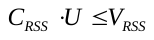

The constraints in *C*_H_ that ensures that it also belongs to H, are given in the full flux space (FFS) with *N* dimensions. To apply those the vector *U* needs to be uplifted to give *U*_SS_ according to

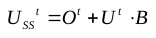

where *O* and *B* refer to Equation . But as the RSS was formed by coincidence capping of the eliminated rays in *R*_e_, an arbitrary point *X* in FFS is formed by adding a suitable ray combination with coefficients *Y*_e_ to *U*_SS_ giving

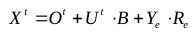

Substituting this into the H constraint set leads to

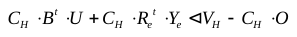

In addition there is a condition to ensure that the last term in equation represents a valid ray, and this simplifies when rearranging C as in Equation with the result that the ray condition merely requires that all components of *Y*_e_ must be nonnegative. Subject to that, Equations and can be combined into a single matrix equation by concatenating the vectors *U* and *Y*_e_ as before to give

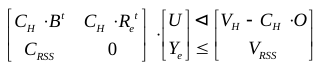

If the constraint set are incompatible, it means that the intersection of H and SS is empty.

To establish that, an LP problem needs to be formulated to see if there exists a feasible vector [*U* *Y*_e_], with the value range of *Y*_e_ restricted to positive values (required by the ray test) and *U* not restricted. An informative objective has unity coefficients for the ray contributions, i.e. the vector *o* = [0 ... 0 1...1] .

This means that the sum of *Y*_e_ values is minimized. Since *Y*_e_ ≥ 0, it will find a solution for *U* with *Y*_e_ = 0, unless no such solution exists. Since *Y*_e_ = 0 means there are no ray contributions, this is a solution within the RSS. Then H intersects the RSS directly. If the LP does find a solution, but with *Y*_e_ mildly positive, it means that H does intersect the SS, but outside of the RSS. In either case the new point found is added to the list of resampled points.

If no LP solution is found, it means H does not intersect the SS at all. In that case the resampling is aborted, and a suitable message delivered.

Note that this is a general test, not reliant on the periphery points and does not assume the PPP to be representative. So it can be used to test that assumption as made in the main calculation; if no sample points are found despite this test finding there is an intersection, the PPP is inadequate and an alarm is raised.

#### 1.6 Resampling points with extreme target flux

A final extension of the resampled set, in order to guarantee the accuracy of the value range calculated for the target metabolite, is to find two additional points that respectively minimizes or maximizes the target flux in the intersection, i.e. subject to the combined SS and H constraints.
