## Supplementary material for "The FBA Solution Space Kernel – Introduction and Illustrative Examples": SSK software manual and code: Resamplebatch.bat

```
@ECHO off
ECHO "%~1"
REM The resampling specification as appropriate for
REM the specific data file that this batch file is
REM invoked for needs to be done in the separate file
REM ResamplingSpec.m available in the same directory.

REM Adjustments to global variables for Configuration.m 
REM are entered directly in the command line as for "verbose" etc. below.

 pushd %~dp0
 START "Commandline SSK Resampling" Resamplescript.wls "%~1" ^
 "{commandline=True, verbose=False, Species=\"Unknown\"}"
 PAUSE
 popd

REM The version above with the pushd/popd is a workaround because the Windows CMD interpreter
REM is unable to process UNC (i.e. network) file paths. It temporarily assigns an unused 
REM drive letter to the current path. The PAUSE is to delay reverting until the script has started.
REM If this fails, e.g. because there is no available drive letter,
REM the version below should work provided this batch file, SSkernelscript.wls and the Packages directory are 
REM all copied to a local file, e.g. on the desktop. Data files can stay on a network. 

REM START "Commandline Version of Resampling" /D %~dp0 SSKernelscript.wls "%~1" "{commandline=True Species=\"Unknown\"}"

REM This version may also be closer to what is required on other operating systems.
```
