## Supplementary material for "The FBA Solution Space Kernel – Introduction and Illustrative Examples": SSK software manual and code: Resamplescript.wls

```
#!/usr/bin/env wolframscript
(* ::Package:: *)

(* THIS SCRIPT RUNS EITHER HYPERSAMPLER OR TARGETVARIATION ON SUPPLIED DATA. *)
(* THE HYPERPLANES NEED TO BE DEFINED IN Hyperspec.m first. *)
(* TargetVariation processes al such hyperplanes together, but only returns the samples, while *)
(* HyperSampler returns both samples and rays for one specific hyperplane chosen below, default is 1 *)

workdir=DirectoryName[$InputFileName,2];
Print["Working directory is ",workdir]
SetDirectory[FileNameJoin[{workdir, "Packages"}]];
Get["Configuration`"]
Get["Centering`"]
Get["ModelProcessing`"]
Get["Resampling`"]
ResetDirectory[];
PackagesDirectory=FileNameJoin[{workdir,"Packages"}];

(* First the defaults and using the TEST directory *)
commandline=True; verbose=False; Species="Unknown";
datadir=workdir<>"TEST";
datafile=FileNames["*SSKernel.dif", datadir];
datafile=If[datafile=={},
 Print["Either the default directory TEST is missing or it contains no *.dif file!\n"];
 "Absent",First@datafile];
modid=If[datafile=="Absent","Unknown",
	StringDelete[FileBaseName[datafile], " SSKernel"~~___]];
modelfile=If[modid=="Unknown","None",
 FileNames[{modid ~~ ".mat", modid ~~ ".m"}, datadir,IgnoreCase->True]];
modelfile=If[modelfile=={},
 Print["No model specification was found in the TEST directory!"];
 "Absent",First@modelfile];
hyperfile=If[modid=="Unknown","None",
	FileNames[{modid ~~ " HyperSpec.m" , modid ~~ " HyperSpec.wl"}, datadir, IgnoreCase->True]];
hyperfile=If[hyperfile=={},
 Print["No resampling hyperplane specification was found in the TEST directory!"];
 "Absent",First@hyperfile];

(* Now process the arguments passed in the command line.*)
(* Print["The command line is: ",$ScriptCommandLine]; *)
   
args = Drop[$ScriptCommandLine, 1];
If[MatchQ[args, {arg__String} /; arg != ""],
(* StringRiffle reassembles an argument string that was split up by blanks *)
 args=StringTrim@StringRiffle[args];
(*  Print["Arguments passed: ",">"<>ToString@args<>"<"]; *)
 controls = StringCases[args, "{" ~~ __ ~~ "}"];
 controls = If[controls == {}, "", First@controls];
(*  Print["Control settings extracted are: ",">"<>controls<>"<"]; *)
 fpath = StringTrim@StringDrop[args, -StringLength@controls];
 datafile=If[StringLength[fpath]>1, fpath,datafile];
	If[FileExtension@datafile!="dif",
		Print["Nope! Datafile of type *."<>FileExtension@datafile<>
		" but I need a file of type *.dif as my primary input."];
		Input["Press x and <Enter> to exit"];Exit[]];
	datadir=DirectoryName[datafile];
	modid = StringDelete[FileBaseName[datafile], " SSKernel"~~___];
	modelfile = First@FileNames[{modid ~~ ".mat" , modid ~~ ".m" , modid ~~ ".wl"}, datadir,IgnoreCase->True];
	(* Where a target has been added, still use the original HyperSpec file *)
	hyperfile = datadir<>StringDelete[modid, " target"]<>" HyperSpec.m";
	hyperfile= FileNames[{StringDelete[modid, " target"] ~~ " HyperSpec.m" ,
						  StringDelete[modid, " target"] ~~ " HyperSpec.wl"}, datadir, IgnoreCase->True];
	hyperfile=If[hyperfile=={},
		Print["No resampling hyperplane specification was found in the data directory!"];
		"Absent",First@hyperfile];
	,
	controls=""];
(* Print["Finished argument processing"];*)
If[controls=="", 
	Print["Defaults used since no control parameters were passed from a batch file."],
	commandstring=StringReplace[controls,"'" -> "\""];
(* Next line executes control settings. *)
	commands=ToExpression[commandstring];
	Print["A total of "<>ToString@Length[commands]<>
	" parameter values are set by the command line, as follows:"];
	Print[commandstring];
];

(* Check that model specification and SSK output data files are found *)
If[!StringQ[datafile]||StringLength[datafile]==0,
	Print["The file path is empty! If you use Resamplebatch.bat, use "Open with " on the desired input file."];
	Input["Press x and <Enter> to exit"];Exit[]]
	
Print["\nI am working on the model identified as "<>modid];	
(*Print["My data directory is ",datadir];*)
Print["The SSK specification is taken from the file ",datafile]; 
Print["The original model stoichiometry is read from the file ",modelfile]; 
Print["The hyperplanes to be sampled are specified in the file ",hyperfile];
If[FileExistsQ[datafile],
	If[FileExistsQ[modelfile],
		If[!FileExistsQ[hyperfile],nospec=True;
			Print["Failed to find the HyperSpec.m input file! Using defaults."];
			dowhat = InputString["Press x and <Enter> to abort, else just <Enter> to proceed."];
			If[dowhat == "x", Exit[]],
			nospec=False],
		Print["Failed to find the model specification file! "];
		Input["Press x and <Enter> to exit"];Exit[]],
	Print["Failed to find the .dif SSK specification file! "];
	Input["Press x and <Enter> to exit"];Exit[]	];
Print[""];

(* Read the model stoichiometry *)
success=True;
If[StringLength[Species]<1,  Species =FileNameTake[datafile, {-2}]]; 
result= CheckAbort[LoadModel[modelfile, verbose], success = False];
If[!success,Print["Load and test of the model file has failed."];Pause[0.5];
			Input["Press x and <Enter> to exit"];Exit[]];  
Print[""];

(* Read the SSK data  *)
Clear[mainhead, SectionHeadings, importdata, KernelSpace, PeriPoints, 
  fixvals, thindirs, rays, insphere, fixdirs, MetaboliteNames, 
  ReactionNames, ReducedSS];
importdata = {KernelSpace, PeriPoints, fixvals, thindirs, rays, 
   insphere, fixdirs, {MetaboliteNames, ReactionNames} , ReducedSS};
Evaluate[Flatten[Prepend[
     Table[{SectionHeadings[i], importdata[[i]]}, {i, Length[importdata]}], mainhead], 1]] = 
		ToExpression /@ Flatten[Import[datafile, "DIF", "IgnoreEmptyLines" -> True]];
Print["SSK data read in successfully.\n"]

(* Read the resampling hyperplane specification and set the target *)
If[nospec,
	(* Default hyperplane used in the absence of a specification file.*)
	target=-1;
	targetfluxname="";
	targetproduced=True;
	cols = Length@ReducedSS[[2, 1]];
	nonfixes = Complement[Range[cols], fixvals[[All, 1]]];
	hyperlist = Partition[nonfixes, 1];
	valuelist = ConstantArray[{0}, Length@hyperlist],
	Get[hyperfile]];
	
(* Set the target column .*)
lastcol=Length@ReactionNames;
targetcol = TargetInsert[target,targetfluxname,targetproduced];
If[NumberQ[targetcol],
	If[targetcol>lastcol, 
		Print["To repeat the SSK calculation, run SSKernelscript on the expanded input file."<>
	"\nThis is being stored in the same directory, as "<>modid<>" target.m"<>
	"\nThen rerun Resamplescript on the resulting target .dif file."];
	newname=modid<>" plus target";
	Export[datadir<>"\\"<>modid<>" target.m",
	{newname,ReactionNames,MetaboliteNames,S,Svals,bounds,objectselector,
	FBAvector,If[maxmin==1,"min","max"]},"List"];
		Pause[0.5];InputString["Press x and <Enter> to exit."];Exit[]
	,
	Print["Target identified successfully.\n"]]
	,
	Print[targetcol];
	Pause[0.5];InputString["Press x and <Enter> to exit."];Exit[]
	]
	
Print["Ready to proceed with resampling the SSK generated from:\n"<>DeleteCases[StringSplit[mainhead, {"(* ", " *)"}], "\n"]];
dowhat = InputString[
   "Press \tt and <Enter> to run TargetVariation,\n\th and <Enter> to run HyperSampler,\n\tx and <Enter> to abort.\n"];
Switch[dowhat,
	"x"|"X", Exit[], 

	"t"|"T",
(* Sample each hyperplane and collect data on the target flux *)
	Print["Working on the resampling..."];
	samples = TargetVariation[hyperlist, valuelist, PeriPoints, rays,  ReducedSS, targetcol, verbose];
	Print["\nSamples collected for "<>ToString@Length@samples<>" hyperplanes, including the full SSK."];
	Pause[0.5];
	whatnow = InputString[
   "Press x and <Enter> to finish, else just <Enter> to store the sample fluxes in a file first."];
	If[whatnow == "x", Exit[],
	Export[datadir<>"\\"<>modid<>" TVarSamples.wl",samples];
	Print["Sample data saved to a file."]],
	
	"h"|"H",
(* Choose which hyperplane to sample. *)
	n=0;
	While[!NumberQ[n] || n<1 || n>Length@hyperlist, 
	n=ToExpression@InputString["Enter the sequence number in the HyperSpec file, of the hyperplane to be sampled."];
	If[!NumberQ[n] || n<1 || n>Length@hyperlist,
		Print["Invalid choice; there are only "<>ToString@Length@hyperlist<>" hyperplanes to choose from. Try again."];Pause[0.1]];
    ];	
(* Sample the chosen hyperplane and find its rays *)
	{Hsample, Hrays} = 
	HyperSampler[hyperlist[[n]], valuelist[[n]], PeriPoints, rays, ReducedSS, targetcol, verbose];
	Print[ToString@Length[Hsample]<>" sample fluxes and "<>ToString@Length[Hrays]<>
			" rays collected from hyperplane no  "<>ToString[n]];
	Pause[0.5];
	whatnow = InputString[
   "Press x and <Enter> to finish, else just <Enter> to store the sample fluxes and rays in a file first."];
	If[whatnow == "x", Exit[],
	Export[datadir<>"\\"<>modid<>" HypSamples.wl",{Hsample, Hrays}];
	Print["Sample fluxes and rays saved to a file."]]	
];

Pause[0.5];
InputString[
   "Press x and <Enter> to finish and close this window."];
Exit[];
```
