## Supplementary material for "The FBA Solution Space Kernel – Introduction and Illustrative Examples": SSK software manual and code: SSKernelbatch.bat

```
@ECHO off
ECHO "%~1"
REM Edit the commandline below to insert further comma separated parameter assignments, 
REM inside the curly brackets.

 pushd %~dp0
 START "Commandline Version of SSKernel" SSKernelscript.wls "%~1" ^
 "{commandline=True, loadreset=False,  verbose=False, Species=\"Unknown\";}"
 PAUSE
 popd

REM The version above with the pushd/popd is a workaround because the Windows CMD interpreter
REM is unable to process UNC (i.e. network) file paths. It temporarily assigns an unused 
REM drive letter to the current path. The PAUSE is to delay reverting until the script has started.
REM If this fails, e.g. because there is no available drive letter,
REM the version below should work provided this batch file, SSkernelscript.wls and the Packages directory are 
REM all copied to a local file, e.g. on the desktop. Data files can stay on a network. 

REM START "Commandline Version of SSKernel" /D %~dp0 SSKernelscript.wls "%~1" "{verbose=False}"

REM This version may also be closer to what is required on other operating systems.
```
