## Supplementary material for "The FBA Solution Space Kernel – Introduction and Illustrative Examples": SSK software manual and code: SSKernelscript.wls

```
#!/usr/bin/env wolframscript
(* ::Package:: *)

workdir=DirectoryName[$InputFileName,2];
Print["Working directory is ",workdir]

SetDirectory[FileNameJoin[{workdir,"Packages"}]];
Get["Configuration`"]
Get["ModelProcessing`"]
Get["Centering`"]
Get["RedundancyRemoval`"]
Get["FixedFluxes`"]
Get["RayFinding`"]
Get["FacetTesting`"]
Get["FBFsearch`"]
Get["Capping`"]
ResetDirectory[];
PackagesDirectory=FileNameJoin[{workdir,"Packages"}];
commandline=True; Species="Test"; verbose=False;

datadir=workdir<>"TEST"; 
mfiles=FileNames[{"*.mat","*.m","*.wl"}, datadir, IgnoreCase->True];
datafile=First@Cases[mfiles, _?(! StringMatchQ[#, __ ~~ "HyperSpec." ~~ __] &)]
(* Print["My data directory is ",datadir];
   Print["My data file is ",datafile];  *)

(* Print["The command line is: ",$ScriptCommandLine]; *)
   
args = Drop[$ScriptCommandLine, 1];
If[MatchQ[args, {arg__String} /; arg != ""],
(* StringRiffle reassembles an argument string that was split up by blanks *)
 args=StringTrim@StringRiffle[args];
 (* Print["Arguments passed: ",">"<>ToString@args<>"<"]; *)
 controls = StringCases[args, "{" ~~ __ ~~ "}"];
 controls = If[controls == {}, "", First@controls];
 (* Print["Control settings extracted are: ",">"<>controls<>"<"]; *)
 fpath = StringTrim@StringDrop[args, -StringLength@controls];
 (* Print["Extracted fpath = >"<>fpath<>"<"]; *)
 datafile=If[StringLength[fpath]>1, fpath,datafile];
 fext=ToLowerCase@FileExtension@datafile;
	If[fext!="mat" && fext!="m" && fext!="wl",
		Print["Nope! Datafile of type *."<>fext<>
		" but I need a file of type *.mat or *.m or *.WL as my primary input."];
		Input["Press x and <Enter> to exit"];Exit[]];
	datadir=DirectoryName[datafile];
	,
	controls=""];

If[controls=="", 
	Print["Control parameters will default to settings in Configuration.m."],
	commandstring=StringReplace[controls,"'" -> "\""];
(* Next line executes control settings. *)
	commands=ToExpression[commandstring];
	Print["A total of "<>ToString@Length[commands]<>
	" parameter values are adjusted by the command line, as follows:"];
	Print[commandstring];
];

If[!StringQ[datafile]||StringLength[datafile]==0,
	Print["The file path is empty! If you use SSKernelbatch.bat,use "Open with " on the desired input file."];
	Input["Press x and <Enter> to exit"];Exit[]]

Print["My data file is ",datafile]; 
If[FileExistsQ[datafile], Print["I found the data file, allright!"],
Print["No file was found at the path "<>datafile];Input["Press x and <Enter> to exit"];Exit[]];
Print[""];

dowhat = 
  InputString[
   "Press x and <Enter> to abort, else just <Enter> to proceed."];
If[dowhat == "x", Exit[]];
Print[""];

(* STAGE 1 *)
success=True;
If[StringLength[Species]<1,  Species =FileNameTake[datafile, {-2}]]; 
result= CheckAbort[LoadModel[datafile, verbose], success = False];
Print[result];
Heading = {"STAGE 1 COMPLETED: Loaded data for " <> Species, "from " <> FileNameTake[datafile] };
If[success,Print[Heading],Print["Load and test of the datafile has failed."];Pause[0.5];
			Input["Press x and <Enter> to exit"];Exit[]];  
Print[""];			

(* STAGE 2*)
SStype = "FacetCone";
Reductiontable = {{"Stage", " Constraints ", " Variables ", " Ray Yield "}};
{RSStime, result} = Timing[
   	result = CheckAbort[Check[DoFBA[Tolerances, True], success = False], success = False];    
   		If[StringQ[result], success = False;
     	Print["LP solution of the imported FBA model has failed. Retry Stage 2 after
 adjusting the LP tolerance, or change LP method in the datafile.  "]];
	result = CheckAbort[Check[
   		ReducedSolutionSpace[S, bounds, objectselector, maxmin, verbose],
   		 success = False],  success = False];
	If[StringQ[result], success=False; Print[result]; PrependTo[feasiblepoints,{}]]];
 Heading = {"STAGE 2 COMPLETED: Reduced Solution Space for " <> Species, 
   	"  Model " <> ModelName <> " of type " <> ToString@SStype <> 
   	"\n calculated on " <> DateString["DateShort"] <> " in " <> 
      Quiet@ToString[NumberForm[RSStime, 3]] <> " seconds."};
If[success,Print[Heading],Print["RSS reduction of the solutionspace has failed."];
		SaveResults[{}, datafile];
		Input["Press x and <Enter> to exit"];Exit[]];
Print[""];		

(* STAGE 3 *)
{Kerneltime, result} =  Timing[CheckAbort[Check[
    	KernelFinder[verbose],
    	 success = False], success = False]];
 Heading = {"STAGE 3 COMPLETED: Capped Solution Space for " <> Species, 
     " Model " <> FileBaseName[datafile] <> " of uncapped type " <> ToString@SStype <>
       "\n calculated on " <> DateString["DateShort"] <> " in " <> 
     Quiet@ToString[NumberForm[Kerneltime, 3]] <> " seconds."};
If[success,
	SSKinRSS = KernelSpace; 
    SSKinsphere = InscribedSphere;
	Print[Heading],
	
	Print["Calculation of a closed SSK has failed."];
		SaveResults[{}, datafile];
		Input["Press x and <Enter> to exit"];Exit[]];  
Print[""];		

(* STAGE 4 *)
If[SStype != "Point",
    {Shapetime, result} = Timing[CheckAbort[Check[KernelShape[maxaspect, maxthin, verbose],
    	 success = False], 	 success = False]]; 
    Heading = {"STAGE 4 COMPLETED: Shape Analysis for " <> Species, 
      " Model " <> FileBaseName[datafile] <> "  with uncapped kernel of type " <> 
      ToString@SStype <> 
       "\n calculated on " <> DateString["DateShort"] <> " in " <> 
       Quiet@ToString[NumberForm[Shapetime, 3]] <> " seconds."};,
    Shapetime = 0; 
    flatplot = Graphics@Text["Point SSK, has no shape."]];
If[success, Print[Heading],Print["Shape analysis of the closed SSK has failed."];
		SaveResults[{}, datafile];
		Input["Press x and <Enter> to exit"];Exit[]];   
Print[""];		

(* STAGE 5 *)
{Chordtime, result} = Timing[CheckAbort[Check[
    	{chordpic, cenpic} = KernelDisplay[ExportResults, verbose],
    	 success = False],  success = False]];
 Heading = {"STAGE 5 COMPLETED: Full SS Kernel Analysis for " <> Species, 
     " Model " <> FileBaseName[datafile] <> "  with uncapped kernel of type " <> 
      ToString@SStype <> "\n calculated on " <> DateString["DateShort"] <> " in " <> 
      Quiet@ToString[NumberForm[RSStime + Kerneltime + Shapetime + Chordtime, 3]] <> 
      " seconds."}; 
If[success, Print[Heading],Print["Display of chords and centering has failed."];
		SaveResults[{}, datafile];
		Input["Press x and <Enter> to exit"];Exit[]];    
Print[""];		
		
(* STAGE 6 *)
Heading = {"Full SS Kernel Analysis for " <> Species, 
     " Model " <> FileBaseName[datafile] <> "  with uncapped kernel of type " <> 
      ToString@SStype <> "\n calculated on " <> DateString["DateShort"] <> " in " <> 
      Quiet@ToString[NumberForm[RSStime + Kerneltime + Shapetime + Chordtime, 3]] <> 
      " seconds."}; 
	  
CheckAbort[success=Check[
   	SaveResults[If[Length@ExportResults==0,{},ExportResults[[1;;9]]], datafile],
   	 False],  success = False];

If[success, Print["Output files have been stored successfully."],
	Print["Export of output files has failed."]; 
	Pause[0.5];
	Input["Press x and <Enter> to exit"];Exit[]];  

Pause[0.5];
Input["Press x and <Enter> to exit"]; 
```
