## Supplementary material for "The FBA Solution Space Kernel – Introduction and Illustrative Examples": SSK software manual and code: Template HyperSpec.m

```
(* NB: This file should be copied and edited for each dataset to 
	specify the target and hyperplanes to be resampled. *)
	
verbose=False;
(* Specify the target in one of three ways: *)
(*	1. A string contained in the name of a metabolite to be *) 
(*		produced/absorbed by the metabolic network. *)
(*	2. An integer, the number of the flux column in the *)
(*		stoichiometry matrix that provides a source/sink for the metabolite. *)
(*	3. An explicit column for S that defines the source/sink flux. *)
(*	The targetfluxname is for use with option 3 but ignored for an existing flux .*)
(*	The targetproduced value controls whether it is a source or sink. *)
target="H2O";
targetfluxname="";
targetproduced=True;

(* Set up the list of hyperplanes to be sampled, as each variable flux is
   in turn fixed to zero*)
cols = Length@ReducedSS[[2, 1]];
nonfixes = Complement[Range[cols], fixvals[[All, 1]]];
hyperlist = Partition[nonfixes, 1];
valuelist = ConstantArray[{0}, Length@hyperlist] ;         

(* Alternatively, hyperplanes can be specified explicitly, *)
(*  e.g.each given as a list of S matrix columns *)
(* The valuelist gives constant values or ranges for hyperplane constraints. *)
(*
hyperlist = {{20}, {20, 22, 27}, {249}};
valuelist = {{-5.3}, {{-5.3, 1}, -0.05, {-0.2, -1}}, {{0.35, 0}}};      
*)
```
