## Supplementary material for "The FBA Solution Space Kernel – Introduction and Illustrative Examples": SSK software manual and code: Capping.m

```
(* ::Package:: *)

(* COPYRIGHT
					© Copyright 2022 Wynand Verwoerd

This file is part of SSKernel.

The SSKernel program is free software; you can redistribute it and/or modify it under the terms of the GNU General Public License as published by the Free Software Foundation; either version 3 of the License, or (at your option) any later version.
 
This program is distributed in the hope that it will be useful, but WITHOUT ANY WARRANTY; without even the implied warranty of MERCHANTABILITY or FITNESS FOR A PARTICULAR PURPOSE . See the GNU General Public License for more details.
 
You should have received a copy of the GNU General Public License along with this program . If not, see http://www.gnu.org/licenses/
*)

(* Wolfram Language Package *)

BeginPackage["Capping`", { "Configuration`", "RedundancyRemoval`", "Centering`", "FBFSearch`","RayFinding`"}]
(* Exported symbols added here with SymbolName::usage *) 
CoincidenceCapper::usage="Function to perform coincidence capping and reduction to a given facet." 
TangentCapper::usage="Tangential capping hyperplanes are applied to truncate remaining rays. "
Deconstructor::usage="The margin by which a given feasible flux falls outside reach of the SSK"
FlatnCentre::usage="Fix periphery points at main diameters and Flattens bounded solutionspace along any thin directions."
WrappingRadii::usage=" Estmate coverage of SSK by periphery point polytope as ratio of enclosing radii."
CoverageInterval::usage=" Estmate % coverage of SSK by periphery point polytope."

(* TESTING *)
directs::usage=""
samplepoints::usage=""

Begin["`Private`"] (* Begin Private Context *) 

SetAttributes[CoincidenceCapper, HoldAll];
CoincidenceCapper[solutionspace_, RayMat_, raydim_, facet_, printresults_: False] :=
 (* This routine condenses the solution space specification by coincidence 
capping to the specified facet (usually FBF progenitor hyperplane). 
 It returns the rays that fall away in the process, as vectors in the variable part of flux space *)
 Module[{constraints, values, SSBase, origin,  cons, vars, centershift, ProgenBasis, ProgenTransform, ProgenRays, 
   ProgenCenter, DiscardRays, keepers, redundancies, reduced, tol = 10^-10},
   
  {{constraints,values},{origin,SSBase}}=solutionspace; 
  {cons, vars} = Dimensions[constraints];
  progresslabel="Perform coincidence capping.";
  Print[Style[progresslabel, Blue, TextAlignment -> Center]];
  (* Separate rays that are in and out of the facet *)
  ProgenRays = Select[RayMat, Total[#[[facet]]] == 0 &][[All, cons + 1 ;;]];
  DiscardRays = Select[RayMat, Total[#[[facet]]] != 0 &][[All, cons + 1 ;;]];
  
  (* STAGE 1: 
  PROJECT CONSTRAINTS TO THE PROGENITOR HYPERPLANE AND TRIM AWAY REDUNDANCIES *)
  
  ProgenBasis = NullSpace[constraints[[facet]],Tolerance->LPtol];
  (* For the origin to lie on the progenitor facet, 
  it must satisfy all progenitor constraints with an = not \[LessEqual] . 
  Calculate the shift to a new origin, as a vector in the current OS *)
  centershift = PseudoInverse[constraints[[facet]]].values[[facet]];
  ProgenTransform = {centershift, ProgenBasis};
  {constraints, values} = 
   Chop[N@DowncastConstraints[constraints, values, ProgenTransform], tol];
  (* Check if the shifted center is inside the progenitor polytope, if not recenter *)
  If[Min[values] < 0,
   ProgenCenter = CLODcentre[constraints, values];
   (*Print["Origin projected to progenitor hyperplane is exterior to feasible polytope - 
   CLOD centering shifts it to the progenitor coordinate location "<>ToString[ProgenCenter]];*)
   values = Chop[values - constraints.ProgenCenter, tol];
   (* Print["After recentering to progenitor, the values vector is : ",values];*)
   centershift = centershift + Chop[ProgenCenter.ProgenBasis, tol];
   ProgenTransform = {centershift, ProgenBasis}
   ];
  (*Print["Location of progenitor center relative to the OS origin: "<>ToString[centershift]]; *)
  keepers = Complement[Range[cons], facet];
  {cons, vars} = Dimensions[constraints];
  If[cons != Length[keepers], 
   Print[" PROJECTION DISCREPANCY - 
fewer than expected constraints remain after projection to progenitor plane. "]; 
   Return[]];
   progressrange={0,cons};progresslabel=" Eliminating redundant progenitor constraints ";	
   Print[Style[progresslabel, Blue, TextAlignment -> Center]];      
   {constraints,values,redundancies} = RedundancyTrimmer[constraints, values,{}, printresults];
   progresslabel=" Progenitor redundant constraint elimination completed."; 
   progresscounter=0;

  
  {cons, vars} = Dimensions[constraints];
  If[Length[redundancies] > 0, 
   keepers = Delete[keepers, Partition[redundancies, 1]]];
   If[printresults,
  Print[" Coincidence capping to the progenitor hyperplane, removing redundant 
  constraints and recentering, reduces the SS specification to " <> 
    ToString[cons] <> " constraints on " <> ToString[vars] <> " variables."]];
  
  (* STAGE 2: PROCESS THE RAY MATRIX AND FBF'S *)
  
  (* The following 3 lines project existing rays and reconstitute a reduced ray matrix *)
  (*
  ProgenRays=ProgenRays.Transpose[ProgenBasis];
  newrayolaps=Chop[ProgenRays.Transpose[constraints],tol];
  RayMat=Join[-newrayolaps,ProgenRays,2];
  *)
  (* But although slower, finding the reduced ray matrix from scratch avoids 
  the possibility of missing any and also gives rays that are more peripheral *)
  {RayMat,raydim} = CompleteRayBasis[constraints,printresults];
  
  (* FBF's already known and deposited in the global array BoundedFacets may 
be reduced to facilitate a subsequent tree search in the reduced space. 
  First reduce the FBF's to the surviving constraint projections,then 
  re-express them in terms of progenitor numbering:*)
  reduced = Map[Intersection[keepers, #] &, BoundedFacets];
  BoundedFacets = 
   Flatten /@ Map[Flatten[Position[keepers, #]] &, reduced, {2}];
  reduced = Map[Intersection[keepers, #] &, Infeasibles];
  Infeasibles = Flatten /@ Map[Flatten[Position[keepers, #]] &, reduced, {2}];
  
  (* STAGE 3: 
  EXPRESS EVERYTHING BACK IN TERMS OF THE FLUX SPACE VARIABLES *)
  solutionspace =  {{constraints,values},{centershift,ProgenBasis}};
  DiscardRays =  Chop[DiscardRays.RSSBasis, tol]
  ]

SetAttributes[TangentCapper, HoldFirst];
TangentCapper[solutionspace_, RayMat_, RSSbasis_, printresults_: False] := 
(* New version of TangentCapper, that introduces a capping hyperplane for
    each ray left in the progenitor; some of these are eliminated if redundant.
    *)
 Module[{constraints, values, Basis, origin, condim, cons, vars, 
   caprays, XValues, XRange, X, ApexPoints, apexintercept, 
   facetcons, capvec, R, redundancies, tempvalues, centershift, center, cshift,
   captype="Tangent", tol = 0.00001},
   {{constraints, values}, {origin, Basis}} = solutionspace;
  condim = Dimensions[constraints];
  If[Length[condim] == 2, {cons, vars} = condim, 
   Print["Invalid - Constraints must be a matrix."]; Return[{}]];
  XRange = ConstantArray[{-Infinity, Infinity}, vars];
  XValues = Map[{#, -1} &, values];
  (* Capping just "interior" rays found by CapRayFinder sometimes still leaves
  long extensions in some directions; better to combine these with the known
  peripheral ray basis. This can give a large number of capping constraints, 
  but redundncy elimination will remove some at least.  *)
  (* For a large dimension count, Caprayfinder can stall. It is a nice-to-have, 
  the ray matrix already gives a complete basis of ray space, so capping
  all rays should be sufficient to ensure a bounded SSK. *)
  caprays = If[vars < 120, TimeConstrained[CapRayFinder[constraints],timeconstraint,{}],{}]; 
  caprays = Join[caprays, RayMat[[All, cons + 1 ;;]]];
  (* Print["Capping ray norms are: "<>ToString[Norm/@caprays]]; *)
  CappingRadii = {};
  If[SStype == "SimpleCone",(*Just add the default radius capping constraints.*)
  	captype="Default";   CappingRadii = ConstantArray[DefaultCap, Length[caprays]];
   ,
   (*Else, calculate a radius from FBF intersections for each capping ray*)
   progressrange = {0, Length@caprays};
   progresslabel =  " Tangent capping: Calculate radii for all progenitor rays ";
   Print[Style[progresslabel, Blue, TextAlignment -> Center]];
   Do[capvec = caprays[[cap]]; progresscounter++;
    (*Loop over all FBF's to find the apex point that maximizes \
overlap with current capping direction*)
    ApexPoints = Reap[Do[facetcons = BoundedFacets[[bf]];
        XValues[[facetcons, 2]] = 0;(*Fix solutions to the facet hyperplane*)
        X = Quiet[Check[LinearProgramming[-capvec, constraints, XValues, 
            XRange, Method -> "Simplex", Tolerance -> tol], 
           "No Solution", {LinearProgramming::lpsnf}], {LinearProgramming::lpsnf}];
        XValues[[facetcons, 2]] = -1;(*Restore the inequality constraints*)
        If[VectorQ[X, NumberQ], Sow[{X.capvec, bf}], 
        (* Print["No apex found for bounded facet " <> ToString[bf]]; *)
          Null ];, {bf, Length[BoundedFacets]}]][[2]];
    (*Find the highest apex and add it to the constraints list*)
    If[Length[ApexPoints] > 0, ApexPoints = ApexPoints[[1]]];
    R = Max[ApexPoints[[All, 1]]];
    AppendTo[CappingRadii, R];
    apexintercept = MaximalBy[ApexPoints, First][[All, 2]];
(*    If[printresults, 
     PrintTemporary["Capping radius " <> ToString[cap] <> 
        " fixed to " <> TextString[R] <>
        " required by intercepting hyperplane " <> ToString[apexintercept]]]; *)
    , {cap, Length[caprays]}];
    progresslabel = " Capping radii calculation completed.";
    progresscounter = 0];

   constraints = Join[constraints, caprays];
   values = Join[values, CappingRadii];
   (*Capping may have excluded the old origin; in this case first move it back inside.*)
   If[Min[values] < 0., centershift = CLODcentre[constraints, values];
    values = values - constraints.centershift;
 (* Print["After capping and recentering , the values vector is : ",values]; *)   
    origin = origin + centershift.Basis];
    CappingRadii=Chop[values[[-Length@caprays;;]],tol];
    
	If[captype == "Tangent",
		If[printresults,Print["The capping radii based on FBF intercepts, are: ",TextString/@Sort[CappingRadii]]];
	AppendTo[Processreport, {"FBF-based capping radii range from " <> TextString[Min@Abs@CappingRadii] <>
  	" to " <> TextString[Max@Abs@CappingRadii] }]];
 
  If[fixflag && Max[CappingRadii] < 4*fixtol,
  	captype="Default"; 
  	CappingRadii = ConstantArray[DefaultCap, Length[caprays]];
  	values[[-Length@caprays;;]] =  CappingRadii;
  	Print[Style["Flux components as well as all capping radii have values in a similar range as the fixed tolerance.",
  		Darker@Green],"This indicates that the SSK is similar to a simple cone, even though its base is not a single point at the origin,\
  but instead a small hypervolume, with extent comparable to the fixed tolerance.
  It is treated similarly as the simple cone, by taking the default value 
  for all capping radii, to allow shape characterisation of the cone.   "];
    AppendTo[Processreport, {Style["Negligible capping radii (compared to fixtol) were replaced by default capping, expanding
 the Kernelspace to a cone for shape characterisation. ",Darker@Green] }]
    ,
    fixflag=False
  ];	
   
    
   (*Where the capping plane intersects an FBF, it will generally make constraints redundant.
   But this cannot happen for a simple cone,so only trim for FacetCone.*)
  If[SStype == "FacetCone" && captype == "Tangent",
   progressrange = {0, Length@constraints};
   progresslabel = 
    " Eliminating redundant constraints after tangent capping ";
    Print[Style[progresslabel, Blue, TextAlignment -> Center]];
   {constraints, values, redundancies} = 
    RedundancyTrimmer[constraints, values, {}, printresults];
   solutionspace =Chop[{{constraints, values}, {origin, Basis}},tol] ;
   progresslabel = 
    " Tangent capping redundant constraint elimination completed.";
   progresscounter = 0;
   If[printresults && Length[redundancies] > 0, 
    Print[" After capping, " <> ToString[Length@redundancies] <> 
    	" constraints were found redundant and were removed."]]];
    	
  (* If MiniMaxCentre returns a zero radius, the centre point falls on a facet. 
  Moreover, this may well be on a higher level facet than the minimum possible. 
  In this case, e.g. for
  a simple cone but where there are still unresolved fixed value directions,
  moving consecutively along each capping ray direction to its midpoint does 
  a better job of driving the facet level down. *)
  
  centershift = MiniMaxcentre[constraints, values, printresults];
  If[InscribedSphere[[1]] < 2*fixtol,  
  	Print[Style[
   "The inscribed sphere radius " <>TextString[InscribedSphere[[1]]]<> 
   " is comparable to the fixed value tolerance.", Darker@Green],
   "\n  This indicates that there may be residual fixed values, 
     that still have to be eliminated in the flattening stage."];
  	tempvalues = values;
	centershift = ConstantArray[0., vars];
	Do[center = Mean[Radii[constraints, tempvalues, caprays[[i]]]]; 
  		cshift = center*caprays[[i]];
  		centershift = centershift + cshift;
  		tempvalues = Chop[tempvalues - constraints.cshift,tol];
  		(* Terminate traversal of caprays if it has wandered outside the ploytope,
  		to avoud trouble with Radii in the next round. Subsequent Retrieval corrects it.*)
  		If[Min[tempvalues]<0,Break[]];
 	 , {i, Length[caprays]}]; 
    centershift=Retrieve[solutionspace,centershift,printresults];
  ];
  values = values - constraints.centershift;
  origin = origin + centershift.Basis;
  solutionspace = Chop[{{constraints, values}, {origin, Basis}},tol];
  InscribedSphere[[2]] = InscribedSphere[[2]]-centershift;
  
  (*Rays are currently in the progenitor facet;
  transfer them upwards first to RSS and then to FS*)
  (*Print[Dimensions/@{caprays,Basis,RSSbasis}];*)
  caprays = Chop[RayMat[[All, cons + 1 ;;]].Basis.RSSbasis, tol];
  {cons, vars} = Dimensions@constraints;
  AppendTo[Kerneltable, {captype<>" capping of " <>ToString@Length[caprays] 
      <>" progenitor rays", cons, vars, Length[caprays]}];

  caprays]
  
Deconstructor[fluxpoint_, Preflat_, Postflat_, deconstruct_:False] :=
  (*This function determines the flattening truncation \
margin for a given fluxpoint in the space Preflat,by deconstructing it into 3 \
components: a vector Q in Postflat; and a ray vector R and a vector T \
(the residue) in Preflat such that the L1-norm of T is minimised. *)

(* NOTE: This function assumes that Postflat is specified relative to Preflat.
	If they are both specified relative to RSS, first reconfigure Preflat. *)
 Module[{m, n, MM, NN, 
 	Preflatcons, Preflatvals, Preflatorigin, Preflatbase, 
 	Postflatorigin, Postflatcons, Postflatvals, Postflatbase, 
 	flatpoint, inplane, interior,
 	INN, Zcon1, Zcon2, CScon, Scon, Addcon, ConMat, bounds, Zobjective,
 	Vals, QRTZ, QRT, trunc, lptol = 0.001},
  
  {{Preflatcons, Preflatvals}, {Preflatorigin, Preflatbase}} = Preflat;
  {{Postflatcons, Postflatvals}, {Postflatorigin, Postflatbase}} = Postflat;
  {m, n} = Dimensions[Postflatcons];
  {MM, NN} = Dimensions[Preflatcons];
(*  Print[{"Deconstructor pre- and post-flattening constraint dimensions: ",{m,n},{MM,NN}}]; *)

(* Test if fluxpoint is already inside the Postflat polytope; 
	if so LP can be skipped as deconstruction  is trivial, R = T = 0 .
	Taking R = 0 assumes that both Pre- and Postflat spaces are bounded. 
	That is true during actual flattening, which asks for just the residue.
	But when full deconstruction is required, as when the nominal Preflat is 
	actually RSS, that is not true; so take the shortcut only in the former case.  *)
 If[!deconstruct,
  flatpoint = DowncastPoint[fluxpoint, {Postflatorigin, Postflatbase}];
  interior = Min@Chop[Postflatvals - Postflatcons.flatpoint] >= 0;
  inplane =  Chop@Norm[fluxpoint - UpliftPoint[flatpoint, {Postflatorigin, Postflatbase}]] == 0;
  If[inplane && interior, 
  (* Print["Deconstructor is skipping the LP "]; *)
  Return[If[deconstruct, {flatpoint, ConstantArray[0., NN], ConstantArray[0., NN]}, 0.]]]
  ];

  INN = IdentityMatrix[NN];
  CScon = Join[Postflatcons, ConstantArray[0., {m, 3 NN}], 2];
  Scon = Join[ConstantArray[0., {MM, n}], Preflatcons, 
    ConstantArray[0., {MM, 2 NN}], 2];
  Addcon = 
   Join[Transpose[Postflatbase], INN, INN, 
    ConstantArray[0., {NN, NN}], 2];
  Zcon1 = 
   Join[ConstantArray[0., {NN, n}], ConstantArray[0., {NN, NN}], 
    INN, -INN, 2];
  Zcon2 = 
   Join[ConstantArray[0., {NN, n}], 
    ConstantArray[0., {NN, NN}], -INN, -INN, 2];
  ConMat = Join[CScon, Scon, Addcon, Zcon1, Zcon2];
  (*Print[MatrixForm@ConMat];*)
  (*Print["CScon,Scon, Addcon,Zcon1,Zcon2"];
  Map[Print[Dimensions[#]]&,{CScon,Scon,Addcon,Zcon1,Zcon2}];*)
  bounds = Join[ConstantArray[{-Infinity, Infinity}, n + 2 NN], 
    ConstantArray[{0, Infinity}, NN]];
  Zobjective = 
   Join[ConstantArray[0., n + 2 NN], ConstantArray[1., NN]];
  Vals = Join[Postflatvals, ConstantArray[0., MM], 
    fluxpoint - Postflatorigin, ConstantArray[0., 2 NN]];
  (*Print[Vals];*)
  Vals = Transpose[{Vals, 
     Join[ConstantArray[-1, m + MM], ConstantArray[0, NN], 
      ConstantArray[-1, 2 NN]]}];
  (*Print["Zobjective,ConMat,Vals,bounds"];
  Map[Print[Dimensions[#]]&,{Zobjective,ConMat,Vals,bounds}];
  *)
  QRTZ = Quiet[
    Check[LinearProgramming[Zobjective, ConMat, Vals, bounds, 
      Method -> "Simplex", Tolerance -> lptol], 
     "Deconstruction failed - LP solver found no solution", \
{LinearProgramming::lpsnf}], {LinearProgramming::lpsnf}];
  
  QRT = If[VectorQ[QRTZ, NumberQ], 
    Chop[{QRTZ[[1 ;; n]], QRTZ[[n + 1 ;; n + NN]], 
      QRTZ[[n + NN + 1 ;; n + 2 NN]]}, lptol], 
      Print[QRTZ]; QRTZ];
  trunc=If[Length[QRT] > 0, 
  	QRT[[1]] = UpliftPoint[QRT[[1]], Postflat[[2]]]; Norm@QRT[[3]],
  	 QRT];
  If[deconstruct, QRT, trunc]
  ]

SetAttributes[FlatnCentre, HoldAll];
FlatnCentre[solutionspace_, maxaspect_, negligible_,   PrintResult_: False] :=
(*This function uses the main diameter directions of the SS polytope as a basis for the flattened SS.
 Any thin directions that it eliminates are returned as a set of vectors in FS,orthogonal both mutually and to the SSK.
 The global array Peripoints are recentered and reduced to any flattened SS.*)
 Module[{constraints, values, OSbase, OSorigin, cons, vars, chords, 
   chordvecs, neglect,   keepers,  thins, thindirs = {}, flatten = True, plotit = True, 
   diametertest, chordtest, FlatBasis, flatcount=0, flatrank, flatcenter, 
   centershift, inscribedcenter, FlattenTransform, 
   redundancies,   poss, negs, lpchords, endpoints, getmargin = False, 
   maxlo, margin, residues, norms, reached = 0, outside,  apexpoint, 
   warn = Style["\nWARNING!", Red, Bold], warning = "",  tol = 0.0001},
  
  {{constraints, values}, {OSorigin, OSbase}} = solutionspace;
  {cons, vars} = Dimensions[constraints];
  
  (*  1. FIND OR LOAD THE MAIN CHORDS. 
  	On subsequent passes of the flattening stage,  only recalculate main chords if 
  the minimum required by LP is more than the ones already known. *)
  
  lpchords = Count[PreflatChordLengths, _?(Positive[#] &)];
  (*Print[{" Already known LP chords ",lpchords,chordmin,vars}];*)
  If[lpchords > 0 && (lpchords >= chordmin || lpchords == vars),
   If[PrintResult,Print["Skipping chord calculation "]];
   chords = PreflatChords; ChordLengths = PreflatChordLengths;
   PeriPoints = PreflatPeris; solutionspace = PreflatSSK,
   If[PrintResult,Print["Doing chord calculation "]];
   progresslabel = "Before flattening: ";
   chords = MainChords[solutionspace, chordmax, chordmin, flipmax, PrintResult];
(* Print["The chords found, are: ",chords]; *)   
  (* If[PrintResult, 
   Print["The main chord lengths before flattening are :" <> TextString[ChordLengths]]]; *)
   {{constraints, values}, {OSorigin, OSbase}} = solutionspace;
   (*Print["After refining, the origin is at: "<>ToString[OSorigin]];*)
   PreflatChords = chords; PreflatChordLengths = ChordLengths;
   PreflatPeris = PeriPoints; PreflatSSK = solutionspace;];
  maxlo = 2.*InscribedSphere[[1]];
  progresslabel = " Main chord search completed. "; 
  Print[Style[progresslabel, Blue, TextAlignment -> Center]];
  progresscounter = 0;
  {{constraints, values}, {OSorigin, OSbase}} = solutionspace;
  poss = Count[ChordLengths, _?(Positive[#] &)]; 
  negs = Count[ChordLengths, _?(Negative[#] &)];
  If[negs > 0, 
   AppendTo[Processreport, {"Only the first " <> ToString[poss] <> 
      " chords were calculated by LP for the SSK, and 
   the remaining " <> ToString[negs] <> " are approximated by diameter values. "}], 
   AppendTo[Processreport, {"All " <> ToString[poss] <> 
      " chords were calculated by LP for the SSK."}]];
  
  	(* 2. THE FLATTENING LOOP *)

   (* Print["Starting the flattening loop, the constraint vectors span "<>ToString[MatrixRank[constraints, Tolerance -> 0.0001]]<>
   	" dimensions for "<>ToString[vars]<>" variables."]; *)
  	
  While[flatten, If[PrintResult,Print["Starting a flattening round number "<>ToString[++flatcount]]];
  	
   			(* i. Decide if any short chords need flattening out. *)
   (*Print["Unflattened chords in this round are: \n",chords];*)
   {keepers, neglect, maxaspect} = Decider[maxaspect, negligible, maxlo, plotit, PrintResult];
   plotit = False; (* Only display the initial decision plot *)
   thins = vars - keepers;
   flatten = thins > 0;
   If[!flatten, Print[" No chords qualified to be flattened out. "]];
(*    Print["Check PeriPoints are interior in FlatnCentre, before flattening: ",
   outside=Chop@Min@Map[(values-constraints.#)&,PeriPoints];
   If[outside\[GreaterEqual]-tol,True,"Outside by margin "<>
   TextString[Abs@outside]]];  *)
   
   			(* ii. Flattening transform, specified relative to current, unflattened space.
   				Recenter using chords, and take chord directions as basis vectors.
   				When flatten=False,this is just an affine transformation.*)
   (*Print[{keepers,Dimensions[PeriPoints],Dimensions[chords]}];*)
   chords = chords[[;; keepers]];
   chordvecs = Map[Subtract @@ # &, chords];
   (*A tricky point -
   the chords and hence chordvecs are supposed to be orthogonal by construction.
   But in a case with extreme aspect ratios,
   the length discrepancy between biggest and smallest chordvecs 
	causes numerical errors if they are just individually normalized. 
   So it is better to do orthogonalization of the whole set instead.
   But since chords were constructed to be orthogonal within tolerance LPtol, 
   the orthogonalization tolerance here needs to be siginficantly smaller
   otherwise it can produce zero vectors i.e. give a rank deficient basis. *)
   FlatBasis = Orthogonalize[chordvecs, Method -> "Householder", Tolerance -> 0.01*LPtol];
   flatrank = Count[Norm/@FlatBasis,_?(# > 0 &)];
   If[flatrank < keepers, 
   Print["The chord set does not produce a full rank basis for the flattened space.
   This is likely caused by an extreme aspect ratio, try flattening more directions. ";Abort[]]];

   (* FlatBasis determines the orientation of the flattened hyperplane.
    To fix its location, need to specify a single point in it; do that
     by choosing the centroid of the chord endpoints, as the coordinate center for now.
     The exception is a simple cone; there, the capping at artificial values means
     that chord endpoints are also artificial; the apex of the cone is the only "real"
     SSK point so use that to fix the location and ensure the apex stays feasible. *)
   
   (* apex is a point in RSS; apexpoint is its projection in SSK coordinates. *)
   flatcenter = If[flatten, 
   		If[SStype=="SimpleCone", apexpoint=DowncastPoint[apex, {OSorigin, OSbase}],
   			 Mean@Flatten[chords, 1]], 
   		ConstantArray[0., Length[OSbase]]
   ];

   (* Print["Constraints will be downcast to the hyperplane passing through "<>
   ToString[flatcenter] <> " as specified in the preflattened coordinates."]; *)
      
   (*Find the thin directions and transfer them to the embedding space.*)
   If[flatten, 
   	 thindirs = Join[thindirs, Chop[NullSpace[FlatBasis, Tolerance -> LPtol].OSbase, tol]]];
   (*Print["Before flattening, there are "<>ToString@Length[thindirs]<>" thin directions"];
   Print["The Flatbasis orthogonality matrix is: \n",MatrixForm@Chop[FlatBasis.Transpose[FlatBasis]]];*)
   FlattenTransform = {flatcenter, FlatBasis};
   OSorigin = OSorigin + flatcenter.OSbase;
   OSbase = FlatBasis.OSbase;
   
   			(* iii. Downcast the constraints, correct the centre, and update SS *)
   {constraints, values} = 
    Chop@N@DowncastConstraints[constraints, values, FlattenTransform];
(*   Print["After downcasting, the constraint vectors span "<>ToString[MatrixRank[constraints, Tolerance -> 0.0001]]<>
   	" dimensions for "<>ToString[keepers]<>" variables."]; *)
    
    (* Print["After downcasting, values are ", values]; *)
    (* Both choices above for the flattening centre can sometimes be marginally exterior.
    Correct this, as both RedundancyTrimmer and MiniMaxcentre rely on an interior origin.*)
    centershift = Retrieve[{{constraints, values}, FlattenTransform}, ConstantArray[0., Length[FlatBasis]], PrintResult];
    flatcenter=flatcenter + centershift.FlatBasis;
    values = values - constraints.centershift;
    OSorigin = OSorigin + centershift.OSbase;
   
   If[flatten,
   	progressrange = {0, Length@constraints};
    progresslabel = " Eliminating constraints that are redundant after flattening. ";
    Print[Style[progresslabel, Blue, TextAlignment -> Center]];
    {constraints, values, redundancies} = RedundancyTrimmer[constraints, values, {}, PrintResult];
    {cons, vars} = Dimensions[constraints];
    progresslabel = " Flattened redundant constraint elimination completed.";
    progresscounter = 0;
    ];

   (* After flattening, the inscribed center is generally a good choice for the origin,
    unless the inscribed radius is zero, in which case just leave it where it is.
    Without flattening, calling MiniMaxcentre just updates its specification to the 
    rotated OSbase vectors.
 *)
   
   inscribedcenter = MiniMaxcentre[constraints, values, PrintResult];
   (* Print[Chop@InscribedSphere]; *)
   If[flatten,
    centershift = If[InscribedSphere[[1]] > tol, 
    	InscribedSphere[[2]] = ConstantArray[0., vars]; inscribedcenter,
    	ConstantArray[0., vars]];
    flatcenter=flatcenter + centershift.FlatBasis;
    values = values - constraints.centershift;
    OSorigin = OSorigin + centershift.OSbase;
    ];
   FlattenTransform = {flatcenter, FlatBasis};
   solutionspace = {{constraints, values}, {OSorigin, OSbase}};
   (* {{TestConstraints, TestValues}, {TestOrigin, TestBasis}}=solutionspace; *)
   
   			(* iv. Exit if not flattened, else find new chords and peripoints and test. *)
   
   If[! flatten,
    (*Note that because PeriPoints vectors are in terms of the old basis,not the updated solutionspace,
    it is FlattenTransform that must be used rather than element 2 of the current solutionspace*)
    PeriPoints = DowncastPoint[PeriPoints, FlattenTransform];
    If[PrintResult, 
     Print["Check PeriPoints are interior in FlatnCentre, after reorienting the basis but without flattening: ",
    outside=Chop@Min@Map[(values-constraints.#)&,PeriPoints];
    If[outside\[GreaterEqual]-tol,True,"Outside by margin "<>
    TextString[Abs@outside]]]]; 
    Break[]
    ];
   
   getmargin = True;
   AppendTo[Kerneltable, {"Flatten out " <> ToString[thins] <> " thicknesses " <>
       TextString[neglect] <> " and below", cons, vars, 0}];
   progressrange = {0, vars}; 
   progresslabel = " After flattening: ";
   chords = MainChords[solutionspace, chordmax, chordmin, flipmax, PrintResult];
   (*Print[{"After chord-based recentering, the flattened SS becomes ",solutionspace}];*)
   {{constraints, values}, {OSorigin, OSbase}} = solutionspace;
(*
apexpoint=DowncastPoint[apex, {OSorigin, OSbase}];
outside=Chop@Min[values-constraints.apexpoint];
Print["Check apex is interior after flattened mainchords: "<>
   If[outside\[GreaterEqual]-tol,"True","Outside by margin "<>
   TextString[Abs@outside]]]; 
*)   
   progresslabel = " Flattened chord search completed. "; 
   progresscounter = 0;
   diametertest = 2*InscribedSphere[[1]] <= negligible;
   chordtest = Min[Abs@ChordLengths] <= negligible;
   (*Print[{"neglect,diametertest,chordtest",neglect,diametertest, chordtest}];*)
   Switch[{diametertest, chordtest},
    {True, True}, flatten = True,
    {True, False}, flatten = False;
    Print[ Style[" CAUTION: The maximal inscribed sphere diameter " <> 
       TextString[2*InscribedSphere[[1]]] <> " suggests that further flattening is possible.
 But no chords shorter than the max thinnning allowed were detected by sampling. 
 Manual repeat of flattening or even earlier stages may be needed to improve this.",
       	 Darker@Green]],
    {False, False}, flatten = False,
    {False, True}, flatten = False; 
    Print[Style["The post flatttening inscribed sphere diameter " <> 
       TextString[InscribedSphere[[1]]] <> " is larger than the 
       allowed flattening " <> TextString[negligible] <> ". 
       Thus flattening was successful, even though some putative shorter chords lengths remain. 
       This reflects the fact that sampled diameters are lower limits of maximal chord lengths.
        To obtain consistency, a larger value may have to be chosen for maxthin.", Darker@Green]]
    ]
   ]; 						(* End while flatten *)
   (* If[PrintResult, Print["The main chord lengths after flattening are :" <> 
      TextString[ChordLengths]]]; *)
   

  
  	(* 3. CALCULATE FLATTENING TRUNCATION MARGINS *)
  (* Preflat chord endpoints are used as a representative sample of points before flattening. 
  Estimate the margin by which flattening truncates the solution space,
  	from the residue after deconstructing these chord endpoints, as located in RSS. *)
  
  margin = If[ getmargin,
    progresslabel = " Calculating deconstruction residue at chord endpoints, introduced by flattening ";
	Print[Style[progresslabel, Blue, TextAlignment -> Center]];
    endpoints = Flatten[PreflatChords, 1];
     norms=Norm/@endpoints; (* Print["The endpoint norms are: ",norms]; *)
    endpoints=UpliftPoint[endpoints,PreflatSSK[[2]]];
   
    progressrange = {0, Length@endpoints}; progresscounter=1;
    residues = Reap[TimeConstrained[
       Do[Sow@Deconstructor[endpoints[[i]], ReducedSS, solutionspace];
        progresscounter++; reached = i, {i, Length@endpoints}], 
       timeconstraint]];
    residues = If[reached == 0, 
      warning = warning <> "Flattening margin could not be calculated within the " <> 
        ToString[timeconstraint] <> " sec time allowed."; {0.}, residues[[2, 1]]];
    progresslabel = " Flattening margin calculation completed.";
	Print[Style[progresslabel, Blue, TextAlignment -> Center]];
    progresscounter = 0;
    (* Print["The endpoint residues are: ",residues]; *)
    norms=norms[[;;reached]];
    residues =100.*Select[residues/norms, NumberQ];
    If[PrintResult, 
     Print["Percentage errors due to flattening at the chord endpoints, calculated within the " <> 
     	ToString[timeconstraint] <> " sec time allowed: " <> TextString[residues]]];
    If[reached > 0, Mean@residues, 0.],
    0.];
  
  	(* 4. UPDATE THE PROGRESS REPORT *)
  
  (* Print["Check PeriPoints are interior in FlatnCentre, after flattening: ",
  outside=Chop@Min@Map[(values-constraints.#)&,PeriPoints];
  If[outside\[GreaterEqual]-tol,True,"Outside by margin "<>TextString[Abs@outside]]]; *)
  poss = Count[ChordLengths, _?(Positive[#] &)]; 
  negs = Count[ChordLengths, _?(Negative[#] &)];
  
  If[getmargin, 
     AppendTo[Processreport, {"The flattening error for points on the Kernelspace margins
 is estimated to be " <> 
       TextString[ NumberForm[margin, 2, ScientificNotationThreshold -> {-2, 3}]] <> 
       "% based on " <> ToString[reached] <> " chord endpoints. "}];
   If[margin > 10, 
    warning = "The estimated flattening margin (over sampled chord endpoints) 
    exceeds 10% of the mean chord length. Inspect validation test and
    reduce maxthin or maxaspect if this is not acceptable.\n" <> warning];
   If[negs > 0, 
    AppendTo[Processreport, {"After flattening, only the first " <> 
       ToString[poss] <> " chords were calculated by LP and \n   the remaining " <> 
       ToString[negs] <> " are approximated by sampled diameter values. "}], 
    AppendTo[Processreport, {"After flattening, all " <> ToString[poss] <> 
       " chords were calculated by LP. "}]];
   ];
  
  If[warning != "", 
   Processreport = Join[Processreport, {{warn}, {warning}}]];
  thindirs]
  
Decider[maxaspect_, negligible_, maxlolimit_, plotit_,  PrintResult_: False] :=
 (* This function decides how many of the chord (arranged in descending order of length) 
 are to be kept while the rest are flattened away. 
 It also optionally produces a plot to visualize the decision implied \
by the parameters maxaspect and negligible.
 Any "thin" diameters that satisfy either of the following criteria,are omitted,
 provided that they are not larger than "negligible":
 i) small enough to give a polytope projection aspect ratio larger than the maxaspect 
 ii) or that drops markedly below an hyperbolic decrease of the "thick" diameters.
 It returns this number as well as the actual maximal thickness that is flattened 
 out by this decision. *)
 Module[{expect, neglect, diameters, aspects, aspectmax, accept, 
   require, keepers, rejects, thicks, thins, counthicks,
   plot1, plot2, aspectmin, cutoff, colorpoints, tol = 0.000001},
  (*To detect a thick-thin gap,
  hyperbolic extrapolation of the last few diameters in the thick group is used below*)
  expect[thics_, n_] := 
   Block[{x, lastfew}, 
    lastfew = Min[4, Ceiling[(1 + Length[thics])/2]];
    If[lastfew == 1, First@thics, 
     Fit[thics[[-lastfew ;;]], {1, 1/x}, x] /. x -> n - Length[thics] + lastfew]];
  
  aspectmax = Max[maxaspect, 1];
  (*Below, diameters are prevented from being smaller than the tolerance tol,
  to avoid numerical difficulties.
  But then, any diameter at this level is spurious and needs to be flattened out.*)
  neglect = Max[negligible, 1.1*tol];
  diameters = Map[Max[#, tol] &, Abs@ChordLengths];
  If[negligible > tol, (* Keep all if negiligible = 0 *)
  (*The following associates with each diameter, 
  the mean value of aspect ratios with all larger diameters.
  That avoidss extreme aspect ratios caused by e.g.the first diameter 
	being exceptionally large as sometimes observed *)
  aspects = Prepend[Table[
     Mean[diameters[[;; i - 1]]]/diameters[[i]], {i, 2, Length[diameters]}], 1];
  (*Limit the diameters to those giving an acceptable aspect ratio,
  while making sure that no diameter larger than the neglect spec is omitted*)
  accept = Position[aspects, _?(# <= aspectmax &)];
  require = Max[1, Position[diameters, _?(# > neglect &)]];
  keepers = Max[accept, require];
  rejects = Length@diameters - keepers;
  (* Print[{"Diameter count, required no, acceptable aspects, keepers", 
  Length@diameters,require,accept,keepers}]; *)
  If[PrintResult && rejects > 0, 
   Print[ToString[rejects] <> " diameters " <> TextString[diameters[[keepers + 1]]] <> 
     " and thinner are dropped to prevent aspect ratios higher than " <> 
     TextString[aspects[[keepers + 1]]]]];
  (*Check if there are diameters significantly thinner than the rest.If so,
  drop any of them that have passed the aspect ratio test above.This 
test is only applied if at least 4 diameters remain,
  otherwise clustering is hardly sensible and it makes more sense to 
avoid approximation.*)

 (* Print[{"keepers,rejects,require",keepers, rejects,require}]; *) 
  If[keepers > 3 && keepers > 1 + require,
  	(*Print[{"diameters", diameters}];*)
  	{thicks, thins} = FindClusters[diameters, 2, DistanceFunction -> (Abs[#1 - #2] &)];
   counthicks = Length@thicks;
   (*Print[{thicks,thins,expect[thicks,1+counthicks]}];*)
   If[Length@thicks > 2 && expect[thicks, 1 + counthicks] > 5*Max[thins], 
    counthicks = Max[counthicks, require];
    keepers = Min[keepers, counthicks];
    (*avoid violating aspect ratio test*)
    rejects = Length@diameters - keepers - rejects;
    If[PrintResult && rejects > 0, 
     Print["Thick-thin gap detected - dropping " <> 
        ToString[rejects] <> " diameters thinner than " <> 
        ToString[diameters[[keepers + 1]]]];]
  ]];
  neglect = If[keepers < Length@diameters,
    If[ChordLengths[[keepers]] > 0 && ChordLengths[[keepers + 1]] < 0,
      Print[Style["\nFlattening threshold coincides with transition from chords to diameters and may be spurious!
Test this by increasing chordmin, or avoid it by decreasing maxaspect or maxthin. \n", Red]]
	];
    diameters[[keepers + 1]]
    , 0.],
    keepers = Length@ChordLengths];
(* Print["Diameters are: ",diameters]; *)
If[Length[diameters]==1, diameters=Join[diameters,diameters]];
  If[plotit,
   (*Display the diameters, flattening criteria and decision for user info*)
   aspectmin = If[keepers > 1,
   	 Mean[diameters[[;; keepers - 1]]]/aspectmax, 
     diameters[[1]]/aspectmax
     ];
   cutoff = Min[negligible, aspectmin];
   colorpoints = Map[Tooltip[
       Which[Abs[#] == 0, Style @@ {Abs[0.0001], Darker[Green, 0.5]}, 
        Abs[#] >= cutoff && # > 0, 
        Style @@ {Abs[#], Darker[Green, 0.5]}, 
        Abs[#] >= cutoff && # < 0, Style @@ {Abs[#], Blue}, 
        Abs[#] < cutoff && # < 0, Style @@ {Abs[#], Purple}, 
        Abs[#] < cutoff && # > 0, Style @@ {Abs[#], Red}], 
       ToString[Abs[#]]] &, ChordLengths];
   plot1 = ListLogPlot[colorpoints, PlotStyle -> PointSize[Large], 
     PlotLegends -> Placed[{"Green/Blue - Kept chord/diameter;\nRed/Purple - \
Flattened chord/diameter."}, Below], PlotRange -> All];
   plot2 = LogPlot[{aspectmin, Min[negligible, 10*Max[diameters]], maxlolimit},
   	 {x, 1, Length[diameters]}, 
     Filling -> {1 -> {Axis, Opacity[0.2, Magenta]}, 2 -> {Axis, Opacity[0.2, Yellow]}}, 
     PlotRange -> {0.5 Min[diameters], 2*Max[diameters]}, 
     PlotLegends -> Placed[{"Aspect ratio limit", "Max negligible", "Inscribed diameter"}, Above]];
   flatplot = Show[plot2, plot1, Frame -> True];
   ];
  
  {keepers, neglect, aspectmax}]
  
 CoverageInterval[solutionspace_, peripoints_, PrintResult_:False] := 
 Module[{constraints, values, OSorigin, OSbase, dims, cons, vars, 
   equalvals, direction, samplespan, sampleprobs, samplefracs, 
   samplecounts, rads, Targets, radii, PointCols, Objective, Coefs, 
   CoefRange, axrads, dircount = 0, count = 0, incount = 0, 
   conf = 0.05, alfa, beta, mean, sdev, hiconf, loconf, wide = 1, 
   minwide = 0.05},
  {{constraints, values}, {OSorigin, OSbase}} = solutionspace;
  dims = Length[OSbase];
  PointCols = Transpose[peripoints];
  {cons, vars} = Dimensions[PointCols];
  Objective = ConstantArray[1, vars];
  CoefRange = ConstantArray[{0, Infinity}, vars];
  (* Find SS radius along axis directions *)
  axrads = Table[axrads = Radii[constraints, values, UnitVector[dims, i]], {i, dims}];
  (*Print[axrads];*)
  samplespan = Median@Abs[axrads];
  {directs, samplepoints} = Reap[
     While[wide > minwide, dircount++; progresscounter=dircount;
      (* Get a pair of random points, within SS. They are random fractions 
      of each distance to the SS boundary, forwards and backwards along a 
      chosen direction. Fully random direction choice is given by:*)
       (*direction=Normalize[RandomReal[{-1,1},Length[OSbase]]];*)
      (*But choosing the components of the vector randomly from an 
      interval equal to the axis radius, aspect ratios of the SS are taken into account
      to give a preference for directions along the long axes. 
      This is equivalent to rescaling all aspect ratios to 1.   *)
      direction = Normalize[Table[RandomReal[axrads[[i]]], {i, Length[OSbase]}]];
      Sow[direction, dirs];
      (*Print[direction];*)
      radii = Radii[constraints, values, direction];
      (* Increase the number of points proportionately to the length of 
		the radius to distribute samples more uniformly *)
      sampleprobs = (Abs@(radii/samplespan));
      samplefracs = sampleprobs - Floor@sampleprobs;
      samplecounts = Floor@sampleprobs + Map[If[RandomReal[] < #, 1, 0] &, samplefracs];
      (*Print["For direction "<>ToString[dircount]<>" I am sampling "<>
      ToString[samplecounts]<>" forwards and backwards points respectively"];*)
      Do[
       (* The random fraction is corrected for radial increase of volume element *)
       rads = radii[[j]]*Map[(#)^(1/cons) &, RandomReal[1, samplecounts[[j]]]];
       Targets = Map[# *direction &, rads];
       (* Decompose each target point as a minimal superposition of the supplied 
       periphery vectors and check if it falls within their convex hull *)
       Do[Sow[Targets[[i]], samplepts]; count++;
        equalvals = Transpose[{Targets[[i]], ConstantArray[0, cons]}];
        Coefs = 
         LinearProgramming[Objective, PointCols, equalvals, CoefRange, Method -> "Simplex"];
        If[Total[Coefs] <= 1, incount++], {i, samplecounts[[j]]}];
       , {j, 2}];
      (* Get width of Jeffreys 95% confidence interval for incount successes 
      in count Bernoulli trials. This interval gives good, symmetric coverage. *)
      (* Print[{"dircount, count, incount",dircount,count,incount}]; *)
      
      alfa = incount + 0.5; beta = count - incount + 0.5;
      (* If either alfa or beta becomes large, approximate Beta distribution by Normal dist. 
      This avoids numerical problems with Beta and is also faster.*)
      mean = alfa/(alfa + beta);
      sdev = Sqrt[(alfa * beta)/((alfa + beta)^2 (alfa + beta + 1))];
      {hiconf, loconf} = If[alfa + beta < 50,
        {Quantile[BetaDistribution[alfa, beta], 1.-conf/2], 
         Quantile[BetaDistribution[alfa, beta], conf/2]}, {Quantile[
          NormalDistribution[mean, sdev], 1.-conf/2], 
         Quantile[NormalDistribution[mean, sdev], conf/2]}];
      (*Print[{loconf,hiconf}];*)
      wide = hiconf - loconf;
     (*  If[dircount > 10 || count > 100, Break[]];       (* Safety valve *) *)
      If[dircount > 1000 || count > 10000, Break[]]       (* Safety valve *)
      ]][[2]];
  If[PrintResult,Print[ToString[count]<>" points were sampled to estimate coverage, and "<>
  ToString[count - incount]<>" were found outside."]];
  
  Round[100. {Max[0., loconf], Min[1., hiconf]}]
  ] 
  
 WrappingRadii[solutionspace_, peripoints_, PrintResult_:False] :=
 (* This function calculates the inscribed radius of a simplex, that just encloses 
 the set of periphery points (the minor simplex), and for a similar simplex with the same 
 orientation that just encloses the complete solution space (the major simplex). 
 	
 	By construction, the orientation of the sides are those of a regular simplex, so the vertex
 	hyperangles for all vertices are equal and so the simplex is a regular simplex.
 	Nevertheless the calculated side displacements hi are unequal, which means that 
 	the origin is not at the center of the inscribed hypersphere. 
 	The true inscribed radius is simply the mean of the side displacements.
 	This follows because any interior point partitions the volume of the simplex into 
 	constituent (non-regular) simplices erected on each side of the original simplex, and
 	their respective volumes are proportional to their heights, which are the distinct hi values.    
    
    It repeats this calculation for several randomly chosen simplex orientations, and returns 
 the mean value of each of these as well as their ratio. 
 	The radius ratio is an indication of the extent to which the periphery points cover 
 the solutionspace, in a way that can be compared between different dimension counts. *)
 Module[{constraints, values, OSorigin, OSbase, cons, vars, 
   SimplexSides, periprojections, target=10, reached=0, simplexradii, MinorSimplexRadii, 
   MajorSimplexRadii, minormean, majormean, rot, minors, majors, 
   XRange, XValues, tol = 0.001},
   
  {{constraints, values}, {OSorigin, OSbase}} = solutionspace;
  {cons, vars} = Dimensions[constraints];

  (* For a 1-dim space, the 2 periphery points lie at SSK vertices so 
  both minor and major reduce to the SSK itself.  *)
  If[vars < 2, Return[{Max[values], Max[values]}]];

  XRange = ConstantArray[{-Infinity, Infinity}, vars]; 
  XValues = Map[{#, -1} &, values];
  
  (*Determine enclosing simplexes and the mean perp radius to its \
sides,a minor one for the periphery points and a major one for the \
complete SSK*)
  SimplexSides = RegSimplexDirections[vars];
  progresscounter=0;
  simplexradii=Reap[TimeConstrained[
  	Do[progresscounter++;
       periprojections = peripoints.Transpose@SimplexSides; 
       MinorSimplexRadii = Max /@ Transpose[periprojections];
       MajorSimplexRadii = Map[#.Quiet@
            LinearProgramming[-#, constraints, XValues, XRange, 
             Method -> "Simplex", Tolerance -> tol] &, SimplexSides];
       minormean = Mean[MinorSimplexRadii]; 
       majormean = Mean[MajorSimplexRadii];
       Sow[{minormean, majormean}];
       (*Randomly reorient the simplex orientation for the next iteration.*)
       rot = RotationMatrix[{UnitVector[vars, 1], RandomReal[{0., 1.}, vars]}]; 
       SimplexSides = SimplexSides.rot; reached = i, {i,target}],
        timeconstraint]];
 (* Only a maximum of target = 10 reorientations were done here, independently of the dimensions. 
 A number proportional to the dimensions, such as target = 5*vars would make sense, but 
 becomes too slow. 
 In practice the mean radius values are fairly similar in different
 orientations and in particular the ratio is not significantly improved by an increase. *)
 
 simplexradii=Which[reached == 0, 
 	Print[Style[" Enclosing simplex could not be calculated in allowed time,
 	 so no coverage estimate is made.",Darker@Green]];{{0.,0.}},
 	 reached < target, Print["Due to time constraint, only "<>ToString[reached]<>
 	 	" reorientations of enclosing simplex were made to estimate coverage."];simplexradii[[2, 1]],
 	 	True,simplexradii[[2, 1]]];
  
 {minors, majors} = Transpose@simplexradii;       
 If[PrintResult && reached >0,
 	Print["The inscribed radius found for a simplex that encloses the periphery points and the complete SSK
 respectively, are as follows for the trial orientations: "]; 
 	Print[{minors,majors}];
  	Print[{"The radius ratios are ",minors/majors}]
  	]; 
  	
  {minors, majors} = Mean /@ {minors, majors}
  ]

End[] 

(* End Private Context *)

EndPackage[]
```
