## Supplementary material for "The FBA Solution Space Kernel – Introduction and Illustrative Examples": SSK software manual and code: Centering.m

```
(* ::Package:: *)

(* COPYRIGHT
					\[Copyright] Copyright 2022 Wynand Verwoerd

This file is part of SSKernel.

The SSKernel program is free software; you can redistribute it and/or modify it under the terms of the GNU General Public License as published by the Free Software Foundation; either version 3 of the License, or (at your option) any later version.
 
This program is distributed in the hope that it will be useful, but WITHOUT ANY WARRANTY; without even the implied warranty of MERCHANTABILITY or FITNESS FOR A PARTICULAR PURPOSE . See the GNU General Public License for more details.
 
You should have received a copy of the GNU General Public License along with this program . If not, see http://www.gnu.org/licenses/
*)

(* Wolfram Language Package *)

BeginPackage["Centering`",{"Configuration`","RayFinding`"}]
(* Exported symbols added here with SymbolName::usage *)  
UpliftPoint::usage="Transform a point from low to high dimension space."
DowncastPoint::usage="Transform a point from high to low dimension space."
CombineTransform::usage="Combine two consecutive transforms into one transform "
DowncastConstraints::usage="Project the constraint set to a lower dimensional subspace."
UpliftConstraints::usage="Express a subspace constraint set as higher dimensional vectors."
SolutionTest::usage="Tests if a given flux or its projection belongs to the solution space."

CLDcentre::usage="Constrained centering by minimizing sum of Linear Distances (facet capable)"
MiniMaxcentre::usage="Find centre and radius of maximal inscribed hypersphere. "
CLODcentre::usage="Constrained centering by minimizing sum of Differences or Ordered Linear distances "
LSDcentre::usage="Center by minmising L2-norm i.e. Least Square Distances using pseudoinverse"
LSDDcentre::usage="Center by minmising L2-norm of differences i.e. Least Square using pseudoinverse"
MainChords::usage="LP calculation of approximate, maximal length, progressively orthogonal chords for polytope "
CenterChords::usage=" Supplements an incomplete LP chord set by sampled approximations. "
Radii::usage="Finds diametrically opposed pair of radii along given vector "
RegSimplexDirections::usage="Finds directions to the sides of a regular simplex "
RefineCentre::usage="Finds a mutually consistent set of peripheral points and their centroid "
Retrieve::usage=" Corrects a point that is marginally infeasible "
Centrality::usage=" Displays a bar plot of diameter pairs"
ChordPairs::usage=" Extracts the indices in the peripehery point list of the chord endpoint pairs"

deviation

Begin["`Private`"] (* Begin Private Context *) 

(* PROJECTION TO/FROM SUBSPACES *)

(* NOTE - Uplift and Downcast configured to operate on Mathermatica convention 
	that use row vectors, while algebraic equations for flux space 
	use column vectors *)

UpliftPoint[Lopoint_, transform_] := 
 Module[{pointset}, 
  pointset = If[VectorQ[Lopoint], {Lopoint}, Lopoint]; 
  pointset = Map[Chop[# . transform[[2]] + transform[[1]], 0.0001] &, pointset];
   If[VectorQ[Lopoint], pointset[[1]], pointset]
  ]
DowncastPoint[Hipoint_, transform_] := 
 Module[{pointset}, 
  pointset = If[VectorQ[Hipoint], {Hipoint}, Hipoint]; 
  pointset = 
   Map[Chop[(# - transform[[1]]) . Transpose[transform[[2]]], 0.0001] &, pointset]; 
    If[VectorQ[Hipoint], pointset[[1]], pointset]
  ]

CombineTransform[HighToMedium_,MediumToLow_]:=Block[{hidim,middim,lodim,HMdims,MLdims},
hidim=Length[HighToMedium[[1]]];
middim=Length[MediumToLow[[1]]];
HMdims=Dimensions[HighToMedium[[2]]];
MLdims=Dimensions[MediumToLow[[2]]];
lodim=MLdims[[1]];
If[MLdims=={lodim,middim} && HMdims=={middim,hidim},
{HighToMedium[[1]]+MediumToLow[[1]] . HighToMedium[[2]],
	MediumToLow[[2]] . HighToMedium[[2]]},
Print["Dimensions of transforms are incompatible!"];{}]
]

DowncastConstraints[convecs_, convals_, transform_] := 
 Module[{bt, norms, keepers, projects, newvecs, newvals, 
   tol = 0.001},
   (* Constraint vectors with projection on the subspace less than tol, are omitted.
   	Such a vector generally produces a constraint hyperplane cutoff on the subspace, at 
   	a distance (conval/projection). If the projection is zero, the intercept is at Infinity,
   	and so for small projections its value is very sensitive to numerical error, but
   	normally falls beyond the cutoffs of other constraints.
   	That means it becomes redundant and would anyway be eliminated by redundancy trimming
   	performed after the downcast, so it is more efficient to eliminate it here.

   	More seriously, in cases where the corresponding conval values is small (i.e., the 
   	coordinate origin is on or near the polytope periphery) the cutoff it produces can be 
   	within the range of other constraint hyperplanes, but spurious because of numerical
   	inaccuracy in the direction of the vector. Generally, the accuracy of a downcast/uplift 
   	cycle is found to be about 10-5 to 10^-6. So choose tol well bigger than this, e.g. 10^ -3
   	so constraints that are kept have reliable cutoffs. 
   	
   	Conceivably, this may cause the downcasted polytope to be open since it misses some constraint,
   	but such cases have not be observed yet. The alternative is worse, cases have arisen where 
   	a downcasted constraint set excluded the current interior origin or became completely infeasible. 
   *)
  bt = Transpose[transform[[2]]];
  projects = convecs . bt; norms = Norm /@ projects;
  keepers = Map[If[# > tol, True, False] &, norms];
  (* Print[{"Keepers",keepers,"Norms",norms}]; *)
  norms = Pick[norms, keepers];
  newvecs = Pick[projects, keepers]/norms;
  newvals=Pick[convals, keepers];
  (*
  Print[{"newvals",newvals}];
  Print[{"Dot",Pick[convecs, keepers].transform[[1]]}];
  *)
  newvals = Chop[(newvals - Pick[convecs, keepers] . transform[[1]])/norms, 0.0001];
  {newvecs, newvals}]


UpliftConstraints[convecs_, convals_, transform_] := 
 Module[{projects, newvals},
  projects = convecs . transform[[2]];
  newvals = convals + projects . transform[[1]];
  {projects, Chop[newvals,0.0001]}
  ]

SolutionTest[flux_, solutionspace_, tol_: 0.001] := 
 Module[{osflux, slacks, satisfy, truncate},
  osflux = DowncastPoint[flux, solutionspace[[2]]];
  slacks = 
   Chop[solutionspace[[1, 2]] - solutionspace[[1, 1]] . osflux, tol];
  (*Print[slacks];*)satisfy = Min[slacks] > -tol;
  truncate = 
   Norm[UpliftPoint[osflux, solutionspace[[2]]] - flux] > tol;
  Which[satisfy && ! truncate, "Full vector in solution space", 
   satisfy, "Projected vector in solution space", True, 
   "Vector violates constraints by " <> 
    TextString[Min[0, #] & /@ slacks]]]

(* FINDING POLYTOPE CENTERS *)
 
CLDcentre[constraints_, values_, facetcols_: {}] := 
Module[{SXConstraints, SObjective, SValues, SX, SXRange, cons, vars, 
   conslack, slackcount, center, tol = 0.00001},
  {cons, vars} = Dimensions[constraints];
  
  (* Try LSD centering first *)
  center = LSDcentre[constraints,values];
  If[Min[values - constraints . center]>= 0.,
  	Return[center], Print["LSD centre infeasible, use CLD to enforce constraints. "]];
  (* But if this turns out to be infeasible, enforce the constraints by LP *)

  conslack = Complement[Range[cons], facetcols];
  slackcount = Length[conslack];
  SValues = Table[{values[[i]], 0}, {i, cons}];
  SXConstraints = 
   Join[IdentityMatrix[cons][[All, conslack]], constraints, 2];
  SObjective = 
   Join[ConstantArray[1, slackcount], ConstantArray[0, vars]];
  SXRange = 
   Join[ConstantArray[{0., Infinity}, slackcount], 
    ConstantArray[{-Infinity, Infinity}, vars]];
  SX = Quiet[
    Check[LinearProgramming[SObjective, SXConstraints, SValues, 
      SXRange, Method -> "Simplex", Tolerance -> tol], 
     "No Solution", {LinearProgramming::lpsnf}],{LinearProgramming::lpsnf}];
  center = If[VectorQ[SX], Drop[SX, slackcount], 
   Print["CLDcentre failed - constraints are incompatible and do not \
define a feasible region."]; {}];
  center]

CLODcentre[A_, b_] := Module[
  {Dmat, OffDiagMinimum, weights, overlaps, w8laps, choice, 
   chosen, pair, pairs, pairlist, isodd = True, newseq, AA, Aall, bb, 
   ball, c, x, lb, m, n , sub, M, DA, Db},
  Dmat[n_] := 
   SparseArray[{Band[{1, 1}] -> 1, Band[{1, 2}] -> -1}, {n - 1, n}];
  (* Note that only the offdiagonal minimum should be found to \
guarantee that a proper opposing constraint pair is produced *)
  OffDiagMinimum[sqmat_] := 
   Module[{dim, updi, updipos}, dim = Length[sqmat]; 
    updi = Table[sqmat[[i, j]], {i, 1, dim - 1}, {j, i + 1, dim}]; 
    updipos = First@Position[updi, Min[updi]]; {updipos[[1]], 
     updipos[[1]] + updipos[[2]]}
    ];
  weights[n_, sub_] := N[sub (n - sub)*(4/n^2)];
  
  (* Reorder the constraints so opposing ones are paired and most \
significant pair differences get highest weights *)
  overlaps = A . Transpose[A];
  (*Print[MatrixForm@overlaps];Print[b];
  Print[MatrixForm@Outer[Plus,b,b]];*)
  w8laps = overlaps*Outer[Plus, b, b];
  (*Print[MatrixForm@w8laps];*)
  choice = Range[Length[overlaps]];
  pairs = Floor[Length[choice]/2];
  isodd = True;
  pairlist = Reap[Do[
      chosen = OffDiagMinimum[w8laps[[choice, choice]]];
      pair = Flatten@Map[choice[[#]] &, chosen];
      (*Print[pair];Print[w8laps[[pair[[1]],pair[[2]]]]];*)
      If[isodd, Sow[pair, odds], Sow[pair, evens]];
      isodd = ! isodd;
      choice = Delete[choice, Partition[chosen, 1]];
      , {pairs}]][[2]];
  (*newseq=Join[choice,Flatten@pairlist];*)
  (*Print[choice];*)
  If[Length[pairlist] == 1,(* Only 2 constraints, so no evens *)
   AppendTo[pairlist, {}]];(*Print[pairlist];*)
  newseq = Join[choice, Flatten@Reverse[pairlist[[2]]], 
    Flatten@pairlist[[1]]];
  (*Print[newseq];*)
  AA = A[[newseq]]; bb = b[[newseq]];
  
  {m, n} = Dimensions[AA]; M = m - 1;
  DA = Dmat[m] . AA;
  Db = Dmat[m] . bb;
  (*Print[{m,n,M}];*)
  Aall = SparseArray@Join[
     Join[IdentityMatrix[m], ConstantArray[0., {m, M}], AA, 2],
     Join[ConstantArray[0., {M, m}], IdentityMatrix[M], -DA, 2],
     Join[ConstantArray[0., {M, m}], IdentityMatrix[M], DA, 2]
     ];
  (*Print[MatrixForm[Aall]];*)
  ball = Join[
    Table[{bb[[i]], 0}, {i, m}],
    Table[{-Db[[i]], 1}, {i, M}],
    Table[{Db[[i]], 1}, {i, M}]
    ];
  (*Print[ball];*)
  c = Join[Table[0, {m}], Table[weights[m, sub], {sub, M}], 
    Table[0, {n}]];
  (*c=Join[Table[0,{m}],Table[1,{M}],Table[0,{n}]];*)
  lb = Join[ConstantArray[0., m], ConstantArray[-Infinity, M + n]];
  (*Print[Dimensions[Aall]];Print[Length/@{c,lb,ball}];*)
  (* Use Simplex - interior point method can be slow or not converge *)
  x = LinearProgramming[c, Aall, ball, lb, Tolerance -> 0.00001, 
    Method -> "Simplex"];
  x = If[VectorQ[x], Chop@Drop[x, m + M], 
    Print["CLODcentre failed - constraints are incompatible and do \
not define a feasible region."]; {}]
  ]
  
 
MiniMaxcentre[constraints_, values_, printresult_:False] :=
 (* This function finds the centre and radius of the maximal inscribed hypersphere. 
 It works by maximising the minimal perp boundary distance using LP.
 It still works for an unbounded prism, but for an unbounded cone \
there is no maximal radius and the centering fails too. *)
 Module[{XZConstraints, PolyConstraints, MinBConstraints, 
   ZeqConstraints, ZObjective, XZValues, XZ, XZRange, cons, vars, 
   tol = 0.00001},
  {cons, vars} = Dimensions[constraints];
  XZValues = Table[{values[[i]], -1}, {i, cons}];
  XZValues = Join[XZValues, XZValues, ConstantArray[{0., 0}, cons - 1]];
  ZObjective = Join[ConstantArray[0., vars], ConstantArray[1., cons]];
  PolyConstraints = Join[constraints, ConstantArray[0., {cons, cons}], 2];
  MinBConstraints = Join[constraints, IdentityMatrix[cons], 2];
  ZeqConstraints = Transpose@Join[ConstantArray[0., {vars, cons - 1}], 
     IdentityMatrix[cons - 1], {ConstantArray[-1, cons - 1]}];
  (*Print[Dimensions/@{PolyConstraints,MinBConstraints, ZeqConstraints}];*)
  XZConstraints = Join[PolyConstraints, MinBConstraints, ZeqConstraints];
  XZRange = Join[ConstantArray[{-Infinity, Infinity}, vars], 
    ConstantArray[{0, Infinity}, cons]];
  (*Print[Dimensions/@{ZObjective,XZConstraints,XZValues,XZRange}];*)
  
  XZ = Quiet[ Check[LinearProgramming[-ZObjective, XZConstraints, XZValues, 
      XZRange, Method -> "Simplex", Tolerance -> LPtol], 
     "Unbounded", {LinearProgramming::lpsub}], {LinearProgramming::lpsub}];
  (*Print[XZ];*)
  Which[
  	VectorQ[XZ],
   InscribedSphere[[1]] = Chop[Last@XZ,tol];
    If[printresult,Print["The inscribed radius is " <> ToString[InscribedSphere[[1]]]]];
   InscribedSphere[[2]]= Chop@Take[XZ, vars],
   StringQ[XZ], 
   Print[Style["MiniMaxcentre failed - the polytope is unbounded and does \
not have a finite inscribed hypersphere.",Red]];Abort[],
   True,
   Print["LP result is: "<>ToString[XZ]];
   Print[Style["MiniMaxcentre failed - constraints are incompatible and do not \
define a feasible region.",Red]]; Abort[]]
  ]  
  
(* The following centering functions are quick, but are not guaranteed 
	to give a point inside the polytope *)
  
LSDcentre[Amat_, bvec_] := Chop[PseudoInverse[Amat] . bvec]

LSDDcentre[Amat_, bvec_] := 
 Module[{Dmat, rows, distmat, distvec, centre}, 
  Dmat[n_] := 
   SparseArray@
    Block[{row = 0}, 
     Flatten@Table[
       row++; {{row, i} -> 1, {row, j} -> -1}, {i, n}, {j, i + 1, n}]];
  rows = Length[Amat]; distmat = Dmat[rows] . Amat; distvec = Dmat[rows] . bvec;
  centre = Chop[PseudoInverse[distmat] . distvec]
  ]
  
MaximalChords[solutionspace_] :=
  
  (* This function finds a set of maximal constraint polytope chords, 
  that are mutually orthogonal and progressively decreasing. 
  The coding below refers to diameters, but as they do not pass through a
  common centre the term chord is a more accurate description. *)
  
  Module[{constraints, values, origin, OSBase, cons, vars, Ymat, 
      XXZBcons, XXZBcons1, XXZBcons2, XXZBcons3, XXZBcons4, XXZBcons5, 
      XXZBcons6, orthocons = {}, XXZBvals, XXZBRange, XXZBDomain, 
      XXZBobjective, xxzb, peripoints = {},  perips, 
       MaxComponent, MM, overlap, Dist,  distable, v1, v2,  
   tol = 0.000001, itcount = 0},
    
    (* Function to calculate the distance from the origin to where \
two hyperplanes intersect *)
    Dist[v1_, v2_, dot_] := Block[{\[Theta]}, 
      	\[Theta] = ArcCos[dot]; 
        Sqrt[(v1^2 + v2^2 - 2 v1 v2 Cos[\[Theta]]) Csc[\[Theta]]^2]];
     
     {{constraints, values}, {origin, OSBase}} = solutionspace;
    {cons, vars} = Dimensions[constraints];
     
    If[vars == 1, peripoints = {DiagonalMatrix[values] . constraints}; 
      Return[peripoints]];
    (*Print[MatrixForm@constraints];Print[values];*)
    
    (* Estimate  an upper limit to SS radius *)
    distable = Table[overlap = constraints[[i]] . constraints[[j]]; 
    v1 = values[[i]];
    v2 = values[[j]]; 
        If[(v1 > tol || v2 > tol) && -1 + tol < overlap < 1 + tol, 
          Dist[v1, v2, overlap], 1.],
          {i, 2, cons}, {j, i - 1}];
    (*Print[distable];*)
     MM = 5. Max@distable; 
    Print[{"MM", MM}];
  
    (* Set up the extended constraints matrix to determine maximal \
Manhattan diameter *)
    XXZBDomain = 
   Join[ConstantArray[Reals, 3*vars], ConstantArray[Integers, vars]];
    (*XXZBDomain = ConstantArray[Reals, 4*vars]; *)
    XXZBRange =  Join[ConstantArray[{-Infinity, Infinity}, 2*vars], 
        ConstantArray[{0., Infinity}, vars], 
    ConstantArray[{0, 1}, vars]];
    (*Print[MatrixForm@XXZBRange];*)
    XXZBcons1 = 
   Join[constraints, ConstantArray[0., {cons, 3*vars}], 2];
    XXZBcons2 = Join[ConstantArray[0., {cons, vars}], constraints, 
        ConstantArray[0., {cons, 2*vars}], 2];
    Ymat = Join[IdentityMatrix[vars], -IdentityMatrix[vars], 2];
    XXZBcons3 = 
   Join[Ymat, -IdentityMatrix[vars], ConstantArray[0., {vars, vars}], 
    2];
    XXZBcons4 = 
   Join[-Ymat, -IdentityMatrix[vars], 
    ConstantArray[0., {vars, vars}], 2];
    (* Loop over orthogonal directions *)
  Do[(*Print[{"MM",MM}];*)(*Loop to find diameter in one direction,
   including incrementing MM if needed.*)
   For[itcount = 1, itcount < 10, itcount++, 
    XXZBcons5 = 
     Join[-Ymat, IdentityMatrix[vars], -MM*IdentityMatrix[vars], 2];
    XXZBcons6 = 
     Join[Ymat, IdentityMatrix[vars], MM*IdentityMatrix[vars], 2];
    XXZBcons = 
     Join[XXZBcons1, XXZBcons2, XXZBcons3, XXZBcons4, XXZBcons5, 
      XXZBcons6, orthocons];
    (*Print[MatrixForm@XXZBcons];*)
    XXZBvals = 
     Join[values, values, ConstantArray[0., 3*vars], 
      ConstantArray[MM, vars]];
    XXZBvals = 
     Transpose@{XXZBvals, ConstantArray[-1, 2*cons + 4*vars]};
    XXZBvals = Join[XXZBvals, ConstantArray[{0., 0}, i - 1]];
    (*Print[XXZBvals];*)
    XXZBobjective = 
     Join[ConstantArray[0., 2*vars], ConstantArray[1., vars], 
      ConstantArray[0., vars]];
    (*Print[XXZBobjective];*)(*Print[Dimensions/@{XXZBobjective,
    XXZBcons,XXZBvals,XXZBRange,XXZBDomain}];*)
    xxzb = Quiet[
      Check[LinearProgramming[-XXZBobjective, XXZBcons, XXZBvals, 
        XXZBRange, XXZBDomain, Tolerance -> 0.01], 
       "Unbounded", {LinearProgramming::lpsub}], \
{LinearProgramming::lpsub}];
    If[StringQ[xxzb], 
     Print["Unbounded LP in chord search - decreasing MM from " <> 
       ToString[MM]];
     MM = .5 MM; MaxComponent = MM,(*Print[{"xxzb",Chop@xxzb}];*)
     MaxComponent = Max@Abs[Ymat . xxzb[[;; 2*vars]]];
     (*Print[{"iteration, Max y component, MM value",itcount,
     MaxComponent,MM}]*)];
    If[MaxComponent < 0.49 MM, Break[], MM = 1.5 MM];];
   
   MM = MaxComponent = 2. Max@Abs[Ymat . xxzb[[;; 2*vars]]];
   perips = {xxzb[[;; vars]], xxzb[[vars + 1 ;; 2*vars]]};
   (*Print[{"Direction , chord",i,Norm@Apply[Subtract,
   perips]}];*)(*Prepare orthogonality constraint for the next \
direction*)
   AppendTo[peripoints, perips];
   AppendTo[orthocons, 
    Join[Normalize[Ymat . Flatten@Last[peripoints]] . Ymat, 
     ConstantArray[0., 2*vars]]];
   , {i, vars (*vars*)}];
       
   Chop@ peripoints]  
  
SetAttributes[MainChords, HoldFirst]; 
MainChords[solutionspace_, chordmax_: 40, chordmin_:10, flipmax_: 50, PrintResult_:False] :=
 (*This function aims to find a set of approximately maximal constraint polytope chords,
 that are mutually orthogonal and progressively decreasing.It does this by maximising 
 the sum of components rather than sum of absolute values,because this avoids 
 using MILP which is intractable for larger dimensions.
 To mitigate underestimation of a chord length because its vector components happen to 
 cancel in the current orientation of the solutionspace relative to the coordinate axes,
 the entire solution space is flipped to multiple different orientations and the maximal 
 value out of these is retained as the estimate.
 Using the chord endpoints, the origin of the solutionspace is improved by a call to RefineCentre.
 Each chord is calculated as a pair of endpoints, and "chords" is an array of such point pairs 
 which is the result returned by this function.
  
 Some coding below refers to diameters,but as they do not pass through a common centre the 
 term chord is a more accurate description.*)
 Module[{constraints, values, origin, OSBase, cons, vars, XXcons, 
   XXcons1, XXcons2, XX12cons, rotcons, orthocons = {}, XX12vals, 
   XXvals, XXRange, XXobjective, xx, maxchords, cmax, chordvec, chordlengths, 
   longchords, toplengths, posquad, quadrant, rotation, flip, mostflips=0,
   maxflips, decrement, totalflips, discard = False, repeats, 
   chords = {}, perips, diameters={}, peripoints = {}, 
   oldcentre, centreshift, descend, tol = 0.00001}, 
   {{constraints, values}, {origin, OSBase}} = solutionspace;
  {cons, vars} = Dimensions[constraints];
  If[vars == 1, chords = {DiagonalMatrix[values] . constraints};
  	 ChordLengths = Map[Norm[Subtract @@ #] &, chords];
  	 PeriPoints=chords[[1]];
   Return[chords]];
  progressrange={0,vars};  
  progresslabel=progresslabel<>" LP Search for main chords. ";
  Print[Style[progresslabel, Blue, TextAlignment -> Center]];
  posquad = ConstantArray[1., vars];
  maxflips = Min[flipmax, 2*vars]; (* The maximal flips performed for the first chord *)
  decrement = N[(maxflips - 3)/vars]; (* linearly reduce maxflips to reach 3 for the last variable *)
  repeats = Max[3, Ceiling@Log[2, vars]]; (* How many times the best value must repeat before 
  											the flipping loop is exited early  *)
  
  (*Set up the constraint set in the paired points or Y space*)
  XXRange = ConstantArray[{-Infinity, Infinity}, 2*vars];
  (*Print[MatrixForm@XXRange];*)
  XXcons1 = Join[constraints, ConstantArray[0., {cons, vars}], 2];
  XXcons2 = Join[ConstantArray[0., {cons, vars}], constraints, 2];
  XX12cons = Join[XXcons1, XXcons2];
  XX12vals = Join[values, values];
  XX12vals = Transpose@{XX12vals, ConstantArray[-1, 2*cons]};
  
  (* Only do this many chords by LP, the rest from periphery points.
  	 When the number of variables exceeds chordmax, the number of LP 
  	 chords is reduced linearly to reach 10 at 2*chordmax variables. *)
  cmax=Max[chordmax, chordmin ]; (* Always find at least chordmin chords *)
  maxchords=Which[vars<cmax,vars,cmax<=vars<=2*cmax-chordmin,2*cmax-vars,True,chordmin];	 
  (* Print[{"maxchords,maxflips",maxchords,maxflips}]; *)
  (* Print[{"chordmin,cmax,vars,maxchords :",chordmin,cmax,vars,maxchords}]; *)

  (*Loop over orthogonal directions to be maximised by LP. *)
  progresscounter = 0;
  Do[progresscounter++; totalflips = 0; 
  	
   XXcons = Join[XX12cons, orthocons];
   (*Print[MatrixForm@XXcons];*)
   XXvals = Join[XX12vals, ConstantArray[{0., 0}, i - 1]];
   (*Print[XX12vals];*)
   XXobjective = 
    Join[ConstantArray[1., vars], ConstantArray[-1., vars]];
   (* Print["LP arguments in MainChords ",Dimensions/@{XXobjective,XXcons,XXvals,XXRange}]; *)
   rotation = IdentityMatrix[vars];
   perips = {};
   (*Print[{"chord nr, maxflips",ToString[i],ToString[maxflips]}];*)
   
   (*The quadrant flipping loop *)
   For[flip = 1, flip <= maxflips, flip++,
    totalflips++;
   (* Print[{"Mainchords flipping: Flip no, Total flips",flip,totalflips}]; *)
    rotcons = 
     Chop[XXcons . ArrayFlatten[{{rotation, 0}, {0, rotation}}]];
    (*Print[Dimensions@rotcons];*)
    (*NOTE:
    If the coordinate center is on the periphery (i.e., the values vector contains one or 
    more 0's) numerical inaccuracy can place it outside, leading to an infeasible or unbounded LP. 
    This happens more often the larger the dimensions,  and when the LP tolerance value 
    is too small or absent or given as 10^-6 instead of 10.^-6.
    Also, the LP sometimes delivers a zero vector as solution -  makes no sense.
    The code below ignores such a trial and flips to a different orientation, which usually
    solves the problem. 
    It would be possible to end up in an infinite loop if this does not work, 
    so as a safety net such unsuccessful flips are limited not to exceed the maximum number
    of flips allowed.
    Note that Simplex with explicit lax tolerance is used, as InteriorPoint might be quicker 
    generally but occasionally stalls because it fails to converge.*)
    
    xx = Quiet[
      Check[LinearProgramming[-XXobjective, rotcons, XXvals, XXRange, 
        Method -> "Simplex", Tolerance -> LPtol], 
       "No LP solution", {LinearProgramming::lpsnf, 
        LinearProgramming::lpsub, 
        LinearProgramming::lpsnfp}], {LinearProgramming::lpsnf, 
       LinearProgramming::lpsub, LinearProgramming::lpsnfp}];  
       
    (*Print[{"xx",Chop@xx}];*)
    If[i > 1 && VectorQ[xx] && Norm[xx[[;; vars]] - xx[[vars + 1 ;;]]] > 5*Last@diameters, 
     xx = "Mainchords encountered an ill-defined LP problem; trying another flip."; Print[xx]];
    (* Discard any quadrant where the LP either fails or delivers a zero vector *)
    If[! VectorQ[xx, NumericQ],
     (* Print["Failure for (variable,flip) = " <> ToString[{i, flip}] <> 
       " - trying a different flip "];*)
     (*Print/@{XXobjective,rotcons,XXvals,XXRange};*)
     discard = True];
    If[xx == ConstantArray[0., 2*vars], 
     (* Print["Discard unsuitable LP solution " <> ToString[xx] <> 
       " for (variable,flip) = " <> ToString[{i, flip}]]; *)
     discard = True;];
    If[discard, flip--;
     quadrant = 
      MapAt[Minus, posquad, 
        Split@RandomSample[1 ;; vars, Floor[vars/2]]] + 
       RandomReal[0.1, vars];
     rotation = RotationMatrix[{posquad, quadrant}]; discard = False;
     (* Protect against infinite loop if no suitable solution is found after many flips *)
     If[totalflips > 2*maxflips, Break[], Continue[]]
     ];
    AppendTo[perips, {rotation . xx[[;; vars]], rotation . xx[[vars + 1 ;;]]}];
    chordlengths = Norm /@ (perips[[All, 1]] - perips[[All, 2]]);
    longchords = Ordering[chordlengths, -Min[repeats, Length[chordlengths]]];
    (* Exit early if the best value (within a 2% spread) was repeated several times *)
    If[flip > repeats,
     toplengths = chordlengths[[longchords]];
     If[First@toplengths/Last@toplengths > 0.98,
     (*	Print[ "Taking an early exit at flip "<>ToString[flip]<>
      " for chord nr "<>ToString[i]<>" with top lengths = "<>ToString@toplengths]; *)
      Break[]]
     ];
    (*choose a quadrant with about half of components negative,randomly chosen*)
    quadrant = MapAt[Minus, posquad, Split@RandomSample[1 ;; vars, Floor[vars/2]]] + 
      RandomReal[0.1, vars];
    rotation = RotationMatrix[{posquad, quadrant}];
    ];
   
   If[Length[perips] == 0, 
    Print["No meaningful chord was found for " <> ToString[i] <> 
      "-th direction; giving up after " <> ToString[totalflips] <> 
      " unsucessful flips."];Break[],
    
    AppendTo[diameters, chordlengths[[Last@longchords]]];
(* Print[{"Direction , chord length",i,chordlengths[[Last@longchords]]}]; *)
    
    (*Prepare orthogonality constraint for the next direction*)
    AppendTo[chords, perips[[Last@longchords]]];
    chordvec = Normalize[Subtract @@ perips[[Last@longchords]]];
    AppendTo[orthocons, Chop@Join[chordvec, -chordvec]];
    ];
    mostflips = Max[mostflips, totalflips]; 
   (* Decrease the number of flips linearly for the next chord, since the number of subsequent 
   chords that are determined by needing to be orthogonal to it, also decreases linearly. *)
   maxflips -= decrement; 
   , {i, maxchords}];
   
   foundchords=Length[diameters];

   If[foundchords< maxchords, Print["Attempt to calculate "<>ToString[maxchords]<>
   	" chords by LP, terminated after "<>ToString[foundchords]<>" successes. "];
   	maxchords=foundchords];
   
   ChordLengths =  Map[Norm[Subtract @@ #] &, chords];
   If[maxchords==0, peripoints={},
   	peripoints = Flatten[Transpose[chords, 1 <-> 2], 1];
   	peripoints = Select[peripoints, Norm[#] > tol &]]; 
   oldcentre=origin;
(* Print["Check inscribed centre is interior in MainChords, before Refinecentre: ",   
		outside=Chop@Min[values-constraints.InscribedSphere[[2]]]; 
   		If[outside >= -tol, True, "Outside by margin "<>TextString[Abs@outside]]]; *)
 (*  Print["Before RefineCentre, the constraint vectors span "<>ToString[MatrixRank[constraints, Tolerance -> 0.0001]]<>
   	" dimensions for "<>ToString[vars]<>" variables."]; *)
   PeriPoints = RefineCentre[solutionspace, peripoints, False, PrintResult];
   If[StringQ[PeriPoints],Print[Style["Aborting, because RefineCentre failed.",Red]]; Abort[]];

   (* Update local copy of solutionspace, the chords and the inscribed sphere
   position to the new centre.
   	Note that the SS origin is a point in the embedding space, so the  
   	shift needs to be projected to the lower dimensional space 
   	in which chords are defined.*)
   {{constraints, values}, {origin, OSBase}} = solutionspace;
   (* Print["After RefineCentre, the constraint vectors span "<>ToString[MatrixRank[constraints, Tolerance -> 0.0001]]<>
   	" dimensions for "<>ToString[vars]<>" variables."]; *)
 	centreshift=(origin-oldcentre) . Transpose@OSBase;
    chords = Map[{#[[1]] - centreshift, #[[2]] - centreshift} &, chords];
    InscribedSphere[[2]] = InscribedSphere[[2]] - centreshift;
(*
 Print["Check centre is interior in MainChords, before SampleChords: ",   
		outside=Chop@Min@values; 
   		If[outside >= -tol, True, "Outside by margin "<>TextString[Abs@outside]]]; 
 Print["Check chord endpoints are interior in MainChords, before SampleChords: ",   
		outside=Chop@Min@Map[(values - constraints.#) &, Flatten[Transpose[chords, 1 <-> 2], 1]]; 
   		If[outside >= -tol, True, "Outside by margin "<>TextString[Abs@outside]]];
 Print["Check inscribed centre is interior in MainChords, before SampleChords: ",   
		outside=Chop@Min[values-constraints.InscribedSphere[[2]]]; 
   		If[outside >= -tol, True, "Outside by margin "<>TextString[Abs@outside]]]; 
*)
  
   If[maxchords < vars, chords = CenterChords[solutionspace, chords, PrintResult];
   {{constraints, values}, {origin, OSBase}} = solutionspace;
	(* Print["Check chord endpoints are interior in MainChords, after SampleChords: ",   
		outside=Chop@Min@Map[(values - constraints.#) &, Flatten[Transpose[chords, 1 <-> 2], 1]]; 
   		If[outside >= -tol, True, "Outside by margin "<>TextString[Abs@outside]]] *)
   		]; 
   If[PrintResult,Print[ToString[maxchords]<>" chords were calculated by LP and "<>
   	ToString[vars-maxchords]<>" were approximated by diameters derived from peripheral points."]];
 
   ChordLengths =  Map[Norm[Subtract @@ #] &, chords];
  (* Flag the chords approximated by Samplechords, by making their lengths negative *)
   If[maxchords < vars, ChordLengths[[maxchords-vars;;]] = -ChordLengths[[maxchords-vars;;]]];
   (* Print["The chord lengths calculated by MainChords are: ",ChordLengths]; *)
   (*Ensure diameters are in descending order, which is not guaranteed for algorithm above.*)
   descend = Ordering[Abs@ChordLengths, All, Greater];
   chords = chords[[descend]];
   ChordLengths=ChordLengths[[descend]];
   
  (* Print["The largest flipcount in MainChords is "<>ToString@mostflips]; *)

  (*
  diams = Length[PeriPoints]/2; 
  pairtest =  Total@Map[Norm,    Normalize /@ PeriPoints[[;; diams]] + 
    Normalize /@ PeriPoints[[-diams ;;]]];
  Print["MainChords testing that PeriPoints are diametrically opposite :", pairtest<0.0001];
  *)

  Chop@chords]
  
SetAttributes[CenterChords, HoldFirst];
CenterChords[solutionspace_, knownchords_: {}, printresults_: False] :=
 (*This function supplements a given set of known LP chords (may be empty), by appending orthogonal spans. 
 The spans are selected from a sample of directions, taken from the periphery points delivered by invoking RefineCentre. 
 The current version only samples the origin and directions connecting the main point pairs. 
 Directions are progressively updated to give orthogonal chords. 
 A cycle of alternate direction sampling and recentering is used to avoid zero chords that can result from an incomplete
 set of directions caused by recentering having only a small number of LP chord points as input.
 
 NB:Run RefineCentre first to populate the global PerPoints array before calling this function.
 Avoid calling RefineCentre after SampleChords, otherwise diameters through a new centre 
 may be larger than the sampled chords.*)
 Module[{constraints, values, origin, OSBase, directions, dirank, 
   peripoints, pericount, cons, vars, inscribed = False, 
   chords, chordcount, chordvecs, chordlengths,
   chordsrequired, lastreq = Infinity, 
   keepsampling=True, recenter, oldcentre, centreshift, newbasis, newchord, 
   spans, bigspan, bigdir, minspan, sloop, tol},
   tol= 0.00001;   
  {{constraints, values}, {origin, OSBase}} = solutionspace;
  {cons, vars} = Dimensions[constraints];
  chords = knownchords;
  
  (*  The inscribed hypersphere diameter is a maximal lower limit to chord lengths.
  So if it has a significant radius, collect as many chords passing through its 
  center as possible. 
  This guarantees that the shortest chord lengths will not be underestimated, 
  leading to spurious flattening. 
  If the radius is zero, the current refined centre has a better chance to be on a low
  level facet and so avoid spurious zero lengths. *)
(*    Print["Check centre is interior in SampleChords, before shift to inscribed center: ",   
		outside=Chop@Min@values; 
   		If[outside >= -tol, True, "Outside by margin "<>TextString[Abs@outside]]]; *)

  If[InscribedSphere[[1]] > LPtol, (* shift the origin to inscribed centre *)
  	inscribed=True; 
  	centreshift = InscribedSphere[[2]];  
  	(* Print["Relocating the origin to inscribed centre at "<>ToString[centreshift]]; *)
  	values = values - constraints . centreshift;
  	origin = origin + centreshift . OSBase;
  	solutionspace = Chop[{{constraints, values}, {origin, OSBase}}];
    chords = Map[{#[[1]] - centreshift, #[[2]] - centreshift} &, chords];
    InscribedSphere[[2]] = InscribedSphere[[2]] - centreshift;
	PeriPoints=Map[(# - centreshift)&, PeriPoints];
  ];
(*  Print["Check centre is interior in SampleChords, after shift to inscribed center: ",   
		outside=Chop@Min@values; 
   		If[outside >= -tol, True, "Outside by margin "<>TextString[Abs@outside]]];  *)

  sloop=0;
  While[keepsampling, sloop++;
   progresslabel = " Supplement LP chords by approximate chords from sampled spans";
   Print[Style[progresslabel, Blue, TextAlignment -> Center]];
   progressrange = {0, vars};
   (* The following statement makes each cycle start from only the known, LP-based, chord set, 
   but with an enlarged set of PeriPoints derived from the recentering. 
   That may appear to be better, because all chords then pass through 
   the "best" known centre point.
   However, leaving it out produces chords in sets, 
   each of which pass through a different common intersection.
   And that is closer to the "proper" chords that are not
   constrained to have a common  intersection. 
   In practice resetting to LP chords only is quite wasteful of time \
(~3 times slower) since all chords calculated in earlier cycles are simply discarded, 
   and the final results are practically identical after sorting chordlengths.    *)
   (*chords=knownchords;*)
   
   progresscounter = chordcount = Length[chords];
   chordvecs = 
    If[chordcount > 0, Map[Normalize, Apply[Subtract, #] & /@ chords],
      ConstantArray[0., {1, vars}]];
   (* Extract a set of directions from peripheral point pairs *)
   {pericount, vars} = Dimensions[PeriPoints];
   directions = 
    Normalize /@ (PeriPoints[[;; pericount/2]] - 
       PeriPoints[[-pericount/2 ;;]]);
   (* Adding constraint directions ensures that the directions are (over-)complete. 
   Including this line slows the calculation by factor 2, 
   but is slightly better at avoiding zero chords. *)
   directions = Join[directions, constraints];

   (* Project directions so they are orthogonal to the known chords*)
   newbasis = NullSpace[chordvecs, Tolerance -> tol];
   directions = 
    DeleteDuplicates[
     Normalize /@ 
       Chop[directions . Transpose[newbasis]] . newbasis, (Abs[#1 . #2] > 
        1 - tol) &];
   directions = Select[directions, Norm[#] > tol &];
   (* Progressively select the longest span, add it to the chords, 
   then repeat with a reduced set *)
   While[Length[directions] > 0, 
    dirank = MatrixRank[directions, Tolerance -> tol];
    (* Print[{"Iteration, Chords required, No of dirs, Rank",
    	sloop,Length[newbasis],Length[directions],dirank}];  *)
    spans = Radii[constraints, values, #] & /@ directions;
    (*Print[{"Iteration,spans",i,spans}];*)
    bigspan = First@MaximalBy[spans, (Total@Abs[#]) &];
    If[Total@Abs@bigspan > tol,
     bigdir = First@directions[[FirstPosition[spans, bigspan]]];
     newchord = {bigspan[[1]]*bigdir, bigspan[[2]]*bigdir};
     AppendTo[chords, newchord]; progresscounter++;
     AppendTo[chordvecs, Normalize@Apply[Subtract, newchord]];
     newbasis = NullSpace[chordvecs, Tolerance -> tol];
     If[Length[newbasis]==0, directions= {}; dirank=0,
     	directions = DeleteDuplicates[Normalize /@ 
         Chop[directions . Transpose[newbasis]] . newbasis, 
         	(Abs[#1 . #2] >  1 - tol) &];
     directions = Select[directions, Norm[#] > tol &]]
     ,
     directions = {}; dirank = 0];
    ];
   (* Decide if recentering and sampling should be repeated because \
chord set is still incomplete. *)
   chordcount = Length[chords];
   (* Print[{vars, chordcount}]; *)
   chordsrequired = vars - chordcount;
   If[printresults,
   	Print[ToString[chordsrequired]<>" additional chords still required after this sampling round."]];
   keepsampling = Which[chordsrequired == 0, False,
     	chordsrequired < lastreq, True,
     chordsrequired >= lastreq,
     (* This round failed to find additional chords; 
     stop cycling and take minimal span chords along remaining directions *)
     chordvecs = Map[Normalize, Apply[Subtract, #] & /@ chords];
     newbasis = NullSpace[chordvecs, Tolerance -> tol];
     minspan=Max[InscribedSphere[[1]],tol];
     (* Print["The minspan value "<>ToString[minspan]<>" is being applied to "<>ToString[chordsrequired]<>
     " missing chords, while the inscribed sphere radius is "<>ToString@InscribedSphere[[1]]]; *)
     chords = Join[chords, Map[{minspan*#, -minspan*#} &, newbasis]];
     chordvecs = Join[chordvecs,newbasis]; False,
     True, 
     Print[Style["Something seriously wrong - can only find " <> 
       ToString[chordcount] <> " chords for " <> ToString[vars] <> 
       " dimensions!",Red]]; Abort[];
     False
     ];
   recenter = keepsampling || inscribed;
      
   If[recenter, lastreq = chordsrequired;
    peripoints = Flatten[Transpose[chords, 1 <-> 2], 1];
    peripoints = Select[peripoints, Norm[#] > LPtol &]; 
    oldcentre=origin;
    PeriPoints = 
     RefineCentre[solutionspace, peripoints, False, printresults]; 
    If[StringQ[PeriPoints], 
     Print[Style["Aborting, because RefineCentre failed.",Red]]; Abort[]];
   (* Update local copy of solutionspace and the chords to the new centre.
   	Note that the SS origin is a point in the embedding space, so the  
   	shift needs to be projected to the lower dimensional space 
   	in which chords are defined.*) 
   	{{constraints, values}, {origin, OSBase}} = solutionspace;
   	centreshift=(origin-oldcentre) . Transpose@OSBase;
    chords = Map[{#[[1]] - centreshift, #[[2]] - centreshift} &, chords];
    InscribedSphere[[2]] = InscribedSphere[[2]] - centreshift;
       ]
   (*; Print["Chord lengths ",Map[Norm,Apply[Subtract,#]&/@ chords]];*)
   ];   (* End while keepsampling *)
   
(* For consistency, check chord lengths through final centre and retain the longest *)
	(* chordvecs = Map[Normalize, Apply[Subtract, #] & /@ chords]; *)
	chordlengths=Map[Norm,Apply[Subtract,#]&/@ chords];
	(* Print[" After sampling, the chord lengths are: ",chordlengths ]; *) 
	spans = Radii[constraints, values, #] & /@ chordvecs;  
	chords=MapThread[If[Total@Abs[#1] > Abs[#2],
		 {#1[[1]]*#4, #1[[2]]*#4}, #3] &, {spans, chordlengths, chords, chordvecs}];
(* In cases where the inscribed radius is so small that its center was not used for sampling,
	it is possible that some chord lengths remain shorter than the inscribed diameter.
	This inconsistency will go away when flattening has removed negligible dimensions so
	that the inscribed radius becomes significant. *)
  progresslabel = 
   "Chord estimation by sampling completed in " <> ToString[sloop] <> 
    " iterations.";
  
  (*Print["Samplechords orthogonality matrix is: \n",MatrixForm@Chop[
  chordvecs.Transpose[chordvecs]]];*)
  Chop@chords]  
    

 (* Older version of Radii, that is not compiled. The code uses Reap/Sow.
  if this version is reactivated, the code without this, as compiled below, 
  should rather be used as it runs faster . *)
 
 (*
 Radii[Amat_, bvec_, vector_?(VectorQ[#, NumericQ] &)] :=
 (* Restricting to  a numeric argument prevents NIntegrate from trying
 symbolic evaluation. *)
 Module[{Avec, aprune, bprune, norm, unitvector, radii = {0, 0}, 
   tol = 0.001},
  If[Or @@ Negative@Chop[bvec], 
   Print["Radii not calculated because origin is not interior"]; 
   Return[{0., 0.}]];
  norm = Norm[vector];
  If[norm < 0.0001, 
  (* Print["Radii not calculated because direction vector " <> 
     ToString[vector] <> " is too short for comfort!"]; *)
   Return[{0., 0.}], unitvector = vector/norm];
  (* The components of all constraint plane normals along vector *)
  Avec = Chop[Amat.unitvector];
  (*Print[Avec];*)
  Do[Avec = i*Avec;
   (* Exclude planes that are not forwards inclined by at least a direction cosine > tol; 
   in effect that limits the aspect ratio of the polytope *)
   {aprune, bprune} = Reap[Do[
       If[Avec[[j]] > tol,
        Sow[Avec[[j]], ap]; Sow[bvec[[j]], bp]
        ]
       , {j, Length[Avec]}], {ap, bp}][[2]];
   (*Print[{"ap",aprune,"bp",bprune,"ratio",bprune/aprune}];*)
   (* If there are no \[LessEqual] constraints left, 
   it means the volume is open in that direction so radius is [Infinity] *)
   radii[[i]] = 
    If[aprune == {},(*Print["No constraint along vector "<>ToString[i*
     vector]];*)Infinity, Chop@Min[bprune/aprune]]
   , {i, 1, -1, -2}];(*Print[radii]*);
  If[radii[[1]] > radii[[2]], {radii[[1]], -radii[[2]]}, {-radii[[2]],
     radii[[1]]}]
  ]
*)

(* Using Compile makes the function faster by a factor 10 to 20 *)

Radii = 
 Compile[{{Amat, _Real, 2}, {bvec, _Real, 1}, {vector, _Real, 1}},
  Module[{Avec = {1.}, norm = 0., rad1 = 0., rad2 = 0., 
    Infinity = 10.^10, tol = 0.00001},
   If[Min[bvec] < -tol,
    Print[Style["Radii not calculated because origin is not interior by margin " <>
     TextString[Min[bvec]],Red]];Abort[];
    Return[{0., 0.}]];
   norm = Norm[vector];
   If[norm < tol, Print[Style[ "Radii not calculated, because the vector of length " <> 
      TextString[norm] <> " has an amnbiguous direction!",Red]]; Abort[];
    Return[{0., 0.}]];
   (*Avec consists of the components of all constraint plane normals, along vector*)
   (* For the compiler, dot product needs two matrices. 
   And change the result back to a row vector so comparison with row \
vector bvec works in compiler *)
   Avec = Flatten[{vector/norm} . Transpose[Amat]];
   (*Only keep hyperplanes that are forwards inclined by at least a direction cosine>tol;
   in effect that limits the aspect ratio of the polytope*)
   rad1 = Min@Table[
      If[Avec[[i]] > tol, bvec[[i]]/Avec[[i]], Infinity], 
      	{i, Length[Avec]}];
   rad2 = Min@Table[
      If[-Avec[[i]] > tol, -bvec[[i]]/Avec[[i]], Infinity],
       {i, Length[Avec]}];
   If[rad1 > rad2, {rad1, -rad2}, {-rad2, rad1}]
   ]
  (*,CompilationTarget\[Rule]"C"*)]
  
RegSimplexDirections[Ndim_]:=Block[{SimplexVertices,Rot},
(* This function returns the vertex coordinates of a regular unit N-simplex *)
(* The set of axis cutoffs in N+1 dims *)
SimplexVertices=IdentityMatrix[Ndim+1];
(* Rotate the last axis to (1,1,...1); note postmultiply because axes are rotated not vector. *)
SimplexVertices=SimplexVertices . RotationMatrix[{UnitVector[Ndim+1,Ndim+1],ConstantArray[1.,Ndim+1]}];
(* Project to N dim and normalise them *)
SimplexVertices=SimplexVertices[[All,;;-2]];
SimplexVertices=SimplexVertices/Norm[First@SimplexVertices]
]

SetAttributes[RefineCentre, HoldAll];
RefineCentre[solutionspace_, fixes_:{}, SimplexOnly_: False, PrintResult_:False] :=
 (* This finds diametrical pairs of peripheral points and the centroid of all these as a radial centre.
 Points are returned with the largest radius in each pair first, 
 and the pairs in descending order of the diameter.
 
 Argument fixes is a list of periphery points to be kept fixed, such as known chord endpoints. 
 These are kept as fixed periphery points, though they cannnot remain as diametrically 
 opposed pairs once the origin is shifted. Instead, each point acquires its own diametrically 
 opposed partner from the Radii function. This tends to neutralise the geometric bias that
 gets introduced by the progressively orthogonal scheme for diametrization so gives a better centre.
 However, it does mean that the diameters connecting the generated periphery points pairs will not
 include the maximal vertex-to-vertex distance of the original first diameter, although both of its 
 endpoints are still guaranteed to be present. 
 The values of fixes is adjusted to the new origin by this function.
 
 Note that SimplexOnly = True does not work well if the maximal aspect ratio is very large or small, 
 because then there are highly specific directions in which the boundary points are far away and 
 unlikely to be found by the small set of simplex directions.
 
 NB: It is essential that the solutionspace specification has normalized constraint vectors!
  *)
 Module[{constraints, values, OSBase, origin,cons, vars, facet, facetcount, nonfacet, 
 radii, order,diameters, chopdiameters, simplexdiameters,  raydiameters = {}, 
   keepers, open, widths, diffs, centrality = 0.0, newcentrality, centflag,shiftflag,
   peripoints, centroid, oldvalues, iterate = True, iteration = 0, CumulativeShift,
   meanrad, newmeanrad, margin, rot, outside,
   convergence = 0.0005,  nearby=0.05, maxcos = 0.99, tol = 0.00001},
  (* All hyperplanes with perp displacements less than a fraction "nearby" 
  of the mean radius, are treated as trappers below. *)

 {{constraints,values},{origin,OSBase}}=solutionspace; 
(* temporary estimate until actual radii are calculated *)
meanrad =Mean[Select[values,# >= Quantile[values,1/4]&]];
(* Print["Starting centre for the set of values "<>ToString[values]<>
	" has a radius estimate "<>ToString[meanrad]]; *)

  {cons, vars} = Dimensions[constraints];
  (* In the trivial 1D case, constraints already define periphery points, 
  so just shift the origin  *)
  If[vars == 1, peripoints = DiagonalMatrix[values] . constraints; 
  	 centroid = Mean[peripoints];
   	 values = Chop[values - constraints . centroid,tol];
 	 origin = Chop[origin + centroid . OSBase];
 	 solutionspace={{constraints,values},{origin,OSBase}}; 
 	 peripoints=Map[(#-origin)&,peripoints];
  Return[peripoints]];
(*
Print["Entering RefineCentre check that fixed points are inside: ",
	outside=Chop@Map[(values - constraints.#) &, fixes];
	If[Min@outside >= -0.001, True, "Outside by margin "<>TextString[Min@outside]]];
*)	
  centroid = CumulativeShift = ConstantArray[0., vars];
 simplexdiameters=RegSimplexDirections[vars];
 progressrange={0,1}; progresslabel="Refining the Kernel Space centering. ";
 Print[Style[progresslabel, Blue, TextAlignment -> Center]];
 While[iterate, iteration++;  (* PrintTemporary["Refining iteration no ",iteration]; *)
   margin=nearby*meanrad;
   facet = Flatten@Position[values, _?(# < margin&)];
   facetcount = Length[facet]; 
   nonfacet = Complement[Range[cons], facet];
   (* To enable escape where the origin lies near a constraint hyperplane intersection 
   (i.e. facet) an interior direction that does not cross any of these is needed. 
   Applying CapRayFinder to just the facet constraint planes, supplies one or more 
   such directions. Replace any constraint directions where the boundary lies closer 
   than the allowed margin, with this single "ray" direction. 
  
   It seems that in large models, especially simple cones, the calculation can get 
   bogged down because CapRayFinder does an LP which won't complete. So only
   do this escape maneuver on the first iteration. *)
 	diameters=  If[iteration < 2 && 1 < facetcount < vars,
    raydiameters = CapRayFinder[constraints[[facet]]]; constraints[[nonfacet]],	constraints];
 	diameters = DeleteDuplicates[ Join[diameters, simplexdiameters, raydiameters], 
 		Abs[#1 . #2] > maxcos &];
    (* Add diameters passing through current centroid and chord endpoints, 
    ensuring they are not be eliminated by DeleteDuplicates *)
   	chopdiameters = Map[Normalize[# - CumulativeShift] &, fixes];
	diameters=Join[chopdiameters,diameters];
	(* Print[{"I am calling Radii with directions ",diameters}]; *)
	radii = Map[Radii[constraints, values, #] &, diameters];

   (* Eliminate diameters along an open direction *)
   (* In the following, < 10.^9 should really be < Infinity, but compiled version of Radii 
   returns 10.^10 instead *)
   keepers = Flatten@Position[radii, _?(Norm[#] < 10.^9 &), {1}];
   open = Length[keepers] < Length[radii];
   radii = radii[[keepers]]; diameters = diameters[[keepers]];
(*  
 	 Print[{iteration,radii}];
 	 Print[{iteration,diameters}];
 *) 
   (* Eliminate diameters that give a nearly zero width, to prevent these 
   from trapping the mean in the corner which slows convergence *)
   widths = Map[Apply[Plus, Abs[#]] &, radii]; 
   radii = Pick[radii, widths, _?(# >2*margin &)];
	(* Print[{iteration,radii}]; *)
   diameters = Pick[diameters, widths, _?(# >2* margin &)];
      
   widths = Map[Apply[Plus, Abs[#]] &, radii];
   diffs = Map[Apply[Subtract, Abs[#]] &, radii];
   progresscounter=newcentrality = 1. - 0.5 Mean[diffs/widths];
   peripoints = 
    Join[diameters*radii[[All, 1]], diameters*radii[[All, 2]]];
   oldvalues = values;
   centroid = Retrieve[solutionspace, Mean[peripoints],PrintResult];
   values = Chop[values - constraints . centroid, tol];
   solutionspace[[1,2]] = values; (* Update for the sake of Retrieve in next iteration *)
  (* The following test has become superfluous now that Retrieve corrects 
  any minor numerical error in the centroid *)
(*  
	If[Or@@Negative[values],
Print["Centroid appears infeasible - Tolerance tol may need adjustment, 
inspect the following values vector for negative entries.\n"<>TextString@values];
Return["RefineCentre has failed."]];
*)
   	CumulativeShift = CumulativeShift + centroid;
	newmeanrad=RootMeanSquare[Flatten@radii];
 	(* Print[{"iteration, newmeanrad, CumulativeShift",iteration, newmeanrad, CumulativeShift}]; *)
	margin =nearby*newmeanrad;
(* Randomly reorient the simplex directions for the next iteration. 
This reorientation allows wider sampling of the available directions, 
but can slow convergence by a factor of 5 or more for small 2D examples. 
In higher dimensions, it seems to have far less effect on convergence.   *)
	rot=RotationMatrix[{UnitVector[vars,1],RandomReal[{0.,1.},vars]}];
	simplexdiameters=simplexdiameters . rot;
(*
  If[PrintResult,    
   Print["Centre refining iteration "<>ToString[iteration]<>" starts near "<>ToString[facetcount]<>
   " facets with "<>ToString[Length[diameters]]<>" diameters. \nIt yields centrality "
   <>ToString[newcentrality]<>" and shifts center by "<>ToString[Norm[centroid]]]];
*)
   (* Convergence based on the centroid appears to be slow, spiralling in to final centre. 
   Letting the mean radius converge is more effective. 
   But in the end it seems that the best convergence criterium is when 
   the fractional change of the centrality measure becomes negligible.   *)
   	shiftflag=Norm[centroid]>0.02meanrad;
   (*shiftflag=Abs[(newmeanrad-meanrad)/newmeanrad]>convergence;*)
   	meanrad=newmeanrad;
 	centflag= If[newcentrality > 0., 
     Abs[(newcentrality - centrality)/newcentrality] > convergence,  True];
 	centrality = newcentrality;
 	iterate =shiftflag||centflag; 
 	(* Safety valve to prevent convergence failure in large models *)
 	If[iteration>=Max[25,vars/4], iterate=False; 		
 	(* Print["Centering iteration loop failed to converge to required tolerance, but was terminated 
because it reached the allowed "<>ToString[Max[25,vars/4]]<>" iterations."] *)];   
   ];
   
   progresslabel=" Kernel space centering completed in "<>ToString[iteration]<>" iterations."; 
   progresscounter=0;
(*   Print[{"peripoints",peripoints}]; *)

(* On loop termination, the origin has been moved to the centroid of the 
latest peripoints, but the peripoints themselves are still specified relative to the centroid
of the previous iteration. There are various options:
1. Respecify the peripoints relative to the new origin. Then the origin will be 
at the centroid of the peripoints; but the peripoint pairs will no longer define
diameters through the origin.
2. Calculate new peripoints using radii from the new origin. This will guarantee
they are diametrically opposed, but the origin will not be at their centroid.
3. Keep the peripoints, but move the origin back to its previous position at 
their centroid. Again this ensures they are diametrically opposed but sacrifices
the origin being at their centroid.
For subsequent use, as in SampleChords, it is important to have diametric pairs.
So can use either 2 or 3; choose option 3 as it is the simplest. 

When the loop terminated by convergence, the origin should then be very close 
to the peripoints centroid, even if not exactly. If it terminated by iteration count,
there may however be a significant difference, so do not rely on this being true.
 *)
 
(* Print[{"Dimensions of origin,CumulativeShift,centroid,OSBase,fixes",
	Dimensions/@{origin,CumulativeShift,centroid,OSBase,fixes}}]; *)
 values = oldvalues;    (* Option 3 *)
 CumulativeShift = (CumulativeShift - centroid);
If[PrintResult, Print[ToString[iteration]<>
	" center refining iterations shifted the center a distance "<>TextString[Norm@CumulativeShift]]];
 origin = Chop[origin + CumulativeShift . OSBase];
 solutionspace={{constraints,values},{origin,OSBase}}; 
 fixes = Map[Subtract[#,CumulativeShift]&,fixes];
 If[open, 
   Print[" Open directions were detected, and are left out of the center refinement!"];
   Abort[]
];
If[SimplexOnly,
diameters=simplexdiameters;radii = Map[Radii[constraints, values, #] &, diameters]];
  (* Order the peripheral points into two sets, 
  forming pairs in descending order of each minor(second) radius  *)
order = Ordering[radii, All,Abs[#1[[2]]] >= Abs[#2[[2]]] &];
radii = radii[[order]]; diameters = diameters[[order]];
 (* Print[radii];Print[diameters]; *)
peripoints = 
   Join[diameters*radii[[All, 1]], diameters*radii[[All, 2]]];
(*
Print["RefineCentre check that peripheral points it found are inside: ",
	outside=Chop@Min@Map[(values - constraints.#) &, peripoints];
	If[outside >= -tol, True, "Outside by margin "<>TextString[Abs@outside]]];
	
Print["Leaving RefineCentre check that fixed points are still inside: ",
	outside=Chop@Map[(values - constraints.#) &, fixes];
	If[Min@outside >= -0.001, True, "Outside by margin "<>TextString[Min@outside]]];
*) 	    
peripoints]

SetAttributes[Retrieve, HoldFirst];
Retrieve[solutionspace_, point_, printresult_: False] := 
 Module[{constraints, values, OSorigin, OSbase, excess, YConstraints, 
   ZupConstraints, ZloConstraints, YZConstraints, vals, y, YZValues, 
   ZObjective, YZRange, YZ, newpoint, cons, vars, 
   tol = 0.00001},
   (*Given a candidate point,  this is tested to be feasible,
  i.e.in the interior of the solutionspace.If it is not,
  the value returned is the interior point closest to the given position vector*)
  {{constraints, values}, {OSorigin, OSbase}} = solutionspace;
  {cons, vars} = Dimensions[constraints];
  newpoint = point;
  excess = constraints . point - values;
  (*Print["Excess : ",excess];*)
  (*For any trivial discrepancies, 
  just move the boundary to include the point as this avoids an LP*)
  Do[If[0 < excess[[i]] < tol, values[[i]] += excess[[i]]], 
  	{i, Length@values}];
  solutionspace = {{constraints, values}, {OSorigin, OSbase}};
  excess = Chop[constraints . newpoint - values];
  (*Print["Excess : ",excess];*)
  If[Or @@ Positive@excess, 
   vals = Join[-excess, ConstantArray[0., 2*vars]];
   YZValues = 
    Transpose@Join[{vals}, {ConstantArray[-1, cons + 2*vars]}];
   (*Print[YZValues];*)
   YConstraints = 
    Join[constraints, ConstantArray[0., {cons, vars}], 2];
   ZupConstraints = 
    Join[IdentityMatrix[vars], -IdentityMatrix[vars], 2];
   ZloConstraints = 
    Join[-IdentityMatrix[vars], -IdentityMatrix[vars], 2];
   YZConstraints = Join[YConstraints, ZupConstraints, ZloConstraints];
   YZRange = 
    Join[ConstantArray[{-Infinity, Infinity}, vars], 
     ConstantArray[{0, Infinity}, vars]];
   ZObjective = Join[ConstantArray[0., vars], ConstantArray[1., vars]];
   (*Print[Dimensions/@{ZObjective,YZConstraints,YZValues,YZRange}];
   Print[{ZObjective,YZConstraints,YZValues,YZRange}];*)
   YZ = LinearProgramming[ZObjective, YZConstraints, YZValues, 
     YZRange, Method -> "Simplex", Tolerance -> tol];
   (*Print[YZ];*)
   If[VectorQ[YZ, NumberQ], y = YZ[[;; vars]];
    If[printresult, 
     Print["Specified point shifted by a distance " <> 
       TextString[Norm[y]] <> " to bring it inside the polytope."]];
    newpoint = point + y, 
    FailedConstraints=constraints;FailedValues=values;
    Print[Style["Retrieve failed - no interior point found which implies \
the constraints are incompatible.",Red]];Abort[]]];
  newpoint]

Centrality[solutionspace_, chordlengths_, periphery_] := 
 Module[{diams, rad1, rad2, difs,(* deviation,*) constraints, values, 
   OSorigin, OSbase, cons, vars, radii, pairtest, maxaspect, 
   meanaspect,logmeanaspect, colors, n, 
   chordpic, cenpic, inspan, insphereline, tol=10.^-6},
  
  maxaspect = Abs[First@chordlengths/Last@chordlengths];
  (* Calculate geometric mean of all aspect ratio's, 
  formed from pairs ordered as (major chord, minor chord) and cancelling factors *)
  n = Length[chordlengths];
  meanaspect=If[n<2, 1.,
  logmeanaspect = Sum[(n + 1 - 2 i) Log@Abs@chordlengths[[i]], {i, n}];
  Exp[logmeanaspect/(n (n - 1)/2)]
  ];
  colors=Sign[chordlengths]/. {1 -> Cyan, -1 -> Magenta};
  chordpic = 
   BarChart[Abs@chordlengths,  ChartStyle -> colors ,
    PlotLabel -> 
     Text[Column[{Style["MAIN ORTHOGONAL CHORD LENGTHS", Bold, Darker@Green], 
     	Style["Cyan chords, magenta diameters ", 10],
         Style["Blue line is max inscribed sphere diameter", 10]}(*, Spacer[15]*)]],
         Frame -> True, 
    FrameLabel -> {" The mean and maximal aspect ratio's are " <> TextString[Round[meanaspect, 0.1]] <> 
       " and " <> TextString[Round[maxaspect, 0.1]]},
    ImageSize -> All, ScalingFunctions -> "Log"];
    
  (*Calculate centrality deviation over all periphery points*)
  diams = Length[periphery]/2;
  pairtest = Max[
   	Map[Norm, Normalize /@ periphery[[;; diams]] + Normalize /@ periphery[[-diams ;;]]]/
   		Map[Norm, periphery[[;; diams]] - periphery[[-diams ;;]]]
   ];
  If[pairtest > 0.01, 
   Print[Style["WARNING: The direction vectors of a diametrically opposed periphery pair
 failed to cancel within 1% of their diameter. This likley reflects inaccuracy of small
 diameters and may require further flattening.",Darker@Green]]];
  rad1 = Norm /@ periphery[[;; diams]];
  rad2 = Norm /@ periphery[[-diams ;;]];
  difs = rad1 - rad2;(*Print[difs];*)
  deviation = 50*Mean[difs/(rad1 + rad2)];
 
  (*But plot only the radii along main chord directions*)
  {{constraints, values}, {OSorigin, OSbase}} = solutionspace;
  {cons, vars} = Dimensions[constraints];
  (*Constraints are expressed in the local coordinate system.This is \
already aligned with the main diameters,
  i.e.the basis vectors are the axis unit vectors,i.e.identity matrix*)
  radii = Map[Radii[constraints, values, #] &, IdentityMatrix[vars]];
  radii=radii/.{0.->tol};
  {rad1, rad2} = Transpose@Abs[radii];
  (*Note that these radii and hence also mindim and maxdim all pass \
through the current origin and hence will generally be smaller than \
the maximal chord lengths (main diameters), even though parallel to them*)
  Diameters=rad1+rad2; (* Store the diameters for later checking against chords *)
  cenpic = PairedBarChart[Transpose[{rad1, rad2}], 
    Transpose[{rad2, rad2}], ScalingFunctions -> "Log", 
    BarOrigin -> "XAxis", PlotRange -> All, 
    ChartLayout -> "Overlapped", 
    BarSpacing -> {0, Automatic, Automatic}, 
    PlotLabel -> 
     Text[Column[{Style["SPLIT DIAMETERS ALONG MAIN CHORD DIRECTIONS", Bold, Darker@Green], 
        Style["Asymmetric overhang shown in light shading.", 10],
        Spacer[15]}(*, Spacer[15]*)]],
    Frame -> True, 
    FrameLabel -> {"Average periphery point non-centrality "<>TextString[Round[deviation, 0.1]] <> 
       "%"},
     ChartStyle -> {{Lighter[Magenta, 0.7], Lighter[Magenta, 0.2]}}, 
    Ticks -> {Automatic, ticks}, ImageSize -> All];
  inspan=Log[2.*InscribedSphere[[1]]];
  insphereline={Blue, Thick, Line[{{0.5, inspan}, {Length[chordlengths]+0.5, inspan}}]};
  chordpic = Show[chordpic, Graphics[insphereline]];
  {chordpic, cenpic}
  ]

ChordPairs[perips_,transform_]:=Module[{diffs,axpairs,SSKperips,chordvecs,count,dim,maxi, nochords,mainchords},
(* Function to extract main chord endpoints from the list of FFS peripheral points. 
The argument <transform> is the one that defines the SSK relative
	to whatever coordinates the point list <perips> is expressed in. 
It returns a list of index pairs that identify chord endpoints in the <perips> list. *)
SSKperips=DowncastPoint[perips,transform];
{count,dim}=Dimensions[SSKperips];
(* Get all chord vectors*)
(* Note. Chord lengths can vary by 5 or 6 orders of magnitude. Normalize
	the difference vectors so Chop can reliably eliminate negligible components.*)
diffs=Table[Chop[Normalize[SSKperips[[i]]-SSKperips[[j]],Norm[#,1]&],0.0001],
				{i,count},{j,i+1,count}];
(* Main chords define the SSK coordinate axes. 
	Select all chords that are along an axis, i.e. has a single non-zero component *)
axpairs=Position[diffs,{0...,Except[0],0...},{2}];
(*Print[axpairs];*)
chordvecs=Extract[diffs,axpairs];
(*Print[chordvecs];*)
If[chordvecs=={},
 MessageDialog["No chords aligned with the coordinate axes were found.
Check that the transform passed to ChordPairs is the one that expresses 
the given periphery points in SSK coordinates, else this is bound to happen! "];
 Return[{}]];

(* Choose just the longest one along each axis. But if the longest one has 
	length 0, it is not a chord; alert the user. *)
mainchords=Flatten@Reap[Do[
	maxi=First@MaximalBy[chordvecs,Abs[#[[i]]]&,1];
	Sow[If[maxi[[i]]==0,0,Position[chordvecs,maxi]]],{i,1,dim}]][[2]];
nochords=Position[mainchords,0];
If[nochords!={},
 MessageDialog["No chords were found along axes "<>ToString[Flatten@nochords]<>
 " of the SSK basis. Most likely these are thin directions not flattened out."];
 mainchords=Delete[mainchords,nochords]];
(* Select out the main chord pairs and convert them from index in the 
	triangular diffs matrix to get them as indices in the periphery points list *)
axpairs=axpairs[[mainchords]];
axpairs=Map[{#[[1]],#[[1]]+#[[2]]}&,axpairs];
(* The actual chord endpoint pairs could be recovered by *)
     (* chords=Map[perips[[#]]&,axpairs] *)
axpairs
]
  
End[] (* End Private Context *)

EndPackage[]
```
