## Supplementary material for "The FBA Solution Space Kernel – Introduction and Illustrative Examples": SSK software manual and code: Configuration.m

```
(* ::Package:: *)

(* COPYRIGHT
					© Copyright 2022 Wynand Verwoerd

This file is part of SSKernel.

The SSKernel program is free software; you can redistribute it and/or modify it under the terms of the GNU General Public License as published by the Free Software Foundation; either version 3 of the License, or (at your option) any later version.
 
This program is distributed in the hope that it will be useful, but WITHOUT ANY WARRANTY; without even the implied warranty of MERCHANTABILITY or FITNESS FOR A PARTICULAR PURPOSE . See the GNU General Public License for more details.
 
You should have received a copy of the GNU General Public License along with this program . If not, see http://www.gnu.org/licenses/
*)

(* Mathematica Package *)

BeginPackage["Configuration`"]
(* Exported symbols added here with SymbolName::usage *)  

(* ADJUSTABLE OPTIONS AND TOLERANCES THAT REPRESENT COMPUTING TIME/ACCURACY TRADEOFFS *)

If[loadreset || !ValueQ[loadreset],
loadreset=True; (* True means adjustable options are reset to values below when a new model is loaded. *)
reshuffle = False; (* Whether to randomly reorder reactions and metabolites before executing stage 2. *)
exempt = False; (* Whether to exempt external metabolites that lack buffering, from flux balance *)
LPmethod=Automatic; (* Method used for initial bounded FBA: Simplex for small models and Interiorpoint for large.*)
(* LPmethod="Simplex"; *)  (* When active, this forces the use of the nominated method.*)
Tolerances=ToString[0.0001];	fixtol=0.002; (*	LoadModel function sets these tolerances according to the number of flux variables in a model *)
artificial = 100.; (* Flux upper limits larger than this are considered artificial and set to Infinity *)
targetcount = 20; (* The number of random FBF's for initial progenitor estimate *)
maxthin=0; (* A thickness less than this may be flattened to zero *)
maxaspect = 50.; (* SSK aspect ratio larger than this is avoided if possible, by flattening
any dimensions thinner than maxthin  *)
samplesize = 500; (* The number of BFBFs found by greedy random search, that is considered an adequate sample *)
mixfraction = 0.80; (* Moderate greedy search by mixing at most the top "mixfraction" score levels *)
greedyfails = 20; (* Allowed number of rejections (see above)  
	before sampling is terminated because of small prospect to find any more BFBF's. *)
treesize = 100000; (* The maximum estimated nodes to be visited by exhaustive tree search, otherwise use greedy search *)
chordmin = 10;  (* The minimal number of chords found by LP. *)
chordmax = 50; (* The maximal number of chords found by LP, any more are approximated from periphery points.  *)
flipmax= 25; (* The maximal number of times quadrants are flipped during main chord calculation.  *)
DefaultCap=1.0;  (* Default capping radius is applied if SSK is a simple cone with no FBF's *)
timeconstraint=60.; (* Time allowed for calculating flattening errors and enclosing simplexes *)
]; 

(* INPUT DATA - VARIABLES THAT DEFINE THE FBA MODEL *)

ModelName; (* An identifier, e.g. BIGG model ID and/or biological species *)
S = {}; (* The MxN stoichiometry matrix, such that S.F = 0 where F is the N-dim flux vector. *)
Svals = {}; (* The input data RHS and equal/inequal for S.F This is forced to  =0 in SSK calculation.*)
bounds = {}; (* A Nx2 matrix, each row gives the lower and upper bound for one flux component *)
objectselector = {}; (* An N-dim (often unit) vector, that selects the flux combination to be optimized *)
maxmin = 1; (* Specifies how the objective is optimized: -1 for maximization, 1 for minimization *)
FBAvector ={}; (* An optimized N-dim FBA flux vector, produced previously in an external LP optimization. *)
ReactionNames = {}; (* Names of the N reactions, in the order of the columns of S; i.e., flux identifiers *)
MetaboliteNames = {}; (* Information only: Names of the M metabolites, in the order of the rows of S *)

(* USER INTERFACE VARIABLES *)

commandline=False; (* Whether the program was run from a script or batch file *)
(* Control panel parameters *)
panelback = ConstantArray[LightYellow,5]; (* Each panel coloured yellow when passive, green when calculating, pink if terminated unsucessfully *)
available = {True, False, False, False, False, False};  (* switches for greying out control panel stage buttons *)
verbose; (* Print details of progress *)
automate; (* Whether to execute all main calculation stages without button prompting *)
Species; (* Biological species; default is parent directory of model input file *)
exemptcount=0; (* The number of metabolites confirmed for flux balance exemption *)
(* Global counter used for progress monitor display *)
progresscounter=0; progressrange={0,1}; 
progresslabel="Progress of current calculation";

(* Displayed results: tables and graphics *)
Heading={"",""}; (* The heading used in reports. *)
Reductiontable={}; Kerneltable={}; Processreport = {}; 
Optiontable := Transpose@{{"Step time limit", "Unbuffered externals exempted","LP tolerance", "Fixed value tolerance", 
	"Progenitor sample size", "Max BFBF tree nodes ", 
    "BFBF random greedy sample size", "Gready search mixing fraction", "Stop sampling when failure rate >",
    "Minimal, Maximal LP chord counts", "Maximal flips to find LP chords", 
     "Aspect ratios \[GreaterEqual] this are flattened",  "Diameters > this not flattened ",
    "Default capping radius","Flux bounds \[GreaterEqual] this are taken as artificial"}, 
    {timeconstraint, exemptcount, ToExpression@StringSplit[Tolerances, ","], fixtol, targetcount,
    	treesize, samplesize, mixfraction, greedyfails, ToString@{chordmin, chordmax}, 
    	flipmax, maxaspect, maxthin, DefaultCap, artificial}};
Validations={}; 
tabheads = {{"Not bounded", "Art. bounded", "Imported"}, 
	{"Flux vector\nlength", "% Flux\nmismatch", "Misalignment\nangle deg"}};
chordpic={}; cenpic={}; flatplot={}; 

(* GLOBAL VARIABLES FOR SSKernel CALCULATION  *)

KernelSpaceVersion = " Version 1.4 Jun 2024 ";
DataDirectory=""; (*Base directory path, datafile specified relative to this in command line  version *)
(* DataDirectory="C:/Users/John Smith/Documents/Data/"; *)       (*EXAMPLE 1: Forward slashes *)
(* DataDirectory="C:\\Users\\John Smith\\Documents\\Data\\"; *)  (*EXAMPLE 2: Note every \ needs to be doubled*)
(* DataDirectory="F:/Lincoln_Backup/Research/Projects/Volume visualisation/"; *)

datafile; PackagesDirectory; 
externals={}; (* A list of row numbers in S for metabolites that are external to the network *)
exemptions={}; (* A list of S matrix rows optionally exempted from flux balance because of a lacking buffer*)
exterior = "extra" | "exter"; (* Compartments with either of these strings in their ID or name,
		 are identified to be extra-cellular and their metabolites are classed as external. *)
outsuffix = "_e" | "[e]" | "(e)"; (* In the absence of compartment allocation, metabolites with
		any of these strings as a suffix to their ID or name, are classed as external.*) 
SStype="FacetCone"; (* Default - a cone with multiple bounded facets *)
apex={}; (*Position vector in RSS of the apex, if the Kernel space is a simple cone *)
augment = {{}}; (* The augmented matrix consisting of stoichiometry constraints and values, 
	as well as constraints to fix the objectives to their optimized values *)
objectcount=1; (* Currently only single objective implemented - can be extended.*)
lowers = {}; uppers = {}; (* Lower and upper bounds for fluxes. *)
feasiblepoints = {}; (* Feasible fluxes collected from FBA and fixed flux determination *)
LPtol; FBAtol;   (* Numeric tolerance used for LP calculations *)
fixvals={{}}; (* Pairs of fixed fluxes, {fluxno, fixed value} *)
fixdirs={}; (* Reversible reactions that get assigned directions from fixed flux search *)
fixflag = False; (* True means that the fix tolerance is more than 10% of the maximal unbounded FBA flux component. *)
NonfixTransform={{},{}}; (* Matrix transform to fixed values subspace *)
progen={}; (* The FBF progenitor facet *)
prismrays={}; (* The set of prismatic rays as vectors in variable part of flux space, or VFS *)
coincrays={}; (* The set of conical rays eliminated by progenitor coincidence capping as VFS vectors  *)
conerays={}; (* The set of conical progenitor rays subjected to tangent capping as VFS vectors *)
(*rays={};     (*The combined set of rays in full flux space *) *)
raymat={}; (* The non-convex complete ray basis and its constraint overlaps *)
RayDim=0;  (* The ray space dimension *)
RSSdims = 180; (* The max RSS dimension count that allows a viable FBF search *)
BoundedFacets={}; (* Lists of constraint plane intersections that form FBF's*)
Infeasibles={}; (* Lists of infeasible facets encountered during tree search *)
FBFcount=0; (* The number of FBF's found so far *)
totalfacets=0; (* The number of facet nodes visited during tree search or sampling *)
emptycycles=0; (* The number of consecutive traceback cycles in progenitor search
					that failed to find an ancestor *)
rejections=0; (* The mean number of duplicate FBF rejections over the last 10 greedy trials *)
CappingRadii={}; (* The radii used for tangent capping at remotest FBF vertex *)
thindirs={}; (* The set of directions that have been compressed *)
InscribedSphere={0.,{0.}}; (* Radius and centre of maximal inscribed hypersphere *)
PeriPoints={}; (* A set of points in FFS that fall on the periphery of SSK *)
ChordLengths={}; (* The lengths of the SSK main orthogonal chords  *)
Diameters={}; (* SSK diameters through the refined centre, along chord directions  *)
ExportResults={}; (* Detailed results, to be exported to *.DIF file *)

(* Interim results, stored to facilitate repetition of earlier stages *)

(* Raw input values, To restore model definition after exemptions or reshuffle *)
{Sraw,rawSvals,rawbounds,rawobject,rawFBAvec,rawreacts,rawmets,rawexternals,rawexempts}; 
rawfeasibles; boundedflux; openflux; FBAdone=False;
RSStype;  RSSraydim; SSKinRSS; SSKinsphere;
LoadReports; RSSReports; KernelReports; ShapeReports; 
Loadtime; RSStime; Kerneltime; Shapetime; Chordtime;
(* When flattening is repeated, without changing chordmin and chordmax, the chord
calculation need not be repeated; so store its results for reuse *)
PreflatSSK={}; PreflatPeris={}; PreflatChords={}; PreflatChordLengths={}; 

(* SOLUTION SPACE DATA STRUCTURES *)

(* The following lines define names for components of solution and Kernel spaces to 
facilitate calling of functions that use them. 
NOTE: Do not assign values to these names, or use them in fuctions that change their arguments 
i.e., have HoldFirst , HoldAll etc. attributes, as that will freeze them to fixed values. *)


(*Reduced space (RSS) implementing stoichiometry and objective constraints, 
after removing fixed fluxes, prismatic rays and linealities *)
ReducedSS={{},{}}; 
RSScons:=ReducedSS[[1,1]];RSSvals:=ReducedSS[[1,2]];
RSSTransform:=ReducedSS[[2]]; (* Affine transform from FS to SS *)
RSSOrigin:=ReducedSS[[2,1]];RSSBasis:=ReducedSS[[2,2]];

(* Bounded Kernel solution space after coincidence and tangent capping, relative to RSS  *)
KernelSpace={{},{}};
Kernelcons:=KernelSpace[[1,1]];Kernelvals:=KernelSpace[[1,2]];
KernelTransform:=KernelSpace[[2]]; (* Affine transform from  RSS to SSK *)
KernelOrigin:=KernelSpace[[2,1]];KernelBasis:=KernelSpace[[2,2]];


(* DIAGNOSTIC VARIABLES *)

 
Begin["`Private`"] (* Begin Private Context *) 


End[] (* End Private Context *)

EndPackage[]
```
