## Supplementary material for "The FBA Solution Space Kernel – Introduction and Illustrative Examples": SSK software manual and code: DataHandler.m

```
(* Wolfram Language Package *)

(* COPYRIGHT
					© Copyright 2022 Wynand Verwoerd

This file is part of SSKernel.

The SSKernel program is free software; you can redistribute it and/or modify it under the terms of the GNU General Public License as published by the Free Software Foundation; either version 3 of the License, or (at your option) any later version.
 
This program is distributed in the hope that it will be useful, but WITHOUT ANY WARRANTY; without even the implied warranty of MERCHANTABILITY or FITNESS FOR A PARTICULAR PURPOSE . See the GNU General Public License for more details.
 
You should have received a copy of the GNU General Public License along with this program . If not, see http://www.gnu.org/licenses/
*)

BeginPackage["DataHandler`",{"Configuration`","StringToDouble`"}]

(* Exported symbols added here with SymbolName::usage *)  
MReader::usage = "Reads input data {modelname, S, bounds, objectvector, FBAvector, 
	MetaboliteNames, ReactionNames, maxmin} from a Mathematica .m or .WL file" 
MatReader::usage="Reads input data from a MatLab .mat file, using COBRA spec as in  https://arxiv.org/pdf/1710.04038.pdf  "
SBMLReader::usage="Reads input data {modelname, S, bounds, objectvector, FBAvector, 
	MetaboliteNames, ReactionNames, maxmin} from a SBML file "
CheckBalance::usage="Inspects Stoichiometry matrix for proper sinks and sources.  "
Shakeup::usage="Reorder reactions and metabolites randomly.  "

Begin["`Private`"] (* Begin Private Context *) 


SetAttributes[MReader, HoldRest];
MReader[file_, S_, Svals_, bounds_, Object_, FBAvector_, 
	 ReactionNames_, MetabNames_, maxmin_, printresult_:False] := 
(* This function reads input data from a Mathematica .m or .WL file that contains the numeric 
arrays as a list of the structure 
		{modelname, S, Svals(Optional), bounds, objectvector, FBAvector, ReactionNames, MetabNames, maxmin}. 
Here modelname is a string, which may include a formal ID and a desciption of the model. It has to appear first.
The remaining items may appear in arbitrary order, and any additional list items are ignored.
maxmin is not an array, but a string containing either "max" or "min"
to indicate whether the objective should be minimized or maximized. 
It is optional, the default is taken as maximisation. 

From within Mathematica, having created the individual arrays somehow, produce the file  by executing: 
	Export[exportfile, {ModelName, S, Svals, bounds, objectselector, FBAvector,  ReactionNames, MetaboliteNames,
  Switch[maxmin, -1, "max", 1, "min", _, "max"]}, "List"];

*)
 Block[{stringimport, eximport, import, indims, candidates, foundvals, Sat, valsat, 
 	boundsat, cons, vars, vectorat,	namesat, FBAvecat, selectorat,SvalsTest},
 	(* Test whether a vector satisfies all constraint equations as determined by Svals *)
  SvalsTest[svals_, vector_] := Block[{equals, smallers, biggers},
  	equals = Position[svals[[All, 2]], 0];
  	smallers = Position[svals[[All, 2]], -1];
  	biggers = Position[svals[[All, 2]], 1];
  	And @@ Join[
    MapThread[Equal, {Extract[vector, equals], Extract[svals[[All, 1]], equals]}], 
    MapThread[LessEqual, {Extract[vector, smallers], Extract[svals[[All, 1]], smallers]}], 
    MapThread[GreaterEqual, {Extract[vector, biggers], Extract[svals[[All, 1]], biggers]}]]
  ];
  Print["Data imported from file " <> file];
(* Using the "List" format takes care of importing numbers written e.g. as 1.23E-10 *)
  eximport = Import[file, "List"];
  ModelName=First@eximport;
  eximport= ToExpression /@ Rest[eximport];
  (* Separate off any non-array, stemming from a string, and convert it back *)
  import=Select[eximport, ArrayQ];
  stringimport=ToString /@ Complement[eximport,import];
  stringimport = If[stringimport == {}, "max", stringimport[[1]]];
  maxmin=Which[StringContainsQ[stringimport, "max", IgnoreCase -> True], -1, 
 	StringContainsQ[stringimport, "min", IgnoreCase -> True], 1, True, -1];
  (* The imported arrays may be in arbitrary order and have 
	superfluous data, such as the FBA optimum value, interspersed. 
  	So identify the required ones by their dimensions and data type. *)
  
  indims = Dimensions /@ import;
  If[printresult, Print["The arrays as imported have the following dimensions \n"<>  ToString[indims]]]; 
  Sat = FirstPosition[indims, {_, _?(# > 2 &)}, {0}][[1]];
  If[Sat > 0, S = import[[Sat]]];
  If[Sat==0 || !MatrixQ[S, NumberQ], 
  	Print[Style["Abort - No 2D numeric array with more than 2 rows could be identified 
   as a stoichiometry matrix in the input data.",Red]]];
  {cons, vars} =indims[[Sat]];
  (* Find arrays with the right dimensions to contain values and bounds *)
  valsat = Position[indims, {cons, 2}, {1}];
  boundsat = Position[indims, {vars, 2}, {1}];
  (* Multiple cases of each may be found, in particular if vars = cons. 
	Isolate the values array by checking that it has pairs with the second element -1, 0, or 1 *)
  candidates = If[valsat != {}, Extract[import, valsat], {}];
  foundvals = FirstPosition[candidates, 
  	{Repeated[{_?NumberQ, _?(Sign[#] == # &)}, {cons}]}, {}];
  Svals = If[foundvals != {}, Extract[candidates, foundvals], {}];
  If[Svals == {}, Svals = ConstantArray[{0, 0}, cons];
  	If[printresult, Print["Default: constraints taken as = 0"]]];
  (* Remove values array, then check remaining bounds candidates for 
	the first one having low-high pairs *)
  boundsat = Complement[boundsat, {Extract[valsat, foundvals]}];
  candidates = If[boundsat != {}, Extract[import, boundsat], {}];
  foundvals = FirstPosition[candidates,
  	 _?(Count[#, {lo_, hi_} /; hi >= lo] == vars &), {}];
  bounds = If[foundvals != {}, Extract[candidates, foundvals], {}];  
  If[bounds == {},
      Print[Style["Abort - A 2D numeric array with "<>ToString[vars]<>
      " rows, each a pair of low and high numbers, to serve as bounds, "<>
      "could not be be identified in the input data.", Red]]; Abort[]];
  (* Next, vectors with reaction names, objective selector and possibly FBA solution *)
  vectorat = Flatten@Position[indims, {vars}]; 
  namesat=Flatten@Position[import, _?(VectorQ[#, StringQ]&)];  
  ReactionNames = Cases[import[[namesat]], _?(Length[#]==vars&)];
  ReactionNames = If[ReactionNames == {}, 
  	Print["No suitable list of reaction names or ID's were detected in the import data!"];
  	ConstantArray["NoName",vars], First@ReactionNames];
  MetabNames = Cases[import[[namesat]], _?(Length[#]==cons&)];
  MetabNames = If[MetabNames == {}, 
  	Print["No suitable list of metabolite names or ID's were detected in the import data!"];
  	ConstantArray["NoName",cons], First@MetabNames];
  externals = Flatten@Position[
   MetabNames, _?(StringContainsQ[#, outsuffix ~~ EndOfString] &), Heads -> False];	

  vectorat=Complement[vectorat,namesat];  
(*  At this point, any numeric vectors in flux space are isolated.
	If any of those satisfies the constraint set, the first of those 
	is taken as the FBA flux; if none, it is set to a zero vector.  *) 
  FBAvecat = Cases[vectorat, _?(SvalsTest[Svals,S.import[[#]]] &)]; 
  FBAvector = If[FBAvecat == {}, ConstantArray[0., vars], import[[First@FBAvecat]]];
  selectorat = Cases[vectorat, _?(0 < Count[import[[#]], Except[0 | 0.]] < 5 &)];
  Object = Switch[Length[selectorat],
    0, Print@
     "Input data invalid - no objective vector that combines fewer 
than 5 fluxes could be identified"; ConstantArray[0., vars],
    1, import[[First@selectorat]],
    _, Print@
     "Multiple apparent objective vectors found in input data, only 
the first one is taken."; import[[First@selectorat]]];
  Print["Dimensions of imported stoichiometry, values, bounds, objective and FBA flux arrays: " <> 
    ToString[Dimensions /@ {S, Svals, bounds, Object, FBAvector}]]; 
  Print["The objective vector selects contributions from the flux set " <> 
  	ToString[Flatten@Position[Object, _?(# > 0 &)]]];
  ]

SetAttributes[MatReader, HoldRest];
MatReader[file_, S_, Svals_, bounds_, Object_,  FBAvector_,
	 ReactionNames_, MetabNames_, maxmin_, printresult_:False] := 
(* This function reads input data from a MatLab .mat file that specifies a metabolic model
according to the COBRA specification (See https://arxiv.org/pdf/1710.04038.pdf). 
The data item "osenseSTR" is optional and if absent, the objective is maximized. *)
 Block[{labels, modelname, description, missing, ruleform, reactionIDs, reactionnames, 
 	reactionECnos, metIDs, metnames, lowers, uppers, ineq, vals, cons, vars},
  maxmin="max";
  Print["Data imported from file " <> file];
  If[printresult,Print@Import[file, "Comments"]];
  (* The following used to work with Mathematica versions up to 12.1*)
(*
  labels = Import[file, "LabeledData"][[1, 2, All, 1]];
  ruleform = First[Import[file, "Labels"] /. Import[file, "LabeledData"]]; 
*)
  (* But since 12.2, it imports as a set of associations. Convert to rules by Normal. *)
  ruleform = Normal[Import[file, "LabeledData"]];
  If[MatchQ[ruleform[[1]], _ -> $Failed], 
  	Print["Reading aborted because data file is invalid. "]; Abort[], 
  	ruleform = ruleform[[1, 2, 1, 1, All]]];
  labels = ruleform[[All, 1]];
  If[printresult,Print["The labeled data items in the input file are: "<>ToString[labels]]];
  missing = Complement[{"S", "lb", "ub", "b", "c"}, labels];
  If[missing!={},
  	Print[Style["Imported data file is incomplete; the following essential data was not found:"<>
  	ToString[missing /. {"S" -> "Stoichiometry matrix", "lb" -> "Lower bounds", 
  		"ub" -> "Lower bounds", "b" -> "S-constraint values", "c" -> "Objective"}],Red]];
  Abort[]];
  {modelname, description, S, lowers, uppers, Object, ineq, vals, maxmin, 
  	reactionIDs, reactionnames, reactionECnos, metIDs, metnames} = 
  	{"modelID", "description", "S", "lb", "ub", "c", "csense", "b","osenseStr",
  		"rxns","rxnNames", "rxnECNumbers","mets", "metNames"} /. ruleform;
  	(* Remove superfluous braces that result from the rules *)
  {modelname, description, ineq, maxmin} = {modelname, description, ineq, maxmin} /. {string_} :> string;
  {reactionIDs, reactionnames, reactionECnos} = 
  		Replace[{reactionIDs, reactionnames, reactionECnos}, t_ /; ArrayQ[t] :> Flatten[t],{1}];
  {metIDs, metnames} = Replace[{metIDs, metnames}, t_ /; ArrayQ[t] :> Flatten[t], {1}];
  modelname= If[modelname=="modelID", FileNameTake[file], modelname];
  description=If[description == "description", "", " : "<>description];
  ModelName = Species<>" - " <> modelname <> description;
  maxmin=Which[StringContainsQ[maxmin, "max", IgnoreCase -> True], -1, 
 	StringContainsQ[maxmin, "min", IgnoreCase -> True], 1, True, -1];
  {cons, vars} =Dimensions@S; 
  
  (* The following gives precedence to reaction IDs/metabolite IDs; 
  		simply rearrange ordering if e.g. full names are preferred *)
  ReactionNames = Which[Length@reactionIDs > 0, Flatten@reactionIDs, Length@reactionECnos > 0, Flatten@reactionECnos,
  	Length@reactionnames > 0, Flatten@reactionnames, True, ConstantArray["NoName",vars]  ];
  MetabNames = Which[ Length@metIDs > 0, Flatten@metIDs, Length@metnames > 0, Flatten@metnames,
  	 True, ConstantArray["NoName",cons]  ];
  externals = Flatten@Position[
   metIDs, _?(StringContainsQ[#, outsuffix ~~ EndOfString] &), Heads -> False];	
  If[externals=={}, externals = Flatten@Position[
   metnames, _?(StringContainsQ[#, outsuffix ~~ EndOfString] &), Heads -> False]]; 
  bounds = Transpose[Flatten /@ {lowers, uppers}];
  Object = Transpose@Object; 
  vals = First@Transpose@vals;
  (* If the csense item is missing, assume all constraints to be equals 
  		Also, provide for the case that ineq comes through as a single string 
  		of characters instead of a list of characters *)
(* Print[{"before processing, ineq =",ineq}]; *)
 If[StringQ[ineq] && ineq=="csense", ineq = ConstantArray[0,Length[vals]],
  				If[StringQ[ineq], ineq = StringPartition[ineq, 1]]];
  ineq = ineq /. {"E" -> 0, "L" -> -1, "G" -> 1};
(* Print[{"after processing, ineq =",ineq}]; *)
  Svals = Transpose[{vals, ineq}];
  If[Svals != ConstantArray[{0, 0}, Length[S]], 
   Print["WARNING: The loaded model contains inequalities and/or 
nonzero RHS vector. The SSK calculation currently does not provide for that! "]];
  If[printresult,Print["Dimensions of imported stoichiometry, bounds and objective arrays: " <> 
  	ToString[Dimensions /@ {S, bounds, Object}]]];
  objectcount=First@Dimensions[Object];
  If[objectcount==1,Object=First@Object;
  If[printresult,Print["The objective vector selects contributions from the flux set " <> 
  	ToString[Flatten@Position[Object, _?(Abs[#] > 0 &)]]]],
  Print["The objective specification contains "<>ToString[objectcount]<>" objective vectors."]]; 
  If[printresult,Print["This objective is to be "<>If[maxmin==1,"minimized.","maximized."]]];

  FBAvector = ConstantArray[0., Length[Object]];
  ]
  
SetAttributes[SBMLReader, HoldRest];
SBMLReader[file_, S_, Svals_, bounds_, Object_, FBAvector_, 
  ReactionNames_, MetabNames_, maxmin_, printresult_ : False] :=
  (*This function reads input data from an XML file containing a metabolic model 
  specification according to SBML conventions.It provides for files using both 
  the the FBC extension to SBML Level 3 and older files that use LOWER_BOUND,UPPER_BOUND 
  and OBJECTIVE_COEFFICIENT parameters in each reaction spec.*)
 Block[{rawdata, FBCprotocol = True, uri, datatags, usedtags, 
   missingtags, comparts, exter, metabolites, exlist, exempts, exnames, 
   reactions, reactants, reactantsat, products, productsat, cons, 
   vars, obj, obrules = {}, reactspeclist, reactspec, reaction, col, 
   xtracts, rules = {}, params, parameterules, lo, up, LowUp, obcof, 
   Srules, outfile, startime, timeused}, 
  If[printresult, Print["Data imported from file " <> file]];
  startime = TimeUsed[];
  progresscounter := Clock[5]; progressrange = Indeterminate;
  (*Check if the file uses the Flux Balance Constraints extension of SBML*)
  rawdata = Import[file, "XML"];
  uri = Cases[rawdata, 
    XMLElement["sbml", {___, {_, "fbc"} -> ur_, ___}, _] -> ur, Infinity];
  uri = If[uri == {}, FBCprotocol = False; "FBC not used", First@uri];
  If[printresult, Print["Flux Balance Constraints protocol: " <> uri]];
  (*Print[{"Time to import ",TimeUsed[]-startime}];
  timeused=TimeUsed[]- startime;*)
  (*Check that all necessary XML elements are present*)
  datatags = Import[file, {"XML", "Tags"}];
  usedtags = {"model", "species", "speciesReference", "reaction", 
    "listOfReactants", "listOfProducts", "listOfParameters","parameter", "compartment"};
  (*Note that the tag fbc:fluxObjective, while used if present,
  may be missing in a simple cone model without defined objective*)
  If[FBCprotocol,  
   usedtags = Join[usedtags, {"listOfObjectives", "objective"(*,"fluxObjective"*)}]];
   missingtags = Complement[usedtags, datatags];
  If[missingtags != {}, 
   MessageDialog["SBML input failed - the following tags were not found in the \
file: \n" <> ToString[missingtags]]; Abort[]];
  (*Extract naming data from the file.*)
  ModelName = FirstCase[rawdata, 
    XMLElement["model", {___,"id" -> id_, ___, ("name" -> nam_) ..., ___}, _] ->
     {id, nam}, {"NoName"}, Infinity];
  ModelName = Species <> " - " <> StringJoin @@ Riffle[ModelName, " : "];
  (* Find the id, "exter", of the compartment to be considered extracellular.
  	Look for the identifying substring "exterior" first in the compartment name, 
  		but if that fails in the compartment id. *)
  comparts = Cases[rawdata, 
    XMLElement["compartment", {OrderlessPatternSequence[
        "id" -> compid_, ("name" -> compname_) ..., ___]}, _] -> 
        {compid, compname}, Infinity];
   exter = FirstCase[comparts, {compid_, _?(StringContainsQ[#, exterior, 
          IgnoreCase -> True] &)} -> compid, "Failed"];
  If[exter == "Failed", exter = FirstCase[comparts, 
  	{compid_?(StringContainsQ[#, exterior, IgnoreCase -> True] &), ___} -> 
  	  compid, "Failed"]];
  If[exter == "Failed", MessageDialog[
    "SBMLReader identified compartments with id's and names\n " <> 
     ToString[comparts] <> "\n but could not recognize an external compartment."]];
  
  metabolites = Cases[rawdata, 
    XMLElement["species", {___, "id" -> ecode_, ___, "name" -> metab_, ___}, _] -> 
    	{ecode, metab}, Infinity];
  cons = Length@metabolites;
  PadRight[metabolites, {cons, 2}, "No name"];
  (*To return metabolite names instead of ID's,change to[[All, 2]] in the next line*)
  MetabNames = metabolites[[All, 1]];
  (* Set up lists of metabolite numbers that are external, and those explicitly exempted from
  	flux balance by boundaryCondition = true.*)
  exlist = Cases[rawdata, 
    XMLElement["species", {OrderlessPatternSequence["compartment" -> cid_, 
        "boundaryCondition" -> bc_, ___]}, _] -> {cid, bc}, Infinity];
  externals = Flatten@Position[exlist[[All, 1]], exter];
  exempts = Position[exlist[[All, 2]], "true"];
  
  reactspeclist = Cases[rawdata, XMLElement["reaction", _, _], Infinity];
  vars = Length@reactspeclist;
  If[cons == 0 || vars == 0, 
   MessageDialog["Invalid SBML input - the number of metabolites and reactions \
found were " <> ToString[{cons, vars}]]; Abort[]];
  (* Recovery of the reversible attribute cancelled because many models set it to true
  	for reversed, but unidirectional, reactions, nstead of bidirectional as per SBML spec. *)
  reactions = Map[Join[Cases[#, ("id" -> id_) -> id, 2],(*Cases[#,("reversible"-> rev_)->rev,2],*)
  	Cases[#, ("name" -> nam_) -> nam, 2]] &, reactspeclist];
  reactions = PadRight[reactions, {vars, 2}, "No name"];
  (*reversibles=reactions[[All,2]];
  reactions=reactions[[All,{1,3}]];*)
  (*To return reaction names instead of ID's,change to[[All,2]] in the next line*)
  ReactionNames = reactions[[All, 2]];
  (*Print[{"Time to read name lists ",TimeUsed[]-startime-timeused}];
  timeused=TimeUsed[]-startime;*)
  
  (*Process the objective section.*)
  If[FBCprotocol, obj = Cases[rawdata, XMLElement[{uri, "objective"}, _, _], Infinity];
   maxmin = Cases[obj, 
     XMLElement[{uri, "objective"}, {OrderlessPatternSequence[{uri, "type"} -> 
          typ_, ___]}, _] -> typ, Infinity];
   maxmin = If[obj == {}, "max", First@maxmin];
   obj = Cases[obj, 
     XMLElement[{uri,"fluxObjective"}, {OrderlessPatternSequence[{uri, "reaction"} -> 
     	reac_, {uri, "coefficient"} -> coef_, ___]}, _] -> {reac, coef}, Infinity];
   obrules = Map[Rule @@ # &, 
     Map[Replace[#, # -> {First@Flatten@Position[reactions[[All, 1]], #[[1]]], 
          StringToDouble[#[[2]]]}] &, obj]], maxmin = "max"];
  maxmin = Which[StringContainsQ[maxmin, "max", IgnoreCase -> True], -1, 
    StringContainsQ[maxmin, "min", IgnoreCase -> True], 1, True, -1];
  (*For FBC,the flux bounds are in a global list of parameters, not with each reaction.*)
  If[FBCprotocol, params = Cases[rawdata, XMLElement["listOfParameters", _, _], Infinity];
   parameterules = Cases[params, 
     XMLElement["parameter", {OrderlessPatternSequence["id" -> id_, 
         "value" -> val_, ___]}, _] -> (id -> val), Infinity]];
  (*Process each reaction in turn.The stoichiometry coefs are converted to rules to give 
  the entries in the applicable column of the S matrix,taking default value as 1. 
  In the old format, flux bounds and objective contribution are also read for each reaction.*)
  progressrange = {0, vars};
  xtracts = Reap[Do[progresscounter = col;
     reactspec = reactspeclist[[col]];
     reactants = Cases[FirstCase[reactspec, 
        XMLElement["listOfReactants", _, _], {}, Infinity], 
       XMLElement[ "speciesReference", {OrderlessPatternSequence[
           "species" -> metab_, ("stoichiometry" -> stoi_) ..., ___]}, _] ->
            {metab, stoi}, Infinity];
     If[reactants != {}, 
      reactants = PadRight[reactants, {Automatic, 2}, 1];
      reactants = Map[ReplaceAt[#, val_String :> StringToDouble[val], 2] &, reactants];
      reactantsat = Flatten[Position[metabolites[[All, 1]], #] & /@ reactants[[All, 1]]];
      reactants[[All, 1]] = reactantsat;
      rules = Map[{#[[1]], col} -> -#[[2]] &, reactants]];
     products = Cases[FirstCase[reactspec, 
        XMLElement["listOfProducts", _, _], {}, Infinity], 
       XMLElement[ "speciesReference", {OrderlessPatternSequence[
           "species" -> metab_, ("stoichiometry" -> stoi_) ..., ___]}, _] ->
            {metab, stoi}, Infinity];
     If[products != {}, 
      products = PadRight[products, {Automatic, 2}, 1];
      products = Map[ReplaceAt[#, val_String :> ToExpression[val], 2] &, products];
      productsat = Flatten[Position[metabolites[[All, 1]], #] & /@products[[All, 1]]];
      products[[All, 1]] = productsat;
      rules = Join[rules, Map[{#[[1]], col} -> #[[2]] &, products]]];
     If[reactants == {} && products == {}, 
      MessageDialog["Failure to parse reaction " <> ToString[reactions[[col, 1]] <> 
          "; neither reactants nor products could be identified."]];
      Abort[]];
     Sow[rules, r];
     
     LowUp = If[FBCprotocol, 
       FirstCase[reactspec, {OrderlessPatternSequence[{uri, 
              "upperFluxBound"} -> ub_, {uri, "lowerFluxBound"} -> 
             lb_, ___]} -> {lb, ub}, {"-Infinity", "Infinity"}, Infinity] /. parameterules, 
       params = Cases[reactspec, XMLElement["listOfParameters", _, _], Infinity];
       params = params /. {("id" -> id_) -> ("id" -> ToUpperCase[id])};
       lo = Cases[params, 
         XMLElement["parameter", {OrderlessPatternSequence[
             "id" -> "LOWER_BOUND" | "LOWERBOUND", 
             "value" -> bound_, ___]}, _] -> bound, Infinity];
       lo = If[lo == {}, {"-Infinity"}, lo];
       up = Cases[params, 
         XMLElement["parameter", {OrderlessPatternSequence[
             "id" -> "UPPER_BOUND" | "UPPERBOUND", 
             "value" -> bound_, ___]}, _] -> bound, Infinity];
       up = If[up == {}, {"Infinity"}, up];
       Join[lo, up]];
     Sow[LowUp, lu];
     
     obcof = Cases[params, 
       XMLElement["parameter", {OrderlessPatternSequence[
           "id" -> "OBJECTIVE_COEFFICIENT", "value" -> ob_, ___]}, _] -> ob, Infinity];
     obcof = If[obcof == {}, 0, StringToDouble@First@obcof];
     If[Abs@obcof > 0, Sow[col -> obcof, ob]];, {col, vars}], {r, lu, ob}];
  (*Print[{"Time to finish reaction loop ",TimeUsed[]-startime- timeused}];
  timeused=TimeUsed[]-startime;*)
  xtracts = xtracts[[2]];
  Srules = Flatten@xtracts[[1]];
  S = SparseArray[Srules, {cons, vars}];
  (*Eliminate any rows that correspond to a listed metabolite that \
does not participate in any reaction, or that are exempt from flux balance *)
  exempts = Join[Position[Normal@S, {(0 | 0.) ..}, 1], exempts];
  S = Delete[S, exempts];
  exnames = MetabNames[[externals]];
  MetabNames = Delete[MetabNames, exempts];
  (* relocate externals according to the new metabolite list, i.e. row numbers of S *)
  externals = Flatten[Position[MetabNames, #] & /@ exnames];
  cons = Length@S;
  Svals = ConstantArray[{0, 0}, cons];
  bounds = Map[StringToDouble[#] &, Flatten[xtracts[[2]], 1], {2}];
  obrules = If[xtracts[[3]] == {}, obrules, Flatten@xtracts[[3]]];
  Object = ReplacePart[ConstantArray[0., vars], obrules];
  FBAvector = ConstantArray[0., vars];
  If[printresult, 
   Print["Dimensions of imported stoichiometry, values, bounds, objective and FBA flux arrays: " <>
   	 ToString[Dimensions /@ {S, Svals, bounds, Object, FBAvector}]]];
  outfile = StringReplace[file, "xml" | "sbml" -> "m"];
  (*Export[outfile,{ModelName,S,Svals,bounds,Object,FBAvector,ReactionNames,MetabNames,maxmin},"List"];*)
  (*Print[{"Time to export ",TimeUsed[]-startime-timeused}];
  timeused=TimeUsed[]-startime;*)
  ]
  
CheckBalance[Stoichio_, Bounds_, Externs_, Printresult_ : False] :=
 (* Function to set up a list of external metabolites which may be \
exempted from the flux balance condition in order to compensate for \
the omission or inefffectiveness of source or sink (i.e. exchange) \
reactions in the model specification. *)
 Module[{stoichio, rows, cols, blockers, exblockers, inblockers, singletons, dirfix, SfSigns, zerows,
   exchangers, blocked, blockedexchangers, singleblockers, singletargets, kickout, notexempt,
   exemptions, exchangedmetabs, exnonexch, intexch},
  (*startime=TimeUsed[];*)
  stoichio = Normal[Stoichio];
  {rows, cols} = Dimensions@stoichio;
  (* Assign the direction value Infinity to reversible reactions. 
  Then when multiplying this by  the stoichio coef,  only 3 values will result: = +/-Infinity or Indeterminate. 
  Indeterminate is the result when the coef is 0, so replace its occurrences by 0.*)
  dirfix = Which[#[[1]] < #[[2]] <= 0 , -1, #[[2]] > #[[1]] >= 0., 1,
  				 #[[1]] == #[[2]] == 0., 0, True, Infinity] & /@ Bounds;
  (* SfSigns is a digitized version of S with values -1,  0 or 1 to indicate if a metabolite is 
  taken up, not inlvolved, or produced by each reaction, and +-Infinity if either.  *)
  (*Print["Time point = "<>ToString[TimeUsed[]-startime]];*)
  SfSigns = Map[Quiet[(Sign[#]*dirfix) /. Indeterminate -> 0] &, stoichio];
  (* NOTE: 
  The previous line takes up about 60% of the processing time for this function.
  	This may be because of the 0*Infinity = Indeterminate generating error messages etc.
  	 The following line avoids this by explcitly testing for a zero before multiplying
  		but it turns out to be only slightly faster. 
  Stick with the simpler version. *)
  (*SfSigns=Map[MapThread[If[#1==0.,0,#1*#2]&,{Sign@#,dirfix}]&, stoichio];*)
  (*Print["Time point = "<>ToString[TimeUsed[]-startime]];*)
  
  (* Determine blocking rows, i.e. metabolites *)
  blockers = DeleteDuplicates@Flatten@Last@
      Reap[Do[If[! MemberQ[SfSigns[[row]], Infinity | -Infinity],
         If[MatchQ[Sign@SfSigns[[row]], {(0 | -1) ..}], Sow[row]];
         If[MatchQ[Sign@SfSigns[[row]], {(0 | 1) ..}], Sow[row]];
         ],
        {row, rows}]];
  zerows = Flatten@Position[SfSigns, {0 ..}];
  singletons = Flatten@Position[Map[Count[#, Except[0]] &, SfSigns], 1];
  (* SfSigns that are all-zero because of 0,0 bounds are picked up. Eliminate them *)
  blockers = Union[Complement[blockers, zerows], singletons];
  exblockers = Intersection[blockers, Externs];
  inblockers = Complement[blockers, Externs];
  (*Print["blockers ",blockers ];*)
  
  (* Find the blocked reactions, the exchange reactions, 
  and then isolate blocked exchanges and allocate as external/internal *)
  blocked = Flatten@Position[Total@Abs@SfSigns[[blockers]], Except[0], 1, Heads -> False]; 
  exchangers = Flatten@Last@Reap[Do[(*Print[Sign@SfSigns[[row]]];*)
       If[MatchQ[Sign@stoichio[[All, col]], {(0 | -1) ..}], Sow[col]];
       If[MatchQ[Sign@stoichio[[All, col]], {(0 | 1) ..}],  Sow[col]];
       , {col, cols}]];
  blockedexchangers = Intersection[exchangers, blocked];
  (*Print["exchangers",exchangers];*)
  
  (*Look at the external blockers that are singletons, and find which reactions they target.
  Omit any that target an exchange reaction from the ones to be exempted. *)
  singleblockers = Intersection[exblockers, singletons]; 
  singletargets =  Flatten@Map[Position[#, Except[0], 1, Heads -> False] &, SfSigns[[singleblockers]]]; 
  kickout = Map[MemberQ[blockedexchangers, #] &, singletargets]; 
  notexempt = Pick[singleblockers, kickout];
  exemptions = Complement[exblockers, notexempt];
  
  (* Check which metabolites have explicit sinks/sources *)
  exchangedmetabs = DeleteDuplicates@Flatten@
  	Map[Position[stoichio[[All, #]], Except[0 | 0.], 1, Heads -> False] &, exchangers]; 
  exnonexch = Complement[Externs, exchangedmetabs];
  intexch = Complement[exchangedmetabs, Externs];
  
  If[Length@exnonexch > 0.5*Length@Externs, 
   Print[Style["WARNING!\n The model appears incomplete; it provides no \
exchange reactions for " <> ToString[Length@exnonexch] <> " out of " <>
       ToString[Length@Externs] <> 
      " external metabolites. \nThis may restrict the solution space \
or even cause the model to be infeasible. \nConsider exempting \
non-buffered externals from flux balance.", Red]]];
  If[Printresult,
   Print[Style["Sink and source analysis of stoichiometry matrix.", "Subsubsection", 16]];
   Print["This model has " <> ToString[Length@Externs] <> " metabolites identified as external, and " <> 
     ToString[Length@exchangers] <> " exchange reactions."];
   If[Length@blockedexchangers > 0, 
    Print[ToString[Length@blockedexchangers] <> " exchange reactions are blocked, because they\n"<>
    	"involve a metabolite for which the flows cannot be balanced. "];
    If[Length@notexempt > 0,
     Print["Such blocking is appropriate for the " <> ToString[Length@notexempt] <> 
       " exchange reactions that target\n the following metabolites, \
since they are not involved in the network: \n" <> ToString[MetaboliteNames[[notexempt]]]]];
	];
   If[Length@exemptions == 0, 
   	Print["All external metabolites have either explicit sink/source reactions, or bidirectional\n"<>
   		" exchange with the enivironment. Ticking the exemptions box in Stage 2 will have no effect."],
    Print["A total of " <> ToString[Length@exemptions] <> 
      " external metabolites are left without a viable source or sink, namely \n" <>
       ToString[MetaboliteNames[[exemptions]]]];
    Print["Ticking the non-buffered exemption option in Stage 2 provides an alternative way to\n"<>
    	"allow free, unbounded exchange of these external metabolites with the environment. "]];
   If[Length@intexch > 0, 
    Print["Environmental exchange is provided for the following internal metabolites: ", 
      MetaboliteNames[[Complement[intexch, inblockers]]]];
    ];
   If[Length@inblockers > 0,
    Print[ "There are also " <> ToString[Length@inblockers] <> " non-buffered internal metabolites, " <> 
      ToString[Length[Intersection[inblockers, singletons]]] <> 
      " of which \neach only participates in a single reaction.\n"<>
      "All reactions that involve them will not be exempted and remain blocked \n"<>
      "in order to preserve the metabolic steady state."]]; 
   Print[Style["_____________________________________________________________", 
     "Subsubsection", 16]];
   ];
  exemptions]

Shakeup[] := Module[{rows, cols, roworder, colorder},
  {rows, cols} = Dimensions[S];
  (* Shuffle rows and columns of the stoichiometry and related matrices 
  	into a randomly different order. *)
  roworder = RandomSample[Range[rows]];
  colorder = RandomSample[Range[cols]];
  ReactionNames = ReactionNames[[colorder]];
  MetaboliteNames = MetaboliteNames[[roworder]];
  (* The externals and exemptions lists are row numbers in original ordering;
  	 adjust them to the new order  *)
  externals = Flatten[Position[roworder, #] & /@ externals];
  exemptions = Flatten[Position[roworder, #] & /@ exemptions];
  S = S[[roworder, colorder]];
  Svals = Svals[[roworder]];
  bounds = bounds[[colorder]];
  objectselector = objectselector[[colorder]];
  (* Reinitialize all data *)
  rawfeasibles = {}; boundedflux = {}; openflux = {};
  augment = {{}}; lowers = {}; uppers = {}; feasiblepoints = {}; 
  ]

End[] (* End Private Context *)

EndPackage[]
```
