## Supplementary material for "The FBA Solution Space Kernel – Introduction and Illustrative Examples": SSK software manual and code: FacetTesting.m

```
(* Wolfram Language Package *)

(* COPYRIGHT
					© Copyright 2022 Wynand Verwoerd

This file is part of SSKernel.

The SSKernel program is free software; you can redistribute it and/or modify it under the terms of the GNU General Public License as published by the Free Software Foundation; either version 3 of the License, or (at your option) any later version.
 
This program is distributed in the hope that it will be useful, but WITHOUT ANY WARRANTY; without even the implied warranty of MERCHANTABILITY or FITNESS FOR A PARTICULAR PURPOSE . See the GNU General Public License for more details.
 
You should have received a copy of the GNU General Public License along with this program . If not, see http://www.gnu.org/licenses/
*)

BeginPackage["FacetTesting`", { "Configuration`", "RayFinding`"}]
(* Exported symbols added here with SymbolName::usage *)  
BoundedQ::usage="Tests boundedness by inspecting ray matrix, then LP if inconclusive "
FeasibleBoundedQ::usage="Combined testing of feasibility and boundedness "
OrthoRayQ::usage=""

Begin["`Private`"] (* Begin Private Context *) 

SetAttributes[BoundedQ, HoldFirst];
BoundedQ[RayMat_, constraints_, facet_, rayfree_: {}, 
  printresult_: False] :=
 (* This function sets slacks only on non-facet constraints and if 
there is an LP solution, this is added to Raymat. 
 Otherwise the facet is bounded and it returns "True".
 The optional argument rayfree is a list of nonfacet rayfree 
constraints inherited from lower level facets *)
 Module[{cons, vars, raysinfacet, noslacks, conslack, slackcount, 
   slackcon, SObjective, SXConstraints, SXRange, SValues, SX, fullSX, 
   bounded, tol = 0.001},
(* For a strict test on boundedness, a slack tolerance value such as 0.001 is needed.
This is because otherwise a false positive can result from failure of the LP test to find
a ray solution. *)

  (* Check the ray matrix first for any rays that are in the facet *)
  raysinfacet = Count[Total[RayMat[[All, facet]], {2}], _?(# < tol &)];
  If[raysinfacet > 0, 
   If[printresult, 
    Print["Facet " <> ToString[facet] <> " is unbounded because of counterexample in ray matrix."]]; 
   Return[False]];
   
   (* Otherwise do LP testing omitting slacks for facet constraints *)
  {cons, vars} = Dimensions[constraints];
  noslacks = Union[facet, rayfree];
  (* To keep track of which constraints have free slacks, use two arrays: 
  conslack is a list of all constraints that have an allocated slack variable, i.e. 
  conslack[[slack]] is the constraint for given slack variable no "slack";
  slackcon is a kind of inverse of this, i.e. slackcon[[con]] is the 
  slack var for given constraint no "con", or 0 if none *)
  conslack = Complement[Range[cons], noslacks];
  slackcount = Length[conslack];
  slackcon = Range[slackcount];
  If[Length[noslacks] > 0, 
   slackcon = Insert[slackcon, 0, 
     Partition[noslacks - Range[Length[noslacks]] + 1, 1]]];
  (*
  Print["The following constraints have slack variables: "<>ToString[conslack]];
  Print["The slack that belongs to each constraint is: "<>ToString[ slackcon]];
  *)
  SValues = ConstantArray[{0., 0}, cons];
  SXConstraints = Join[IdentityMatrix[cons][[All, conslack]], constraints, 2];
  SObjective = Join[ConstantArray[1, slackcount], ConstantArray[0, vars]];
  (* Set range constraints for MINIMISATION *)
  SXRange = Join[ConstantArray[{0., Infinity}, slackcount], ConstantArray[{-Infinity, Infinity}, vars]];
  (* Minimisation requires removing the trivial solution by setting a minimum objective value,
  say 0.5 . This value is arbitrary, as long as it is significantly nonzero, 
  because scaling of the SX vector is fixed later anyway.   Add a dummy constraint to fix this *)
  SXConstraints = Append[SXConstraints, SObjective];
  SValues = Append[SValues, {0.5, 1}];
  (*Print[{Dimensions[SObjective],Dimensions[SXConstraints],Dimensions[SValues],Dimensions[SXRange]}];*)
  SX = Quiet[
    Check[LinearProgramming[SObjective, SXConstraints, SValues, SXRange, Method -> "Simplex", Tolerance -> tol], 
     "No Solution", {LinearProgramming::lpsnf}], {LinearProgramming::lpsnf}];
     bounded = If[VectorQ[SX, NumberQ],
    (* Normalize the rays to avoid problems with rank calculation ? *)
    SX = SX/Norm[SX[[-vars ;;]]];
    (* Reintroduce the skipped "noslacks" zero slack values *)
    fullSX = Join[ConstantArray[0, cons], SX[[slackcount + 1 ;;]]];
    fullSX[[conslack]] = SX[[;; slackcount]];
    AppendTo[RayMat, Chop[fullSX]]; False, 
    True];
  
  If[printresult,
   If [bounded, Print["Facet " <> ToString[facet] <> " is bounded "], 
     Print["Facet " <> ToString[facet] <> 
       " is unbounded and yields a new ray."]];
   ];
  bounded
  ]

SetAttributes[FeasibleBoundedQ, HoldFirst]; 
FeasibleBoundedQ[RayMat_, constraints_, values_, FacetCols_, 
  DoBounded_: True] := 
  Module[{cons, vars, NonFacetCols, Fcon, NFcon, Finv, Phi, 
      Fproj, exist, SPhi, Kset, facetrays, Gconstraints, 
      Objective, Values, XRange, X, feasible, rayfree, bounded, 
      tol = 0.00001},
  (* Here a fairly strict tolerance 0.00001 to avoid false positive on feasibility, 
  but not too strict as that may produce false negative .
  Note that final bounded test is done by call to BoundedQ which sets its own tolerance *)
    {cons, vars} = Dimensions[constraints];
    
    (* The full polytope is assumed to be feasible but unbounded *)
    If[FacetCols == {}, Return[{True, False}]];
    (* Separate facet and non-facet constraints, 
    then solve facet constraints with pseudoinverse *)
    NonFacetCols = Complement[Range[cons], FacetCols];
    Fcon = constraints[[FacetCols]]; 
    NFcon = constraints[[NonFacetCols]];
    (* A non-empty intersection of facet constraint hyperplanes exists if a 
  pseudoinverse can be calculated and it gives a solution for the 
  corresponding values vector; else facet is infeasible. *)
    Finv = Quiet@Check[PseudoInverse[Fcon], Return[{False, False}]];
    Phi = Finv.values[[FacetCols]];
    exist = Norm[Fcon.Phi - values[[FacetCols]]] < tol;
    If[! exist,(*Print["Infeasible because pseudoinverse gives no intersection"];*)
      Return[{False, False}]];
    
    (* Prepare slack values for the facet hyperplane reference point Phi *)
    SPhi = values[[NonFacetCols]] - NFcon.Phi;(*Print[SPhi];*)
    Kset = Extract[NonFacetCols, 
        Position[Chop[SPhi, tol], _?(# < 0 &)]];(*Print[Kset];*)
  
  (* Check the ray matrix *)  
    facetrays = Flatten@Position[Total[RayMat[[All, FacetCols]], {2}], _?(# < tol &)];
    (*Print[facetrays];*)
    
    (* Classify the cases by inspection *)
    {feasible, rayfree} = Which[
        facetrays == {} && Kset == {}, {True, True},	(* Case 1a *)
        facetrays == {}, {None, True},                  (* Case 1b *)
        Kset == {}, {True, False},                      (* Case 2 *)
        Count[Total[RayMat[[facetrays, Kset]]],
          	 _?(# < tol &)] == 0, {True, False},      	(* Case 3 *)
        True, {None, False}                             (* Case 4 *)
        ];
    
    (* Check infeasibility by LP testing *)
    If[feasible == None, (*Print["Doing the LP feasibility test"];*)
      (* Set up the reduced LP feasibility test *)
      Fproj = IdentityMatrix[vars] - Finv.Fcon;
      Gconstraints = Chop[NFcon.Fproj];
      {cons, vars} = Dimensions[Gconstraints];
      Objective = ConstantArray[0, vars];
      Values = Transpose[{SPhi, ConstantArray[-1, cons]}];
      XRange = ConstantArray[{-Infinity, Infinity}, vars];
      (* Simplex works best; 
      Interior Point and RevisedSimplex sometimes fail to converge *)
      X = Quiet[          
     Check[LinearProgramming[Objective, Gconstraints, Values, XRange, 
              Method -> "Simplex", Tolerance -> tol],           
      "No Solution", {LinearProgramming::lpsnf}], {LinearProgramming::lpsnf}];
      (*X=LinearProgramming[SObjective,SXConstraints,SValues,SXRange,
      Method\[Rule]"Simplex",Tolerance\[Rule]tol];*)
      feasible = VectorQ[X, NumberQ]
      ];
    
    (* Feasibility is settled, now check boundedness *)
  bounded = If[DoBounded && feasible && rayfree, 
    BoundedQ[RayMat, constraints, FacetCols], 
        rayfree];
  If[! DoBounded, bounded = False];
    {feasible, bounded}
    ]
  
OrthoRayQ[RayMat_, constraints_, facet_, printresult_: False] :=
 (* This function tests the existence of a ray that is orthogonal to 
the facet specified as the intersection of the constraint hyperplanes 
listed in the argument facet *)
 Module[{cons, vars, FacetBasis, facetdim, rayoverlaps, orthorays, 
   SObjective, SXConstraints, SXRange, SValues, SX, rayfound, 
   tol = 0.00001},
  {cons, vars} = Dimensions[constraints];
  (* If the facet list is empty, we are dealing with the zero level facet
  	i.e. the polytope itself, and no ray can be orthognal to that*)
  FacetBasis = If[facet=={},{},NullSpace[constraints[[facet]],Tolerance-> tol]];
  If[FacetBasis=={},  If[printresult,
        Print["There is no orthogonal ray for facet " <> ToString[facet]]];
   		Return[False]];
  (* Check the ray matrix first for any rays that are orthogonal to the facet *)
  If[Length[RayMat] > 0,
   rayoverlaps = 
    Chop[RayMat[[All, cons + 1 ;;]].Transpose[FacetBasis], tol];
   orthorays = Count[Total[rayoverlaps, {2}], 0];
   If[orthorays > 0, 
    If[printresult, 
     Print["There are " <> ToString[orthorays] <>" rays in the ray matrix orthogonal to facet " <> 
       ToString[facet]]]; Return[True]];
   ];
  {facetdim, vars} = Dimensions[FacetBasis];
  FacetBasis = 
   Join[ConstantArray[0., {facetdim, cons}], FacetBasis, 2];
  SXConstraints = Join[IdentityMatrix[cons], constraints, 2];
  SXConstraints = Join[SXConstraints, FacetBasis];
  SValues = ConstantArray[{0., 0}, cons + facetdim];
  SObjective = Join[ConstantArray[1, cons], ConstantArray[0, vars]];
  (* Set range constraints for MAXIMISATION *)
  SXRange = 
   Join[ConstantArray[{0., Infinity}, cons], 
    ConstantArray[{-1, 1}, vars]];
  (* Simplex works best; Interior Point and RevisedSimplex sometimes fail to converge *)
  SX = Quiet[
    Check[LinearProgramming[-SObjective, SXConstraints, SValues, 
      SXRange, Method -> "Simplex", Tolerance -> LPtol], 
     "No Solution", {LinearProgramming::lpsnf}],{LinearProgramming::lpsnf}];
  (*Print[Chop@SX];*)
  If[SX == "No Solution", 
   If[printresult, Print[" Lin Prog failed to find a ray for slack variables " <> 
     ToString[Flatten@Position[SObjective, 1]]]]; Return[False]];
  (*Print["Objective value is "<>ToString[SX.SObjective]];*)
  rayfound = If[SX.SObjective > tol, True, False];
  If[printresult,
   If [rayfound, 
     Print["Facet " <> ToString[facet] <> " has an orthogonal ray."], 
     Print["There is no orthogonal ray for facet " <> 
       ToString[facet]]];
   ];
  rayfound
  ]

End[] (* End Private Context *)

EndPackage[]
```
