## Supplementary material for "The FBA Solution Space Kernel – Introduction and Illustrative Examples": SSK software manual and code: FBFSearch.m

```
(* Wolfram Language Package *)

(* COPYRIGHT
					© Copyright 2022 Wynand Verwoerd

This file is part of SSKernel.

The SSKernel program is free software; you can redistribute it and/or modify it under the terms of the GNU General Public License as published by the Free Software Foundation; either version 3 of the License, or (at your option) any later version.
 
This program is distributed in the hope that it will be useful, but WITHOUT ANY WARRANTY; without even the implied warranty of MERCHANTABILITY or FITNESS FOR A PARTICULAR PURPOSE . See the GNU General Public License for more details.
 
You should have received a copy of the GNU General Public License along with this program . If not, see http://www.gnu.org/licenses/
*)

BeginPackage["FBFSearch`", { "Configuration`","RayFinding`", "FacetTesting`"}]
(* Exported symbols added here with SymbolName::usage *)  
GreedySearch::usage="Randomised greedy search for feasible bounded facets based on ray matrix scoring "
FBFfinder::usage="Searches for FBF's by exhaustive tree search or semi-random sampling "
FBFsampler::usage="Does multiple greedy searches with backtracking for feasible bounded facets "
Backtrack::usage="Backtracks a given FBF to its lowest level bounded, feasible ancestor or BFBF"
TracebackSearch::usage="Simplified randomized greedy search for a single feasible bounded facet "
Progenitor::usage="Extract the progenitor facet listing from repeated random searches "
TreeSearch::usage="Systematic search for feasible bounded facets by traversing the facet tree"
MemberList::usage=" Find essential columns in the ray matrix and sort the remaining possibles "
SubFacet::usage="Determines if a given facet is a subfacet of the known BFBF's "
BranchAllocate::usage="Allocates the list of FBFs to branches of the tree resulting from given node ordering"

Begin["`Private`"] (* Begin Private Context *) 

(* SEPARATING ESSENTIALS, POSSIBLES AND DISPENSABLES, AND SORT BY RAY ELIMINATION score *)

MemberList[RayMat_, constraints_, values_] := 
  Module[{cons, vars, raycount, UnitS, InOutRayList, dispensables, 
      rowsum, singlerows, essentials, possibles, scores, scoreorder, 
      tol = 0.00000001}, {cons, vars} = Dimensions[constraints];
    raycount = Length[RayMat];
    UnitS = Unitize[RayMat[[All, ;; cons]], tol];
    (* Identify dispensable (PRF) columns as wel as those that are 
  essential,from all rows that have a single nonzero. 
    The rest are possibles; score and sort them.*)
    dispensables = Flatten@Position[Total[UnitS], 0];
    rowsum = Total[UnitS, {2}];
    singlerows = Flatten@Position[rowsum, 1];
    essentials = Last /@ Position[UnitS[[singlerows]], 1];
    possibles = Complement[Range[cons], essentials, dispensables];
    (* Set indicators for all rays that were not ruled out by Essentials *)
    InOutRayList = Map[1 - Unitize@Total[#[[essentials]]] &, UnitS];
    (* Sort possibles according to their power to eliminate known rays *)
    scores = Map[InOutRayList.UnitS[[All, #]] &, possibles];
    scoreorder = Ordering[scores, All, Greater];
    possibles = possibles[[scoreorder]]; 
    scores = scores[[scoreorder]];
  (*Print[
    "Facet columns selected from the following list of possibles, in \
descending order of score:"];
    Print[Transpose[{possibles,scores}]];*)
    {possibles, essentials}
    ]

(* THE RANDOMISED GREEDY SEARCH *)

SetAttributes[GreedySearch, HoldAll];
GreedySearch[RayMat_, constraints_, values_, BFset_,  memberlist_, 
  PrintResult_: False] := 
  Module[{cons, vars, raycount, UnitS, possibles, essentials, 
   esscount, posscount,
    	InOutRayList, scores, scoreorder, scoreseq,  
   mixscore, mixcount, mixorder, mixupto, topmix, facetcols, next, 
   feasible = True, bounded = False, trialfacet, infeasibles = {},   
   tol = 0.000001},
     (*This function uses a moderately greedy heuristic to get a somewhat random,feasible bounded facet,
    based on scoring according to the ray matrix passed to it in the first argument*)
    {cons, vars} = Dimensions[constraints];
    raycount = Length[RayMat];
    {possibles, essentials} = memberlist;
    esscount = Length[essentials]; posscount = Length[possibles];
    If[PrintResult,
    Print["There are "<>ToString[esscount]<>
  " essential columns, leaving "<>
    ToString[posscount]<>" columns that can be allocated."]];
  
  (* Calculate the score for each possible, 
  then shuffle the ones on the highest score levels randomly *)
  facetcols = essentials; 
   UnitS = Unitize[RayMat[[All, ;; cons]], tol];
  InOutRayList = Map[1 - Unitize@Total[#[[facetcols]]] &, UnitS];
  scores = Map[InOutRayList.UnitS[[All, #]] &, possibles];
  scoreorder = Ordering[scores, All, GreaterEqual];
  possibles = possibles[[scoreorder]]; scores = scores[[scoreorder]];
  scoreseq = DeleteDuplicates[scores];
  (* Mix at most the top "mixfraction" score levels, but keep the 2 lowest levels unmixed. *)
  topmix =  Min[IntegerPart[mixfraction*Length[scoreseq]], Length[scoreseq] - 1];
  topmix = Max[topmix, 1];
  mixscore = scoreseq[[topmix]];
  mixupto = Last@Flatten@Position[scores, mixscore];
  mixcount = Count[scores, _?(# >= mixscore &)];
  (*Print[{topmix,mixscore,mixupto,mixcount}];*)
  mixorder = RandomSample[Range[mixcount], mixcount];
  mixorder = Join[mixorder, Range[mixcount + 1, Length[possibles]]];
  possibles = possibles[[mixorder]]; scores = scores[[mixorder]];
    next = 1;
    While[next <= Length[possibles] > 0 && scores[[next]] > 0, 
   If[MemberQ[infeasibles, possibles[[next]]], next++; Continue[]]; 
   trialfacet = Append[facetcols, possibles[[next]]];
   totalfacets++;
   If[SubFacet[trialfacet, BFset],
    (* Print["Subfacet found!"];*)
    next++; Continue[]];
      {feasible, bounded} = 
        FeasibleBoundedQ[RayMat, constraints, values, trialfacet, True];
      If[Length[RayMat] > raycount, raycount = Length[RayMat];
        If[PrintResult, 
          Print[" A new ray was added to the ray matrix at level " <> 
              ToString[Length[facetcols]]]];
        UnitS = Unitize[RayMat[[All, ;; cons]], tol];];
   
      Which[
    feasible && bounded, facetcols = trialfacet; Break[], 
    feasible,(*not bounded,progress to the next tree level*)
    facetcols = trialfacet; 
    InOutRayList = 
     Map[1 - Unitize@Total[#[[facetcols]]] &, UnitS];
     (*
     Print["I select column "<>ToString[possibles[[next]]]<>
        " at position "<>ToString[next]<>" with score "<>ToString[
    scores[[next]]]<>" leaving "<>ToString[Total[InOutRayList]]<>
    " rays in the facet."];
    *)
    possibles = Delete[possibles, next]; 
    scores = Map[InOutRayList.UnitS[[All, #]] &, possibles];
    scoreorder = Ordering[scores, All, GreaterEqual];
    possibles = possibles[[scoreorder]]; 
    scores = scores[[scoreorder]];
    
    If[Length[facetcols] - esscount <= mixupto,
     scoreseq = DeleteDuplicates[scores];
     topmix = 
      Min[IntegerPart[mixfraction*Length[scoreseq]], 
       Length[scoreseq] - 2];
     topmix = Max[topmix, 1];
     mixscore = scoreseq[[topmix]];
     mixcount = Count[scores, _?(# >= mixscore &)];
     (*Print["At level "<>ToString[Length[facetcols]]<>" there are "<>
     ToString[mixcount]<>" possibles available for the random choice."];*)
     mixorder = RandomSample[Range[mixcount], mixcount];
     mixorder = Join[mixorder, Range[mixcount + 1, Length[possibles]]];
     possibles = possibles[[mixorder]];
     scores = scores[[mixorder]];
     (*Print[Transpose[{possibles,scores}][[1;;5]]];*)
     ];
    next = 1;,
    
    True,
    (*Print["Skip possible "<>ToString[possibles[[next]]]<>" as it is infeasible."];*)
    (*not feasible, traverse horzontally to the next branch*)
    infeasibles = 
     DeleteDuplicates@Append[infeasibles, possibles[[next]]];
    next++]
   ];
  
  (* Test the facetcols that came out of the loop to confirm boundedness. 
  Without the test, false positives (i.e., 
  an unbounded facet is returned as if it is bounded) are sometimes 
observed. This is unexplained. 
  The only way that the loop can be exited with a "bounded" result \
supposed to be the explicit Which case; 
  however the loop terminated otherwise, bounded should remain False. 
  So how can the test here give unbounded if it previously gave bounded? Granted, 
  the ray matrix may have been extended in the meantime. 
  But that is only supposed to increase the discrimination of the ray-
  based boundedness test; if it is positive, 
  FeasibleBoundedQ alaway confirms this with an LP test.
  Whatever, this works and should not cost much time!
   *)
    {feasible, bounded} =  
   FeasibleBoundedQ[RayMat, constraints, values, facetcols, True];
  
  (*Print[infeasibles];*)
    If[PrintResult, 
      If[bounded, Print[
     "GreedySearch found a feasible and bounded facet at level " <> 
            ToString[Length[facetcols]] <> " and with " <> 
            ToString[raycount] <> " rays."];
    Print[facetcols], 
    Print["GreedySearch terminating on an unbounded facet at level " <> 
            ToString[Length[facetcols]] <> ", because there remain " <> 
            ToString[Total[InOutRayList]] <> " rays and all " <> 
            ToString[Count[scores, _?Positive]] <> 
            " viable remaining columns fail to produce a new and feasible facet."]
    ]];
    If[bounded, facetcols, {}]]

SubFacet[facetlist_, BFset_] := 
  Block[{BFcount, facetlength, bflength, subfacet = False},
    (* If BFset contains any facet of which facetlist is a subfacet 
  (i.e., it is a superset) this function returns True, otherwise False *)
    BFcount = Length[BFset]; 
    facetlength = Length[facetlist]; 
    Do[bflength = Length[BFset[[i]]]; 
      If[facetlength >= bflength && 
     Length[Intersection[BFset[[i]], facetlist]] == bflength, 
        subfacet = True;(*Print[i];*)Break[]
        ], {i, BFcount}];
    subfacet]
    
SetAttributes[FBFfinder, HoldFirst];
FBFfinder[RayMat_, RayDim_, constraints_, values_, samplesize_, treelimit_, Printresult_: False] :=
(*This function does an exhaustive tree search for FBF's if it estimates that 
the mumber of facets is less than treelimit.
 Otherwise it does a randomised greedy search to find up to targetcount FBF's.
 Either way, the FBF's found are put in the BoundedFacets array and 
it returns True/False for whether the list is exhaustive, as well as 
the number of FBF's found or sampled respectively. *)
 Module[{possibles, essentials, posscount, esscount, cons, vars, 
 	particount, nodecount, levelcounts, minlevel, toplevel, maxlevel,
    constraintfreqs, participants, nonparticipants, treesearch, count},
  {cons, vars} = Dimensions[constraints];
  {possibles, essentials} = MemberList[RayMat, constraints, values];
  {posscount, esscount} = Length /@ {possibles, essentials};
  totalfacets = 0;
  (*In case the set of possibles is empty,there is only one BFBF, given by the list of essentials.*)
	(* Print[{"posscount,esscount",posscount,esscount}]; *)
  If[posscount == 0 , If[Printresult, 
   Print["Analysis of the ray matrix shows that all non-dispensable hyperplanes are essential,
          	so there is only a single BFBF which is its own progenitor."]];
   BoundedFacets = {essentials};
   FBFcount = 1; totalfacets = 1;
   Return[{True, 1}]];
  
  (* For progenitor search, no FBF's have been sampled. Then only do tree
  	search for up to 14 possibles, i.e. tree size max 16384 *)
  treesearch = If[FBFcount==0, 
  	If[posscount > 14, False,
  		minlevel = 1;	maxlevel = Infinity; 
  		possibles = Reverse@possibles; True],
  (* Else use existing BFBF sample to decide which search to use *)
  	particount=Length@DeleteDuplicates[Flatten@BoundedFacets];
  	particount=particount/(1 - 0.4 Exp[-0.055*FBFcount]);
  	levelcounts = Tally[Length /@ BoundedFacets];
  	toplevel = MinimalBy[levelcounts, 
   				Abs[First[#] - particount/2] &][[1, 1]];
	nodecount = N@Binomial[particount, toplevel];
	If[nodecount > treelimit, False,
		(* Rearrange possibles using ascending frequencies in known BFBF's*)
		constraintfreqs = SortBy[Tally[Join @@ BoundedFacets], Last];
		participants = Select[constraintfreqs[[All, 1]], ! MemberQ[essentials, #] &];
		nonparticipants = Reverse@Select[possibles, ! MemberQ[participants, #] &];
		possibles = Join[nonparticipants, participants];
		maxlevel = Max[levelcounts[[All, 1]]] + 2;
		minlevel = Min[levelcounts[[All, 1]]] - 2;
		If[Printresult,Print["Estimated tree nodes to visit: "<>ToString[nodecount]]];
		True]
  ];
  
  (* Call the appropriate FBF search. *)
  If[treesearch,
    progressrange={0,treelimit}; progresscounter=0;
    progresslabel= " Exhaustive tree search for FBF's"; 
    Print[Style[progresslabel, Blue, TextAlignment -> Center]];   
    count = TreeSearch[RayMat, constraints, values, {possibles, essentials}, 
     	{minlevel, maxlevel}, Infinity, treelimit, Printresult];
    progresslabel= " Tree search completed finding"<>ToString[count]<>" BFBF's."; 
    progresscounter=0;
  (*  Monitor[
     count = TreeSearch[RayMat, constraints, values, {Reverse@possibles, essentials}, 
     	{esscount+minposs, esscount+maxposs}, Infinity, Infinity, Printresult];,
     Labeled[ProgressIndicator[totalfacets/treesize], " Exhaustive tree search for FBF's"]];*)
     ,
     
    progresscounter=0; progressrange={0,samplesize}; 
    progresslabel= " Randomised greedy FBF sampling";
    Print[Style[progresslabel, Blue, TextAlignment -> Center]];
    count = FBFsampler[RayMat, constraints, values, {possibles, essentials}, samplesize, Printresult];
    progresslabel= " Greedy search completed finding" <>ToString[count]<>" BFBF's."; 
    progresscounter=0;

(*    Monitor[
     count = FBFsampler[RayMat, constraints, values, {possibles, essentials}, samplesize, Printresult];,
     Labeled[ProgressIndicator[FBFcount/samplesize], " Randomised greedy FBF sampling"]]; *)
     
    ];
  
  {treesearch, count}]

    

FBFsampler[RayMat_, constraints_, values_, memberlist_, targetcount_: Infinity, PrintResult_: False] := 
  Module[{possibles, essentials, rayma, lowfacet, backlevels,  
      countback = 0, count = 0, trialcount = 0, rejects = 0, 
   rejectlist = {0}, consecutive = 10},
    {possibles, essentials} = memberlist;
  (* Disable the greedyfails termination test if an explicit targetcount was given *)
  (* If[targetcount < Infinity, greedyfails = Infinity]; *)
  
    (* BoundedFacets = {};*) (*DISABLED FOR TESTING - MAY NEED TO BE RESTORED *)
    
    (*
    This function performs a sequence of semi-random greedy FBF searches, 
    and backtracks each to find its lowest level bounded parent or BFBF. 
  Resulting BFBF's are stored in the global array BoundedFacets.
  The sampling terminates when either the sample size has reached the value specified by "targetcount", 
  or the greedy search has failed "greedyfails" times  before finding the next FBF, 
  taken as average over the last "consecutive" FBF's it found.
    It returns the number of greedy trials done in total.
    *)
    While[count < targetcount && Mean[rejectlist] < greedyfails, trialcount++;
   (* GreedySearch and Backtrack both extend the ray matrix they are passed, as they find new rays. 
   But this extension seems counterproductive in the progenitor calculation as it reduces the success 
   rate of the greedy search and this is a limiting factor in traceback. 
   So just pass them a local disposable copy of the ray matrix *)
      rayma = RayMat;
      lowfacet = 
    GreedySearch[rayma, constraints, values, BoundedFacets, memberlist, False];
    countback = If[Length[lowfacet] > 0, 
     rejectlist = PadLeft[Append[rejectlist, rejects], consecutive]; 
     rejects = 0;
     Backtrack[rayma, constraints, values, lowfacet, essentials, False], 
     rejects++; 0];
     (*
     If[PrintResult,
        If[Length[lowfacet] == 0, 
          Print["Greedy search failed to deliver a new FBF."], 
          Print[ToString[countback] <> " BFBF's found by backtracking."]]
        ]; *)
   (* New entries are added to rejectlist only when the next new FBF is found. 
        If there are no further new FBFs, this can cause an infinite loop. 
        So terminate also when the current search has failed 3 times the greedyfails limit  *)
     If[rejects>3*greedyfails, Break[]];
      (* Update the set of backtrack levels *)
      backlevels = Union[Length /@ BoundedFacets];
      rejections = N@Mean@rejectlist;
      progresscounter = count = Length[BoundedFacets];
      ];
      FBFcount = count;
     If[PrintResult,
     Print["Search terminated with rejection counts for recent BFBF's given by\n"<>ToString[rejectlist]];
     Print[" Backtracked random search has determined the BFBF levels to be " <> ToString[backlevels]]];
    trialcount
    ]

SetAttributes[Backtrack, HoldFirst];
Backtrack[RayMat_, constraints_, values_, facetlist_, essentials_, PrintResult_: False] := 
  Module[{BFancestor, members, candidates, memberlist, maxlevel, oldfbfs, newfbfs},
    oldfbfs = Length[BoundedFacets]; 
    (* Print["Doing a Backtrack "]; *)
    candidates = Reap[Do[
      BFancestor = BoundedQ[RayMat, constraints, DeleteCases[facetlist, i]];
            If[BFancestor, Sow[i]], {i, Complement[facetlist, essentials]}]][[2]];
   (*Print[candidates];*)
    candidates = If[Length[candidates] > 0, candidates[[1]], 
        If[PrintResult, Print["No backtrack candidates found, the facet has no lower level ancestors."]];
        AppendTo[BoundedFacets, facetlist]; Return[1]];
    maxlevel = Length[facetlist];
    members = Complement[facetlist, candidates];
    memberlist = {candidates, members};
    newfbfs = TreeSearch[RayMat, constraints, values,  memberlist, {0, maxlevel}, 
        Infinity, Infinity, PrintResult];
    newfbfs - oldfbfs
    ]

SetAttributes[TracebackSearch, HoldFirst];
TracebackSearch[commons_, RayMat_, constraints_, values_] := 
 Module[{possibles, essentials, nonessentials, removals, 
   removables = {}, candidates, rayma, FBF},
   (*This function tests each non-essential item in the commons list by 
   removing it from the list of possibles,and doing a greedy search for
   an FBF descendant.
   If one is found, all elements absent from the FBF (which will include 
   the probed item)  are removed from the commons list and added to the
   removables list that the function returns.*)
  {possibles, essentials} = MemberList[RayMat, constraints, values];
  nonessentials = Complement[commons, essentials];
  (*Print[nonessentials];*)
  nonessentials = nonessentials[[RandomSample[Range[Length[nonessentials]]]]];
  While[Length@nonessentials > 0,
  	(*Print["Remaining: ",Length@nonessentials];*)
   rayma = RayMat;
   candidates = DeleteCases[possibles, First@nonessentials];
  (*Print["Probing hyperplane ",First@nonessentials];*)
   FBF = GreedySearch[rayma, constraints, values, {}, {candidates, essentials}, False];
   If[Length@FBF > 0,
    (*Print[{"FBF found in Greedy Search",First@nonessentials,FBF}];*)
    removals = Complement[commons, FBF];
    commons = Complement[commons, removals];
    nonessentials = Complement[nonessentials, removals];
    removables = Join[removables, removals],
    (*Print[" Removing common member "<>ToString[First@
    nonessentials]]<>" yielded no descendants "];*)
   If[Length@nonessentials > 0, nonessentials = Rest@nonessentials]];
   ];
  
   removables]

(* DETERMINING THE PROGENITOR FACET *)

SetAttributes[Progenitor, HoldFirst];
Progenitor[RayMat_, RayDim_, constraints_, values_, samplesize_, Printresult_: False] := 
 Module[{startime, treesearch, FBFlevels, common, possibles, essentials, posscount, esscount,
 	discards, removables, totest, removed, misfrac = 1, maxmiss = 0.005, cycle = 0, count},
 (* This function finds an unbounded (but feasible) progenitor facet from which all (known) feasible,
  bounded facets (FBF's) are descended.
  Parameters:  samplesize of initial FBF sample to get starting common for traceback. 
  It returns the facetlist defining the progenitor facet.
  *)
   startime = TimeUsed[];
  progresslabel="Finding a progenitor facet";
  Print[Style[progresslabel, Blue, TextAlignment -> Center]];

    
  (* STEP 1:   COLLECT EITHER A SAMPLE OR ALL BFBF's, AND EXTRACT A SHORTLIST OF COMMON HYPERPLANES  *)
  BoundedFacets = {}; Infeasibles = {}; FBFcount = 0; totalfacets = 0;
  {treesearch,count}=FBFfinder[RayMat, RayDim, constraints, values, samplesize, treesize, Printresult];
  
  (* The following test should never be satisfied. At worst, the combination of all constraint columns
  should eliminate all rays and so give a bounded facet. If not, it means there is a ray that is orthogonal
  to all constraint vectors, but such a ray would define a lineality and those were explicitly eliminated
  by PrismDrop during the SS reduction step. 
  If it happens, either the greedy search or the tree search failed and may need parametr adjustment.  *)
  If[Length@BoundedFacets==0,Print[Style["Catastrophe! FBFfinder failed to deliver any FBF's!",Red]];
  	Abort[]];

  FBFlevels = Length /@ BoundedFacets;
  If[Printresult, Print["The BFBF levels and counts are: "<>TextString@SortBy[Tally[FBFlevels], First]]];
  common = Intersection @@ BoundedFacets;
  If[Printresult, If[treesearch,
  	   Print["Progenitor exhaustive tree search yielded " <> ToString[count] <> 
     " FBF's with " <> ToString[Length[common]] <> 
     " common entries in " <> ToString[TimeUsed[] - startime] <> 
     " seconds."],
   Print["Progenitor random backtracked search with " <> ToString[count] <> 
     " samples yielded " <> ToString[Length[BoundedFacets]] <> 
     " FBF's with " <> ToString[Length[common]] <> 
     " common entries in " <> ToString[TimeUsed[] - startime] <> 
     " seconds."]
     ]];
     
   If[treesearch, Return[{True,common}]];
  
  (* STEP 2: IN CASE OF A RANDOM SAMPLE, DO THE TRACEBACK LOOP TO TRIM THE COMMONS LIST *)
    progressrange={0.8, 1};
    progresslabel= " Traceback sampling to find progenitor ";
    Print[Style[progresslabel, Blue, TextAlignment -> Center]];
    {possibles, essentials} = MemberList[RayMat, constraints, values];
    {posscount, esscount} = {Length@possibles, Length@essentials};
    discards = posscount + esscount - Length@common;
    While[misfrac > maxmiss, cycle++;
    	removables = TracebackSearch[common, RayMat, Kernelcons, Kernelvals];
    	totest = Length[common] - esscount;
    	removed = posscount - discards - totest;
    	If[Printresult && Length@removables > 0, 
    		Print["Found removables " <> ToString[removables] <> " in cycle " <>
    			ToString[cycle] <> " leaving a common of size " <> ToString[esscount + totest]]];
    	misfrac = (8*totest)/(cycle^2 (2*discards + removed) + 2*cycle*removed + 8*totest);
    	progresscounter = 1-misfrac;
    ];
    
   progresslabel=" Traceback sampling completed after "<>ToString[cycle]<>" cycles."; 
   progresscounter=0;
   If[Printresult, 
   	Print["Progenitor search terminated at " <> ToString[cycle] <> 
    " cycles because it is estimated that more than " <> TextString[PercentForm[1 - misfrac, 3]] <> 
    " of all FBF's are descended from it."]];
   
  {False,common}]

(* THE TREE SEARCH *)

SetAttributes[TreeSearch, HoldFirst]; 
TreeSearch[RayMat_, constraints_, values_, Memberlist_, LevelRange_: {1, Infinity},
	 maxcount_: Infinity, maxfacets_: Infinity, PrintResult_: False] := 
  Module[{cons, vars, raycount, startime, rootlist, rootcount, 
      InOutRayList, feasible, bounded, choice, facet, level, minlevel,
    maxlevel, possibles, posscount, possremain, NonElim, viablehi = 0, outcome,
    descriptions =
     {"Viable", "Non-viable", "Rayfree", "Feasible", "Infeasible", "Bounded", "Unbounded", "Subfacet"}, 
      PrintNode = False, tol = 0.00000001},
     
    {cons, vars} = Dimensions[constraints];
    {minlevel, maxlevel} = LevelRange;
    raycount = Length[RayMat];
    startime = TimeUsed[];
    FBFcount = Length[BoundedFacets];
    (* Variables shared between Treesearch and NodeAction *)
    facetcount = 0;
    UnitS = Unitize[RayMat[[All, ;; cons]], tol];
    (*Print[MatrixForm[UnitS]];*)
    
    {possibles, rootlist} = Memberlist;
    posscount = Length[possibles];
    
    (* Check if just the rootlist already gives a rayfree facet *)
    level = Length[rootlist];
    {feasible, bounded} = FeasibleBoundedQ[RayMat, constraints, values, rootlist];
    Which[
    	feasible && bounded, FBFcount++;
      If[PrintResult, Print["Single lowest bounded facet is at root, level "<>ToString[level]]];
      AppendTo[BoundedFacets, rootlist]; Return[FBFcount],
      bounded, 
    If[PrintResult, Print["The lowest bounded facet at root, level "<>ToString[level] <>            
      ", is infeasible, so this facet has no bounded subfacets."]]; Return[0],
      feasible, If[Length[RayMat] > raycount,
        Print[" A new ray was added to the ray matrix at the root level"];
        UnitS = Unitize[RayMat[[All, ;; cons]], tol];]
      ];
    If[posscount == 0, 
      Print["The root facet is feasible but unbounded, and there are no \
    viable subfacets, so the facet has no bounded subfacets."]; 
      Return[FBFcount]];
      
    rootcount = Length[rootlist];
    If[PrintResult, 
      Print["All facets found, share the following " <> 
          ToString[rootcount] <> " hyperplane intersection "<>ToString[rootlist]];
      Print["Additional entries are chosen from the following list of " <>
           ToString[posscount] <> " candidates  " <>ToString[possibles]];
      ];
    
    (* Remove branches that are non-viable, after rays removed by rootlist, 
  before ascending to the next level *)
    
    InOutRayList = Map[1 - Unitize@Total[#[[rootlist]]] &, UnitS];
    Do[possremain = Drop[possibles, i - 1];
      (* Flag the rays that will not be eliminated by remaining possible cols*)
      NonElim = 1 - Sign@Total[UnitS[[All, possremain]], {2}];
      If[NonElim.InOutRayList > 0, Break[], viablehi++],
      {i, posscount}];
    If[PrintResult, 
      Print["The " <> ToString[viablehi] <>" viable subtrees at level 1 are " <> 
          ToString[possibles[[;; viablehi]]]]];
  (* Also remove branches that are not deep enough to reach minimal level *)
   viablehi = Min[viablehi, level + posscount + 1 - minlevel];
  
    Do[
      If[(totalfacets += facetcount) > maxfacets || FBFcount >= maxcount, Break[]];
      facetcount = 0; 
      choice = possibles[[i]]; facet = Join[rootlist, {choice}];
      If[PrintResult,
        PrintTemporary["Starting top level branch " <> ToString[choice] <> 
              " at time " <> ToString[TimeUsed[] - startime] <> 
              " and facet count " <> ToString[totalfacets]];];
      (* Only display full-blown tree searches on progress monitor. 
      Exclude the Backtrack use of Treesearch; recognize that 
      because it starts from minlevel = 0. *)
      If[minlevel>0, progresslabel="Tree search of main branch number "<>ToString[i]];
      outcome = NodeAction[RayMat, constraints, values, facet, Drop[possibles, i],
           rootcount, descriptions, LevelRange, maxcount, maxfacets, PrintNode];
      If[PrintNode && outcome == {4, 6},
        Print[{Drop[facet, rootcount], descriptions[[outcome]],facetcount}]];
      , {i, viablehi, 1, -1}];
    FBFcount
    ]

SetAttributes[NodeAction, HoldFirst]; 
NodeAction[RayMat_, constraints_, values_, Facet_, Possibles_, 
    Esscount_, Descriptions_, LevelRange_, maxcount_, maxfacets_, PrintResult_: False] := 
  Block[{cons, vars, raycount, feasible, bounded, Stop = False, 
      Stop1 = False,  Stop2 = False, choice, facet, RaysInFacet, 
      remains, viablehi = 0, NonElim, level, minlevel, maxlevel, outcome, tol = 0.00000001},
     
    {cons, vars} = Dimensions[constraints];
    {minlevel, maxlevel} = LevelRange;
    raycount = Length[RayMat];
    level = Length[Facet]; 
    nowat = Facet; facetcount++;
    (* Only display full-blown tree searches on progress monitor. 
      Exclude the Backtrack use of Treesearch; recognize that 
      because it starts from minlevel = 0. *)
	If[minlevel>0, progresscounter=totalfacets+facetcount];
    (*Print[{level, facetcount}];*)
    (* Do the subset tests first, then feasibility  *)
    If[SubFacet[Drop[Facet, Esscount], Infeasibles], Return[{5}]];
    If[SubFacet[Facet, BoundedFacets], Return[{8}]];
    (* Below the known minimum FBF, it cannot be bounded. Just test feasibility*)
    {feasible, bounded} = If[level < minlevel, 
    FeasibleBoundedQ[RayMat, constraints, values, Facet, False],
       FeasibleBoundedQ[RayMat, constraints, values, Facet, True]];
    If[level < minlevel, bounded = False];
    Which[
      feasible && bounded, FBFcount++;
      	AppendTo[BoundedFacets, Facet]; Return[{4, 6}],
      bounded,(*Print["Infeasible but bounded found at facet no "<>ToString[facetcount]];*)
      	AppendTo[Infeasibles, Drop[Facet, Esscount]]; 
      	Return[{5, 6}],
      feasible, 
       If[Length[RayMat] > raycount, raycount = Length[RayMat];
        If[PrintResult, Print[" A new ray was added to the ray matrix at node " <>
            	 ToString[Drop[Facet, Esscount]]]];
        UnitS = Unitize[RayMat[[All, ;; cons]], tol]],
      True,(*Print["Unbounded and Infeasible at facet no "<>ToString[facetcount]];*)
      	AppendTo[Infeasibles, Drop[Facet, Esscount]]; 
      	Return[{5, 7}]
      ];
    
    If[level >= maxlevel,
    	(*Print["NodeAction bailing out at level "<> ToString[level]];*)
      Return[{4, 7}]];
    
    If[FBFcount >= maxcount, Return[{4, 7}]];
    
    (* Viability pruning usually prevents Possibles from being empty. 
    However, when the LP bounded test failed, it can happen. 
    There is no harm, neither of the following loops are executed in that case, 
    so as appropriate it just returns with no action taken. *)
    RaysInFacet = Map[1 - Unitize@Total[#[[Facet]]] &, UnitS];
    Do[remains = Drop[Possibles, i - 1];
      (* Flag the rays that will not be eliminated by remaining possible cols*)
      NonElim = 1 - Sign@Total[UnitS[[All, remains]], {2}];
      (*Print[{RaysInFacet, NonElim}]*)
      If[NonElim.RaysInFacet > 0, Break[], viablehi++],
      {i, Length[Possibles]}];
  (* Also remove branches that are not deep enough to reach minimal level *) 
  viablehi = Min[viablehi, level + Length[Possibles] + 1 - minlevel];
   (* Print["Only the first "<>ToString[viablehi]<>
    " of the available branches are viable and reach the minimal level."]; *)

    (* Process the viable nodes at the next level of the search tree *)
    (*
    Print[{Facet[[Esscount+1;;]],Possibles[[;;viablehi]]}];
    *)
    Do[If[totalfacets+facetcount > maxfacets || FBFcount >= maxcount, Break[]];
      choice = Possibles[[i]]; facet = Join[Facet, {choice}]; 
      outcome = NodeAction[RayMat, constraints, values, facet, Drop[Possibles, i],
           Esscount, Descriptions, LevelRange, maxcount, maxfacets, PrintResult];
      If[outcome == {4, 6}(*||Length[BoundedFacets]\[Equal]6*),
        If[PrintResult, 
          Print[{Drop[facet, Esscount], Descriptions[[outcome]],facetcount}]]
        ];
      , {i, viablehi, 1, -1}];
     
    {4, 7}
    ]

BranchAllocate[BFBF_, nodelist_: {}] := 
 Module[{treelist, tofind, found, minlevel, participants, branchfbfs},
  treelist = If[nodelist == {}, SortBy[Tally[Join @@ BFBF], Last][[All, 1]], nodelist];
  minlevel = Min[Length /@ BFBF];
  tofind = Range[Length[BFBF]];
  branchfbfs = Reap[Do[
      participants = treelist[[-branch ;;]];
      found = Flatten@Position[
         BFBF[[tofind]], _?(Quiet@Complement[#, participants] == {} &)];
      found = tofind[[found]];
      Sow[found];
      tofind = Complement[tofind, found];
      , {branch, minlevel, Length[treelist]}]][[2]];
  branchfbfs = Join[ConstantArray[{}, minlevel - 1], branchfbfs[[1]]];
  Reverse@branchfbfs
  ]


End[] (* End Private Context *)

EndPackage[]
```
