## Supplementary material for "The FBA Solution Space Kernel – Introduction and Illustrative Examples": SSK software manual and code: FixedFluxes.m

```
(* Wolfram Language Package *)

(* COPYRIGHT
					© Copyright 2022 Wynand Verwoerd

This file is part of SSKernel.

The SSKernel program is free software; you can redistribute it and/or modify it under the terms of the GNU General Public License as published by the Free Software Foundation; either version 3 of the License, or (at your option) any later version.
 
This program is distributed in the hope that it will be useful, but WITHOUT ANY WARRANTY; without even the implied warranty of MERCHANTABILITY or FITNESS FOR A PARTICULAR PURPOSE . See the GNU General Public License for more details.
 
You should have received a copy of the GNU General Public License along with this program . If not, see http://www.gnu.org/licenses/
*)

BeginPackage["FixedFluxes`",{"Configuration`"}]
(* Exported symbols added here with SymbolName::usage *)  

HopSkipnJump::" The Hop Skip and Jump algorithm for removing fixed fluxes."
SSKfixvals::usage=" Mop up any fixed values missed by HSnJ by inspecting SSK periphery points"

Begin["`Private`"] (* Begin Private Context *) 

HopSkipnJump[tol_: 10.^-3,PrintResult_:False] :=
 (* Global Variables: The equals constraint augmented matrix and nonnegative variable upper limits;
 and a known solution vector i.e. vertex for this constraint problem. 
  Return value: A list of frozen variables found by extreme point sampling. 
  *)
 Module[{realuppers, tempmax,orthos = {}, nonorthos, neworthos = {0}, 
   SpanningRows, norms, keepers, nonzerolows, feasible, constraints, 
   fixlist, fixvals, floats, differ, rows, cols, jumpto},
   (* The algorithm operates on feasible, which is a VERTEX found by Simplex FBA. 
   This point is progressively reduced in dimension (by Skipper) by removing 
   fixed value variables, while leaving the global array feasiblepoints 
   untouched, until the end when all feasible points are transferred to the 
   variable flux subspace together.  *)
  feasible = feasiblepoints[[1]];
  realuppers=uppers;
  {rows, cols} = Dimensions[augment];

  (* THE FIRST SKIP STEP  - 
  FIND AND ELIMINATE ORTHOGONAL VARIABLES *)
  
  If[PrintResult,Print["Starting Hop, Skip and Jump."]];
  
  fixlist = Flatten@Reap[
      Do[If[Abs[uppers[[col]] - lowers[[col]]] < tol, Sow[col]],
      	 {col, cols - 1}]][[2]];
  constraints = Join[augment[[All, ;; -2]], Map[UnitVector[cols - 1, #] &, fixlist]];
  nonorthos = Range[cols - 1];
  progresslabel="Checking for orthogonal flux axes that can be skipped.";
  Print[Style[progresslabel, Blue, TextAlignment -> Center]];
  progressrange={0,cols};
  progresscounter=0;
  While[neworthos != {},
  	(* Note the small tolerance 0.0000001 in the call to NullSpace. 
  	Bigger tolerances seem to give inaccurate results and are not recommended. *)
   SpanningRows = N@NullSpace[constraints,Tolerance-> 0.0000001];
   (* If a column of the spanning row matrix is a zero vector, it means that
   the corresponding axis of the row space is orthogonal to it and is a fixed variable *)
   norms = Norm /@ Transpose[SpanningRows];
   keepers = Map[If[# > tol, True, False] &, norms];
   neworthos = Flatten@Position[keepers, False];
   progresscounter+=Length[neworthos];
   If[neworthos != {},
    Skipper[feasible, neworthos]; 
    {rows, cols} = Dimensions[augment];
    orthos = Join[orthos, nonorthos[[neworthos]]];
    nonorthos = Delete[nonorthos, Partition[neworthos, 1]];
    constraints = augment[[All, ;; -2]]];
   ];
   If[PrintResult, Print["A total of "<>ToString[Length[orthos]]<>
   	" orthogonal flux axes Skipped."]];
  progresslabel="Skip step completed. ";
  progresscounter=0;

   (* A shortlist of candidate fixed variables is found below, by jumping 
   to a maximally different solution compared to the one previously found.
   That hits a snag in the  case of a solution space that is partly unbounded, 
   since then the max distance is infinite.
 
 	Deal with that by replacing all infinite variable bounds by a temporary maximum. It 
 	does not matter too much what this maximum is, since its only function is to encourage
 	component values to become different from the existing ones, so simply choose it 
 	as twice the maximum known flux component (this preserves a reasonable scale for SS).
 	
 	To also provide for the case where the supplied solution is close to zero flux,
 	as e.g.happens when there is no FBA objective and RSS is a simple cone. 
 	In that case, simply take the default capping radius.
 	*)
    tempmax=Max[ 2.Max@Abs@feasiblepoints[[1]], DefaultCap];
    uppers = uppers /. {_?(# > tempmax &) -> tempmax};
 
 
  (* Variables with lower limits larger than zero, 
  are temporarily shifted as required by Hopper and Jumper.
  Modify the last column of the augmented matrix, 
  the feasible solution and the limits accordingly. *)
  nonzerolows = Flatten@Position[lowers, low_ /; low > 0];
 (* Print[{"Non-zero lower limits for ", ToString@nonzerolows}]; *)
  augment[[All, -1]] -= augment[[All, nonzerolows]].lowers[[nonzerolows]];
  uppers[[nonzerolows]] -= lowers[[nonzerolows]];
  feasible[[nonzerolows]] -= lowers[[nonzerolows]];
  (* Do row reduction on augment and put it in canonical form as required by Hopper  *)
  ExtendedBasis[feasible];
  {rows, cols} = Dimensions[augment];
(*
   Print["After row reduction"];
   Print[Max@Abs@Chop[augment[[All,-1]]-augment[[All,;;-2]].feasible,  tol]]; 
 *) 
  (* JUMP TO A NEW FEASIBLE POINT AND GET CANDIDATES FOR FIXED VARIABLES *)
  
  jumpto = Jumper[feasible, tol];
  differ = Chop[feasible - jumpto, 5*tol];
  fixvals = 
   Complement[Range[cols - 1], Flatten@Position[differ, _?(# != 0 &)]];
   If[PrintResult,Print[ToString[Length[fixvals]]<>
  " Candidate fixes after long Jumping . "]];
  (* jumpto[[nonzerolows]] += lowers[[nonzerolows]]; *)
  
  (* USE HOPPING TO FILTER CHANGEABLE VARIABLES FROM THE LIST, 
  THEN RESTORE LOWER LIMITS *)
  
  progresslabel="Trimming down shortlist of fixed values by hopping.";
  Print[Style[progresslabel, Blue, TextAlignment -> Center]];
  progressrange={0,Length[fixvals]};
  
  fixvals = Hopper[feasible, tol, fixvals];
   If[PrintResult,Print[ToString[Length[fixvals]]<>
  " Confirmed fixes after doing the Hops. "]];
  uppers[[nonzerolows]] += lowers[[nonzerolows]];
  augment[[All, -1]] += 
   augment[[All, nonzerolows]].lowers[[nonzerolows]];
  (* Map[Print[Max@Abs@Chop[augment[[All,-1]]-augment[[All,;;-2]].#,
  tol]]&,feasiblepoints]; *)
  
  (* SECOND SKIP -  REMOVING THE NEWLY FOUND FIXED VARIABLES *)
  Skipper[feasible, fixvals];
  (* Return a consolidated list of all fixed variables and their values, after
  transferring the global feasible points array to variable flux space *)
   fixvals = Union[orthos, nonorthos[[fixvals]]];
   uppers=realuppers[[Complement[Range[Length@realuppers], fixvals]]];
   floats = Complement[Range[Length[First@feasiblepoints]], fixvals];
   fixvals = Chop[Transpose[{fixvals, feasiblepoints[[1,fixvals]]}],tol];
   feasiblepoints = feasiblepoints[[All, floats]];
 (* 
  {rows, cols} = Dimensions[augment];
  Print["A total of " <> ToString[Length[fixvals]] <> 
    " fixed variables were eliminated."];
  Print["The problem now has " <> ToString[rows] <> 
    " equals constraints and " <> ToString[--cols] <> 
    " variables. "];
*)    
  progresslabel="Hop, Skip and Jump search for fixed values completed.";
  progresscounter = 0;

  fixvals
  ]

Hopper[feasible_, tol_, shortlist_: All] := 
 Module[{rows, cols, nonzerolows, cons, vals, values, limits, 
   discard = {0}, groups, keepers, bots, mids, tops, obvector, hopto},
  (* This function uses a cycle of hops to filter all remaining \
changeable variables from the shortlist.*)
  (* Equality constraints are taken from the global matrix augment. *)

  (* Note that for this routine to function correctly, 
  it is essential that augment should be in canonical
     form corresponding to the hopfrom point (i.e. 
  feasible) as produced by function ExtendedBasis  *)
  {rows, cols} = Dimensions[augment];
  nonzerolows = Flatten@Position[lowers, low_ /; low > 0];
  {cons, vals} = Normal@{augment[[All, ;; -2]], augment[[All, -1]]};
  values = Transpose[{vals, ConstantArray[0, rows]}];(* {value, 
  type} pairs; type=0 => equal *)
  limits = Transpose[{ConstantArray[0., cols - 1], uppers}];
  keepers = shortlist;
  While[discard != {},
   groups = 
    Reap[Do[j = keepers[[i]]; 
       Which[feasible[[j]] < tol, Sow[j, lows], 
        uppers[[j]] - feasible[[j]] < tol, Sow[j, ups], True, 
        Sow[j, ibtw]], {i, Length[keepers]}], {lows, ups, ibtw}][[2]];
   {bots, tops, mids} = Flatten[groups /. {} -> {{}}, 1];
(* Print[{bots,tops,mids}]; *)
   obvector = ConstantArray[0., cols - 1]; 
   obvector[[bots]] = -1.;   obvector[[tops]] = 1.;
   hopto = 
    LinearProgramming[obvector, cons, values, limits, 
     Method -> "InteriorPoint", Tolerance -> 0.1 tol]; 
   If[!VectorQ[hopto],
    	Print[Style["Aborting - no valid LP solution found by Hopper.",Red]];Abort[]];

   discard = 
    Position[(feasible - hopto)[[keepers]], _?(Abs[#] > tol &)];
    progresscounter+=Length[discard];
   (*Print[discard];
   Map[Print[{#,feasible[[#]],solution[[#]],Abs[feasible[[#]]-
   solution[[#]]]}]&,keepers[[Flatten@discard]]];*)
 
   keepers = Delete[keepers, discard]];
  keepers]

SetAttributes[Skipper, HoldFirst];
Skipper[feasiblepoint_, fixes_] := 
 Module[{rows, cols, fixcount, floats, shift, constraints, values, 
   nonzerorows},
  (* This function skips over all the axes listed in the 'fixes' argument.
  It operates on the augmented constraints matrix supplied as a global variable. 
  It updates the global variables (i.e., the augmented matrix, 
  the solution set and lower/upper limits)to a reduced space 
  formed by removing the 'fixes' *) 
  constraints = augment[[All, ;; -2]]; values = augment[[All, -1]];
  {rows, cols} = Dimensions[constraints];
  fixcount = Length[fixes];
  If[fixcount > 0,
   floats = Complement[Range[cols], fixes];
   (* Substitute fixed values, 
   then remove those columns by shifting to the RHS *)
   shift = Chop[feasiblepoint]; 
   shift[[floats]] = ConstantArray[0., cols - fixcount];
   values -= constraints.shift;
   constraints = constraints[[All, floats]];
   lowers = lowers[[floats]]; uppers = uppers[[floats]];
   feasiblepoint = feasiblepoint[[floats]];
   (* Remove constraints that are trivial, 
   due to removal of columns or a silly original model *)
   nonzerorows = 
    Flatten@Position[Normal@constraints, _?(Norm[#] > 0 &), {1}];
   If[nonzerorows != {},
    constraints = constraints[[nonzerorows]]; 
    values = values[[nonzerorows]];
    augment = 
     SparseArray@Join[constraints, Transpose[{values}], 2]];
   ];
  ]

SetAttributes[VertexHopper, HoldFirst];
VertexHopper[feasible_] := Module[{rows, cols, values, change, lims},
  (* Feasible is a solution for the equality constraint set specified 
by global matrix augment *)
  {rows, cols} = Dimensions[augment];
  (* Minimise the closest distance to each limit *)
  change = 
   MapThread[
    If[#1 > #2, -1., 1.] &, {uppers - feasible, 
     feasible - lowers}];(*Print[Dimensions@change];*)
  (* Or just give it a trivial objective, sometimes converges faster *)
  (*change=ConstantArray[0.,cols-1];*)
  values = 
   Transpose[{augment[[All, -1]], ConstantArray[0, rows]}];
   (* {value, type} pairs; type=0 => equal *)
  (*Print[Dimensions@values];*)
  lims = Transpose[{lowers, uppers}];(*Print[Dimensions@lims];*)
  (* Don't use the Tolerance option with Simplex - 
  strangely that often gives infeasible *)
  feasible = 
   LinearProgramming[change, augment[[All, ;; -2]], values, lims, 
    Reals, Method -> "Simplex"]
  ]

Jumper[feasible_, tol_, vertex_: False] := 
 (* This function finds a feasible point that is markedly different from
 the feasible point given as argument, so that as many as possible flux components
 can be ruled out from being fixed since they differ between this pair of points.
 It achieves this by a variation on minimising the Manhattan distance; it maximizes, but 
 inserts an explicit upper limit for the auxiliary variables zi. While the maximized z 
 is always at this assumed upper limit, the flux point which delivers this tends to be
 well displaced from the given feasible point. . 
 *)
 
 Module[{rows, cols, Imat, constraints, values, 
   bounds, objectvector, xz, solution, jumptol},
   (* jumptol is the LP tolerance that determines how accurately the far side endpoint
   of the jump is determined. This needs to be stricter than the tolerance tol within which
   changes are ignored, otherwise actual fixed fluxes may be wrongly rejected.
   *)
    jumptol = LPtol ; 
    {rows, cols} = Dimensions[augment]; cols--;
   (* 
    Print[Abs@Chop[augment[[All,-1]]-augment[[All,;;-2]].feasible, tol]];*)
    Imat = SparseArray[{{i_, i_} -> 1}, {cols, cols}];
    constraints = 
   SparseArray@
    ArrayFlatten[{{augment[[All, ;; -2]], 0}, {-Imat, Imat}, {Imat, 
       Imat}}];
    (*Print[Dimensions[constraints]];*)
    (* Print@MatrixForm@Normal[constraints];*)
    (* characterise constraint types by {value, type} pairs; 
    type=0 => equal, 1 => greater/eq *)
    values = 
   Transpose@
    SparseArray[{Flatten[{augment[[All, -1]], -feasible, feasible}], 
      Flatten[{ConstantArray[0., rows], ConstantArray[1, 2*cols]}] }];
    (*Print@Normal[values];*)
    objectvector = 
   SparseArray@
    Flatten[{ConstantArray[0., cols], -ConstantArray[1., cols]}];
    (*Print@Normal[objectvector];*)
 
     (* The next line is one where the flux values are assumed to be
     positive (0 lower limit). That would need to be changed if HSnJ 
     is applied to reversible reactions. There may be other places that 
     also need changing in that case. *)
     bounds = Transpose[{ConstantArray[0., 2*cols], 
    	 Flatten[{uppers, uppers}]}];
     (* Notice how the upper constraint is applied to the auxiliary vector z as well,
     this avoids an unbounded LP while still producing a jump!*)
    (*Print@Normal[bounds];*)
   
    (*Note that here, for Jumper, it matters whether Simplex or InteriorPoint is used. 
    Both produce the same (trivial) maximum for the sum of z components, but
    jump to different points to achieve that.
    Interiorpoint is both faster and generally gives a better jump. *)
    xz = LinearProgramming[objectvector, constraints, values, bounds, 
    Method -> "InteriorPoint", Tolerance -> jumptol]; 
    
    (*Print[Chop[(values[[All,1]]-constraints.xz)[[;;rows]],tol]];*)
    If[VectorQ[xz], solution = Take[xz, cols],Print[xz];
    	Print[Style["Aborting - no valid LP solution found by Jumper.",Red]];Abort[]];
    solution = If[vertex, VertexHopper[solution], solution];
    Chop[solution, jumptol]]


SetAttributes[ExtendedBasis, HoldFirst];
ExtendedBasis[feasible_] :=
 (* From the feasible solution that satisfies nonnegativity, the global augmented matrix and upper limits
 this function calculates a Simplex extended basis and corresponding row reduced augmented form 
 The feasible solution must be a vertex of the solution space, not interior point!? 
 Global variables: augment, base, uppers
 *)
 Module[{reorder, groups, targetpairs, restore, discard, rows, cols, 
   tol = 10.^-6},
  {rows, cols} = Dimensions[augment];
   (* To choose basis columns, 
   first priority are nonzero solution values below their limits; 
   then the ones at upper limits, and last the zero values. 
   Express this as a reordering list *)
   groups = 
    Flatten /@ 
     Reap[Do[Which[feasible[[i]] < tol, Sow[i, nuls], 
         uppers[[i]] - feasible[[i]] < tol, Sow[i, ups], True, 
         Sow[i, ibtw]], {i, cols - 1}], {ibtw, ups, nuls}][[2]];
         
   If[Length[groups[[1]]] > rows, 
    MessageDialog[
     " The feasible point has too many non-boundary variables for a \
valid Simplex basis - it is replaced by a similar vertex, which can take some time."]; 
	VertexHopper[feasible]; 
    groups = Flatten /@ 
      Reap[Do[Which[feasible[[i]] < tol, Sow[i, nuls], 
          uppers[[i]] - feasible[[i]] < tol, Sow[i, ups], True, 
          Sow[i, ibtw]], {i, cols - 1}], {ibtw, ups, nuls}][[2]]];
          
   (* Reorder columns to put preferred basis columns first, then do row reduction *)
   (*Print[groups];*)
   reorder = Append[Flatten[groups], cols];
   targetpairs = Transpose[{Range[cols], reorder}];
   restore = 
    Sort[targetpairs[[reorder]], #1[[2]] < #2[[2]] &][[All, 1]];
   augment = RowReduce[augment[[All, reorder]], Tolerance -> tol];
   (* Construct the full base from pivot columns, 
   then restore the ordering and discard empty rows*)
   (*
   base = Flatten@Map[Position[#, 1 | 1., 1, 1] &, augment[[All, ;; -2]]];
   base = reorder[[base]];
   *)
   augment = augment[[All, restore]];
   discard = Flatten@Position[augment, ConstantArray[0, cols], 1];
   (*Print[ToString[Length[discard]]<>
   " constraint dependencies were detected and eliminated."];*)
   augment = Chop[augment[[Complement[Range[rows], discard]]], tol];
   augment = SparseArray[augment];(* Saves about 25% computing time in one trial *);
  ]

SSKfixvals[SSKpoint_,PeriPoints_, rays_ , fixtol_] := 
 Module[{valuecounts, fixes, rayzeroes},
  (* This function finds fixed flux components by inspection of SSK periphery points.
  Except when the periphery points coincide with the SSK vertices (e.g. when the SSK 
  is a single point,   such as for a cone) it is a heuristic assumption 
  that the periphery points adequately cover all degrees of freedom of the SSK. *)
  valuecounts = 
   Map[Length@Tally[#, Abs[#1 - #2] < fixtol &] &, 
    Transpose@PeriPoints];
  fixes = Flatten@Position[valuecounts, 1];
  rayzeroes = If[rays=={}, fixes, Flatten@Position[Total@Abs@rays, 0]];
  fixes = Intersection[fixes, rayzeroes];
  Transpose[{fixes, SSKpoint[[fixes]]}]
  ]


End[] (* End Private Context *)

EndPackage[]
```
