## Supplementary material for "The FBA Solution Space Kernel – Introduction and Illustrative Examples": SSK software manual and code: ModelProcessing.m

```
(* ::Package:: *)

(* Wolfram Language Package *)

(* COPYRIGHT
					\[Copyright] Copyright 2022 Wynand Verwoerd

This file is part of SSKernel.

The SSKernel program is free software; you can redistribute it and/or modify it under the terms of the GNU General Public License as published by the Free Software Foundation; either version 3 of the License, or (at your option) any later version.
 
This program is distributed in the hope that it will be useful, but WITHOUT ANY WARRANTY; without even the implied warranty of MERCHANTABILITY or FITNESS FOR A PARTICULAR PURPOSE . See the GNU General Public License for more details.
 
You should have received a copy of the GNU General Public License along with this program . If not, see http://www.gnu.org/licenses/
*)

BeginPackage["ModelProcessing`", { "Configuration`","DataHandler`","Centering`","FixedFluxes`",
	"RayFinding`","RedundancyRemoval`","FBFSearch`","Capping`","FacetTesting`"}]
(* Exported symbols added here with SymbolName::usage *)  

LoadModel::usage = "Loads data from a datafile and performs integrity checks. "
DoFBA::usage = "Performs reference FBA on input data with artificial bounds."
ReducedSolutionSpace::usage="Function for preliminary reduction of FBA solution space dimensions "
KernelFinder::usage="Finds the progenitor facet and performs coincidence and tangent capping "
KernelShape::usage="Finds chords, centre and peripheral points for a closed kernel polytope. "
KernelDisplay::usage="Performs validation test, displays chord plots and transfers all results to the full flux space."
SaveResults::usage=" Exports a verbal report and the calculated SSK and other results to disk files."
StageManager::usage=" Manages the main stages of the calculation in response to button press by user "
UserInterface::usage=" Creates the control- and reportwindow contents. "
(*
Reform::usage=" Reformulate model with fluxes positive and without artificial limits "
Revert::usage=" Reconsolidate opposing reaction pairs and revert to space of variable fluxes, reversible reactions last"
*)

Begin["`Private`"] (* Begin Private Context *) 

LoadModel[DataFile_, printresult_]:=
Module[{cons,vars, bigmodel = 1500 },
progresslabel="Loading data from input file.";
Print[Style[progresslabel, Blue, TextAlignment -> Center]];
(* Print["File "<>DataFile<>" with extension "<>FileExtension[DataFile]]; *)
Switch[ToLowerCase@FileExtension[DataFile],
  "m"|"wl", MReader[DataFile, S, Svals, bounds, objectselector, 
   FBAvector, ReactionNames, MetaboliteNames, maxmin, printresult],
  "mat", MatReader[DataFile, S, Svals, bounds, objectselector, 
    FBAvector, ReactionNames, MetaboliteNames, maxmin, printresult],
  "xml"|"sbml", SBMLReader[DataFile, S, Svals, bounds, objectselector, 
    FBAvector, ReactionNames, MetaboliteNames, maxmin, printresult],
  _, Print["Invalid data file of type " <> FileExtension[DataFile]]];

(* Having read a .mat or  SBML file, create a .m file for testing purposes *)
(*
exportfile = StringReplace[datafile, ".mat" | ".sbml" | ".xml" -> ".m"]
Export[exportfile, {ModelName, S, bounds, objectselector, FBAvector, ReactionNames, 
	MetaboliteNames, Switch[maxmin, -1, "max", 1, "min", _, "max"]}, "List"];
*)

exemptions = CheckBalance[S, bounds, externals, True];

 If[printresult, Print[Show[MatrixPlot[{S . objectselector},
		PlotLabel->"Metabolites contributing to the objective"],ImageSize->Full]]];
 artificial = ChooseArt[bounds, printresult];
 progresscounter=0.5;		

(* LP tolerance can be relaxed in case of convergence problems. *)
 {cons, vars} = Dimensions[S];
 LPmethod = If[StringQ[LPmethod], LPmethod, 
	If[vars < bigmodel, "Simplex",  "InteriorPoint"]];
 If[vars >= bigmodel, timeconstraint=200.];
 If[loadreset, 
	Tolerances = ToString@If[vars < bigmodel, 0.0001,  0.001];
	fixtol = If[vars < bigmodel, 0.002,  0.05];
	maxthin = Max[maxthin,fixtol];
];

(* Discard imported solution if invalid or in conflict with explicit solution *)
  If[Chop@Norm[FBAvector] == 0. || Chop@Norm[S . FBAvector] != 0. ||
   Position[MapThread[#2[[1]] <= #1 <= #2[[2]] &, {Chop[FBAvector], bounds}], False] != {}, 
  Print["The imported FBA vector is absent, zero or violates its constraints, so is ignored!"];
  feasiblepoints = {},
  feasiblepoints = {FBAvector}];
(*Print[Dimensions[feasiblepoints]];*)
If[objectcount != 1, 
  Print["The current implementation only provides for a single objective vector."]];
If[Norm[objectselector] == 0., 
 Print[Style["This model has no valid objective and so lacks an objective space.
 Reduction of the stoichiometrically feasible flux space is done instead.", Darker[Green]]]];
(* Calculate a solution with artificial boundaries and check its objective value 
	Use InteriorPoint method here, even though Simplex is used for subsequent calculations, 
	because artificial upper limits can give convergence failure for large models. *)

  progresscounter=1.;
  progresslabel="Data input and testing completed.";
]

ChooseArt[bounds_, printresult_ : False] := 
 Module[{maxpower,lobins,hibins, counts,gap,power,loart,labels,lowchart,
 	hiart,hichart,choice, artificial,scaletop, lineat, showlo, showhi}, 
  (*Function to display bounds histogram and choose artificial limit value.
  It proposes a value based on inspecting histogram for gaps.*) 
  artificial = 100.; (* lower limit for cases without a gap*)
  (* Setting artificial to a reasonable value independent of the global setting
  	avoids the situation that a low value inherited from a previous run causes
  	proper upper bounds to be eliminated. This is only done in stage 1, the user
  	remains free to choose a value before stage 2 is run. *)	
  lowers = bounds[[All, 1]]; 
  uppers = bounds[[All, 2]];
  maxpower = Ceiling@Log10@Max[1, Cases[Abs@lowers, Except[Infinity]]];
  maxpower = Min[maxpower, 6];
  (*Set up logarithmic bins,negative intervals for each power of 10,
  all positve values lumped together*)
  lobins = Reverse@Table[-10.^power, {power, 0, maxpower}];
  lobins = Append[Riffle[lobins, 5*Rest[lobins]], 100];
  counts = BinCounts[lowers, {lobins}];
  (*Now look for the last of up to 2 gaps,
  counting back from 0. This is taken as the value where artificial bounds start*)
  gap = SequencePosition[Reverse@counts, {0, _?Positive}, 2];
(* Print[{"Lower bounds: counts and gaps ", lobins, counts, gap}]; *)
  loart = Switch[Length@gap, 0, artificial, _, Abs@lobins[[-gap[[-1, 2]] - 1]]];
(* Print[{"The lower bound gap, and lower artificial bound",gap,loart}]; *)
  labels = Map[ScientificForm[#, ScientificNotationThreshold -> {0, 4}] &, lobins];
  scaletop = Log@Max[counts];
  lowchart = BarChart[counts, PlotRange -> {{Automatic, Automatic}, {0.1, Automatic}},
  	ScalingFunctions -> "Log",  PlotLabel -> "Histogram of Lower Limits", 
    ChartLabels -> Placed[labels, Axis, Rotate[#, Pi/4] &],
    ImagePadding -> {{30, 0}, {40, 0}}, PlotRange -> {{Automatic, Automatic}, {0.1, Automatic}}];
  maxpower = Ceiling@Log10@Max[100.,Cases[uppers, Except[Infinity]]];
  maxpower = Min[maxpower, 6];
  (*Set up logarithmic bins,intervals for each power of 10,
  all negative values lumped together*)
  hibins = Table[10.^power, {power, 0, maxpower}];
  hibins = Prepend[Riffle[hibins, 5*hibins], -1000.];
  counts = BinCounts[uppers, {hibins}];
  (*Now look for the last of up to 2 gaps.This skips any gap if there \
are no negative values.This is taken as the value where artificial bounds start*)
  gap = SequencePosition[counts, {0, _?Positive}, 2];
(* Print[{"Upper bounds: counts and gaps ", hibins, counts, gap}]; *)
  hiart = Switch[Length@gap, 0, artificial, _, Abs@hibins[[gap[[-1, 1]] + 1]]];
  choice = Max[artificial, Min[loart, hiart]]; (* Protect small models with  no gap *)
  lineat = First@FirstPosition[lobins, n_ /; n >= -choice] + 0.5;
  showlo = lobins[[1]] <= -choice <= lobins[[-1]];
  lowchart=Show[lowchart, If[showlo, 
  	Graphics[{Blue, Thickness[0.02], Line[{{lineat, -2.5}, {lineat, scaletop}}]}], {}]
  	 ];  

  labels = Map[ScientificForm[#, ScientificNotationThreshold -> {0, 4}] &,Rest@hibins];
  scaletop = If[Total@counts == 0, 6, Log@Max[counts]];
  lineat = First@FirstPosition[hibins, n_ /; n >= choice] - 0.5;
  showhi = hibins[[2]] <= choice <= hibins[[-1]];
  hichart = BarChart[counts, PlotRange -> {{Automatic, Automatic}, {0.1, Automatic}},
  	ScalingFunctions -> "Log",  PlotLabel -> "Histogram of Upper Limits", 
    ChartLabels -> Placed[labels, Axis, Rotate[#, Pi/4] &],
    ImagePadding -> {{30, 0}, {40, 0}}, PlotRange -> {{Automatic, Automatic}, {0.1, Automatic}}];
  hichart=Show[hichart, If[showhi, 
  	Graphics[{Blue, Thickness[0.02], Line[{{lineat, -2.5}, {lineat, scaletop}}]}], {}]
  	];  
  If[printresult, 
   Print[GraphicsRow[{lowchart, hichart}, ImageSize -> Full, 
     AspectRatio -> 0.35, Frame -> All, Spacings -> 0]];
   Print["Proposed lowest absolute boundary value taken as artificial is " <> TextString[choice]]];
  choice]

SetAttributes[DoFBA, HoldFirst];  
DoFBA[tolerances_, printresult_ : False] := 
	(* Function to do the FBA calculation with artificial flux bounds. *)
Module[{tolerance, (* FBAtol,*) FBAflux, objective, optimum, Svals},
	
  	If[exempt,
  		exemptcount=Length@exemptions;
  		Print[ToString[exemptcount]<>" unbuffered externals are exempted from flux balance."];
  		MetaboliteNames=Delete[MetaboliteNames,Partition[exemptions,1]];
  		S=Delete[S,Partition[exemptions,1]],
  		exemptcount=0;	
  	];

  (* Take homogenous constraints S.f = 0 irrespective of imported values *)  
  Svals = ConstantArray[{0, 0}, Length[S]]; (*{value,type} pairs; type=0 =>equal*); 
  tolerance=ToExpression@StringSplit[tolerances, ","];
  {FBAtol,LPtol}=If[Length@tolerance==1,{tolerance[[1]],tolerance[[1]]},{tolerance[[1]],tolerance[[2]]}];
  If[!FBAdone,
  progresslabel="LP calculation for imported model with artificial boundaries. ";
  Print[Style[progresslabel, Blue, TextAlignment -> Center]];
  progresscounter:=Clock[5]; progressrange=Indeterminate;
(*  Print["maxmin,objectselector, S, Svals, bounds",maxmin," ",Length@objectselector, 
  	Dimensions@S, Dimensions@Svals, Dimensions@bounds]; *)
(*
  {LPtime, result} = Timing[ FBAflux = Quiet[Check[
     LinearProgramming[maxmin*objectselector, S, Svals, bounds, Reals,
       Tolerance -> FBAtol, Method -> LPmethod], 
     "No LP solution was found! ", {LinearProgramming::lpsnf, 
      LinearProgramming::lpsub, 
      LinearProgramming::lpsnfp}], {LinearProgramming::lpsnf, 
     LinearProgramming::lpsub, LinearProgramming::lpsnfp}]];
*)     
     {LPtime, result} = Timing[ FBAflux = Check[
     LinearProgramming[maxmin*objectselector, S, Svals, bounds, Reals,
       Tolerance -> FBAtol, Method -> LPmethod], 
     "No LP solution was found! ",
      {LinearProgramming::lpsnf, LinearProgramming::lpsub, LinearProgramming::lpsnfp}]
      	];
   If[printresult,Print["Bounded LP calculation took "<>ToString[LPtime]<>" seconds"]]; 
	,
  FBAflux = boundedflux];
     
  If[VectorQ[FBAflux], objective = objectselector . FBAflux;
   If[printresult, 
    Print["The objective value calculated here for imported model is " <> TextString[objective]]];
   If[Abs[objective] < lptol, 
    Print[" The optimized objective value is zero and since this is \
consistent with zero flux, the full solution space is a cone and the SSK is just the origin. "]];

   If[Length[feasiblepoints] == 1,
    optimum = objectselector . feasiblepoints[[1]];
    If[Abs[optimum - objective] < 3*lptol, 
     Print["Imported and calculated objectives agree."];
     PrependTo[feasiblepoints, FBAflux], 
     Print["Discrepancy between imported and calculated objective values " <>
     	 ToString[{optimum, objective}] <> 
       " .\nImported FBA solution discarded for now,       	but this \
may indicate invalid input data and should be checked."];
     feasiblepoints = {FBAflux}],
    feasiblepoints = {FBAflux}];
   boundedflux = FBAflux (* store the result in case of repeated reduction stage *)
   ,
   (* LP calculation unsuccessful; put a temporary zero vector for the flux, 
   it will give zero objective and so be eliminated again if the unconstrained LP is successful. *)
   feasiblepoints = PrependTo[feasiblepoints,ConstantArray[0., Dimensions[S][[2]]]];
      ];
      
   progresslabel="Bounded LP calculation is completed. ";
   FBAflux
  ]
  
SetAttributes[ReducedSolutionSpace, HoldAll];
ReducedSolutionSpace[Stoichiometry_, Bounds_, objectselector_, maxmin_,PrintResult_: False] := 
 Module[{optimum, reversibles, irreversibles, negatives, 
 	revcount, irrevcount, splitcons, splitvals,	shift, feasible, 
 	ExpandedFeasibles, Svals, FBAflux, objective, cons,
 	 vars, obvars, ObSpaceTransform, ObSpaceOrigin, ObSpaceBasis, bounds, ups, downs, 
   uppercons, lowercons, OScons, OSvals, keepers, newcons, newvals, 
   redundancies, maxflux, tol = 0.0001},
 
  (* Given an FBA model, this function finds its reduced solution space 
  after applying mass balance, fixed objectives, non-  trivial range constraints
   and removing prismatic rays. 
  The result is directly stored in the global variable ReducedSS. *)
  bounds=Bounds; (* leave the bounds in the raw model untouched *)
  {cons, vars} = Dimensions[Stoichiometry];
  cons = cons + objectcount + 2*Length@bounds;
  AppendTo[Reductiontable, {"Stoichio, objective and range constraints", cons, vars, 0}];
    
  (*    STAGE 1:     REMOVE FIXED FLUXES AND SEPARATE IRREVERSIBLE/REVERSIBLE FLUXES   *)
  (*If[Norm[objectselector]>0,*)
  (* Remove artificial limits, sort irreversible reactions first and split reversibles. *)
    {irreversibles, reversibles, negatives} = Reform[Stoichiometry, bounds, objectselector, artificial];
	(* Print[{"irreversibles, reversibles, negatives",irreversibles, reversibles, negatives}];  *)
    {cons, vars} = Dimensions[Stoichiometry];
    cons = cons + objectcount + Count[Flatten@bounds, _?(Abs[#] < Infinity &)];
    AppendTo[Reductiontable, {"Remove artificial bounds, split reversibles", cons, vars, 0}];
  Svals = ConstantArray[{0, 0}, Length[Stoichiometry]];(* {value, type} pairs; type=0 => equal *)
  If[Length[openflux]>0 && Length[openflux]!=vars, FBAdone = False];
  If[!FBAdone,
  progresslabel="LP calculation for unbounded, unidirectional model. ";
  Print[Style[progresslabel, Blue, TextAlignment -> Center]];
  progresscounter:=Clock[5]; progressrange=Indeterminate;
  (* Simplex needs to be used here, because HsnJ called subsequently relies on the feasible
   point it uses being located at a vertex, not interior, so it can construct the LP basis. *)
(* Print[{"maxmin*objectselector, Stoichiometry, Svals, bounds,",maxmin*objectselector, Stoichiometry, Svals, bounds}]; *)
(*
  {LPtime, result} = Timing[FBAflux = Quiet[Check[
    LinearProgramming[maxmin*objectselector, Stoichiometry, Svals, bounds, Reals, Method-> "Simplex",
     Tolerance -> LPtol ], 
    "No LP solution for irreversible expansion of model!", 
    	{LinearProgramming::lpsnf, LinearProgramming::lpsub, LinearProgramming::lpsnfp}],
    	{LinearProgramming::lpsnf, LinearProgramming::lpsub, LinearProgramming::lpsnfp}]
    ];
*)
      {LPtime, result} = Timing[FBAflux = Check[
    LinearProgramming[maxmin*objectselector, Stoichiometry, Svals, bounds, Reals, Method-> "Simplex",
     Tolerance -> LPtol ], 
    "No LP solution for irreversible expansion of model!", 
    	{LinearProgramming::lpsnf, LinearProgramming::lpsub, LinearProgramming::lpsnfp}]
    ];
  If[PrintResult, Print["Unconstrained LP calculation took "<>ToString[LPtime]<>" seconds"]]; 
  If[StringQ[FBAflux], Return[FBAflux], 
  	 openflux = FBAflux; (* store the result in case of repeated reduction stage *)];
  progresslabel="Unbounded LP calculation is completed. ",
  FBAflux = openflux];
  objective = If[VectorQ[FBAflux], objectselector . FBAflux, 0.0];
  optimum = objectselector . feasiblepoints[[1]];
	(* Print[{"Optimum, objective",optimum,objective}]; *)
  If[Abs[optimum - objective] < 3*Max[ToExpression@StringSplit[Tolerances, ","]], 
  	(* Some models have a ridiculously large artificial upper limit, such as 10^6.
  	This leads to FBAflux having flux components with this value.
  	Prepending the properly calculated infinite limit FBAflux that avoids such components,
  	avoids numerical difficulties arising when the first feasible point is used in HSnJ 
  	to construct a Simplex basis. *)
  	PrependTo[feasiblepoints,FBAflux];
  	FBAdone=True;
    If[PrintResult,Print["Confirm that unbounded FBA calculation objective value "<>
    TextString[objective]" reproduces the artificial bounds value, while the respective 
    flux vectors differ by "<>TextString[Norm[feasiblepoints[[1]]-feasiblepoints[[2]]]]]]
     , 
    feasiblepoints={FBAflux};
    Print["Discrepancy between unbounded FBA calculation "<>TextString[objective]<>
    " and bounded objective value "<>TextString[optimum]<>
    " too large compared to LP tolerances; Continuing with new solution only."];
   ];
   If[reshuffle, AppendTo[Processreport, 
   	{Style["CAUTION - The stoichiometry matrix was randomly reshuffled, 
   		so reactions and metabolites are now in a different order from the input data!  ", Blue ]}];];
   AppendTo[Processreport, 
   	{"All points in the solution space share the objective value " <> TextString[objective]}];

   
   maxflux=Max@Abs@FBAflux;
   If[fixtol > 0.1 maxflux, fixflag=True; Print["WARNING: the fixed flux tolerance exceeds 10% of the "<>
   TextString[maxflux]<>" maximal flux component of the known, unbounded, optimal feasible point! "]];  
    
  (*
  Print["Flux space separations between feasible points "];
  Print[TableForm@Table[Chop@Norm[feasiblepoints[[i]]-feasiblepoints[[j]]],
  	{i,Length[feasiblepoints]},{j,i}]];
  *)
  (* Initialize global variables, then remove fixed fluxes *)
  {lowers, uppers} = Transpose[bounds];
   augment =  If[Norm[objectselector]>fixtol,
   	Join[Append[Stoichiometry, objectselector], 
    Transpose@{Append[ConstantArray[0, Length[Stoichiometry]], objective]}, 2],
    
    Join[Stoichiometry, Transpose@{ConstantArray[0, Length[Stoichiometry]]}, 2]
   ];
  ExpandedFeasibles=feasiblepoints;

(*******************************************************************************
  (* In case there are reversible reactions, add constraints to remove
  spurious degrees of freedom introduced by splitting them. These constraints
  fix the sampled reversible flux pairs to the diagonal line of the rectangle formed by 
  their respective upper in the 2D plane that the pair defines. 
  This allows the pair fluxes to individually vary over its full range 
  but with a fixed sum (i.e. circulation) flux.   *)  
  revcount=Length[reversibles]; irrevcount=Length[irreversibles];
  If[revcount>0,
  	splitcons = Join[Join[ConstantArray[0., {revcount, irrevcount}], 
    	IdentityMatrix[revcount], IdentityMatrix[revcount], 2]];
  	splitvals = ConstantArray[0., revcount];
  	ups = uppers /. Infinity -> Max@feasiblepoints[[1]];
  	feasible = feasiblepoints[[1]];
  	Do[	j = irrevcount + i; 
  		splitcons[[i, j]] = ups[[j + revcount]]; 
  		splitcons[[i, j + revcount]] = ups[[j]]; 
  		splitvals[[i]] = ups[[j]]*ups[[j + revcount]];
  		(* find a flux increment that when added to each reversible half, will put the 
  		known flux value on the constraint line. *)
    	shift = (splitvals[[i]]
     		- feasible[[j]]*splitcons[[i, j ]]
     		- feasible[[j + revcount]]*splitcons[[i, j + revcount]])
     	  /(splitcons[[i, j]] + splitcons[[i, j + revcount]]);
  		feasible[[j]] += shift; feasible[[j + revcount]] += shift,
   	{i, revcount}];
 	feasiblepoints[[1]]=feasible;		
 	augment=Join[augment,Join[splitcons, Transpose@{splitvals}, 2]];
  ];
*******************************************************************************)
     
 (*
   Print["Check that feasible points satisfy expanded augment matrix conditions."];
   Map[Print[Max@Abs@Chop[augment[[All,-1]]-augment[[All,;;-2]].#, tol]]&,feasiblepoints]; 
 *)
  
  HSnJtime=Timing[fixvals =HopSkipnJump[fixtol,PrintResult];];
  If[PrintResult,Print["Time taken by Hop, Skip and Jump = "<>ToString[First@HSnJtime]]];
  (*
     Print["Check that after Hop, Skip and Jump, feasible points satisfy augment matrix conditions."];
   Map[Print[Max@Abs@Chop[augment[[All,-1]]-augment[[All,;;-2]].#, tol]]&,feasiblepoints]; 
	*)
   (* Revert to reversible reactions and set up the FFS to FS affine transform. *)
  NonfixTransform = 
  Revert[Stoichiometry, objectselector, bounds, ExpandedFeasibles, objective, 
  	reversibles, irreversibles, negatives];
 (*
   Print["Check that after reversion, feasible points satisfy contracted augment matrix conditions."];
   Map[Print[Max@Abs@Chop[augment[[All,-1]]-augment[[All,;;-2]].#, tol]]&,feasiblepoints]; 
 	*)
  {cons, vars} = Dimensions[augment]; vars--;
  cons = cons + Count[Join[uppers,lowers], _?(Abs[#] < Infinity &)];
  AppendTo[Reductiontable, {"Fix " <> ToString[Length[fixvals]] <>
  	 " fluxes, revert reversibles", cons, vars, 0}];
 
  (*    STAGE 2:     
  APPLY MASS BALANCE AND OBJECTIVE CONSTRAINTS AND ELIMINATE PRISMATIC RAYS   *)
  
  (* HSnJ may have added feasible points, but discard them as they are not vertices *)
  feasiblepoints = Take[feasiblepoints, Length[ExpandedFeasibles]];
  (* Set up the objective space transform *)
  ObSpaceOrigin =feasiblepoints[[1]];
  ObSpaceBasis = Chop@N@Orthogonalize[NullSpace[augment[[All, ;; -2]], Tolerance -> LPtol], Method -> "Householder"];
  ObSpaceTransform = {ObSpaceOrigin, ObSpaceBasis};
  obvars=Length[ObSpaceBasis];
  AppendTo[Reductiontable, {"Apply mass balance and fixed objective value  ", 0, obvars, 0}]; 
  	(* Print[{"ObSpaceTransform",ObSpaceTransform}]; *)  
  If[obvars == 0,
   ObSpaceTransform[[2]] = {0.,{{0.}}}; 
   ReducedSS = {{{{1.}}, {0.}}, ObSpaceTransform};
   AppendTo[Processreport,{" No remaining degrees of freedom so RSS and SSK are both a single point. "}];

    ,
   (* Apply the non-trivial (i.e. non-infinite) flux range constaints  *)
   ups = Flatten@Position[uppers, _?(# < Infinity &), {1}];
   uppers = uppers[[ups]];
   uppercons = UnitVector[vars, #] & /@ ups;
   downs = Flatten@Position[lowers, _?(# > -Infinity &), {1}];
   lowers = lowers[[downs]];
   lowercons = -UnitVector[vars, #] & /@ downs;
   {OScons, OSvals} = DowncastConstraints[Join[lowercons, uppercons], Join[-lowers, uppers], ObSpaceTransform];
   {OScons, OSvals, keepers} = DropColinears[OScons, OSvals];
   ReducedSS = {{OScons, OSvals}, ObSpaceTransform};
   {cons, vars} = Dimensions[OScons];
   AppendTo[Reductiontable, {"Apply nontrivial range constraints ", cons, vars, 0}];
   (* Print[{"SS dimensions before Prismdrop",Dimensions/@Flatten[ReducedSS,1]}]; *)

   (* Project out prismatic ray dimensions and linealities *)
   prismrays = PrismDrop[ReducedSS,PrintResult];
   {cons, vars} = Dimensions[RSScons];
	 (*Print[{"SS dimensions after Prismdrop",Dimensions/@Flatten[ReducedSS,1]}];  *)
   If[Norm[RSSBasis]>0.,
   (*	Monitor[
   	{newcons, newvals, redundancies} = RedundancyTrimmer[RSScons, RSSvals, {}, PrintResult];
   	      , Labeled[ProgressIndicator[progresscounter/cons], " Eliminating redundant constraints "]]; *)
   progressrange={0,cons};progresslabel=" Eliminating redundant RSS constraints ";
   Print[Style[progresslabel, Blue, TextAlignment -> Center]];	 
   {newcons, newvals, redundancies} = RedundancyTrimmer[RSScons, RSSvals, {}, PrintResult];
   progresslabel=" RSS redundant constraint elimination completed."; 
   progresscounter=0;

   ReducedSS[[1]] = {newcons, newvals};
   {cons, vars} =  Dimensions[newcons];
   
   (* If all constraint values are 0, all constraint hyperplanes intersect at the current 
   origin, i.e. they define a cone without bounded facets and the SSK is simply its apex 
   i.e. a single point at the RSS origin. *)
   AppendTo[Reductiontable, {"RSS after removing redundant constraints ", cons, vars, 0}]];
   If[Length[newvals] > 0 && Total[newvals] < tol,    SStype = "SimpleCone";
   	Print["All constraint hyperplanes intersect at the origin, 
 giving a conical RSS and the SSK is its apex point."]];
  ];
  
  ] ; 
  
  SetAttributes[Reform, HoldAll];
  Reform[Stoichiometry_, bounds_, objective_, artificial_] :=
 (* Function to reformulate model with fluxes positive and without artificial limits.
 Reactions are rearranged and the original positions of irreversibles and reversibles are returned. *)
 Module[{negatives, reversibles, irreversibles},
  (* Replace all artificial limits with infinity *)
  bounds = bounds /. {_?(# >= artificial &) -> Infinity, _?(# <= -artificial &) -> -Infinity};
  (* Reverse any reaction with both bounds negative *)
  negatives = Flatten@Position[ bounds, {lb_, ub_} /; Sign[lb] < 0 && Sign[ub] <= 0];
  If[negatives!={}, 
  (*Print["The following reactions are reversed to ensure positive flux values:"<>ToString[negatives]];*)
  Stoichiometry[[All, negatives]] = -Stoichiometry[[All, negatives]];
  feasiblepoints[[All, negatives]] = -feasiblepoints[[All, negatives]];
  objective[[negatives]] = -objective[[negatives]];
  bounds[[negatives]] = Reverse[-bounds[[negatives]], 2];
  ];
   (* In case of reversible reactions, split them into two opposing reactions.
  The stoichiometry matrix columns are rearranged so the \
irreversibles are first, followed by the set of reversibles in original directions, 
  follwed by the set of their reverses. *)
  reversibles = Flatten@Position[bounds, {lb_, ub_} /; Sign[lb]*Sign[ub] < 0];
  irreversibles = Complement[Range[Length@objective], reversibles];
  If[Length[reversibles] > 0,
   Stoichiometry = Join[Stoichiometry[[All, irreversibles]], 
     Stoichiometry[[All, reversibles]], -Stoichiometry[[All, reversibles]], 2];
   objective = Join[objective[[irreversibles]], 
     objective[[reversibles]], -objective[[reversibles]]];
   feasiblepoints = Join[feasiblepoints[[All, irreversibles]], 
     feasiblepoints[[All, reversibles]] /. _?(# < 0 &) -> 0., 
     -feasiblepoints[[All, reversibles]] /. _?(# < 0 &) -> 0., 2];
   bounds = Join[bounds[[irreversibles]], bounds[[reversibles]] /. _?(# < 0 &) -> 0., 
     Reverse[-bounds[[reversibles]] /. _?(# < 0 &) -> 0., 2]]
   ];
    {irreversibles, reversibles, negatives}
  ]

SetAttributes[Revert, HoldAll];
Revert[Stoichiometry_, objectselector_, bounds_, feasibles_, 
  objective_, reversibles_, irreversibles_, negatives_] := 
 Module[{irrcount, revcount, fixes, nonfixes, order, inverse, 
   fluxcount, revfixvals, revfixes, fixpairs, singles, pairs, rect, 
   shift, constraints, values, nonzerorows, nonfixorigin, nonfixbasis},
  (* Function to reconsolidate opposing reaction pairs and revert to 
	a variable flux space of original reactions, 
  	but with irreversible reactions first followed by reversible ones. 
  	It returns the transformation from FFS to this FS. *)
  
  order = Join[irreversibles, reversibles];
   (* inverse is the ordering required to regain the unsorted order *)
  inverse = Ordering[order];
  {irrcount, revcount} = Length /@ {irreversibles, reversibles};
  (*Print[{irrcount,revcount,irrcount+revcount}];*)
  fluxcount = irrcount + revcount;
  fixes = fixvals[[All, 1]];
  nonfixes = Complement[Range[fluxcount], fixes];
  (*Print[{fluxcount,irrcount,revcount,fixvals}];*)
  
  (* As implemented, HSnJ progressively reduces the flux space by eliminating 
  fixed fluxes, modifying augment and bounds matrices as it goes.
  That is fine for irreversibles reactions but in the case of reversibles, 
  particularly where only one leg of it is found fixed, the elimination upsets 
  the assumed pairing and hence ordering of fluxes. To deal with that, the code below
  in effect discards the matrices delivered by HSnJ and reconstitutes augment etc. 
  from the stoichiometry matrix. The feasibles argument needs to be the feasible fluxes 
  before HSnJ was let loose on them. Note that after step 6, it gets rearranged if the 
  compatibility test between S and feasibles is carried out.  *)
  
  If[revcount > 0,
   (* Some of the fixed fluxes found, belong to reversible reactions. 
   Step 1: reunite the reversible pairs in the expanded matrices. *)
   Stoichiometry = Drop[#, -revcount] & /@ Stoichiometry;
   bounds = Join[bounds[[;; irrcount]], 
     bounds[[irrcount + 1 ;; irrcount + revcount]] - 
      Reverse[bounds[[-revcount ;;]], 2]];
   objectselector = Drop[objectselector, -revcount];
   feasibles = Join[feasibles[[All, ;; irrcount]], 
     feasibles[[All, irrcount + 1 ;; irrcount + revcount]] - 
      feasibles[[All, -revcount ;;]], 2];
 (*     
   Print["Check that after Step 1, feasible points satisfy S matrix conditions."];
   Map[Print[Max@Abs@Chop[Join[ConstantArray[0.,Length[Stoichiometry]],{objective}]-
   	Join[Stoichiometry,{objectselector}].#]]&,feasibles]; 
 *)       
   (* Step 2: separate reversible fixed pairs and singles 
   		and apply consolidation rules *)
   If[Max[fixes] > irrcount,
    revfixvals = Select[fixvals, #[[1]] > irrcount &];
    revfixes = revfixvals[[All, 1]];
    fixpairs = Map[Which[ 
    	MemberQ[revfixes, # + revcount],
    	 {FirstCase[revfixvals, {#, _}], FirstCase[revfixvals, {# + revcount, _}]}, 
        MemberQ[revfixes, # - revcount], Nothing, 
        True, FirstCase[revfixvals, {#, _}]] &, revfixes]; 
    singles = Select[fixpairs, Dimensions[#] == {2} &];
    pairs = Select[fixpairs, Dimensions[#] == {2, 2} &];
    (*Print[{singles,pairs}];*)
    
    If[Length[singles] > 0,
     If[Norm[singles[[All, 2]]] > 0, 
      Print[Style["A nonzero fixed value was found for only one member of a reversible flux pair.
       That is not plausible, tolerance for fixed values probably too lax. - Revert quits!",Red]];
        Abort[]];
     (* If the forward direction is fixed to zero flux, 
     reaction proceeds backwards and upper bound is zero , 
     else just put lower bound zero *)
     Map[(rect = #[[1]]; If[rect <= irrcount + revcount,
         bounds[[rect,2]] = 0., bounds[[rect - revcount, 1]] = 0.]) &, singles];
     fixdirs = Map[If[#[[1]] <= irrcount + revcount, 
     			  {#[[1]], "backwards"},
     			  {#[[1]] - revcount, "forwards"}] &, singles];
     rect = fixdirs[[All, 1]];
     revfixvals = {}];
     
    (* Step 3: Consolidate fixed values in pairs and 
    	set them into the global list of fixed values*)
    If[Length[pairs] > 0,
    	(* The next line was an attempt to filter out spurious fixed fluxes that result from adding 
    	line constraints to remove the futile fluxes introduced by reaction splitting. 
    	But it doesnot seem to achieve the goal.  *)
    (*	pairs=Select[pairs, 
    		#[[1, 2]]*#[[2, 2]] < 10.^-4 ||Abs[#[[1, 2]] - #[[2, 2]]] < 0.01 &];*)
     	revfixvals = Map[{#[[1, 1]], #[[1, 2]] - #[[2, 2]]} &, pairs]];
   
    fixvals = Join[Select[fixvals, #[[1]] <= irrcount &], revfixvals];
    fixes = fixvals[[All, 1]];
    nonfixes = Complement[Range[fluxcount], fixes];
    (*Print[fixvals];*)
    (*Print[fixdirs];*)
    ];
   
     (* Step 4: Revert all reactions that were reversed because their fluxes were negative  *)
  If[negatives!={}, 
  negatives=inverse[[negatives]]; (* convert the original negatives to the sorted order *)
  Stoichiometry[[All, negatives]] = -Stoichiometry[[All, negatives]];
  feasibles[[All, negatives]] = -feasibles[[All, negatives]];
  objectselector[[negatives]] = -objectselector[[negatives]];
  bounds[[negatives]] = Reverse[-bounds[[negatives]], 2];
  fixvals=Replace[fixvals, fv_?(MemberQ[negatives, #[[1]]] &) :> 
  	{fv[[1]], -fv[[2]]}, {1}];
  (* Note that only irreversibles can be a negative, so reverting negatives do not affect 
  fixdirs that can only apply to a reversible reaction. *)
  ];
  
   (* Step 5: 
   Recalculate global arrays after removal of fixed fluxes *)
   (*Substitute fixed values, then remove those columns by shifting to the RHS*)
   shift = ConstantArray[0., fluxcount];
   Map[(Part[shift, #[[1]]] = #[[2]]) &, fixvals];
   If[Norm[objectselector]>10^-6,
   	constraints = Append[Stoichiometry, objectselector];
   	values = Append[ConstantArray[0, Length[Stoichiometry]], objective],
   	constraints = Stoichiometry;
   	values = ConstantArray[0, Length[Stoichiometry]]
   ];
   values -= constraints . shift;
   constraints = constraints[[All, nonfixes]];
   (*Remove constraints that are trivial,
   due to removal of columns or a silly original model*)
   nonzerorows = 
    Flatten@Position[Normal@constraints, _?(Norm[#] > 0 &), {1}];
   If[nonzerorows != {}, constraints = constraints[[nonzerorows]];
    values = values[[nonzerorows]]];
   augment = 
    SparseArray@Chop[Join[constraints, Transpose[{values}], 2], 10^-6];
   {lowers, uppers} = Transpose[bounds[[nonfixes]]];
   feasiblepoints = feasibles[[All, nonfixes]];
     	
   (* Step 6: Revert the 3 model matrices to their origininal flux ordering and 
   reassign fixvals and fixdirs, according to the original FFS ordering *)
   bounds=bounds[[inverse]];
   objectselector=objectselector[[inverse]];
   Stoichiometry=Stoichiometry[[All,inverse]];
   fixvals[[All, 1]] = order[[fixes]];
   If[Length[fixdirs]>0,fixdirs[[All, 1]] = order[[rect]]];

   ];
  (* 
  (* Note that the local variable feasibles is in FFS, while the global feasiblepoints
  is in FS with rearranged flux ordering and fixed fluxes removed *)
  feasibles=feasibles[[All,inverse]]; 
   Print["Check that after Step 6, feasible points satisfy S matrix conditions."];
   Map[Print[Max@Abs@Chop[Join[ConstantArray[0.,Length[Stoichiometry]],{objective}]-
   	Join[Stoichiometry,{objectselector}].#]]&,feasibles]; 
 *)  
  (* Step 7:  set up the transformation from FFS to FS *)
  nonfixorigin = ConstantArray[0., fluxcount];
  nonfixorigin[[fixes]] = fixvals[[All, 2]];
  nonfixorigin = nonfixorigin[[inverse]];
  nonfixbasis = IdentityMatrix[fluxcount][[order]];
  nonfixbasis = nonfixbasis[[nonfixes]];
   
  {nonfixorigin, nonfixbasis}
  ]
  
(* SetAttributes[KernelFinder,HoldAll]; *)
KernelFinder[PrintResult_: False] := 
 Module[{ProgenTreesearch=None, FBFTreesearch=None, count, levelcounts, 
   cons, vars, ssdim, raydim, centershift, result = "c"},
   Kerneltable={};
   (*This function controls the calculation of the Solution Space Kernel or SSK*)
  
  (*  STAGE 1: DIAGNOSE THE NATURE OF THE RSS AND DEAL WITH SIMPLER CASES FIRST*)
  
  If[Norm[RSSBasis] ==  0.,                 (* CASE 1 - a single point *)
   SStype = "Point";
   (*Print["No degrees of freedom remain after removing prismatic rays from the SS.
      So the SSK is just the single point at the SS origin."];*)
   AppendTo[Kerneltable, {"SSK = RSS, a single point at the origin", 0, 0, 0}];
   Return[]
   ];
  
  {raymat, raydim} = CompleteRayBasis[RSScons, PrintResult];
  (*Print["Ray dimensions after CompleteRayBais "<>ToString[
  Dimensions@raymat]];*)
  RayDim = RayDim + raydim;
  {cons, vars} = Dimensions[RSScons];
  KernelSpace = ReducedSS;
  (*KernelSpace initially the same as RSS, but respecify it relative to RSS,rather than FS,
  to reduce dimensions for capping and centering procedures*)
  ssdim = Length[RSSBasis];
  KernelSpace[[2]] = {ConstantArray[0., ssdim], IdentityMatrix[ssdim]};
  
  If[raydim == 0,                              (*  CASE 2 - a bounded polytope *)
   SStype = "Compact";
   (* Just recenter the Kernel space to an interior point *)
   centershift=MiniMaxcentre[Kernelcons,Kernelvals, PrintResult];
   KernelSpace[[1,2]] = Kernelvals - Kernelcons . centershift;
   KernelSpace[[2,1]] = KernelOrigin + centershift . KernelBasis;
   AppendTo[Kerneltable, {"RSS is closed, no capping done", cons, vars, 0}];
   ];
  
  apex = PseudoInverse[Kernelcons] . Kernelvals;
  (* Print[{"Test for simple cone ",apex,Norm[Kernelcons.apex-Kernelvals]}]; *)
  If[Norm[Kernelcons . apex - Kernelvals] < LPtol,        (* CASE 3 - a simple cone *)
   SStype = "SimpleCone";
   KernelSpace[[2, 1]] = apex;
   KernelSpace[[1, 2]] = ConstantArray[0., cons];
   AppendTo[Kerneltable, {"RSS a simple cone, capped with default radius", cons, vars, 0}]
   ];
  
  (*STAGE 2: DETERMINE THE PROGENITOR FACET AND DO COINCIDENCE CAPPING *)
  
  If[SStype == "FacetCone",              (*  CASE 4 - a cone with bounded facets *)
  	
  	If[vars > RSSdims, result = ChoiceDialog[
   "The RSS has more  than " <> ToString[RSSdims] <> 
    " variables, making FBF search unviable. Either:\n
a) Abort, then repeat stage 2 with an increased fixed tolerance to reduce the \
variable count.\n
b) Drastically reduce the sample sizes, e.g. to \
a value between 1 and 3 for the progenitor, and 1 to 10 for the BFBF \
search, then press Continue.\n
c) Adjust the default capping radius to \
a plausible value, then Skip FBF to apply default tangent capping.", \
{"Abort" -> "a", "Continue" -> "c", "Skip FBF" -> "s"}]];
	Switch[result, 	"a", Abort[], 
					"s", SStype = "SimpleCone", 
			  		"c", If[vars > RSSdims, treesize=1000; 
			  				Print[Style["Viable tree size reduced to 1000",Darker@Green]]];
   {ProgenTreesearch, progen} = 
   		Progenitor[raymat, raydim, Kernelcons, Kernelvals, targetcount, PrintResult];
   (*Print["Ray dimensions after progen "<>ToString[Dimensions@raymat]];*)
   levelcounts = SortBy[Tally[Length /@ BoundedFacets], First];
   If[PrintResult, Print["Relative to RSS, the BFBF levels and their member counts
   are as follows:\n" <> ToString[levelcounts]]];
   If[PrintResult, Print["Their progenitor is at level " <> ToString[Length@progen]]];
   (* Note: CoincidenceCapper updates raymat,raydim, BoundedFacets and Infeasibles*)
   If[OrthoRayQ[raymat, Kernelcons, progen], 
    coincrays = CoincidenceCapper[KernelSpace, raymat, raydim, progen, PrintResult];
    {cons, vars} = Dimensions[Kernelcons];
    AppendTo[Kerneltable, {"Coincidence capped to level " <> ToString[Length@progen] <> 
    	" progenitor facet ", cons, vars, Length[coincrays]}]
    , 
    coincrays = {}; 
    {cons, vars} = Dimensions[Kernelcons];
    Print["Coincidence capping was skipped because there is no ray
    orthogonal to the progenitor found at level " <> ToString[Length@progen]];]
    ];
   (*Print["Ray dimensions after coinc "<>ToString[Dimensions@raymat]];
   Print["Dimensions of ray space = "<>ToString[raydim]];*)
   
   (*STAGE 3:   CHECK IF COINCIDENCE CAPPING REDUCED THE SSK TO CASES 2 OR 3;
   	OTHERWISE FIND (MOST) FBF's. THEN DO TANGENT CAPPING*)
   	(* Note that coincidence capping cannot produce case 1, a single point. 
   	   For that to happen, the progenitor would have to be a single point; 
   	   but a point has no subfacets, so it cannot be the progenitor of FBF's. *)
   
   {cons, vars} = Dimensions[Kernelcons];
   If[raydim == 0, SStype = "Compact";              (* CASE 2 - a bounded polytope *)
   	   (* Recenter the Kernel space to an interior point *)
  	centershift=MiniMaxcentre[Kernelcons,Kernelvals, PrintResult];
   	KernelSpace[[1,2]] = Kernelvals - Kernelcons . centershift;
   	KernelSpace[[2,1]] = KernelOrigin + centershift . KernelBasis;
    AppendTo[Kerneltable, {"SSK is closed, no tangent capping needed", cons, vars, 0}]
   ,
   apex = PseudoInverse[Kernelcons] . Kernelvals;
   If[Norm[Kernelcons . apex - Kernelvals] < LPtol,        (* CASE 3 - a simple cone *)
    SStype = "SimpleCone";
    KernelSpace[[2, 1]] = apex;
    KernelSpace[[1, 2]] = ConstantArray[0., cons];
    AppendTo[Kerneltable, {"SSK a simple cone, capped with default radius", cons, vars, 0}]]
    ];
   ];
   
  Switch[ProgenTreesearch,
  	True, AppendTo[Processreport, {"Exhaustive search visiting "<>ToString[totalfacets]<>" facets found "<>
  		ToString[FBFcount]<>" BFBF's, to establish the progenitor facet. "}],
   	False, AppendTo[Processreport,{"Randomised greedy search sampled "<>ToString[totalfacets]<>" facets to find "<>ToString[FBFcount]<>
  		" BFBF's, in establishing the progenitor facet. "}],
   	_,Null];
  If[PrintResult,Print["Proceed to tangent capping."]];
 
  If[SStype == "FacetCone",
   {FBFTreesearch, count} = FBFfinder[raymat, raydim, Kernelcons, Kernelvals, samplesize, 
     treesize, PrintResult];
   If[PrintResult, Print["Ray dimensions after full BFBF search " <> 
      ToString[Dimensions@raymat]]];
   levelcounts = SortBy[Tally[Length /@ BoundedFacets], First];
   If[PrintResult, If[FBFTreesearch, 
    Print["Exhaustive list of "<>ToString[FBFcount]<>" BFBF's was found by a tree search."], 
    Print[ToString[FBFcount] <> " BFBF's were collected by random sampling.  "]]];
   ];
  
  If[SStype == "SimpleCone" || SStype == "FacetCone",
     conerays = TangentCapper[KernelSpace, raymat, RSSBasis, PrintResult];
   (*Print["Ray dimensions after TangentCapper "<>ToString[Dimensions@raymat]];*),
  	conerays={};
   ];
   {cons, vars} = Dimensions[Kernelcons];
    
  (* BY WHICHEVER ROUTE, THE KernelSPACE IS NOW BOUNDED AND CENTERED AT INSCRIBED HYPERSPHERE. *)
  
  Switch[FBFTreesearch,
  	True, AppendTo[Processreport, {"The tangent capping search found "<>ToString[FBFcount]<>
  		" BFBF's by an exhaustive search, covering "<>ToString[totalfacets]<>" facets."}],
   	False, AppendTo[Processreport,{"The tangent capping search found "<>ToString[FBFcount]<>
  		" BFBF's, by a random greedy search, sampling "<>ToString[totalfacets]<>" facets."}],
   	_,Null];
   	
  If[fixflag, AppendTo[Processreport, {"\nCAUTION: The chosen fixed tolerance " <> ToString[fixtol] <>
  	" is more than 10% of the largest flux component of the
  	 known (unbounded) optimal flux vector, which has length "<>TextString[Norm@First@feasiblepoints],
	"This may be plausible, if the tolerance is small compared to the chord lengths and enclosing radii. \n"   }]];
  
 ]

SetAttributes[KernelShape, HoldAll];
KernelShape[ maxaspect_, maxthin_, PrintResult_: False] := 
 Module[{cons,vars, enclose, ratio, volratio, coverage, thintext, mindiam, diams, 
 	rad1, rad2, meandiam, meanchord, EnvelopeIndiam, EnvelopeMaxdiam, diamrange,
 	aspectmin, tol = 0.00001},
 	(* This function characterises the shape of a closed SSK polytope. 
 	Using the main chords/diameters, the coordinates are recentered.
 	When appropriate, minor directions are flattened out.
 	The final recentering delivers a set of peripheral points.
 	The coverage of the complete SSK that the peripheral point polytope 
 	achieves, is estimated.
 	Space specifications and reference points are transferred up to the FFS.
 	Finally, the results achieved are entered into the summary report.
 	 *)
 	{cons, vars} = Dimensions[Kernelcons]; 
 	 
  thindirs = FlatnCentre[KernelSpace, maxaspect, maxthin, PrintResult];
 Print["Check that periphery points satisfy interior requirement: ",
	outside=Chop@Min@Map[(Kernelvals - Kernelcons . #) &, PeriPoints];
	If[outside >= -tol, True, "Outside by margin "<>TextString[Abs@outside]]]; 
  progressrange={0,10};progresslabel=" Reorienting enclosing simplex. ";
  Print[Style[progresslabel, Blue, TextAlignment -> Center]];
  enclose = WrappingRadii[KernelSpace, PeriPoints, PrintResult];
  progresslabel=" Completed estimation of enclosing simplex radii."; 
  progresscounter=0;
  
  If[enclose[[2]] > 0. ,
  ratio=Divide@@enclose;
  (* Print[{"Kernelspace counts of constraints and variables are "<>ToString[{cons,vars}]}]; *)
  volratio = Chop[ratio^vars,0.000001];
  coverage = If[volratio > 0.01, 
  	progressrange={0,1000};
  	progresslabel=" Sampling random SSK points to determine coverage. ";
  	Print[Style[progresslabel, Blue, TextAlignment -> Center]];
    CoverageInterval[KernelSpace, PeriPoints], {0, 0}];
   progresslabel=" SSK coverage determination completed."; 
   progresscounter=0;
  ]; 
  
  (* COMPLETE THE SUMMARY REPORT *)
  mindiam = 2*First@InscribedSphere;
  thintext=If[Length[thindirs]>0,", after flattening "<>ToString@Length[thindirs]<>" dimensions,",""];
  AppendTo[Processreport,{"The maximal inscribed hypersphere diameter"<>thintext<>" is "<>
  	TextString[mindiam]}];
  aspectmin=Min[maxthin, Mean[Abs@ChordLengths]/maxaspect];
  If[mindiam < aspectmin, AppendTo[Processreport,
  	{Style["This value is below the threshold targeted to be flattened out.", Blue],
  	"So some thin diameters may remain unidentified and flattening may need to be repeated or made more stringent. "}]];
  If[enclose[[2]] > 0. ,
  AppendTo[Processreport, {"The diameter and volume coverage ratios, between mutually similar simplices
  that encloses the  periphery points or the complete SSK, are "  <> 
     TextString@Quiet@NumberForm[100*{ratio, volratio}, 3] <> "% respectively."}]];
     
  diams = Length[PeriPoints]/2;
  rad1 = Norm /@ PeriPoints[[;; diams]];
  rad2 = Norm /@ PeriPoints[[-diams ;;]];
  Diameters=rad1+rad2;
  meandiam = Mean[Diameters];
  meanchord = Mean[Abs@ChordLengths];
  {EnvelopeIndiam, EnvelopeMaxdiam} = If[enclose[[2]] > 0. ,
 		{2*enclose[[2]], (vars+1)*enclose[[2]]}, {0.,0.}];
  diamrange = {Max[meandiam,EnvelopeIndiam/vars], meandiam/ratio, Min[meanchord, EnvelopeMaxdiam]};
  If[diamrange[[1]]<=diamrange[[2]]<=diamrange[[3]],Null,
  	diamrange[[3]]=EnvelopeMaxdiam;
  	(* A negative chordlength was estimated using diameters, then this test does not apply *)
  	If[Min[ChordLengths] >= 0, 
  		AppendTo[Processreport,{Style["Caution - diameter estimate is larger than mean chord length.
  Flattening may resolve this.",Blue]}]]];
  AppendTo[Processreport,{"The mean SSK diameter is estimated to be in the range "<>
  	TextString[{diamrange[[1]],diamrange[[3]]}] <> "\n and the best value estimate is " <>
  	TextString[diamrange[[2]]]}];
     
  If[coverage != {0, 0}, 
   AppendTo[Processreport, {"The sampled fraction of the SSK spanned by the
   peripheral point polytope, is in the 95% confidence interval " <> 
      ToString[coverage]<>"%"}]
   ];
   
 If[Length[thindirs]>0, 
 	"Flattened "<>ToString[Length[thindirs]]<>" dimensions.",
 	"No flattening performed."]
 ]
 
SetAttributes[KernelDisplay, HoldFirst];
KernelDisplay[exportdata_, Printresult_:False] := 
 Block[{exfeasibles, exrays, exSolutionSpace, exSSTransform, 
   exKernelSpace, exKernelTransform, exPeriPoints, exISphereCentre, 
   exthindirs, exfixvals, fixdirnames, BaseDim, chordpic, cenpic, discrepancies, 
   warn = Style["\nWARNING!", Red, Bold], warning = ""},
    (*This function finalizes the report and produces plots of chords and centrality.
  	It also transfers results back to its representation \
	in the full flux space (FFS) and stores these in the ExportData structure.*)

  (* PERFORM THE VALIDATION TEST ON THE VARIABLE FLUX SPACE FEASIBLE POINTS *)
  If[SStype != "Point", 
  	progressrange={0,2*Length[feasiblepoints]+1};
  	progresscounter = 1;
  	progresslabel = InsertLinebreaks["Performing Validation Tests "];
  	Print[Style[progresslabel, Blue, TextAlignment -> Center]];
   	If[Printresult,Print["Vector norms of flux, reconstitued flux, kernel part, raypart and shortfall: "]];
  	Validations = TimeConstrained[If[Length[thindirs]==0,
  		{Map[KernelDiscrepancy[#, KernelSpace, Printresult] &, feasiblepoints]},
  		{Map[KernelDiscrepancy[#, SSKinRSS, Printresult] &, feasiblepoints], 
   		Map[KernelDiscrepancy[#, KernelSpace, Printresult] &, feasiblepoints]}],
   		timeconstraint];
   	If[Validations==$Aborted, Print["Validation step aborted as it exceeded the step time limit."]];
  ];
  
  (*
  Print["Diagnostic: Peripoints, dimensions of SS, SSK and NonfixTransform"];
  Print[Norm/@PeriPoints];
  Print[Dimensions/@Flatten[ReducedSS,1]];
  Print[Dimensions/@Flatten[KernelSpace,1]];
  Print[Dimensions/@NonfixTransform];
  *)
  
  (*TRANSFER SPECS BACK TO FULL FLUX SPACE IN ORDER TO UPDATE FIXED VALUES*)
  (* There are 4 spaces used internally: Full Flux Space (FFS), variable space (VS) 
  	 i.e. fixed fluxes eliminated, Reduced Solution Space (RSS) and the Solution Space
  	 kernel (SSK). Each of these is a subspace of the previous in this sequence.
  	 
  	 Internally, RSS and SSK specs are relative to VS because it cuts the dimension requirement.
  	 For export, however, only SSK and perhaps RSS are relevant and shoud be given relative to FFS.
  	 Note that to change the reference frame, it is just the Transform part of the space spec that
  	 needs to be changed, the constraints and values refer to their local coordinate origin (by 
  	 design!) and dimension count and are NOT changed.
  	
  	 NonfixTransform gives the transformation from FFS to VS.
  *)
  exfeasibles = Chop[UpliftPoint[feasiblepoints, NonfixTransform], 10.^-6];
  exrays =DeleteDuplicates[Join[prismrays, coincrays, conerays], (1 - Abs[#1 . #2] < 10.^-6) &];
  If[Length@exrays>0,
  	exrays = Chop[exrays . NonfixTransform[[2]], 10.^-6];
    BaseDim = MatrixRank[exrays, Tolerance -> LPtol],
  	BaseDim = 0];
  exSolutionSpace = ReducedSS;  exKernelSpace = KernelSpace;
  exSSTransform = CombineTransform[NonfixTransform, RSSTransform];
  exSolutionSpace[[2]] = Chop[exSSTransform, 10^-6];

  If[SStype == "Point",
   exKernelSpace = exSolutionSpace;
   exKernelTransform = exSSTransform;
   exPeriPoints = {exKernelTransform[[1]]};
   ,
   exKernelTransform = CombineTransform[exSSTransform, KernelTransform];
   exKernelSpace[[2]] = Chop[exKernelTransform, 10^-6];
   exPeriPoints = 
    Chop[UpliftPoint[PeriPoints, exKernelTransform], 0.000001];
   ];
  exISphereCentre = 
   Chop[UpliftPoint[Last@InscribedSphere, exKernelTransform], 0.000001];
  exthindirs=If[thindirs != {},  Chop[thindirs . exSSTransform[[2]], 10^-6], {}];
  (*Print[Dimensions/@{exthindirs,exKernelBasis,exSSBasis,
  NonfixTransform[[2]]}];*)
  fixdirnames=fixdirs;
  If[Length[fixdirs] > 0, 
   fixdirnames[[All,1]] = ReactionNames[[fixdirs[[All, 1]]]];
   Processreport = Join[Processreport,
   	{{"The following reactions acquired fixed directions:"},
     {TextGrid@Partition[fixdirnames,UpTo@5]}}]
     ];
  exfixvals = 
   SSKfixvals[exKernelTransform[[1]], exPeriPoints, exrays, fixtol];
  AppendTo[
   Processreport, {ToString[Length[PeriPoints]] <> " peripheral points were found. 
   Assuming these to be representative, and combining with rays, extends
     	 the fixed value list to " <> ToString[Length[exfixvals]] <> " items"}];
  AppendTo[
   Processreport, {"The combined set of " <> 
     ToString[Length[exrays]] <> " rays span " <> ToString[BaseDim] <>
      " of the total " <> ToString[RayDim] <> " ray dimensions."}];
 
  If[Length[chordpic] == 0 && Length[PeriPoints] > 1,
     {chordpic, cenpic} = Centrality[KernelSpace, ChordLengths, PeriPoints];
    discrepancies = Count[Diameters/Abs@ChordLengths, _?(# > 1.1 &)];
    If[discrepancies > 0, warning = 
     "There are " <> ToString[discrepancies] <> 
      " diameters through the current origin, that exceed their corresponding 
putative chord lengths by more than 10%.
Chord approximation (especially by diameters) gives a lower limit only, underestimating in these cases. 
Repeating and/or reducing flattening, or using more LP chords, may avoid this.\n";
    Processreport = Join[Processreport, {{warn}, {warning}}]];
   , 
     {chordpic, cenpic} =
     	{Graphics@Text["No chords to show."],Graphics@Text["No diameters to show."]}
    ];

    exportdata = {exKernelSpace, exPeriPoints, exfixvals, exthindirs, exrays, 
    	{InscribedSphere[[1]], exISphereCentre}, 
    	fixdirnames,{MetaboliteNames, ReactionNames}, exSolutionSpace };
  
   progresslabel =  " Validation, preparing export data and display completed. ";
   Print[Style[progresslabel, Blue, TextAlignment -> Center]];
   progresscounter=0;

  
  {chordpic, cenpic}]
 
 KernelDiscrepancy[feasible_, Kernelspace_, Printresult_:False] :=
  (* Function to evaluate the discrepancy between a feasible point in Variable Flux Space
  and its optimal reconstitution from a Kernelspace vector and a ray vector, as calculated 
  by LP in Deconstructor. 
  It returns the percentage difference in total flux, and the angle in degrees 
  between  the given and reconstituted flux drections. *)
  
  (* NOTE: This code assumes that Kernelspace is specified relative to RSS. So it has 
  	to be called before the specification is transferred to FSS *)
  Module[{RSSfeasible, OrthoRay, result, Kernelpart, raypart, shortfall, 
    reconstitute, fluxnorm, reconorm, fluxdif, deviation
    (*,meanflux, meanrecon, nonzerofluxes*) },

   (* Do the deconstruction in low-dimensional Kernel space *)
   RSSfeasible = DowncastPoint[feasible, RSSTransform];
   OrthoRay = Chop[feasible - UpliftPoint[RSSfeasible, RSSTransform]];
   
   result= Deconstructor[RSSfeasible, ReducedSS, Kernelspace, True];
  (* Print[{"In KernelDiscrepancy, on return from Deconstructor, the result is: ",result}]; *)
   If[Length[result]==3,{Kernelpart, raypart, shortfall} = result,Return[{0.,"Failed","Failed"}]];
   (* Transfer the Kernel flux up to the RSS; 
   note that the ray part is not a vector fixed to the origin, 
   so is just recast as a vector in the higher dimension*) 
   Kernelpart = UpliftPoint[Kernelpart, RSSTransform];
   raypart = raypart . RSSBasis;
   (*shortfall= shortfall.RSSBasis;*)
   reconstitute = Kernelpart + raypart + OrthoRay;
   progresscounter++;
   If[Printresult,
   	Print[Round[Norm[#],0.01]&
   			/@{feasible,reconstitute,Kernelpart,raypart+OrthoRay,shortfall}]];
   fluxnorm = Max[Norm@feasible,10.^-4]; 
   reconorm = Max[Norm@reconstitute,10.^-4];
   fluxdif = Chop[100*Abs[fluxnorm-reconorm]/Sqrt[fluxnorm*reconorm], 10.^-6];
   deviation = If[fluxnorm > LPtol && reconorm > LPtol,
    Chop[VectorAngle[feasible, reconstitute]*(180/Pi), 10.^-6],0.];
    
    (* Old version
        (* The norm of the flux vector is not a good measure of total flux, e.g. for a
        linear pathway consisting of many reactions they will all add to the norm, but
        are really the same flux flowing in sequence through them all. Mean flux per reaction
        seems better. But for just comparing original and reconstituted, comparing the
        vectors directly (i.e., norm and direction) seems OK.
        
    Mean flux taken over all fluxes that are  NONZERO in the given feasible point. 
    Including zero components in the mean does not give a reasonable account of the flux in e.g. a 
    pathway that only covers a fraction of all model fluxes. And also, taking the 
    nonzero criterion independentlly in both feasible and reconstitute, does not work well, because
    if reconstitute picks up negligible additional fluxes, these contribute to the 
    denominator in the mean so they can distort the comparison of mean values. *)
   nonzerofluxes=Position[feasible,_?(Abs[#] > 0.1*fluxnorm &)]; 
   {meanflux, meanrecon, fluxdif} = If[Length@nonzerofluxes==0, {0.,0.,0.},
   	   {Mean[Abs@Extract[feasible, nonzerofluxes]],
   		Mean[Abs@Extract[reconstitute, nonzerofluxes]], 
   		Chop[100*Abs[meanflux-meanrecon]/Sqrt[meanflux*meanrecon], 10.^-6]}];
   {meanflux, fluxdif, deviation} = Round[{meanflux, fluxdif, deviation}, 0.1];
   If[meanflux==0, fluxdif=If[fluxdif<0.1,"Zero","NonZero"]; deviation="Indeterminate"];
   {meanflux, fluxdif, deviation}
   *)
   
   {fluxnorm, fluxdif, deviation} = Round[{fluxnorm, fluxdif, deviation}, 0.1];
   If[fluxnorm == 0, fluxdif=If[NumberQ[fluxdif] && fluxdif<0.1,"Zero","NonZero"]; deviation="Indeterminate"];
   {fluxnorm, fluxdif, deviation}
   ];
 
 SaveResults[results_, datafile_] := 
 Module[{MainHeading, SectionHeadings, exportfile, savepic, pages, 
   table, reportnb, success = True},
  progressrange = {0, 2};
  exportfile = FileBaseName[datafile] <> " SSKernelReport.pdf";
  exportfile = FileNameJoin[{DirectoryName[datafile], exportfile}];
   progresscounter = 1;
   progresslabel = InsertLinebreaks["Exporting report to file " <> exportfile];
   Print[Style[progresslabel, Blue, TextAlignment -> Center]];
   (* Getting the ImageSizes right in the following, so that none of the text 
   gets cut off in the PDF file  that is eported, is very tricky. It seems to work 
   best to get actual numerical values. The final Scaled value for the compound graphic
   is mainly to keep the whole picture on a single page. *)
   savepic =  GraphicsColumn[{Show[flatplot, FrameLabel -> 
     Style["FLATTENING DECISION PLOT ", Bold, Darker[Green], 
      FontSize -> 12], ImageSize -> { 480, 160}], 
   Show[chordpic, ImageSize -> {720, 320}], 
   Show[cenpic, ImageSize -> {720, 320}]},(*AspectRatio->1.4,*)
  ImageSize -> Scaled[1], Frame -> All];
    
   pages = {ExpressionCell[table = Join[Reductiontable, Kerneltable];
      If[Length[Processreport] > 0 || Length[table] > 0, 
       Column[{Style[Heading[[1]], "Subtitle"],
         Style[Heading[[2]], "Subsubtitle"], 
         Row[{TableForm[
            Optiontable[[;; Ceiling[Length[Optiontable]/2]]], 
            TableSpacing -> {1, 3}], 
           TableForm[Optiontable[[-Floor[Length[Optiontable]/2] ;;]], 
            TableSpacing -> {1, 3}]}, "      "], 
         If[Length[table] > 1, 
          TableForm[table[[2 ;;, 2 ;;]], 
           TableHeadings -> {table[[2 ;;, 1]], table[[1, 2 ;;]]}], 
          Null], 
          TableForm[Flatten@Processreport],
	 If[Length[Validations] > 0, 
 		Labeled[Grid[{
 		{Style["FULL\nSS KERNEL ", Bold], 
     		TableForm[Validations[[1]], TableHeadings -> tabheads]}, 
     	If[Length[Validations]==1,{},
     		{Style["FLATTENED\nSS KERNEL ", Bold], 
     		TableForm[Prepend[Validations[[2]],	{"            ","        ","          "}], 
      	  TableHeadings -> {Prepend[tabheads[[1]], " "], None}]}]
      	}, 
   		Background -> {Automatic, {LightBlue, LightYellow}}, 
   		Dividers -> {{{1}}, {{-1}, {1}}}, Alignment -> Left], 
  "\nVALIDATION TEST: Deconstruct FBA solutions (with/without artificial bounds) 
    	into the sum of a Kernel space flux, and a flux along a ray direction. 
    	Agreement between actual and reconstituted solutions are indicated by % discrepancy
    	of total flux, and angle in degrees between their directions in flux space.", Top],
       Null]
          	 }, 
        Dividers -> Center, Spacings -> 2], 
       Style["          Nothing to report          ", "Subsection"]], 
      "Output", FontSize -> 11]};
   AppendTo[pages, savepic];

   If[commandline, Export[exportfile, pages, OverwriteTarget->"KeepBoth"],
   		reportnb =  CreateDocument[pages,  
   		PrintingOptions -> {"PrintingMargins" -> 30}, PageBreakBelow -> True, Visible -> False];
   		Export[exportfile, reportnb, OverwriteTarget->"KeepBoth"];
   		NotebookClose[reportnb]]; 
    
  If[Length[results]>0,
  MainHeading = 
   "(* " <> Heading[[1]] <> " *)\n(* " <> Heading[[2]] <> " *)";
  SectionHeadings = {"\n(* Kernel Space Data Structure *)", 
    "\n(* Periphery points in flux space *)", 
    "\n(* Fixed flux numbers and values *)", 
    "\n(* Thin directions in flux space *)", 
    "\n(* Ray vectors in flux space *)", 
    "\n(* Maximal inscribed sphere radius and centre *)", 
    "\n(* Reversible reactions becoming fixed in direction *)",
    "\n(* Names of metabolites, and reactions that form the flux space basis vectors *)",
    "\n(* Reduced Solution Space Data Structure *)" 
    };
  exportfile = FileBaseName[datafile] <> " SSKernel.dif";
  exportfile = FileNameJoin[{DirectoryName[datafile], exportfile}];
  progresscounter = 2;
  progresslabel = InsertLinebreaks["Exporting data to file " <> exportfile];
  Print[Style[progresslabel, Blue, TextAlignment -> Center]];
  Export[exportfile, 
    Prepend[Riffle[InputForm /@ SectionHeadings, 
      InputForm /@ results], InputForm@MainHeading], "DIF", OverwriteTarget->"KeepBoth"], 
  Print["Interim report saved, but export data unavailable - \
first execute the Display stage before that can be saved. "]];

   progresslabel = If[success, " Export of output files completed ",
   	" One or both output files failed to be exported. "];
   progresscounter=0;
   success
  ]

SetAttributes[StageManager, HoldFirst];
StageManager[mainstage_, repwindow_] :=
 (*For each button press, this function prepares the report and \
commentary windows, stores/restores interim results, regulates button \
availability,and calls the appropriate function to execute the \
required calculations. *)
 Module[{skipstages = 0, success = True, result},
  (* Print["I have arrived in StageManager."]; *)
  (* Delete any existing plots from the report window *)
  If[mainstage != "Export" && Length[chordpic] > 0, 
  	(* Print["Removing chord plots"]; *)
   Validations={};
   SelectionMove[repwindow, Previous, CellGroup, 1];
   NotebookDelete[repwindow];
   chordpic = {}; cenpic = {}];
  If[mainstage != "Display" && mainstage != "Export" && Length@flatplot > 0, 
  	(* Print["Removing flatplot"]; *)
   SelectionMove[repwindow, Previous, CellGroup, 1];
   NotebookDelete[repwindow];
   flatplot = {}];
   
  skipstages = Switch[mainstage, "Loading", 0, "Reducing", 1, "Capping", 2, 
    "Shaping", 3, "Display", 4, "Export", 5];
  
  (* Perform functions appropriate for the current stage*)
  If[mainstage == "Loading" || (automate && skipstages < 1),
		(* New model-initialize settings,delete previous commentary and reports  *)
(* If the control window has a docked cell, everything below it is deleted.
   If not, the last cell or cellgroup is selected and tested if it contains
   any buttons. If not, it is deleted. 
    This normally removes any output, while protecting a non-docked control box. 
    If the user inserted additional cells, it may fail to delete all output. *)
	If[Length@CurrentValue[controlwindow, "DockedCells"] == 0, 
 		SelectionMove[controlwindow, After, Notebook]; 
 		SelectionMove[controlwindow, Previous, CellGroup];
 		If[! MemberQ[NotebookRead[controlwindow], ButtonBox, Infinity, 
    		Heads -> True], NotebookDelete[controlwindow]],
 		SelectionMove[controlwindow, All, Notebook];
 		NotebookDelete[controlwindow]];
 	Get[FileNameJoin[{PackagesDirectory, "Configuration.m"}]];
   		panelback = {LightGreen, LightYellow, LightYellow, LightYellow, LightYellow};
   	If[StringQ@Quiet@Check[FindFile[datafile], $Failed],
   (* Species = If[Species == "" || StringMatchQ[Species, Whitespace], 
      FileNameTake[datafile, {-2}], Species]; *)
    Print@Grid[{{"Stage 1"}}, ItemSize -> 40, Spacings -> {Automatic, 0}, 
      Frame -> True, Background -> Hue[0, 1, 1, .5]];
    {Loadtime, result} = Timing[CheckAbort[LoadModel[datafile, verbose], success = False]];
    SelectionMove[EvaluationNotebook[], After, Notebook, 1],
    success = False; MessageDialog[" Input file is invalid! "]];
    Heading = {"Loaded data for " <> Species, "from " <> FileNameTake[datafile] <>
    	 " in " <> Quiet@ToString[NumberForm[Loadtime, 3]] <> " seconds."};
    Reductiontable={{}}; (* Poke the table so report window gets updated *)
  	If[success,
    LoadReports = {Reductiontable, Kerneltable, Processreport};
    panelback[[1]] = LightCyan;
    available = {True, True, False, False, False, False};
    {Sraw,rawSvals,rawbounds,rawobject,rawFBAvec,rawreacts,rawmets,rawexternals,rawexempts} = 
     	{S,Svals,bounds,objectselector,FBAvector,ReactionNames,MetaboliteNames,externals,exemptions}; 
    rawfeasibles = feasiblepoints;
    ,
    panelback[[1]] = LightRed;
    available = {True, True, False, False, False, False};
    Return[]];
   ];
  
  If[mainstage == "Reducing" || (automate && skipstages < 2),
   panelback[[2;;]] = {LightGreen, LightYellow, LightYellow, LightYellow};
   (* NOTE: The LP solver sometimes (!) gives "not a matrix" error message for sparse matrix, so 
   			convert to the normal form to prevemnt this.*)
   {S,Svals,bounds,objectselector,FBAvector,ReactionNames,MetaboliteNames,externals,exemptions} = 
    	{Normal@Sraw,rawSvals,rawbounds,rawobject,rawFBAvec,rawreacts,rawmets,rawexternals,rawexempts} ; 
   feasiblepoints = rawfeasibles; 
   SStype = "FacetCone";
   Tolerances = StringTrim[Tolerances, (" " | PunctuationCharacter) ...];
   {Reductiontable, Kerneltable, Processreport} = LoadReports;
   Reductiontable = {{"Stage", " Constraints ", " Variables ", " Ray Yield "}};
   Print@Grid[{{"Stage 2"}}, ItemSize -> 40, Spacings -> {Automatic, 0}, 
   	Frame -> True, Background -> Hue[0.16, 1, 1, .5]];    
    {RSStime, result} = Timing[
     If[reshuffle,Shakeup[]];	
   	 result = CheckAbort[Check[DoFBA[Tolerances, True], success = False], success = False];    
   		If[StringQ[result], success = False;
     	Print[Style["LP solution of the imported FBA model has failed. 
     	Tick the reshuffle box and/or adjust the LP tolerance before retrying Stage 2.", Red]]];
   	result = CheckAbort[Check[ReducedSolutionSpace[S, bounds, objectselector, maxmin, verbose],
   		 success = False], 	 success = False];
    If[StringQ[result], success=False; 
    	Print[Style[result<>"\nCheck that artificial bound value is large enough to
    	avoid excluding proper bounds.",Red]]; 
    	PrependTo[feasiblepoints,{}]]
     ];
     
     (* Could also use FileBaseName[datafile] instead of ModelName *)
   Heading = {"Reduced Solution Space for " <> Species, 
   	"  Model " <> ModelName <> " of type " <> ToString@SStype <> 
   	"\n calculated on " <> DateString["DateShort"] <> " in " <> 
      Quiet@ToString[NumberForm[RSStime, 3]] <> " seconds."};
   SelectionMove[EvaluationNotebook[], After, Notebook, 1]; 
   available = If[success || Length@feasiblepoints > 0, {True, True, True, False, False, True}];
  	RSStype = SStype; RSSraydim = RayDim;
    RSSReports = {Reductiontable, Kerneltable, Processreport};
   If[success,
	panelback[[2]] = LightCyan,
    panelback[[2]] = LightRed; Return[]];
   ];
  
  If[mainstage == "Capping" || (automate && skipstages < 3),
   panelback[[3;;]] = {LightGreen, LightYellow, LightYellow};
   SStype = RSStype; RayDim = RSSraydim;
   {Reductiontable, Kerneltable, Processreport} = RSSReports;
   (* Force chord calculation in stage 4 if the SSK is recalculated *)
   PreflatSSK={};PreflatPeris={}; PreflatChords={}; PreflatChordLengths={}; 
   Print@Grid[{{"Stage 3"}}, ItemSize -> 40, Spacings -> {Automatic, 0}, 
   	Frame -> True,  Background -> Hue[0.32, 1, 1, .5]];
   {Kerneltime, result} = 
    Timing[CheckAbort[Check[KernelFinder[verbose],
    	 success = False],	 success = False]];
   Heading = {"Capped Solution Space for " <> Species, 
     " Model " <> ModelName <> " of uncapped type " <> ToString@SStype <>
       "\n calculated on " <> DateString["DateShort"] <> " in " <> 
     Quiet@ToString[NumberForm[Kerneltime, 3]] <> " seconds."};
   SelectionMove[EvaluationNotebook[], After, Notebook, 1];
   If[success,
   	panelback[[3]] = LightCyan;
    SSKinRSS = KernelSpace; 
    SSKinsphere = InscribedSphere;
    KernelReports = {Reductiontable, Kerneltable, Processreport};
    available = {True, True, True, True, False, True},
    panelback[[3]] = LightRed; Return[]]
   ];
  
  If[mainstage == "Shaping" || (automate && skipstages < 4),
  	panelback[[4;;]] = {LightGreen, LightYellow};
   (*Restore unflattened SSK*)
   KernelSpace = SSKinRSS;     InscribedSphere = SSKinsphere;
   {Reductiontable, Kerneltable, Processreport} = KernelReports;
   SelectionMove[repwindow, After, Notebook];
   NotebookWrite[repwindow, Cell["Flattening Decision Plot ", "Subsubsection"]];
   Print@Grid[{{"Stage 4"}}, ItemSize -> 40, Spacings -> {Automatic, 0}, 
   	Frame -> True, Background -> Hue[0.48, 1, 1, .5]];
   If[SStype != "Point",
    {Shapetime, result} = Timing[CheckAbort[Check[KernelShape[maxaspect, maxthin, verbose],
    	 success = False], 	 success = False]]; 
    Heading = {"Shape Analysis for " <> Species, 
      " Model " <> ModelName <> "  with uncapped kernel of type " <> 
      ToString@SStype <> 
       "\n calculated on " <> DateString["DateShort"] <> " in " <> 
       Quiet@ToString[NumberForm[Shapetime, 3]] <> " seconds."};
    SelectionMove[EvaluationNotebook[], After, Notebook, 1]; ,
    Shapetime = 0; 
    flatplot = Graphics@Text["Point SSK, has no shape."]];
   NotebookWrite[repwindow, ToBoxes[Show[flatplot, ImageSize -> All]]]; 
   SelectionMove[repwindow, After, Cell, 1];
   If[success,
	panelback[[4]] = LightCyan;   	
    ShapeReports = {Reductiontable, Kerneltable, Processreport}; 
    available = {True, True, True, True, True, True},
    panelback[[4]] = LightRed; Return[]]
   ];
  
  If[mainstage == "Display" || (automate && skipstages < 5),
  	panelback[[5]] = LightGreen;
   {Reductiontable, Kerneltable, Processreport} = ShapeReports;
   SelectionMove[repwindow, After, Notebook];
   NotebookWrite[repwindow, 
    Cell["Chords, diameters, centrality and aspect ratios.", "Subsubsection"]];
   Print@Grid[{{"Stage 5"}}, ItemSize -> 40, Spacings -> {Automatic, 0}, 
   	Frame -> True, Background -> Hue[0.64, 1, 1, .5]];
   {Chordtime, result} = 
    Timing[CheckAbort[Check[
    	{chordpic, cenpic} = KernelDisplay[ExportResults, verbose],
    	 success = False], 	 success = False]];
   Heading = {"Full SS Kernel Analysis for " <> Species, 
     " Model " <> ModelName <> "  with uncapped kernel of type " <> 
      ToString@SStype <> 
      "\n calculated on " <> DateString["DateShort"] <> " in " <> 
      Quiet@ToString[NumberForm[Loadtime + RSStime + Kerneltime + Shapetime + Chordtime, 3]] <> 
      " seconds."}; 
   NotebookWrite[repwindow, ToBoxes[chordpic]];
   SelectionMove[repwindow, After,Notebook]; 
   NotebookWrite[repwindow, ToBoxes[cenpic],After];
   SelectionMove[repwindow, After,Notebook]; 
   If[success, 
   	panelback[[5]] = LightCyan;
   	available = {True, True, True, True, True, True}, 
    panelback[[5]] = LightRed; Return[]]
   ];
  
  If[mainstage == "Export" (* || automate *) ,
  	panelback[[5]] = LightGreen;
   CheckAbort[success=Check[
   	SaveResults[If[Length@ExportResults==0,{},ExportResults], datafile],
   	 False],
      success = False];
   If[success,
    panelback[[5]] = LightCyan,
    panelback[[5]] = LightRed
   ]
   ];
  ]

UserInterface[controlwindow_, reportwindow_] := 
 Module[{stage1, stage2, stage3, stage4, stage5, monitor, progress, 
   Controlbox, StatusReport}, 
  stage1 = Panel[Grid[{{FileNameSetter[Dynamic[datafile, (datafile = #;
           Species = FileNameTake[datafile, {-2}]) &], 
        "Open", {"Metabolic models" -> {"*.mat", "*.sbml", "*.xml", 
           "*.m", "*.WL"}, "All files" -> {"*"}}], 
       InputField[Dynamic[datafile], String, 
        FieldHint -> "Choose the input file.", FieldSize -> 38],
        SpanFromLeft, SpanFromLeft,
        "Option reset", Checkbox[Dynamic[loadreset]]},  
        {"Organism: ", 
       InputField[Dynamic[Species], String, 
        FieldHint -> "Default: parent directory name.", 
        FieldSize -> 15], Optiontable[[1, 1]], 
       InputField[Dynamic[timeconstraint], Number, FieldSize -> 3], 
       
       Button[Style[" Load ", 10, Bold], 
        StageManager["Loading", reportwindow], ImageSize -> {80, 16}, 
        Background -> Lighter[Green, 0.95], 
        Enabled -> Dynamic@available[[1]], Method -> "Queued"],SpanFromLeft}}, 
     Spacings -> {1.7, 0}], 
    Style["Stage 1:   Load and check FBA model", "Subsubtitle", 
     FontSize -> 12], ImageSize -> {Automatic, 56}, 
    Background -> Dynamic[panelback[[1]]]];
  stage2 = 
   Panel[Grid[{{(*Optiontable[[15,1]],*)"Flux bounds larger", 
       "than this", "are taken as artificial:", 
       InputField[Dynamic[artificial], Number, 
        FieldSize -> 5]}, {Tooltip[Optiontable[[3, 1]], 
        "A single value used throughout, or a pair of values in \
braces; 
        the first is used for the initial bounded FBA only."], 
       InputField[Dynamic[Tolerances], String, FieldSize -> 5], 
       Optiontable[[4, 1]], 
       InputField[Dynamic[fixtol], Number, 
        FieldSize -> 3]}, {"Exempt unbuffered externals", 
       Checkbox[Dynamic[exempt, (FBAdone = False; exempt = #) &]], 
       "Reshuffle stoichiometry first", 
       Checkbox[
        Dynamic[reshuffle, (FBAdone = False; reshuffle = #) &]], 
       Button[Style[" Reduce ", 10, Bold], 
        StageManager["Reducing", reportwindow], ImageSize -> {80, 16},
         Background -> Lighter[Green, 0.95], 
        Enabled -> Dynamic@available[[2]], Method -> "Queued"]}}, 
     Spacings -> {2.0, 0}], 
    Style["Stage 2:   Reduction - Eliminate fixed fluxes and \
prismatic rays.", "Subsubtitle", FontSize -> 12],
    ImageSize -> {Automatic, 60}, 
    Background -> Dynamic[panelback[[2]]]];
  stage3 = 
   Panel[Grid[{{Optiontable[[5, 1]], 
       InputField[Dynamic[targetcount], Number, FieldSize -> 3], 
       Optiontable[[7, 1]], 
       InputField[Dynamic[samplesize], Number, 
        FieldSize -> 3]}, {Optiontable[[6, 1]], 
       InputField[Dynamic[treesize], Number, FieldSize -> 5], 
       Optiontable[[9, 1]], 
       InputField[Dynamic[greedyfails], Number, 
        FieldSize -> 3]}, {Optiontable[[14, 1]], 
       InputField[Dynamic[DefaultCap], Number, FieldSize -> 3], Null, 
       Null, Button[Style["  Cap  ", 10, Bold], 
        StageManager["Capping", reportwindow], ImageSize -> {80, 16}, 
        Background -> Lighter[Green, 0.95], 
        Enabled -> Dynamic@available[[3]], Method -> "Queued"]}}, 
     Spacings -> {2.4, 0}], 
    Style["Stage 3:   Ray matrix and Progenitor;   Coincidence and \
tangent capping.", "Subsubtitle", FontSize -> 12], 
    ImageSize -> {Automatic, 64}, 
    Background -> Dynamic[panelback[[3]]]];
  stage4 = 
   Panel[Grid[{{Optiontable[[10, 1]], 
       InputField[Dynamic[{chordmin, chordmax}], Expression, 
        FieldSize -> 5], Optiontable[[11, 1]], 
       InputField[Dynamic[flipmax], Number, 
        FieldSize -> 3]}, {Optiontable[[12, 1]], 
       InputField[Dynamic[maxaspect], Number, FieldSize -> 3], 
       Tooltip[Optiontable[[13, 1]], 
        "This overrides the aspect ratio; choose 0 for no flattening, \
Infinity to use aspect ratio only. "], 
       InputField[Dynamic[maxthin], Expression, FieldSize -> 3], 
       Button[Style[" Flatten ", 10, Bold], 
        StageManager["Shaping", reportwindow], ImageSize -> {80, 16}, 
        Background -> Lighter[Green, 0.95], 
        Enabled -> Dynamic@available[[4]], Method -> "Queued"]}}, 
     Spacings -> {1.3, 0}], 
    Style["Stage 4:   Analyze SS Kernel shape;   Flatten large aspect \
ratios.", "Subsubtitle", FontSize -> 12], 
    ImageSize -> {Automatic, 50}, 
    Background -> Dynamic[panelback[[4]]]];
  stage5 = 
   Panel[Grid[{{CancelButton[Style["Exit SS Kernel", 10, Bold], 
        NotebookClose[reportwindow];
        NotebookClose[controlwindow], ImageSize -> {80, 16}, 
        Background -> Red], Null, Null, Null, 
       Button[Style[" Export ", 10, Bold], 
        StageManager["Export", reportwindow], ImageSize -> {80, 16}, 
        Background -> Lighter[Green, 0.95], 
        Enabled -> Dynamic@available[[6]], Method -> "Queued"], 
       Button[Style[" Display ", 10, Bold], 
        StageManager["Display", reportwindow], ImageSize -> {80, 16}, 
        Background -> Lighter[Green, 0.95], 
        Enabled -> Dynamic@available[[5]], Method -> "Queued"]}}, 
     Spacings -> {4.6, 0}], 
    Style["Stage 5: Validation, Centering and Chord display ;    \
Export results.", "Subsubtitle", FontSize -> 12], 
    ImageSize -> {Automatic, 28}, 
    Background -> Dynamic[panelback[[5]]]];
  monitor = 
   Panel[Labeled[
     Tooltip[ProgressIndicator[Dynamic[progresscounter], 
       Dynamic[progressrange], ImageSize -> {260, Automatic}], 
      HoldForm[Dynamic[progresscounter]/Dynamic[progressrange[[2]]]]],
      Dynamic[progresslabel]], Background -> LightYellow];
  (*progress=Row[{Style["Running Commentary : verbose","Subsubtitle",
  FontSize->12],Checkbox[Dynamic[verbose]],
  "                                                 ",Style[
  "Automate stages","Subsubtitle",FontSize->12],Checkbox[Dynamic[
  automate]]},Spacer[15]];
  Controlbox=Panel[Column[{stage1,stage2,stage3,stage4,stage5,monitor,
  progress},Spacer[20]],Background->LightGray(*,
  ImageSize\[Rule]Full*)];*)
  Controlbox = 
   Panel[Grid[{{stage1, Column[
       	{Panel[Checkbox[Dynamic[automate]], "Automate", Appearance -> "Frameless"], 
         Panel[Checkbox[Dynamic[verbose]], "Commentary",Appearance -> "Frameless"]}]}, 
       {stage2, Rotate[monitor, Pi/2]}, {stage3, SpanFromAbove}, 
       {stage4, SpanFromAbove}, {stage5, SpanFromAbove}}, Alignment -> Top, Spacings -> {1, 0.3}],
         Background -> LightGray, FrameMargins -> 1];
  StatusReport = 
   ExpressionCell[
    Dynamic[Module[{combinedreport, table}, 
      table = Join[Reductiontable, Kerneltable];
      combinedreport = 
       If[Length[Processreport] > 0 || Length[table] > 0, 
        Column[{Style[Heading[[1]], "Subtitle"], 
          Style[Heading[[2]], "Subsubtitle"], 
          Row[{TableForm[
             Optiontable[[;; Ceiling[Length[Optiontable]/2]]], 
             TableSpacing -> {1, 3}], 
            TableForm[Optiontable[[-Floor[Length[Optiontable]/2] ;;]],
              TableSpacing -> {1, 3}]}, "      "], 
          If[Length[table] > 1, 
           TableForm[table[[2 ;;, 2 ;;]], 
            TableHeadings -> {table[[2 ;;, 1]], table[[1, 2 ;;]]}], 
           Null], TableForm[Flatten@Processreport], 
          If[Length[Validations] > 0, 
           Labeled[Grid[{{Style["FULL\nSS KERNEL ", Bold], 
               TableForm[Validations[[1]], 
                TableHeadings -> tabheads]}, 
              If[Length[Validations] == 
                1, {}, {Style["FLATTENED\nSS KERNEL ", Bold], 
                TableForm[
                 Prepend[Validations[[2]], {"            ", 
                   "        ", "          "}], 
                 TableHeadings -> {Prepend[tabheads[[1]], " "], 
                   None}]}]}, 
             Background -> {Automatic, {LightBlue, LightYellow}}, 
             Dividers -> {{{1}}, {{-1}, {1}}}, Alignment -> Left], 
            "\nVALIDATION TEST: Deconstruct FBA solutions \
(with/without artificial bounds) 
                	into the sum of a kernel flux, and a flux along a \
ray direction. 
                	Agreement between actual and reconstituted solutions \
are indicated by % discrepancy
                	of total flux, and angle in degrees between their \
directions in flux space.", Top], Null]}, Dividers -> Center, 
         Spacings -> 2], 
        Style["          Nothing to report          ", 
         "Subsection"]]]], "Output", FontSize -> 10];
  {Controlbox, StatusReport}]
  
  
End[] (* End Private Context *)

EndPackage[]
```
