## Supplementary material for "The FBA Solution Space Kernel – Introduction and Illustrative Examples": SSK software manual and code: RayFinding.m

```
(* ::Package:: *)

(* COPYRIGHT
					© Copyright 2022 Wynand Verwoerd

This file is part of SSKernel.

The SSKernel program is free software; you can redistribute it and/or modify it under the terms of the GNU General Public License as published by the Free Software Foundation; either version 3 of the License, or (at your option) any later version.
 
This program is distributed in the hope that it will be useful, but WITHOUT ANY WARRANTY; without even the implied warranty of MERCHANTABILITY or FITNESS FOR A PARTICULAR PURPOSE . See the GNU General Public License for more details.
 
You should have received a copy of the GNU General Public License along with this program . If not, see http://www.gnu.org/licenses/
*)

(* Wolfram Language Package *)

BeginPackage["RayFinding`",{"Configuration`"}]
(* Exported symbols added here with SymbolName::usage *)  
PrismDrop::usage="Reduces solution space by removing all prismatic rays."
SingletonRays::usage="Find all singleton rays by pseudoinverse of the constraint matrix."
CapRayFinder::usage="Find a minimal capping ray set by LP maximisation of the sum of slacks."
CompleteRayBasis::usage="Calculates a set of rays to form a (non-convex) complete basis for RS"

Begin["`Private`"] (* Begin Private Context *) 
 
SetAttributes[PrismDrop, HoldFirst];
PrismDrop[solutionspace_, PrintResult_:False] :=
 (* This routine directly condenses the constraint equations, objective space basis and ray vectors 
 by removal of prismatic rays as well as any linealities (double rays). 
 It returns the removed rays as vectors in the variable part of flux space *)
 Module[{constraints, values, OSBase, origin,cons, vars, overlaps, linrays,
 	prismrays, prismraysat, raycount, keepcon, centershift, newbase, tol = 10.^-6},
  {{constraints,values},{origin,OSBase}}=solutionspace;
  {cons, vars} = Dimensions[constraints];
  progresslabel="Eliminate prismatic rays and linealities.";
  progresscounter=0; progressrange={0,1};
  Print[Style[progresslabel, Blue, TextAlignment -> Center]];
  (* STAGE 1: Find and eliminate prismatic rays. *)
  
  overlaps = constraints.Transpose[constraints];
  (* Find mutually orthogonal constraints, i.e. overlap
  row vectors that are zero except for a diagonal 1*)
  prismraysat = 
   Reap[Do[If[Max@Abs[overlaps[[i]] - UnitVector[cons, i]] < tol, 
       Sow[i]], {i, cons}]][[2]]; 
  raycount=If[prismraysat=={}, 0, prismraysat =First@prismraysat; Length[prismraysat]];      
 (* Print[ToString[raycount]<> " prismatic rays for "<>ToString[vars]<>" variables."]; *) 
  If[raycount==vars,
  	(*Print["All remaining degrees of freedom are rays, so the SSK reduces to a single point. "];*)
  	 RayDim=raycount;
  	 AppendTo[Reductiontable, {"Point SSK after removing prismatic rays", 0, 0, raycount}];
     solutionspace={{{{1.0}},{0.}},{origin,{ConstantArray[0.,Length[origin]]}}};
     Return[-Chop[constraints.OSBase]]];
     
  If[raycount > 0,
   prismrays = -constraints[[prismraysat]];
   keepcon = Complement[Range[cons], prismraysat];
   (* Print[keepcon]; *)
   (* Move the OS origin to its new position in FS *)
   centershift = values[[prismraysat]].constraints[[prismraysat]];
   origin = Chop[origin + centershift.OSBase];
   (* Get the base of the lower dimensional subspace remaining after prismatic rays are eliminated *)
   newbase = 
    N@Orthogonalize[NullSpace[prismrays], Method -> "Householder",Tolerance->tol];
   (* Keep only the constraints that do not oppose prismatic rays. 
   Since these vectors are by construction orthogonal to the prismatic projection, 
   they retain their normalisation and can be simply projected, 
   and their constraint values also remain unaffected. *)
   constraints = constraints[[keepcon]];
   values = values[[keepcon]];
   If[Length[keepcon]>0, 
   	constraints =constraints.Transpose[newbase];
   	{cons, vars} = Dimensions[constraints],
   	
   	constraints ={ConstantArray[0.,Length[newbase]]}; values={0};
   	{cons,vars}={0,Length[newbase]}
   	];
   (* Finally, express prismatic rays as well as the basis 
   of the reduced SS in terms of the embedding space*)
   prismrays = Chop[prismrays.OSBase];
   OSBase = Chop[newbase.OSBase];
   
   AppendTo[Reductiontable, {"Remove prismatic rays", cons, vars, raycount}];
  
   If[PrintResult, Print["Eliminating " <> ToString[raycount] <> 
     " prismatic rays reduces the SS specification to " <> 
     ToString[cons] <> " constraints on " <> ToString[vars] <> 
     " variables."]];
     
     , prismrays={}];
       
     
  (* STAGE 2: Find and eliminate linealities. *)
  
  linrays=NullSpace[constraints, Tolerance -> tol];
  raycount = Length[linrays];
	(* Print[{"Number of prismatic rays and linealities ",Length@prismrays,Length@linrays}]; *)  
  (* Note that by construction prismrays are mutually orthogonal, as are linrays, so
   define the dimensions of their respective spaces *)
  RayDim=Length[prismrays]+raycount;
  If[PrintResult, Print[ToString[raycount]<> " lineality directions were found for "<>ToString[vars]<>" variables."]]; 
  If[raycount==vars,
  	(*Print["All remaining degrees of freedom are rays, so the SSK reduces to a single point. "];*)
  	 AppendTo[Reductiontable, {"Point SSK after removing lineality rays", 0, 0, 2*raycount}];
     solutionspace={{{{1.0}},{0.}},{origin,{ConstantArray[0.,Length[origin]]}}};
     linrays = Chop[linrays.OSBase];
   	 prismrays=Join[prismrays,linrays,-linrays];
     Return[prismrays]];
     
  If[Length[linrays]>0,
  	   (* Get the base of the lower dimensional subspace remaining after linealities are eliminated *)
   newbase = 
    N@Orthogonalize[NullSpace[linrays], Method -> "Householder",Tolerance->tol];
    constraints = constraints.Transpose[newbase];
      (* Add opposing rays for each lineality to the ray set and rexpress in terms of the embedding space*)
   linrays = Chop[linrays.OSBase];
   prismrays=Join[prismrays,linrays,-linrays];
   OSBase = Chop[newbase.OSBase];
   {cons, vars} = Dimensions[constraints];
   AppendTo[Reductiontable, {"Linealities yielding ray pairs", cons, vars, 2*Length[linrays]}];
  ];
     
   solutionspace={{constraints,values},{origin,OSBase}};
   prismrays
  ]
  
  
SingletonRays[constraints_] := 
 Module[{cons, vars, SValues, SConstraints, SRays, normray, 
   slack, SingleRays, tol = 10.^-6},
  {cons, vars} = Dimensions[constraints];
  SValues = ConstantArray[0, cons];
  SConstraints = 
   Join[Transpose[{ConstantArray[0., cons]}], constraints, 2];
  SingleRays = Reap[Do[
      SRays = NullSpace[ReplacePart[SConstraints, {c, 1} -> 1.],Tolerance->tol];
      If[Length[SRays] > 0,
       Map[(normray = #/Norm[#[[-vars ;;]]]; 
       	    slack = First@normray; 
            Sow[Sign[slack]*Join[ReplacePart[SValues, c -> First@normray], Rest[normray]]])&,
          SRays]],
    {c, cons}]][[2]];
   If[Length[SingleRays] > 0, 
   	Chop@DeleteDuplicates[First@SingleRays,Abs[#1.#2 - 1] < 0.00001 &],
   	{}]
  ]

CapRayFinder[constraints_] := 
 Module[{cons, vars, Rays = {}, SConstraints, scons, SValues, SXRange,
    SObjective, SX, tol = 0.001, count = 0},
  {cons, vars} = Dimensions[constraints];
  If[vars == 0, Return[{}]];
  SConstraints = Join[IdentityMatrix[cons], constraints, 2];
  SConstraints = SparseArray@SConstraints;
  scons = cons;
  (* Each iteration produces a new capping ray. 
  A maximum number of iterations needs to be set to avoid an infinite loop if LP solver fails. 
  But even if it does not fail it can stall and produce no result.
  So for a purpose like escaping from a corner in centering, a single capping ray should
  be enough and so set it to find a single ray for that purpose. *)
  While[count++ < 2,
   SObjective = 
    Join[ConstantArray[1., scons], ConstantArray[0., vars]];
   (*SObjective=Join[ob,ConstantArray[0.,vars+scons-cons]];*)
   SValues = ConstantArray[{0., 0}, scons];
   SXRange = 
    Join[ConstantArray[{0., Infinity}, scons], 
     ConstantArray[{-1, 1}, vars]];
   (* Print[count];
   Print[Dimensions/@{SObjective,SConstraints,SValues,SXRange}]; *)
   (* Simplex works best; 
   Interior Point and RevisedSimplex sometimes fail to converge *)
   SX = Quiet[
     Check[LinearProgramming[-SObjective, SConstraints, SValues, 
       SXRange, Method -> "Simplex", Tolerance -> tol], 
      "No Solution", {LinearProgramming::lpsnf}],{LinearProgramming::lpsnf}];
  (* If[VectorQ[SX],Print[Length@SX],Print@SX;Break[]]; *)
   If[Norm[SX] < tol || SX == "No Solution", Break[]];
   (*Print[SObjective.SX];*)
   SX = SX/Norm[SX[[-vars ;;]]];
   (* Return the full SX vector, but skip any slacks for temporary constraints  *)
   (*Rays=Append[Rays,Join[SX[[;;cons]],SX[[-vars;;]]]];*)
   (* Return only the actual ray vectors   *)
   Rays = Append[Rays, SX[[-vars ;;]]];
   (* Add all the found composite ray capping planes as temporary constraints and repeat *)
   scons++;
   SConstraints = 
    Join[IdentityMatrix[scons], 
     Join[constraints, Rays[[All, -vars ;;]]], 2];
   (*Print[Length/@SConstraints]*)
   ];
  If[Rays == {}, 
   Print[" CapRayFinder failed to find a ray - The constraint set may define a closed polytope."]];
  
  Chop@Rays]
  
SetAttributes[ZeroSlackTester, HoldFirst];
ZeroSlackTester[RayMat_, constraints_, facet_, inherit_, zerocols_: {}, printresult_: False] :=
 (* This routine is given two sets of constraint numbers with known \
zero slack values:
 
 facet: The set of constraints that characterise the facet under \
consideration. All rays in the facet are orthogonal to these \
constraint vectors, so the constraint equation reduces to equality \
and its slack is zero.
 
 inherit: The ray-free space of the lower level facet in which the \
current one is embedded. Higher level facets inherit RF constraints \
from lower levels, so zero slack values for these are also given.
 
 If non-empty, no slack variable at all is allocated to members of \
facet and inherit (noslacks), equivalent to them being set to zero.
 
 Optionally, it can be given a candidate set (zerocols) of slack \
columns
  that are ostensibly zero, to test. If absent or empty, it will \
itself set up a list that were consistently zero in all rays found so \
far that belong to the facet.
 Then it probes all candidates that do not belong to noslacks,by LP \
testing. 
 If a counterexample is found, it is added to the ray list 
 and the probe columns correspondingly reduced. 
 When the LP test fails, any columns that are left are thus confirmed
 as zero and are returned.
 
    *)
 Block[{cons, vars, noslacks, facetrays, conslack, slackcount, 
   slackcon, probecols, probecount, maxprobe, probe, confirmed = {}, SObjective,
    SXConstraints, SXRange, SValues, SX, fullSX, newrays, candidates, 
   zeroslacks, zerocons, lpcount = 0, tol = 0.00001},
   (* Tol value 0.00001 and LPmethod -> Simplex used previously. *)
  {cons, vars} = Dimensions[constraints];
  (* Slack columns of preassigned zero constraints will come out zero so skip them. 
  Note that "probecols" are numbers of columns in the full S-matrix, 
  not in the one reduced by leaving out "noslacks"! *)
  noslacks = Union[facet, inherit];
  candidates = If[zerocols != {}, zerocols,
    (* Look for zero columns only in known rays that belong to the \
facet, otherwise just the PRF will be found *)
    facetrays = 
     Flatten@Position[RayMat[[All, facet]], _?(Total[#] == 0 &), {1}, 
       Heads -> False];
    If[facetrays == {}, Range[cons], 
     Flatten@Position[
       Total[RayMat[[facetrays, ;; cons]]], _?(# < 0.00001 &)]]];
  candidates = probecols = Complement[candidates, noslacks]; 
  
  If[printresult, 
   Print["For facet " <> ToString[facet] <>  " a total of "<> 
     ToString[Length@probecols] <>
     " columns qualify for zero slack probing. "]];  
     
  maxprobe= probecount = Length[probecols];
  (* No probe columns means that counterexamples are known for all columns. 
  So the facet does not add any RF constraints to those known from the lower level. *)
  If[probecount == 0, Return[{}]];
  (* To keep track of which constraints have free slacks, use two arrays: 
  conslack is a list of all constraints that have an allocated slack variable,
  i.e. conslack[[slack]] is the constraint for given slack variable nr "slack";
  slackcon is a kind of inverse of this, i.e. slackcon[[con]] is the slack var 
  for given constraint nr "con", or 0 if none *)
  conslack = Complement[Range[cons], noslacks];
  slackcount = Length[conslack];
  slackcon = Range[slackcount];
  If[Length[noslacks] > 0, 
   slackcon = 
    Insert[slackcon, 0, 
     Partition[noslacks - Range[Length[noslacks]] + 1, 1]]];
  (*
  Print["The following constraints have slack variables: "<>ToString[conslack]];
  Print["The slack that belongs to each constraint is: "<>ToString[slackcon]];
  *)
  SValues = ConstantArray[{0., 0}, cons];
  SXConstraints = 
   Join[IdentityMatrix[cons][[All, conslack]], constraints, 2];
  SObjective = 
   Join[ConstantArray[1, slackcount], ConstantArray[0, vars]];
  (* Set range constraints for MINIMISATION .
    Limit the range of flux components to avoid  extreme ray lengths *)
  SXRange = 
   Join[ConstantArray[{0., Infinity}, slackcount], 
    ConstantArray[{-1000., 1000.}, vars]];
  (* Minimisation requires removing the trivial solution by fixing \
one slack to a finite value,say 0.5 . This value is arbitrary, 
  as long as it is significantly nonzero, 
  because scaling of the SX vector is fixed later anyway. 
  Add a dummy constraint to fix this *)
  SXConstraints = 
   Append[SXConstraints, ConstantArray[0, slackcount + vars]];
  SValues = Append[SValues, {0.5, 1}];
  newrays = Reap[
     While[probecount > 0, lpcount++;
      probe = slackcon[[First@probecols]];
 (*     If[printresult, PrintTemporary["ZeroSlackTester probing column "<>
    	ToString[probe]<>" in LP round "<>ToString[lpcount]]];      *)
      SXConstraints[[-1]] = UnitVector[slackcount + vars, probe];
      (*Print[{Dimensions[SObjective],Dimensions[SXConstraints],
      Dimensions[SValues],Dimensions[SXRange]}];*)
      SX = 
       Quiet[Check[
         LinearProgramming[SObjective, SXConstraints, SValues, 
          SXRange, Method -> "Simplex", Tolerance -> tol], 
         "No Solution", {LinearProgramming::lpsnf}], {LinearProgramming::lpsnf}];
      (* Since LP above searched for a ray (with probe column slack nonzero at 0.5) the norm 
      of the vector found is essentially unlimited. In practice that does not happen 
      with Simplex, since it only tries vertices and even the unbounded facets only 
      have vertices with moderate flux values. But InteriorPoint is not restricted 
      to vertices, so can produce huge ray vetors. When this is normalised below, it
      can reduce the probe slack to less than tol. In this case the probe col will 
      in effect have zero slack so does not solve the tested condition.
      This "solution" has to be disqualified otherwise an infinite loop results where 
      the same probe col is probed repeatedly.
       *)
     (* Print[{"Checking slack total ",Total[SX[[;;slackcount]]],1./LPtol,Norm[SX[[-vars ;;]]]}]; *)
      If[VectorQ[SX] && Norm[SX[[-vars ;;]]] > 1./LPtol, SX="No Solution"]; 

      If[VectorQ[SX],
       (* Normalize the rays to avoid problems with rank calculation ? *)
     (* Print[{"Probe value and normalisation ",SX[[probe]],Norm[SX[[-vars ;;]]]}]; *)
       SX = SX/Norm[SX[[-vars ;;]]];
       (* Reintroduce the skipped "noslacks" zero slack values *)
       fullSX = Join[ConstantArray[0, cons], SX[[slackcount + 1 ;;]]];
       fullSX[[conslack]] = SX[[;; slackcount]];
       Sow@Chop[fullSX, LPtol];
       progresscounter= ++RayCount;
       (* Keep track of columns that are zero/nonzero for all rays so far found *)
       zeroslacks = 
        Flatten@Position[SX[[;; slackcount]], _?(# < LPtol &)];
     (* Print[" Zeroes found for slacks no "<>ToString[zeroslacks]]; *)
       zerocons=conslack[[zeroslacks]];
     (*  Print["In probing round "<>ToString[lpcount]<>" zeroes are found for constraints no "<>
       	ToString[zerocons]]; *)
       probecols = Intersection[probecols, zerocons]; 
       probecount = Length[probecols];
     (*  Print[ToString[probecount]<>" Zero columns to be further tested "<>ToString[probecols]]; *)
       ,
       AppendTo[confirmed, conslack[[probe]]];
       probecols = Rest[probecols]; probecount--;
       ];
       (* If[printresult, Pause[0.1];NotebookDelete@temp]; *)
      ]
     ][[2]];
  
  If[Length[newrays] > 0,
   newrays = newrays[[1]];
   RayMat = Join[RayMat, newrays]];
  If[printresult, probecount = Length[confirmed]; 
   Print[" ZeroSlackTester required " <> ToString[lpcount] <> 
     " LP rounds to " <>
     Which[probecount == 0, "eliminate all probed constraints", 
      probecount == Length[candidates], 
      "confirm without change the probe columns ", True, 
      "reduce the probed constraints to " <> ToString[probecount] <> " confirmed members."]];
   ];
  confirmed
  ]
    
  CompleteRayBasis[constraints_, printresult_: False] := 
 Block[{cons, vars, RayMat = {}, basedim, raydim, PRF, targets, 
   targetcols, augmentedcons, augmentedrays, newrays, tol = 10.^-6}, 
   {cons, vars} = Dimensions[constraints];
  progresslabel = "Calculating the ray matrix. ";
  Print[Style[progresslabel, Blue, TextAlignment -> Center]];
  progressrange={0,vars};
  progresslabel=" Get singleton rays using pseudoinverse ";
  progresscounter=RayCount=0;
 
  (*Use linear algebra for a quicker and exhaustive calculation of all singleton rays*)
  RayMat = SingletonRays[constraints];
  progresscounter=RayCount=Length@RayMat;
  If[printresult, Print[ToString[RayCount] <> 
     " singleton rays are used as the starting set."]];
  
  (*Find the polytope rayfree subset and the ray space dimensions*)
  progresslabel=" Rays resulting from zero column probes ";
  PRF = ZeroSlackTester[RayMat, constraints, {}, {}, {}, printresult];
  If[Length[RayMat] == 0, raydim=0; basedim=0;
   If[printresult,  Print["Specified constraints define a closed polytope with no rays."]];
  ,
   
  raydim = If[Length[PRF] == 0, vars, 
    vars - MatrixRank[constraints[[PRF, All]], Tolerance -> tol]];
  basedim = MatrixRank[RayMat[[All, -vars ;;]], Tolerance -> tol];
  If[printresult, 
   Print["After rayfree determination, there are " <> 
     ToString[Length[RayMat]] <> " rays that span " <> ToString[basedim] <> 
     " dimensions out of the rayspace total of " <> ToString[raydim]];
   Print[" The polytope rayfree constraints are " <> ToString[PRF]]];
  
  progresslabel=" Find rays from base vector alignment   ";
  While[basedim<raydim,
  targets = NullSpace[Join[constraints[[PRF, All]], RayMat[[All, -vars ;;]]], 
    Tolerance -> tol];
  augmentedcons = Join[constraints, targets];
  augmentedrays = Join[RayMat[[All, ;; cons]], 
    ConstantArray[0., {Length@RayMat, Length@targets}], RayMat[[All, -vars ;;]], 2];
  targetcols = Range[cons + 1, cons + Length@targets];
  newrays = 
   AlignedRayFinder[augmentedrays, augmentedcons, {}, PRF, targetcols, printresult];
  If[Length@newrays != Length@targets, 
   Print[ToString[Length@targets]<>" rays required to complete the ray basis, but only "<>
   	ToString[Length@newrays]<>" were found, try another iteration."]];
  newrays = Join[newrays[[All, ;; cons]], newrays[[All, -vars ;;]], 2];
  RayMat = Join[RayMat, newrays];
  basedim = MatrixRank[RayMat[[All, -vars ;;]], Tolerance -> tol]];
  ];
  progresslabel=" Ray matrix determination completed. ";
  progresscounter=0;

  
  If[printresult, 
   Print["The final ray matrix has " <> ToString[Length[RayMat]] <> " rays that span " <> 
   	ToString[basedim] <> " dimensions out of the ray space total of " <> 
      ToString[raydim]]];
  {RayMat, raydim}]

AlignedRayFinder[RayMat_, constraints_, facet_, inherit_, targetcols_: {}, printresult_: False] :=
(*This routine constructs rays that are aligned with (i.e., have non-zero overlap with}
 the constraint vectors specified by argument targetcols.
 If argument facet is not {}, only rays falling in the specified facet are found.
 If argument inherit is not {}, the rays are also excluded from the subspace spanned by 
 the listed constraint vectors, usually the RF list of the parent of the facet.
 The new rays that were found is returned as a ray matrix.  *)
 
 Block[{cons, vars, noslacks, facetrays, conslack, slackcount, 
   slackcon, probecols, probecount, maxprobe, probe, rejected = {}, 
   SObjective, SXConstraints, SXRange, SValues, SX, fullSX, newrays, 
   zeroslacks, zerocons, lpcount = 0}, 
   {cons, vars} = Dimensions[constraints];
  (*Slack columns of preassigned zero constraints will come out zero so skip them.
  Note that "probecols" are numbers of columns in the full S-matrix,
  not in the one reduced by leaving out "noslacks"!*)
  noslacks = Union[facet, inherit];
  probecols = Complement[targetcols, noslacks];
  (*If[printresult,Print["For facet "<>ToString[facet]<>
  ",\naligned rays are to be found for columns "<>ToString[probecols]]];*)
  maxprobe = probecount = Length[probecols];
  If[probecount == 0, Return[{}]];
  (*To keep track of which constraints have free slacks, use two arrays:
  conslack is a list of all constraints that have an allocated slack variable,
  i.e.conslack[[slack]] is the constraint for given slack variable nr "slack";
  slackcon is a kind of inverse of this,
  i.e.slackcon[[con]] is the slack var for given constraint nr "con", or 0 if none*)
  conslack = Complement[Range[cons], noslacks];
  slackcount = Length[conslack];
  slackcon = Range[slackcount];
  If[Length[noslacks] > 0, 
   slackcon = Insert[slackcon, 0, 
     Partition[noslacks - Range[Length[noslacks]] + 1, 1]]];
  (*Print["The following constraints have slack variables: "<>ToString[conslack]];
  Print["The slack that belongs to each constraint is: "<>ToString[slackcon]];*)
  SValues = ConstantArray[{0., 0}, cons];
  SXConstraints = Join[IdentityMatrix[cons][[All, conslack]], constraints, 2];
  SObjective =    Join[ConstantArray[1, slackcount], ConstantArray[0, vars]];
  (*Set range constraints for MINIMISATION. 
    Limit the flux component range to avoid extreme ray lengths. *)
  SXRange = Join[ConstantArray[{0., Infinity}, slackcount], 
    ConstantArray[{-1000., 1000.}, vars]];
  (*Minimisation requires removing the trivial solution by fixing one \
slack to a finite value,say 0.5.This value is arbitrary,
  as long as it is significantly nonzero,
  because scaling of the SX vector is fixed later anyway.Add a dummy \
constraint to fix this*)
  SXConstraints = 
   Append[SXConstraints, ConstantArray[0, slackcount + vars]];
  SValues = Append[SValues, {0.5, 1}];
  newrays = Reap[
  	While[probecount > 0, lpcount++;
        probe = slackcon[[First@probecols]];
        (*Print["AlignedRayFinder probing column "<>ToString[probe]<>
        " in round "<>ToString[lpcount]];*)
        SXConstraints[[-1]] = UnitVector[slackcount + vars, probe];
        (*Print[{Dimensions[SObjective],Dimensions[SXConstraints],
        Dimensions[SValues],Dimensions[SXRange]}];*)
        SX = Quiet[
          Check[LinearProgramming[SObjective, SXConstraints, SValues, 
            SXRange, Method -> "Simplex", Tolerance -> LPtol], 
           "No Solution", {LinearProgramming::lpsnf}], {LinearProgramming::lpsnf}];
        (*Since LP above searched for a ray (with probe column slack \
nonzero at 0.5) the norm of the vector found is essentially unlimited.
In practice that does not happen with Simplex, since it only tries vertices and 
even the unbounded facets only have vertices with moderate flux values.
But InteriorPoint is not restricted to vertices, so can produce huge ray vetors.
When this is normalised below, it can reduce the probe slack to less than tol.
In this case the probe col will in effect have zero slack so does not solve the tested condition.
This "solution" has to be disqualified otherwise an infinite loop results where 
the same probe col is probed repeatedly.*)
(*Print[{"Checking slack total ",Total[SX[[;;slackcount]]],1./LPtol,Norm[SX[[-vars;;]]]}];*)
        If[VectorQ[SX] && Norm[SX[[-vars ;;]]] > 1./LPtol, SX = "No Solution"];
        If[VectorQ[SX],
    (*Normalize the rays to avoid problems with rank calculation?*)
    (*Print[{"Probe value and normalisation ",SX[[probe]], Norm[SX[[-vars;;]]]}];*)
    SX = SX/Norm[SX[[-vars ;;]]];
         (*Reintroduce the skipped "noslacks" zero slack values*)
    fullSX = Join[ConstantArray[0, cons], SX[[slackcount + 1 ;;]]];
    fullSX[[conslack]] = SX[[;; slackcount]];
    Sow@Chop[fullSX, LPtol];
    progresscounter=++RayCount;
         ,
    AppendTo[rejected, conslack[[probe]]]];
        probecols = Rest[probecols]; probecount--;
        ]][[2]];
   
  If[Length[newrays] > 0, newrays = newrays[[1]]];
  If[printresult,
   Print[" AlignedRayFinder required " <> ToString[lpcount] <> 
     " LP rounds to find " <>
     ToString[Length@newrays] <> 
     " rays aligned to constraint vectors out of the requested total of " <> ToString[Length@targetcols] ]];
  newrays]
  
 
End[] (* End Private Context *)

EndPackage[]
```
