## Supplementary material for "The FBA Solution Space Kernel – Introduction and Illustrative Examples": SSK software manual and code: RedundancyRemoval.m

```
(* Wolfram Language Package *)

(* COPYRIGHT
					© Copyright 2022 Wynand Verwoerd

This file is part of SSKernel.

The SSKernel program is free software; you can redistribute it and/or modify it under the terms of the GNU General Public License as published by the Free Software Foundation; either version 3 of the License, or (at your option) any later version.
 
This program is distributed in the hope that it will be useful, but WITHOUT ANY WARRANTY; without even the implied warranty of MERCHANTABILITY or FITNESS FOR A PARTICULAR PURPOSE . See the GNU General Public License for more details.
 
You should have received a copy of the GNU General Public License along with this program . If not, see http://www.gnu.org/licenses/
*)

BeginPackage["RedundancyRemoval`",{"Configuration`"}]
(* Exported symbols added here with SymbolName::usage *)  
DropColinears::usage="Remove redundant colinear constraint vectors by inspection of overlap matrix "
RedundancyTrimmer::usage="Full elimination of redundant constraints by LP testing."

Begin["`Private`"] (* Begin Private Context *) 

DropColinears[constraints_, values_] :=
(* Filter out constraints that are redundant due to colinearity. *)
 Module[{concount,  keepers, overlaps, colipairs, 
   colinearows, colinears, minat, discard}, 

   If [Max[Abs[1 - Norm/@constraints]] > 10^-6, 
	 Print["Invalid input to DropColinears - constraint vectors not normalised."]; 
 	 Return[{"DropColinears failed!"}]];
   concount = Length[values];
   keepers = Range[concount];
  overlaps = constraints.Transpose[constraints]; 
  overlaps = MapIndexed[Drop[#1, First[#2] - 1] &, overlaps];(* 
  Only use upper triangle *)
  colipairs = Position[overlaps, _?(# > 0.999 &)]; 
  colinearows = GatherBy[colipairs, First][[All, All, 2]];
  Do[If[keepers[[i]] > 0,
    colinears = colinearows[[i]] + i - 1; (* 
    Restore counting for full matrix *)
    (*Print[colinears];*)
    If[Length[colinears] > 1, 
     minat = Position[values[[colinears]], Min[values[[colinears]]]][[
       1, 1]]; discard = Drop[colinears, {minat}];
     keepers[[discard]] = 0]], {i, concount}];
  keepers = Select[keepers, # > 0 &];
  {constraints[[keepers]], values[[keepers]], keepers}
  ]

RedundancyTrimmer[constraints_, values_, excluded_: {}, printresults_: False] := 
 Module[{cons, vars, testset, keepers, delta = 0.01, Objective, 
   LPvalues, XRange, NewCons, NewVals, ActiveCons, ActiveVals, con, Test, redundancies},
  (* NB: perform centering before calling this function. *)
  (* Parameter delta is the outwards displacement of a constraint \
plane being tested, to make sure that tangent planes are detected.
  It needs to be large enough to deal with constraint planes not \
exactly coinciding due to numerical inaccuracy, 
  as well as to get rid of a constraint that has a genuine but \
trivially small intersection with the polytope. 
  But note that that creates a risk for centering producing an \
infeasible point. *)

 (* Reduce the number of constraints to be subjected to full LP testing, by
 	eliminating any that are colinear *)
 	
  {NewCons, NewVals, keepers} = DropColinears[constraints, values];
  redundancies = Complement[Range[Length@keepers], keepers];
  progresscounter=Length@redundancies;
  If[printresults, Print["Constraints detected as colinear and removed: " <> 
  	ToString[redundancies]]];

  {cons, vars} = Dimensions[NewCons];
  (* Use a trivial objective since only feasibility is tested. Simplex is fastest *)
  Objective = ConstantArray[0., vars];
  testset = Complement[Range[cons], excluded];
  keepers = Range[cons];
  LPvalues = Map[{#, -1} &, NewVals];
  XRange = ConstantArray[{-Infinity, Infinity}, vars];

  redundancies = Flatten@Reap[
  	Do[progresscounter++;
       ActiveCons = NewCons[[keepers]];
       (* Set the constraint to equals,and displace outwards by delta. 
       Note that for this to work, the coordinate center has to be inside the polytope, 
       otherwise positive delta produces an inwards displaceent *)
       con = testset[[k]];
       LPvalues[[con]] = LPvalues[[con]] + {delta, 1};
       ActiveVals = LPvalues[[keepers]];
       Test = 
        Quiet[Check[
          LinearProgramming[Objective, ActiveCons, ActiveVals, XRange,
            Method -> "Simplex", Tolerance -> 0.1 delta], 
          "No Solution", {LinearProgramming::lpsnf,LinearProgramming::lpdinf}],
           {LinearProgramming::lpsnf,LinearProgramming::lpdinf}];
       (* restore the constraint *)
       (*Print["Constraint no "<>ToString[testset[[k]]]<>" gives result "<>ToString[Test]];*)
       LPvalues[[con]] = LPvalues[[con]] - {delta, 1};
       If[! ArrayQ[Test], Sow[con]; 
        keepers = DeleteCases[keepers, con];(*Print[keepers];*)
        ];
       , {k, Length[testset]}]][[2]];
  If[printresults,
   If[redundancies == {}, Print["No redundancies were detected."], 
    Print["The following constraints are redundant and are removed: " <> ToString[redundancies]]
    ]];
  {NewCons[[keepers]],NewVals[[keepers]],redundancies}]

End[] (* End Private Context *)

EndPackage[]
```
