## Supplementary material for "The FBA Solution Space Kernel – Introduction and Illustrative Examples": SSK software manual and code: Resampling.m

```
(* Wolfram Language Package *)

(* COPYRIGHT
					© Copyright 2023 Wynand Verwoerd

This file is part of SSKernel.

The SSKernel program is free software; you can redistribute it and/or modify it under the terms of the GNU General Public License as published by the Free Software Foundation; either version 3 of the License, or (at your option) any later version.
 
This program is distributed in the hope that it will be useful, but WITHOUT ANY WARRANTY; without even the implied warranty of MERCHANTABILITY or FITNESS FOR A PARTICULAR PURPOSE . See the GNU General Public License for more details.
 
You should have received a copy of the GNU General Public License along with this program . If not, see http://www.gnu.org/licenses/
*)

BeginPackage["Resampling`", { "Configuration`", "Centering`","ModelProcessing`"}]
(* Exported symbols added here with SymbolName::usage *)  

TargetInsert::usage = "Identifies and/or inserts a target flux to source/sink a metabolite of interest. "
HyperSampler::usage = "Uses existing SSK periphery points to generate sample points located on a particular hyperplane. "
TargetVariation::usage="Function to display the variation of the target flux on a series of hyperplanes. "
Formula::usage="Function to display a reaction by number or name "

Begin["`Private`"] (* Begin Private Context *) 

TargetInsert[target_, targetname_ : "Target", produce_ : True] := 
 Module[{modeldir,modelfile, columntarget, rows, cols, targetrow, targetcol, prod, drains, sink,sinks,oldtargets},
  (* Function to define a bioengineering target metabolite(s) and if needed extend stoichiometry matrix.*)
  
  (* The target is specified in the first argument in one of 3 ways:
    1) It can be a string, contained in the name of an existing metabolite. 
  In this case, if there is an existing sink reaction in S for the metabolite 
  produced (or a source if it is absorbed) that is taken as the target flux.
  If not, an appropriate extra sink/source flux is added to S, to serve as target.
    2) If a single number,this is taken the number of a column in S to be the target flux.
  	3) If a vector, this is added as an additional column to S and taken as the target. 
  It has to have an entry for each metabolite, negative for those produced and positive for those absorbed.
  The last argument sets whether the target metabolite is to be produced (the default) or absorbed by the network. 
  This is optional and used only with a named target. 
  
  It returns the number of the target column, or an error message.
  
  NOTE: TargetInsert modifies various global arrays defined in Configuraion.m  *)
  modeldir=If[ValueQ[datafile],FileNameDrop[datafile],Directory[]];
  Species="ToBeLoaded";  (* Avoids error message in reader function *)
  {rows, cols} = If[Dimensions[S] == {0}, {0, 0}, Dimensions[S]];
(* Print["Modeldir is "<>modeldir<>" and I have S dimensions "<>ToString[{rows, cols}]]; *)
  If[{rows, cols} != {Length@MetaboliteNames, Length@ReactionNames}, 
   modelfile = SystemDialogInput["FileOpen", {modeldir, 
   	{"Metab model, text" -> {"*.mat", "*.m", "*.WL"}, "Metab model, sbml" -> {"*.sbml", "*.xml"}, 
       "All files" -> {"*"}}},  WindowTitle -> 
      "Stoichiometry inconsistent; select the original data file for reloading."];
   If[StringQ[modelfile], LoadModel[modelfile, False]]
   ];
   Species = If[ValueQ[modelfile], FileNameTake[modelfile, {-2}], "Unknown Species"];
  {rows, cols} = If[Dimensions[S] == {0}, {0, 0}, Dimensions[S]];
  If[{rows, cols} != {Length@MetaboliteNames, Length@ReactionNames}, 
   MessageDialog["SSKernel results and current S matrix have incompatible \
dimensions.\nReload the model or refresh the SSKernel data first!"]; 
   Return["Incompatible dimensions for name lists and current stoichiometry matrix. "]];
  
  prod = If[produce, 1., -1.];
  columntarget = target;
  targetcol = target;
  
  If[StringQ[columntarget],
   targetrow = Flatten@Position[MetaboliteNames, _?(StringContainsQ[#, columntarget, 
          IgnoreCase -> True] &), Heads -> False];
   (*Print[{"targetrow = ",targetrow}];*)
   Switch[Length@targetrow,
    0, MessageDialog[
     "Invalid target - given string is not recognised as a metabolite in the model!"];
     Return["Invalid target."],
    1,(*Metabolite identified. Check if a sink column exists, else create one. *)
    drains = Flatten@Position[First@S[[targetrow]], _?(prod*# < 0 &)];
     sink = Extract[drains, Position[Transpose[S][[drains]],
     	 {RepeatedNull[0 | 0.], _?(prod*# < 0 &), RepeatedNull[0 | 0.]}]];
    targetcol = If[Length[sink] == 1, sink[[1]], 
      columntarget = -prod*UnitVector[rows, First@targetrow]; 0],
    _,(* Metabolite name not unique, let user choose which one to use. *)
    sinks = Reap[Do[
    	drains = Flatten@Position[S[[targetrow[[i]]]], _?(prod*# < 0 &)]; 
        sink = Extract[drains, Position[Transpose[S][[drains]],
        	 {RepeatedNull[0 | 0.], _?(prod*# < 0 &), RepeatedNull[0 | 0.]}]];
        Sow[If[Length@sink == 1, {True, First@sink}, {False, 0}]]
        , {i, Length@targetrow}]][[2, 1]];
    MessageDialog["Several metabolites have <" <> target <> 
      "> as part of its name, in row numbers as follows:\n" <> 
      ToString[Transpose[{targetrow, MetaboliteNames[[targetrow]]}]] <>
       "\n\nA unique source/sink reaction column exists only for the cases as follows:\n" <> 
      ToString[Transpose[{MetaboliteNames[[Pick[targetrow, sinks[[All, 1]]]]],
          Pick[sinks[[All, 2]], sinks[[All, 1]]]}]] <> 
      "\n\nRerun TargetInsert with either a more explicit name, or the unique column number, or an explcit column vector with " <> ToString[rows] <> " components."];
    Return["User to choose between columns " <> 
      ToString[Pick[sinks[[All, 2]], sinks[[All, 1]]]]]
    ]
   ];
  (*Print["Failed the string test"];*)
  (* Remove target designations that were assigned in previous TargetInsert runs *)
  oldtargets = Flatten@Position[ReactionNames,
  	 _?(StringContainsQ[#, "Target"] &), Heads -> False]; 
  ReactionNames = If[Length@oldtargets > 0, 
    StringTrim@First@Map[ReplacePart[ReactionNames, # -> 
          Last@StringSplit[ReactionNames[[#]], ":"]] &, oldtargets], 
    ReactionNames];
  
  If[VectorQ[columntarget],
   If[Length@columntarget != rows, 
    MessageDialog["Invalid target - it should either be a string contained in the \
name of an existing metabolite, the number of an existing \
stoichiometry column, or a vector of " <> ToString[rows] <> 
      " values that identify target metabolite proportions and can be \
added as an additional column of S !"];
	Return["Invalid target."], 
    targetcol = 0];
   (*Print["Passed the valid column test"];*)
   (* Add the new column to S if no sink was found, else just the new name *)
   If[targetcol == 0,
    S = Join[S, Transpose@{columntarget}, 2];
    {rows, cols} = Dimensions[S];
    FBAdone = False;
    feasiblepoints = {};
    Svals = ConstantArray[{0, 0}, Length[S]];(* {value, type} pairs; type=0 => equal *)
    AppendTo[bounds, {0., 999.99}];
    AppendTo[objectselector, 0];
    AppendTo[ReactionNames, targetname];
    MessageDialog[
     "An additional flux was added to the model.\n
     Rerun the SSK calculation before analysing this target."];
    targetcol = cols,
    
    ReactionNames[[targetcol]] = "Target: "<> ReactionNames[[targetcol]]],
   
   (*Print["Target spec is only a number "];*)
   ReactionNames[[targetcol]] = "Target: " <> ReactionNames[[targetcol]]];
  {Sraw, rawSvals, rawbounds, rawobject, rawFBAvec, rawreacts, 
    rawmets} = {S, Svals, bounds, objectselector, FBAvector, 
    ReactionNames, MetaboliteNames};
  targetcol
  ]

HyperSampler[hypercon_, hypervals_, PeriPoints_, Rays_, ReducedSS_, 
  targetcol_ : -1, PrintResult_ : False] :=
 (* This function returns a set of sample points and rays that fall on 
 the hyperplane specified by the constraints and values set, hypercon and hypervals.
 Each list element of argument hypercon specifies a hyperplane that \
will be sampled. This can have one of two forms: either a list of \
flux numbers that are constant on the hyperplane, or a 2D array that 
specifies the constraint vectors that define the hyperplane.
 Each list element of argument hypervals gives the set of constant \
values for the corresponding hyperplane.
 This can also have two forms, allowing either a strict hyperplane, \
or using inequalities to define a subregion bounded by the hyperplane, to be given. 
 If just a single number, the constraint is taken as "equal" with the value given. 
 If it is a number pair (in braces} the first is the number, the second can be -1,0 or 1
  to indicate respectively "less or equal","equal" or "greater or equal". 
 The remaining arguments specify SSK periphery points, the known rays and the RSS, 
 	all specified relative to FFS, and the number of the column of S for the target flux.
 Each periphery point is projected to the hyperplane by interpolation 
 	and/or extrapolation using another periphery point or a ray.
 NOTE that there may be SS regions that cannot be reached from the PPP 
 by either interpolation or ray extrapolation. 
 To compensate, up to three additional points calculated by LP are added to the sample. 
 First, an intercept between H and the SS is calculated. If any point in this 
 intercept exists, it is included in the sample. 
 Then two more points that respectively minimize/maximize the target flux value in \
the intercept are found. In case either is unbounded, a nominal \
sample point at +/- Infinity is used. *)
 Module[{hyperc, hyperv, pointcount, concount, varcount, 
   emptyh = False, hcons,
   ord, eqs, ineqs, i, j, zerohyps, overlapat, pairs, insample = {}, 
   exsample = {}, extremes, slacks, regionpat, inregion, onboundary, 
   onsingle, sample, focusgroup = {}, focuspoints, CHBT, CHPT,  
   RSSorig, RSSbasis, RSScons, RSSvals, A, U, UU, objective, 
   rayweights, pointcoef, rayinregion, Hrays = {}, 
   Brays = {}, sortrays, BH, Q, raytest, isray, overlaps, elims, 
   elimrays = {}, remrays = {}, raycount, rayresult, elimcount = 0, 
   remcount = 0, CHRRT, CRRRT, CHRET, row1, row2, row3, row4, 
   intercept, ExtBasis, minpoint, maxpoint, UXrange, XYrange},
  hyperv = hypervals;
  pointcount = Length@PeriPoints;
  raycount = Length@Rays;
  RSScons = ReducedSS[[1, 1]];
  RSSvals = ReducedSS[[1, 2]];
  RSSorig = ReducedSS[[2, 1]];
  RSSbasis = ReducedSS[[2, 2]];
  {concount, varcount} = Dimensions@RSScons;
  hcons = Length@hypercon;
  
  (* STAGE 0:  Convert mixed equalities and inequalities to standard form *)
  
  If[((hypercon == {} || hypercon == {{}}) && hypervals == {} ),
   hcons = 0; emptyh = True,
   If[( hcons != Length@hypervals), 
    Print["Invalid hyperplane specification; number of constraints \
and values do not match!"]; Return[{{}, {}}]]];
  (* Convert hyperplane constraints given as a list of numbers
   to the actual constraint vectors *)
  hyperc = If[MatrixQ[hypercon], hypercon,
    UnitVector[Length@RSSorig, #] & /@ hypercon];
  (* Constraint vectors normalised and \[GreaterEqual] changed into \[LessEqual] ;
  	inequalities counted and sorted first; value/ineq pairs converted back to simple list*)
  hyperv = Replace[hyperv, val_?NumberQ :> {val, 0}, {1}];
  Do[If[hyperv[[i, 2]] > 0, hyperc[[i]] = -hyperc[[i]];
  		hyperv[[i]] = -hyperv[[i]]], {i, Length@hyperv}];
  ord = OrderingBy[hyperv, Last];
  hyperv = hyperv[[ord]];
  hyperc = Normalize /@ hyperc[[ord]];
  ineqs = Count[hyperv[[All, 2]], _?Negative];
  hyperv = hyperv[[All, 1]];
  eqs = hcons - ineqs;
  overlapat = Position[hyperc . Transpose[hyperc], _?(Abs[#] == 1 &)];
   pairs = Cases[overlapat, {a_, b_} /; (a > b && a > ineqs)]; 
  If[pairs != {}, 
   Print["ABORT: Parallel hyperplanes specified, at least one of \
which is a strict equality. \nThese are contradictory or at least \
redundant; please correct!"]; Return["Failed"]];
  If[PrintResult,
   If[ineqs == 0, 
    Print["The given constraints define a hyperplane at facet level " <> ToString[eqs]], 
    Print["The given constraints define an SS subregion at facet \
level " <> ToString[eqs] <> " and bordered by " <> ToString[ineqs] <> 
      " additional hyperplanes."]]
   ];
  (* Downcast the hyperplane constraints to RSS *)
  CHBT = If[emptyh, {}, hyperc . Transpose[RSSbasis]];
  
  (* STAGE 1: Starting sample and sampling focus group. *)
  
  If[emptyh, sample = PeriPoints; focusgroup = {},
   slacks = Map[(# - hyperv) &, PeriPoints . Transpose[hyperc]];
   regionpat = {Repeated[_?(# <= 10.^-6 &), {ineqs}], _?(Abs[#] <=10.^-6 &) ...}; 
   inregion = Map[MatchQ[#, regionpat] &, slacks];
   onboundary = If[ineqs > 0,
     onsingle = Apply[Or, #] & /@ 
       Map[(Abs[#] <= 10.^-6) &, slacks[[All, ;; ineqs]], {2}];
     MapThread[#1 && #2 &, {inregion, onsingle}],inregion];
   (*Print[{"inregion, onboundary: ",inregion, onboundary}];*)
   sample = Pick[PeriPoints, inregion];
   focusgroup = Pick[Range[pointcount], Not /@ onboundary]];
  focuspoints = Length@focusgroup;
  If[PrintResult, 
   Print[ToString[Length@sample] <> 
     " given periphery points already satisfy hyperplane constraints \
and are kept as sample points.\nPoint no's " <> ToString[focusgroup] <>
      " are to be resampled."]];
  
  (* STAGE 2: RAY PROCESSING. Put non-RSS rays last and 
  find subregion rays to be exported. *)
  
  (*  Identify any rays that were eliminated in forming the RSS, 
  and rearrange the rays to put those last, 
  as that is required for setting up the LP arrays.*)
  If[raycount > 0,
   (* The SS is unbounded. 
   Get a ray basis for the H and SS intersection by projection.  *)
   overlaps = Rays . Transpose[RSSbasis];
   elims = Flatten@Position[overlaps, {_?(Abs[#] < 10.^-6 &) ..}];
   elimcount = Length@elims;
   elimrays = If[elimcount == 0, {}, Rays[[elims]]];
   remrays = Complement[Rays, elimrays];
   (*Print["elimrays: ",elimrays,"remrays: ",remrays];*)
   remcount = Length@remrays;
   sortrays = 
    Which[elimcount == 0, remrays, remcount == 0, elimrays, 
    	True, Join[remrays, elimrays]];
   If[PrintResult, 
    Print["Previous RSS reduction had eliminated " <> 
      ToString[elimcount] <> " rays orthogonal to RSS while " <> 
      ToString[remcount] <> " rays belong to RSS itself. "]];
   (*Print[{"sortrays: ",sortrays}];*)
   (* Start the intersection ray set as those among the SS rays that satisfy H constraints *)
   Hrays = If[emptyh, sortrays,
     slacks = sortrays . Transpose[hyperc];
     (*Print[{"Ray intersection slacks: ",slacks}];*)
     regionpat = {Repeated[_?(# <= 10.^-6 &), {ineqs}], _?(Abs[#]<=10.^-6 &) ...}; 
     rayinregion = Map[MatchQ[#, regionpat] &, slacks];
     Pick[sortrays, rayinregion]];
   
   (* Construct rays aligned along the intersection and the new \
boundary and add them to the Hrays set *)
   (* All rays, even those that fall in H,  are included in this. 
   They will be unchanged when projected to H itself, 
   but may project to boundary rays when there are inequalities. *)
   i = 0;
   Until[i > ineqs,
    zerohyps = If[i == 0,
      If[eqs > 0, hyperc[[-eqs ;;]], {{}}],
      Which[
       ineqs == 0, hyperc,
       ineqs < hcons, Prepend[hyperc[[-eqs ;;]], hyperc[[i]]],
       ineqs == hcons, {hyperc[[i]]}]
      ];
    (*Print[{"i, zerohyps: ",i,zerohyps}];*)
    BH = NullSpace[zerohyps];
    Q = If[BH == {}, {}, 
      Select[Rays . Transpose[BH] . BH, Norm[#] > 10.^-6 &]];
    (*Print[{"Q: ",Q}];*)
    If[Length@Q > 0,
     Q = Normalize /@ DeleteDuplicates[Q, Norm[#1 - #2] < 10.^-6 &]; 
     raytest = Join[CHBT, RSScons] . (RSSbasis . Transpose[Q]);
     isray = Position[Transpose[raytest], _?(Max[#] <= 10.^-6 &), 1];
     Brays = Join[Brays, Extract[Q, isray]]];
    i++;
    ];
   (*Print[{"Number of rays found by projection: ",Length@Brays}];*)
   Brays = 
    Complement[Brays, Hrays, SameTest -> (Norm[#1 - #2] < 10.^-6 &)];
   Hrays = Join[Hrays, Brays];
   If[PrintResult,
    rayresult = If[Hrays == {{}} || Hrays == {},
      " containing no rays.",
      " containing " <> ToString[Length@Hrays] <> " rays of which " <>
        ToString[Length@Brays] <> 
       " are boundary rays constructed by projection."];
    If[! emptyh, 
     If[ineqs == 0,
      Print[
       "Sampling a " <> ToString[varcount - eqs] <> 
        "-dimensional strict hyperplane," <> rayresult], 
      Print["Sampling a " <> ToString[varcount - eqs] <> 
        "-dimensional region, with " <> ToString[ineqs] <> 
        " additional boundaries and" <> rayresult]]]]
   ];
   
  (* STAGE 3: Deal with an empty H spec, 
  set up matrices for LP and make sure there is an intercept *)
  
  If[emptyh, (* Use RSS itself, extended by prismatic rays *)
   sample = PeriPoints;
   onboundary = ConstantArray[True, Length@PeriPoints];
   If[elimcount > 0,
    A = Join[RSScons, ConstantArray[0., {concount, elimcount}], 2];
    UU = Transpose@{RSSvals, ConstantArray[-1., concount]};
    UXrange = 
     Join[ConstantArray[{-Infinity, Infinity}, varcount], 
      ConstantArray[{0., Infinity}, elimcount]],
    
    A = RSScons;
    UU = Transpose@{RSSvals, ConstantArray[-1., concount]};
    UXrange = ConstantArray[{-Infinity, Infinity}, varcount];
    ];
   A = Chop[A,10.^-6];
   (*Print["Matrix Dimensions for RSS alone: ",Dimensions/@{A,UU,UXrange}];*)
   
   , (* Else it is the intersection of the hyperplane with the extended RSS *)
   CHPT = hyperc . Transpose[PeriPoints];
   (* First task is to set up the LP matrices for the intersection test. *)
   If[elimcount > 0,
    CHRET = hyperc . Transpose[elimrays];
    row1 = Join[CHBT, CHRET, 2];
    row2 = Join[RSScons, ConstantArray[0., {concount, elimcount}], 2];
    A = Join[row1, row2];
    UU = Transpose@{Join[hyperv - hyperc . RSSorig, RSSvals], 
       Join[ConstantArray[-1., ineqs], ConstantArray[0., eqs], 
        ConstantArray[-1., concount]]};
    objective = Join[ConstantArray[0., varcount], ConstantArray[1., elimcount]];
    UXrange =   Join[ConstantArray[{-Infinity, Infinity}, varcount], 
      ConstantArray[{0., Infinity}, elimcount]],
    
    A = Join[CHBT, RSScons];
    UU = Transpose@{Join[hyperv - hyperc . RSSorig, RSSvals], 
       Join[ConstantArray[-1., ineqs], ConstantArray[0., eqs], 
        ConstantArray[-1., concount]]};
    objective = ConstantArray[0., varcount];
    UXrange = ConstantArray[{-Infinity, Infinity}, varcount];
    ];
   A = Chop[A,10.^-6];
   (*Print["Matrix Dimensions for intersection test: ",Dimensions/@{A,UU,UXrange}];*)
   
   (* If known points not feasible, 
   check for the existence of an intercept, else no sample points *)
   If[Count[inregion, True] == 0,
    (*Print["{A,UU,objective,UXrange}",{A,UU,objective,UXrange}];*)
    intercept = Quiet[Check[
       LinearProgramming[objective, A, UU, UXrange, Method->"Simplex", Tolerance->10.^-3], {},
        {LinearProgramming::lpsnf, LinearProgramming::lpsnfp, LinearProgramming::lpdinf, 
        	LinearProgramming::lpdinfp}],
        {LinearProgramming::lpsnf, LinearProgramming::lpsnfp, LinearProgramming::lpdinf, 
        	LinearProgramming::lpdinfp}];
    (*Print[{"intercept: ",intercept}];*)
    If[intercept == {}, If[PrintResult,
      Print["No intercept of the specified hyperplane and the SS was found."]];
       Return[{{}, {}}],
     AppendTo[sample,
      UpliftPoint[intercept[[;; varcount]], {RSSorig, RSSbasis}] + 
       intercept[[varcount + 1 ;;]].elimrays];
     If[PrintResult, 
      Print["No periphery point falls on the hyperplane, but an \
interception point was found and added to the sample set."]; 
      intercept = True]]
    ,
    intercept = True];
   ];
  (* STAGE 4: There is an intercept, 
  so find two sample points that have max/min target flux value. *)
  
  ExtBasis = Join[RSSbasis, elimrays];
  objective = ExtBasis[[All, targetcol]];
  (*Print["Matrix Dimensions for min/max search: ",
  Dimensions/@{objective, A,UU,UXrange}];*)
  minpoint = Quiet[Check[
     LinearProgramming[objective, A, UU, UXrange, Method->"Simplex", Tolerance->10.^-3], -Infinity, 
      {LinearProgramming::lpsub, LinearProgramming::lpdinf, LinearProgramming::lpdinfp}], 
      	{LinearProgramming::lpsub, LinearProgramming::lpdinf, LinearProgramming::lpdinfp}];
  maxpoint =  Quiet[Check[
     LinearProgramming[-objective, A, UU, UXrange, Method->"Simplex", Tolerance -> 10.^-3],Infinity, 
      {LinearProgramming::lpsub, LinearProgramming::lpdinf, LinearProgramming::lpdinfp}], 
      	{LinearProgramming::lpsub, LinearProgramming::lpdinf, LinearProgramming::lpdinfp}];
  (*Print[{"Min/max points: ",minpoint,maxpoint}];*)
  (* In case the LP was unbounded, 
  form a nominal sample point by adding the 
  implied ray vector along the flux axis of the target flux *)
  minpoint = 
   If[VectorQ[minpoint], UpliftPoint[minpoint, {RSSorig, ExtBasis}], 
    ReplacePart[RSSorig, targetcol -> minpoint]];
  maxpoint = 
   If[VectorQ[maxpoint], UpliftPoint[maxpoint, {RSSorig, ExtBasis}], 
    ReplacePart[RSSorig, targetcol -> maxpoint]];
  extremes = {minpoint, maxpoint};
  (*Print[{"Full range as per uplifted Min/max points: ",
  extremes[[All,targetcol]]}];*)
  If[Length@focusgroup == 0, 
   If[PrintResult, 
    Print["All periphery points fall on the hyperplane. The required \
sample are those plus extreme target points."]];
   sample = 
    Chop@DeleteDuplicates[Join[sample, extremes], 
      Norm[#1 - #2, \[Infinity]] < 10^-6 &];
   Return[{sample, Hrays}]];
  
  (* STAGE 5: INTERPOLATION. IF the SS is bounded, 
  just interpolate and exit. *)
  
  (* If there are no rays, extrapolation is not applicable so just 
  interpolate additional sample points on the PPP/hyperplane 
  intersection from the remaining offplane periphery points.  *)
  If[raycount == 0 ,
   If[PrintResult, 
    Print["Interpolation only, since the RSS is bounded."]];
   A = Append[CHPT, ConstantArray[1., pointcount]];
   U = Transpose@{Append[hyperv, 1.], 
      Join[ConstantArray[-1., ineqs], ConstantArray[0., eqs + 1]]};
   (*Print["Matrix Dimensions for point interpolation: ",
   Dimensions/@{objective,A,U}];*)
   (* The sampling loop over the periphery points in the focus group *)
   insample = Reap[Do[
       objective = -UnitVector[pointcount, focusgroup[[i]]];
       (*Inner loop to address H subspace as a whole and then each boundary facet in turn *)
       j = 0;
       Until[j > ineqs,
        pointcoef = 
         Quiet[Check[
           LinearProgramming[objective, Chop[A,10.^-6], U], {},
            {LinearProgramming::lpsnf, LinearProgramming::lpsnfp, LinearProgramming::lpdinf, 
        	LinearProgramming::lpdinfp}], 
           {LinearProgramming::lpsnf,LinearProgramming::lpsnfp,LinearProgramming::lpdinf, 
        	LinearProgramming::lpdinfp}];
       (*Print[{"Interpolation coefficients: ", pointcoef}];*)
        If[pointcoef != {}, Sow[pointcoef.PeriPoints]];
        j++;
        (* Change the next inequality to equality, and restore the present one *)
        UU[[j, 2]] = 0;
        If[j > 1, UU[[j - 1, 2]] = -1]], {i, focuspoints}]][[2]];
   If[insample == {}, 
    If[PrintResult, 
     Print["Periphery points outside the hyperplane did not yield any \
interpolated points on it."]], 
    insample = Chop@DeleteDuplicates[insample[[1]], Norm[#1 - #2, \[Infinity]] < 10^-6 &]; 
    If[PrintResult, 
     Print[ToString[Length@insample] <> 
       " distinct sample points were interpolated to the hyperplane."]]];
   sample = Chop@DeleteDuplicates[Join[sample, insample, extremes], 
      Norm[#1 - #2, \[Infinity]] < 10^-6 &];
   Return[{sample, {}}]];
  
  (* STAGE 6: Combined interpolation and extrapolation.*)
  
  (* STEP 1: 
  set up the constraint matrices for the various cases of how many rays in each class *)
  U = Transpose[{Append[hyperv, 1.], 
     Join[ConstantArray[-1., ineqs], ConstantArray[0., eqs + 1]]}];
  Which[
  	elimcount == 0,
   CHRRT = hyperc . Transpose[remrays];
   CRRRT = RSScons . ( RSSbasis . Transpose[remrays]);
   row1 = Join[CHPT, CHRRT, 2];
   row2 = {Join[ConstantArray[1., pointcount], ConstantArray[0., remcount]]};
   row3 = Join[ConstantArray[0., {concount, pointcount}], CRRRT, 2];
   A = Join[row1, row2, row3];
   UU = Join[U, ConstantArray[{0., -1.}, concount]];
   XYrange = Join[ConstantArray[{0., Infinity}, pointcount], 
     ConstantArray[{-Infinity, Infinity}, remcount]],
     
   remcount == 0,
   CHRET = hyperc . Transpose[elimrays];
   row1 = Join[CHPT, CHRET, 2];
   row2 = {Join[ConstantArray[1., pointcount], 
      ConstantArray[0., elimcount]]};
   row3 = Join[ConstantArray[0., {elimcount, pointcount}], -IdentityMatrix[elimcount], 2];
   A = Join[row1, row2, row3];
   UU = Join[U, ConstantArray[{0., -1.}, elimcount]];
   XYrange = Join[ConstantArray[{0., Infinity}, pointcount], 
     ConstantArray[{-Infinity, Infinity}, elimcount]],
     
   True,
   CHRRT = hyperc . Transpose[remrays];
   CRRRT = RSScons . ( RSSbasis . Transpose[remrays]);
   CHRET = hyperc . Transpose[elimrays];
   row1 = Join[CHPT, CHRRT, CHRET, 2];
   row2 = {Join[ConstantArray[1., pointcount], 
      ConstantArray[0., remcount + elimcount]]};
   row3 = Join[ConstantArray[0., {concount, pointcount}], CRRRT, 
     ConstantArray[0., {concount, elimcount}], 2];
   row4 = Join[ConstantArray[0., {elimcount, pointcount + remcount}],
   	 -IdentityMatrix[elimcount], 2];
   A = Join[row1, row2, row3, row4];
   UU = Join[U, ConstantArray[{0., -1.}, concount + elimcount]];
   XYrange = Join[ConstantArray[{0., Infinity}, pointcount], 
     ConstantArray[{-Infinity, Infinity}, (remcount + elimcount)]];
   ];
  A = Chop[A,10.^-6];
 (* Print["Matrix Dimensions before extrapolation loop: ", 
  	Dimensions /@ {objective, A, UU, XYrange}]; *)
 (* Print["Matrix dimensions of row groups ",Dimensions /@ {row1, row2, row3, row4}];*)
 
  (* STEP 2: The sampling loop over the periphery points in the focus group *)
  exsample = Reap[Do[
      (* Inner loop to address H subspace as a whole and then 
      each boundary facet in turn *)
      j = 0;
      Until[j > ineqs,
       (* Assign random low, positive weights to rays, to counteract 
       bias to rays generally and any specific ray in particular *)
       rayweights = RandomReal[1, raycount];
       objective = Join[-UnitVector[pointcount, focusgroup[[i]]], 
         rayweights/(5*Total@rayweights)];
        
        (* MATHEMATICA 13 BUG? 
        Quiet in the next line still lets "No solution" messages through.
        Completely switching these off with the Off instruction is no solution,
        because then the Check does not work either. 
        Might improve in future Mathematica version? *)
        
       pointcoef = Quiet[Check[
       	LinearProgramming[objective, A, UU,XYrange], {},
       		{LinearProgramming::lpsnf, LinearProgramming::lpsnfp, LinearProgramming::lpdinf, 
        	LinearProgramming::lpdinfp}],
       		 {LinearProgramming::lpsnf, LinearProgramming::lpsnfp, LinearProgramming::lpdinf, 
        	LinearProgramming::lpdinfp}];
       (*Print["Coefficients for peri point no "<>ToString[i]<>" :", pointcoef];*)
       If[pointcoef != {}, 
        Sow[Transpose[pointcoef[[;; pointcount]].PeriPoints + 
           pointcoef[[pointcount + 1 ;;]].sortrays]]];
       j++;
       (* Change the next inequality to equality, and restore the present one *)
       UU[[j, 2]] = 0;
       If[j > 1, UU[[j - 1, 2]] = -1]]
      ,
      {i, focuspoints}]][[2]];
  If[exsample == {}, 
   If[PrintResult, 
    Print["Combined interpolation/extrapolation attempted, but no new \
points were found. "]],
   exsample = DeleteDuplicates[exsample[[1]],Norm[#1 - #2, \[Infinity]] < 10^-6 &]; 
   If[PrintResult, 
    Print[ToString[Length@exsample] <> 
      " distinct sample points were found by combined interpolation and extrapolation."]]];
  (*Print["Sample counts ",Length/@{sample,insample,exsample, extremes}];*)
  sample = Chop@DeleteDuplicates[Join[sample, exsample, extremes], 
     Norm[#1 - #2, \[Infinity]] < 10^-6 &];
  {sample, Hrays}
  ]
  
TargetVariation[hyperlist_, valuelist_, perips_, rays_, 
  ReducedSS_, targetcol_ : -1, PrintResult_ : False] :=
 (* This function generates descriptive statistics of the variation of a target flux.
 This is taken over a set of hyperplane intersections with the FBA solution space, 
 calculated for an optimized objective. 
 The function returns the complete set of the SS sample points it generates, as
 a nested list containing the samples for each hyperplane intersection separately.
 
 Each element of argument hyperlist specifies a hyperplane that will \
be sampled. This can have one of two forms: 
either a list of flux numbers that are constant on the hyperplane, 
 or a 2D array that specifies the constraint vectors that define the hyperplane.
 Each element of argument valuelist gives the set of constant values \
for the corresponding hyperplane. 
 The next 3 arguments give the SSK specification,as used by HyperSampler 
 to find the set of sample points on the requested hyperplane. 
 The target flux column can be given as an optional last argument, \
otherwise it is assumed to have been appended to the stoichiometry matrix 
and so forms the last column. 
 *)
 Module[{fluxcount, SSMedian, SSMean, SSstd, SSRange, SStats, jlist, hypercount,
   samplecount, medlist, meanlist, stdlist, rangelist,
    Hsample, Hrays, SampleValues, SampleMean, SuperSample, SampleMedian, SampleStd,
    SampleRange, ord, stats, title, table, numberedstats, samps, meds, means, stds, ranges},

  If[! NumberQ[targetcol], MessageDialog[
  "The target column argument has to be a single number!"]; 
  Return["Invalid argument."]];
  hypercount = Length@valuelist;
  If[Length@hyperlist != hypercount, 
  MessageDialog[
   "Invalid arguments: hyperlist has " <> ToString[Length@hyperlist] <>
     " members while valuelist has " <> ToString[hypercount]]; 
  Return["Invalid argument."]];
  fluxcount = Length@ReducedSS[[2, 1]];
  If[fluxcount != Dimensions[S][[2]], MessageDialog[
  "Incompatible data. The loaded stoichiometry has " <> 
   ToString[Dimensions[S][[2]]] <> " fluxes while the RSS from the SSK run has " <> 
   ToString[fluxcount] <> " flux dimensions.\n
   If a target column was added, SSKernel has to be run on this extended model first."];
   Return["Invalid data."]];
   If[!commandline,
   	PrintTemporary[Labeled[ProgressIndicator[Dynamic[j], {0, hypercount}, 
    ImageSize -> {260, Automatic}], 
   "Sampling " <> ToString[hypercount] <> " listed hyperplanes"]]];
   j = 0.5;
  (* Get sample points and statistics for the RSS extended by rays *)
  {Hsample, Hrays} = 
   HyperSampler[{}, {}, perips, rays, ReducedSS, targetcol, PrintResult];
  SampleValues = Select[Hsample[[All, targetcol]], Abs[#] < Infinity &];
  SSMedian = Median[SampleValues];
  SSMean = Mean[SampleValues];
  SSstd = If[Length@SampleValues < 2, 0, Chop@StandardDeviation[SampleValues]];
  SSRange = MinMax[Hsample[[All, targetcol]]];
  SStats = {"SS Kernel", "--", Length@Hsample, SSMedian, SSMean, SSstd,Splice@SSRange};
  (*Print[{"SSMean,SSRange ",SSMean,SSRange}];*)
  (* Collect the sampled points collected from all the listed hyperplanes together. *)
  SuperSample = {Hsample};
  
  {jlist, samplecount, medlist, meanlist, stdlist, rangelist} = 
   Reap[Do[ If[commandline,WriteString[OutputStream["stdout",1] ,"."]];
      {Hsample, Hrays} = 
       HyperSampler[hyperlist[[j]], valuelist[[j]], perips, rays, ReducedSS, 
        targetcol, PrintResult];
       SuperSample = Append[SuperSample, Hsample];
      If[Hsample == {},
       SampleMedian = "--";
       SampleMean = -Infinity; (* To get proper ordering below *)
       SampleStd = "--";
       SampleRange = {"--", "--"},
       
       SampleValues = Select[Hsample[[All, targetcol]], Abs[#] < Infinity &];
       SampleMedian = Median[SampleValues];
       SampleMean = Mean[SampleValues];
       SampleStd = If[Length@SampleValues < 2, 0, 
         Chop@StandardDeviation[SampleValues]];
       SampleRange = MinMax[Hsample[[All, targetcol]]];
       (*If[Hrays!={}&& Max@Abs[Hrays[[All,targetcol]]]>0,
       SampleRange={"None found","None found"}]*)];
      If[MatrixQ[hyperlist[[j]]], 
       Sow[Table["HypPlane " <> ToString[i], {i, Length@hyperlist[[j]]}], 
        jlist], Sow[ReactionNames[[#]] & /@ hyperlist[[j]], jlist]
        ];
      Sow[Length@Hsample, samps];
      Sow[SampleMedian, meds];
      Sow[SampleMean, means];
      Sow[SampleStd, stds];
      Sow[SampleRange, ranges]
      , {j, hypercount}]][[2]];
  Print["\nNumber of distinct sample points, across all " <> ToString[hypercount] <>
  	 " probed hyperplanes: ", Length[DeleteDuplicates@Flatten[SuperSample, 1]]];

  (*   Sort by mean and present the results in a table form. *)
  ord = Ordering[meanlist, All, Greater];
  SuperSample[[2;;]]=SuperSample[[1+ord]];
  stats = Transpose@{jlist, valuelist, samplecount, medlist, 
     meanlist /. -Infinity -> "--", stdlist, rangelist[[All, 1]], 
     rangelist[[All, 2]]};
  stats = stats[[ord]];
  stats[[All, 1]] = Style[ToString[#], Small] & /@ stats[[All, 1]];
(* Convert value sets to a single string, to avoid deeper nested list \
that does not display well *)
  stats[[All, 2]] = 
  Replace[stats[[All, 2]], 
  	{{val_, sign_} :> "\[GreaterEqual] " <> ToString[val] /; sign > 0, 
     {val_, sign_} :> "\[LessEqual] " <> ToString[val] /; sign < 0, 
     {val_, sign_} :>  ToString[val] /; sign == 0, 
              val_ :> ToString[val]}, 
     {2}];
  stats[[All, 2]] = Map[Style[#, Small] &, stats[[All, 2]], {1}];
  stats = Join[{SStats}, {ConstantArray[" ", 8]}, stats];
  numberedstats = 
   Join[Transpose@{Insert[Range[Length@hyperlist + 1], " ", 2]}, stats, 2];
   title = "Variation of target flux (" <> ReactionNames[[targetcol]] <> 
   ") in response to assigning flux values.";
  table = NumberForm[
   TableForm[numberedstats[[All, {1, 2, 3, 4, 6, 8, 9}]], 
    TableHeadings -> {None, {"TRIAL\nNo", "ASSIGNED \nFluxes", 
       "\nValues", "SAMPLE\nCount"(*,"Median"*), "TARGET\nMean"(*,
       "Std dev"*), "\nLower", "\nUpper"}}, TableSpacing -> {0,3}], 3];
  Print[""];
  Print[Style[Capitalize[title, "TitleCase"], Bold, Blue]];
  If[commandline,Print["\n"<>ToString@table],Print[table]];
  SuperSample
  ]
  
Formula[react_] := 
 Module[{reactlist, Scol, inflows, outflows, left, right},
  (* This function can be given a reaction number, 
  or a list of numbers, 
  or a character string contained in a reaction name.
  It prints the reaction as a chemical formula using the metabolite names.*)
  reactlist = 
   If[StringQ[react], 
    Flatten@Position[
      ReactionNames, _?(StringContainsQ[#, react, 
          IgnoreCase -> True] &), Heads -> False], Flatten[{react}]];
  Do[
   Scol = reactlist[[i]];
   inflows = Flatten@Position[S[[All, Scol]], _?(# < 0 &)]; 
   left = MapThread[
     If[FractionalPart@Abs[#1] == 0., IntegerPart@Abs[#1], Abs[#1]] #2 &,
      {S[[inflows, Scol]], MetaboliteNames[[inflows]]}]; 
   left = StringRiffle[ToString /@ left, " + "];
   outflows = Flatten@Position[S[[All, Scol]], _?(# > 0 &)];
   right = MapThread[
     If[FractionalPart@Abs[#1] == 0., IntegerPart@Abs[#1], Abs[#1]] #2 &,
      {S[[outflows, Scol]], MetaboliteNames[[outflows]]}]; 
   right = StringRiffle[ToString /@ right, " + "];
   Print[ReactionNames[[Scol]]<>": "<>left<>" \[LongRightArrow] "<>right],
   {i, Length@reactlist}]
  ]

End[] (* End Private Context *)

EndPackage[]
```
