## Supplementary material for "The FBA Solution Space Kernel – Introduction and Illustrative Examples": SSK software manual and code: StringToDouble.m

```
(* ::Package::*)
(*Author:Hrvoje Abraham 
Date:19 Jun 2016 
Version:3 
License:MIT 

Floating point string to correctly rounded double precision value conversion based on the AlgorithmM 
available in http://www.cesura17.net/~will/professional/research/papers/howtoread.pdf and in 
the simplified form at http://www.exploringbinary.com/correct-decimal-to-floating-point-using-big-integers/
Values are rounded according to IEEE-754 round-half-to-even rule.This is currently the only rounding mode.
It's rigorously tested against the most pathological inputs and halfway cases,
such as specially constructed 1000+bytes long strings and other known-to-be-problematic inputs.
But one never knows... 
https://github.com/ahrvoje/numerics/blob/master/strtod_tests.toml 

Implementation is not speed optimized,safety and transparency was the priority.
Scaling approximation step was introduced in version 2 which considerably improves
 performance for extreme values,but that's about it.
Valid inputs are integers,C99 language standard decimal double precision floating 
constants (section 6.4.4.2) and special values[-]inf and[-]nan (section 7.19.6.1.8).
The only non-standardized constraint is that NaN value string can't contain closing parenthesis ")" 
inside optional parentheses,e.g.input "nan(type_0)_123)" is not allowed,while "nan(type_0_123)" is valid.
This practice is widely spread among standard-compliant implementations.
NaN sign is handled and preserved by all the methods.
Other NaN payload is discarded,in binary representation mantissa is set to 1.
 
 About the methods:
-StringToDoubleKernel:returns double precision value sign,significand and exponent
-StringToDouble:returns double precision value based on StringToDoubleKernel output
-StringToDoubleBin:returns IEEE-754 64-bit binary representation of double precision value
-StringToDoubleHex:returns IEEE-754 64-bit hex representation of double precision value
-StringToDoubleInt:returns unreduced integer representation of double precision value*)

BeginPackage["StringToDouble`"]

StringToDouble::usage="StringToDouble[s] returns the double precision value of the input string."
StringToDoubleBin::usage="StringToDoubleBin[s] returns binary representation of the IEEE-754 double precision value of the input string."
StringToDoubleHex::usage="StringToDoubleHex[s] returns hexadecimal representation of the IEEE-754 double precision value of the input string."
StringToDoubleInt::usage="StringToDoubleInt[s] returns exact integer representation of the double precision value of the input string."

Begin["`Private`"]

StringToDoubleKernel[str_]:=Module[{fpRegex,s,sign,mantissa,exponent,int,frac,decimals,a,scale,n,d,sd,r},(*check the formatting*)fpRegex=RegularExpression["(?i)^[-+]?((((\\d+\\.?)|(\\d*\\.\\d+))(e[-+]?\\d+)?)|(inf(inity)?)|(nan(\\([^)]*\\))?))$"];
If[Length@StringCases[str,fpRegex]==0,Throw["Invalid floating point string format."]];
s=ToLowerCase[str];
(*extract the sign*)sign=1;
If[StringTake[s,1]=="-",sign=-1;s=StringDrop[s,1]];
If[StringTake[s,1]=="+",s=StringDrop[s,1]];
(*check for NaN*)If[StringContainsQ[s,"nan"],Return[{sign,Indeterminate,0}]];
(*check for infinity*)If[StringContainsQ[s,"inf"],Return[{sign,Infinity,0}]];
(*determine the exact value*)If[StringContainsQ[s,"e"],{mantissa,exponent}=StringSplit[s,"e"],{mantissa,exponent}={s,"0"}];
If[StringContainsQ[mantissa,"."],If[StringTake[mantissa,1]==".",{int,frac}={"0",StringDrop[mantissa,1]},If[StringTake[mantissa,-1]==".",{int,frac}={StringDrop[mantissa,-1],"0"},{int,frac}=StringSplit[mantissa,"."]]],{int,frac}={mantissa,"0"}];
decimals=StringLength[frac];
{int,frac,exponent}=ToExpression[{int,frac,exponent}];
(*the exact value of the input string*)a=(int*10^decimals+frac)*10^(exponent-decimals);
(*underflow if at or below halfway between 0 and 2^-1074*)If[a<=2^-1075,Return[{sign,0,0}]];
(*overflow if at or over 1/2 ULP past 2^971(2^53-1)*)If[a>=2^971*(2^53-1+1/2),Return[{sign,Infinity,0}]];
(*scale'a' into[2^52,2^53) interval*)(*scaling approximaton step based on the fast approx for Log2[a]*)scale=Round[3.3*(IntegerLength[int]-1+exponent)]-52;
If[scale<-1074,scale=-1074];
a*=2^-scale;
(*scaling correction loops*)(*if the exact value is below the interval*)While[a<2^52&&scale>-1074,a*=2;scale-=1];
(*if the exact value is above the interval*)While[a>=2^53,a/=2;scale+=1];
(*significand integer part and reminder*)n=Numerator[a];
d=Denominator[a];
sd=IntegerPart[a];
r=n-sd*d;
(*significand rounding according to IEEE-754 round-half-to-even rule*)If[(2*r>d)||(2*r==d&&OddQ[sd]),sd+=1];
(*if rounding got significand out of[2^52,2^53) interval*)If[sd==2^53,sd=2^52;scale+=1];
(*sign,significand,power of 2 scale bias*)Return[{sign,sd,scale}];]

StringToDouble[str_]:=N[#1*#2*2^#3]&@@StringToDoubleKernel@str

StringToDoubleBin[str_]:=Module[{sign,sd,scale,signBin,exponentBin,mantissaBin},{sign,sd,scale}=StringToDoubleKernel[str];
signBin=If[sign==1,{0},{1}];
Which[SameQ[sd,Infinity],exponentBin=IntegerDigits[2047,2,11];mantissaBin=IntegerDigits[0,2,52],SameQ[sd,Indeterminate],exponentBin=IntegerDigits[2047,2,11];mantissaBin=IntegerDigits[1,2,52],sd>=2^52,(*normal*)exponentBin=IntegerDigits[scale+1075,2,11];mantissaBin=Drop[IntegerDigits[sd,2,53],1],True,(*subnormal*)exponentBin=IntegerDigits[0,2,11];mantissaBin=IntegerDigits[sd,2,52]];
Return[IntegerString[FromDigits[Join[signBin,exponentBin,mantissaBin],2],2,64]];]

StringToDoubleHex[str_]:=IntegerString[FromDigits[StringToDoubleBin[str],2],16,16]

StringToDoubleInt[str_]:=Module[{sign,sd,scale,signStr,sdStr,exponentStr},{sign,sd,scale}=StringToDoubleKernel[str];
signStr=If[sign==1,"","-"];
Which[SameQ[sd,Infinity],sdStr="inf";exponentStr="",SameQ[sd,Indeterminate],sdStr="nan";exponentStr="",True,sdStr=ToString[sd];exponentStr=If[scale!=0,StringJoin["*2^",ToString[scale]],""]];
Return[StringJoin[signStr,sdStr,exponentStr]];]

End[]

EndPackage[]
```
