## Supplementary material for "The FBA Solution Space Kernel – Introduction and Illustrative Examples": SSK software manual and code: README.md

# SSKernel
Extraction of the compact kernel of an FBA model of cellular metabolism: Mathematica implementation

This repository implements the Solution Space Kernel (SSK) analysis of FBA solution spaces, as fully documented in the book:

W. S. Verwoerd, The Compact Kernel of a Metabolic Flux Balance Solution Space, World Scientific Publishing Co., Singapore, October 2022 ; ISBN 978-981-12-5583-0, doi:10.1142/12821 .

as descibed in the following summary.

SUMMARY

This monograph offers a fundamentally new approach to facilitate the study of metabolic networks in cells. Two central methods in this field are flux balance analysis (FBA) and extreme pathway analysis, and together form the basis for bioengineering cells to produce desirable metabolic products. The overwhelming number of extreme pathways in a realistic metabolic network presents severe obstacles to this. By contrast, this book focusses on the FBA solution space and simplifies the task of describing it, to extracting just a bounded subspace: the Solution Space Kernel or SSK. 

This reduces the relevant number of flux space dimensions by orders of magnitude, and allows its location, size and shape to be characterised. It is a multi-stage process, requiring many new concepts and algorithms for manipulating polytopes in high dimensional spaces. It identifies directions in which the solution space is unbounded as ray directions, which are partitioned off. This allows a focus on the more significant bounded facets of the SSK, which represent the physically meaningful constraints imposed by the interplay of fluxes in a genome scale metabolic network. 

The book introduces and develops these concepts in a pragmatic way that takes into account the difficulties of performing analyses in a flux space with dimensions counting in the hundreds or thousands. It emphasizes the details of implementation in computational code and applications to realistic models are demonstrated. 
For many cases, the resulting SSK has only single or double-digit dimensions, allowing the range of metabolic states accessible to a cell to be geometrically interpreted in terms of a manageable set of orthogonal diameters and aspect ratios. In addition, explicit representative fluxes, giving the centre and periphery of the solution space kernel, become available for further exploration.
