## Supplementary material for "The FBA Solution Space Kernel – Introduction and Illustrative Examples": SSK software manual and code: README.txt

TESTING SSKERNEL SOFTWARE INSTALLATION

SSKernel is implemented in Wolfram Language and the prerequisite to utilise its interactive user interface, is that the licensed Mathematica software package (Wolfram Research, Inc., Mathematica, Version 14.0, Champaign, IL (2024)) has been installed.

If so, the SSKernel installation can be tested by simply opening and executing the "SSKernel Launcher.nb" notebook in the Mathematica environment. Alternatively the commandine test described below should also work with Mathematica installed.

In the absence of a full Mathematica installation, a command line version can still be run using just the Wolfram Engine software. This is available, free of charge but with usage restrictions, from https://www.wolfram.com/engine/. 

The Wolfram Engine has to have a file association with files having the extension *.wls (Wolfram Language Script). This should automatically happen during installation of the engine.   

The SSKernel operation can then be tested by double clicking either of the files SSKernelscript.wls or SSKernelbatch.bat in Windows Ecplorer. These files are located in the CommandlineFiles subdirectory of the SSKernel download. Alternatively, either of these files can be run without an argument from a command line window. 
If the file association failed, it is still possible to invoke the correct script interpreter using a command such as "wolframscript -f XXX ..." (see the User Manual, Chapter 6, for details).

In either case, follow the prompts in the command line window, which will display a progress report as text message lines. Successful completion will end with the message "Output files have been stored successfully." and the two output file locations are shown just before the success message. 

These files are named "iAB_RBC_283 SSKernelReport.pdf" and "iAB_RBC_283 SSKernel.dif" and can respectively be opened by a PDF reader and a text file editor for inspection, to fully confirm successful completion of the SSKernel analysis.
