## Supplementary material for "The FBA Solution Space Kernel – Introduction and Illustrative Examples": SSK software manual and code: iAB_RBC_283 SSKernelReport.pdf

### Full SS Kernel Analysis for Test

Model iAB_RBC_283 with uncapped kernel of type Compact

calculated on Wed 29 May 2024 in 1.33 seconds.

| | |
|---|---|
| Step time limit | 60. |
| Unbuffered externals exempted | 0 |
| LP tolerance | 0.0001 |
| Fixed value tolerance | 0.002 |
| Progenitor sample size | 20 |
| Max BFBF tree nodes | 100 000 |
| BFBF random greedy sample size | 500 |
| Gready search mixing fraction | 0.8 |

| | |
|---|---|
| Stop sampling when failure rate > | 20 |
| Minimal, Maximal LP chord counts | {10, 50} |
| Maximal flips to find LP chords | 25 |
| Aspect ratios ≥ this are flattened | 50. |
| Diameters > this not flattened | 0.002 |
| Default capping radius | 1. |
| Flux bounds ≥ this are taken as artificial | 1000. |

| | Constraints | Variables | Ray Yield |
|---|---|---|---|
| Stoichio, objective and range constraints | 1281 | 469 | 0 |
| Remove artificial bounds, split reversibles | 1127 | 645 | 0 |
| Fix 271 fluxes, revert reversibles | 368 | 198 | 0 |
| Apply mass balance and fixed objective value | 0 | 12 | 0 |
| Apply nontrivial range constraints | 33 | 12 | 0 |
| Remove prismatic rays | 30 | 9 | 3 |
| Linealities yielding ray pairs | 30 | 8 | 2 |
| RSS after removing redundant constraints | 14 | 8 | 0 |
| RSS is closed, no capping done | 14 | 8 | 0 |

{All points in the solution space share the objective value 2.93556

All 8 chords were calculated by LP for the SSK.

The maximal inscribed hypersphere diameter is 0.0771845

The diameter and volume coverage ratios, between mutually similar simplices that encloses the periphery points or the complete SSK, are {79.7, 16.3}% respectively.

The mean SSK diameter is estimated to be in the range {2.33531, 4.55858} and the best value estimate is 2.92861

The sampled fraction of the SSK spanned by the peripheral point polytope, is in the 95% confidence interval {21, 26}%

The following reactions acquired fixed directions:

{EX_h2o2_e, backwards} {EX_nh4_e, forwards}

62 peripheral points were found.

Assuming these to be representative, and combining with rays, extends the fixed value list to 408 items

The combined set of 4 rays span 4 of the total 4 ray dimensions.

VALIDATION TEST: Deconstruct FBA solutions (with/without artificial bounds) into the sum of a Kernel space flux, and a flux along a ray direction. Agreement between actual and reconstituted solutions are indicated by % discrepancy of total flux, and angle in degrees between their directions in flux space.

| FULL SS KERNEL | | Flux vector length | % Flux mismatch | Misalignment angle deg |
|---|---|---|---|---|
| | Not bounded | 21.5 | 0. | 0. |
| | Art. bounded | 1732.1 | 0. | 0. |

,

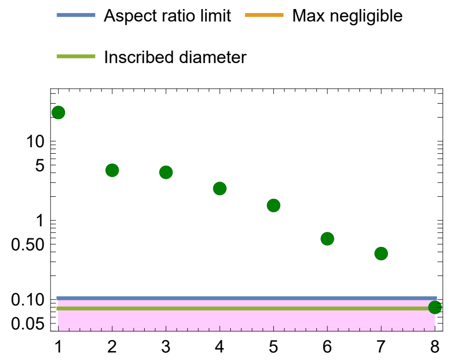


FLATTENING DECISION PLOT

Green/Blue – Kept chord/diameter;
Red/Purple – Flattened chord/diameter.

MAIN ORTHOGONAL CHORD LENGTHS

Cyan chords, magenta diameters

Blue line is max inscribed sphere diameter

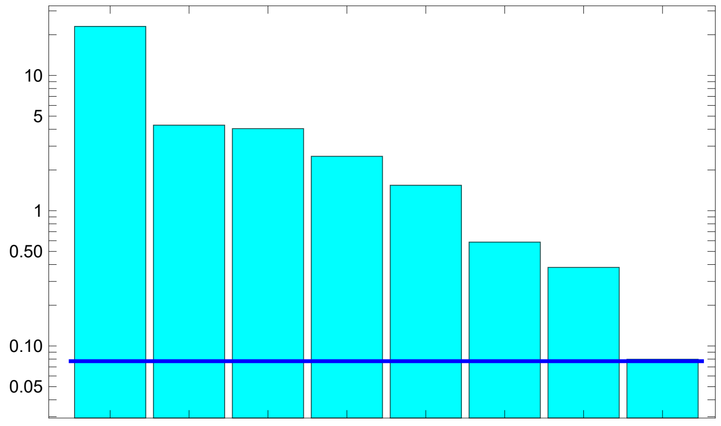

The mean and maximal aspect ratio's are 8.0 and 288.8

SPLIT DIAMETERS ALONG MAIN CHORD DIRECTIONS

Asymmetric overhang shown in light shading.

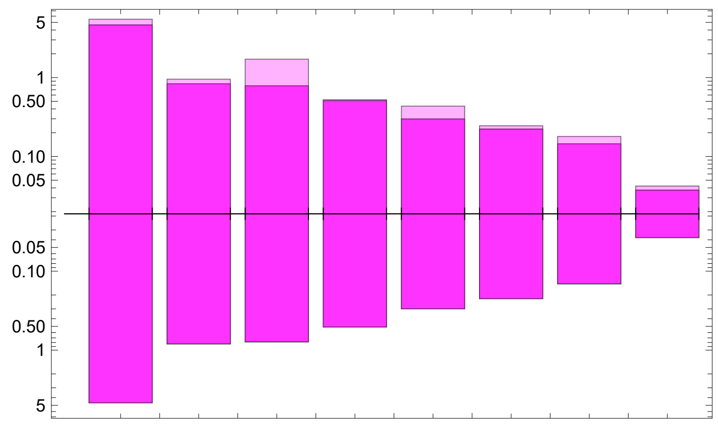

Average periphery point non–centrality 13.5%

}
