## Supplementary material for "The FBA Solution Space Kernel – Introduction and Illustrative Examples": SSK software manual and code: Toy Network SSKernelReport.pdf

### Full SS Kernel Analysis for SolutionSpaceKernel

Model Toy network with uncapped kernel of type SimpleCone
calculated on Tue 19 Sep 2023 in 0.313 seconds.

| | |
|---|---|
| Step time limit | 60. |
| Unbuffered externals exempted | 0 |
| LP tolerance | 0.0001 |
| Fixed value tolerance | 0.002 |
| Progenitor sample size | 20 |
| Max BFBF tree nodes | 100 000 |
| BFBF random greedy sample size | 500 |
| Gready search mixing fraction | 0.8 |

| | |
|---|---|
| Stop sampling when failure rate > | 20 |
| Minimal, Maximal LP chord counts | {10, 50} |
| Maximal flips to find LP chords | 25 |
| Aspect ratios ≥ this are flattened | 50. |
| Diameters > this not flattened | 0.002 |
| Default capping radius | 1. |
| Flux bounds ≥ this are taken as artificial | 100. |

| | Constraints | Variables | Ray Yield |
|---|---|---|---|
| Stoichio, objective and range constraints | 26 | 9 | 0 |
| Remove artificial bounds, split reversibles | 17 | 9 | 0 |
| Fix 0 fluxes, revert reversibles | 15 | 9 | 0 |
| Apply mass balance and fixed objective value | 0 | 3 | 0 |
| Apply nontrivial range constraints | 6 | 3 | 0 |
| RSS after removing redundant constraints | 3 | 3 | 0 |
| RSS a simple cone, capped with default radius | 3 | 3 | 0 |
| Default capping of 3 progenitor rays | 7 | 3 | 3 |

All points in the solution space share the objective value 0.0
All 3 chords were calculated by LP for the SSK.
The maximal inscribed hypersphere diameter is 0.270358
The diameter and volume coverage ratios, between mutually similar simplices
that encloses the periphery points or the complete SSK, are {99.2, 97.6}% respectively.
The mean SSK diameter is estimated to be in the range {0.461784, 0.659141}
and the best value estimate is 0.465581
The sampled fraction of the SSK spanned by the
peripheral point polytope, is in the 95% confidence interval {95, 100}%
32 peripheral points were found.
Assuming these to be representative, and combining with rays, extends
the fixed value list to 0 items
The combined set of 3 rays span 3 of the total 3 ray dimensions.

VALIDATION TEST: Deconstruct FBA solutions (with/without artificial bounds)
into the sum of a Kernel space flux, and a flux along a ray direction.
Agreement between actual and reconstituted solutions are indicated by % discrepancy
of total flux, and angle in degrees between their directions in flux space.

| | | Flux vector length | % Flux mismatch | Misalignment angle deg |
|---|---|---|---|---|
| **FULL SS KERNEL** | Not bounded | 0. | Zero | Indeterminate |
| | Art. bounded | 0. | Zero | Indeterminate |

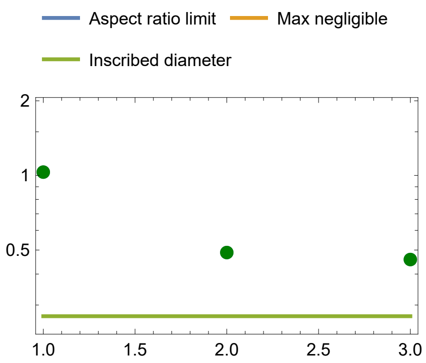

#### FLATTENING DECISION PLOT

Green/Blue – Kept chord/diameter;
Red/Purple – Flattened chord/diameter.

#### MAIN ORTHOGONAL CHORD LENGTHS

Cyan chords, magenta diameters

Blue line is max inscribed sphere diameter

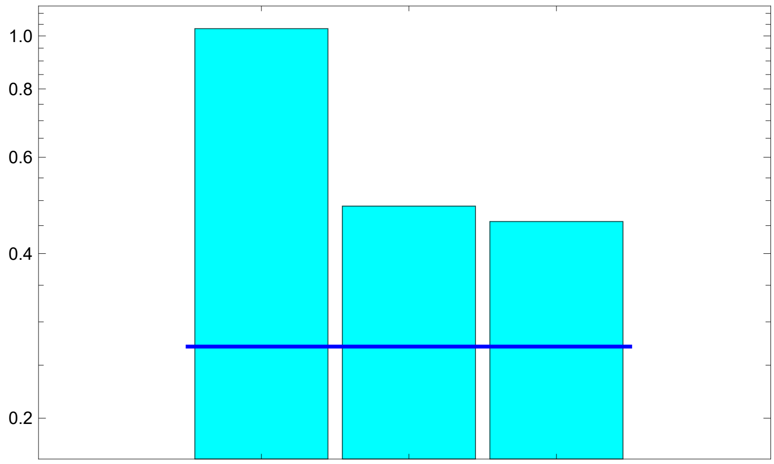

The mean and maximal aspect ratio's are 1.7 and 2.3

#### SPLIT DIAMETERS ALONG MAIN CHORD DIRECTIONS

Asymmetric overhang shown in light shading.

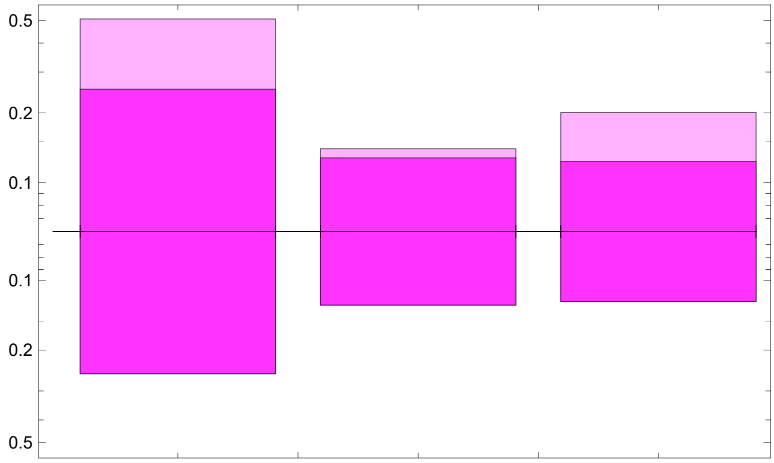

Average periphery point non–centrality 11.0%
