## Supplementary material for "The FBA Solution Space Kernel – Introduction and Illustrative Examples": SSK software manual and code: USER GUIDE.pdf

### SSKernel V 1.4

### June 2024

*Copyright: Wynand Verwoerd*

Centre for Advanced Computational Solutions
Dept WF & Molecular Bioscience
Faculty of Agriculture & Life Sciences
Lincoln University
Ellesmere Junction Road
CHRISTCHURCH 7647
New Zealand

### 1 Copyright

Copyright © 2022 Wynand S. Verwoerd

This file is part of SSKernel.

The SSKernel program is free software; you can redistribute it and/or modify it under the terms of the GNU General Public License as published by the Free Software Foundation; either version 3 of the License, or (at your option) any later version.

This program is distributed in the hope that it will be useful, but WITHOUT ANY WARRANTY; without even the implied warranty of MERCHANTABILITY or FITNESS FOR A PARTICULAR PURPOSE. See the GNU General Public License for more details.

You should have received a copy of the GNU General Public License along with this program. If not, see http://www.gnu.org/licenses/.

The program may be freely used in accordance with the GPL but it is required that in any publication of results based on its use, the following reference should be cited:

W. S. Verwoerd, *The Compact Kernel of a Metabolic Flux Balance Solution Space*, World Scientific Publishing Co., Singapore, 2022; ISBN 978-981-12-5583-0, doi:10.1142/12821 .

### 2 System requirements

The notebook file *SSKernel Launcher.nb* runs on *Mathematica* version 12 or higher; it may also work on earlier versions. *Mathematica* is obtainable from Wolfram Research, Inc, for various operating system environments. Because of its notebook implementation the *SSKernel* program can be run on any operating system for which *Mathematica* is available.

In addition, a Wolfram Language Script file *SSKernelscript.wls* is supplied. This is essentially a command-line version of *SSKernel*. It sacrifices the interactive functionality available in the notebook version, but only requires access to the Wolfram Engine, either on a network or cloud server (such as Wolfram Cloud, https://www.wolfram.com/cloud/) or a locally installed engine. To run the script file, the *wolframscript* interpreter is also required.

Wolfram allows free registration and subsequent installation of the Wolfram Engine for the purpose of software development and open code projects, including the *wolframscript* interpreter. See https://www.wolfram.com/engine/ for more details.

### 3 Program Concepts

The purpose of the *SSKernel* program is to facilitate the further characterization and comparison of constraint-based steady state models of metabolic networks, once a flux balance analysis (FBA) of such a models has been performed.

This section gives a brief synopsis of how this works; the full explanation, with details of the many algorithms needed for this task, is found in the book *The Compact Kernel of a Metabolic Flux Balance Solution Space* referred to in the copyright section above.

A common feature of FBA models is that they are underdetermined. Even after the optimization of one or more objectives that are meant to broadly represent the effects of cellular metabolic regulation, metabolic fluxes are not uniquely determined but only confined to lie in a subspace of the high-dimensional flux space. This subspace is denoted as the **solution space** (SS) and may itself still have dimensions in the order of 100's or 1000's. Since FBA constraints and objectives are normally linear in terms of fluxes, the SS is in principle a polytope, but may be unbounded in some directions and so may not have a finite volume. To avoid this, artificial maximum flux constraints are routinely introduced into FBA models, leading to a finite (closed) polytope for the SS of such a model. Such artificial termination of the SS is discarded in the SSKernel analysis.

An FBA calculation using linear programming, gives only a single point (often a vertex) of the SS polytope. But the polytope is mathematically fully described by the set of vectors in flux space that specify all its vertices, since they form a convex basis. That is the idea underlying extreme flux or elementary mode analysis[1], which essentially specify these vertices. However, this is impractical for a large dimensional polytope, because the number of vertices tends to increase exponentially with the number of dimensions. An extension of FBA, called flux variability analysis (FVA), can give some idea of the extent of the SS. This is compared in section 7.1 to the SS characterization achieved here, for a simple network example.

The premise of the analysis performed by *SSKernel*, is that even for large networks it is nevertheless possible to extract a much smaller dimensional subspace, which contains the biochemically most significant part of the flux variation that the model allows. This is called the **solution space kernel** (SSK) and is a closed polytope, even where the SS is not fully bounded.

The approach taken, allows consideration of the SSK as a geometric object in N dimensions, with a particular shape that can be characterized even if not visualized. An aspect of this is that various directions in flux space are found to have special

---

[1] Papin, J., Stelling, J., Price, N., et al. (2004) Comparison of network-based pathway analysis methods. Trends Biotechnol, 22, 400–405. doi:10.1016/j.tibtech.2004.06.0101;
Schilling, C. H., Letscher, D., & Palsson, B. O. (2000) Theory for the systemic definition of metabolic pathways and their use in interpreting metabolic function from a pathway-oriented perspective. Journal of Theoretical Biology, 203, 229–248. doi:10.1006/jtbi.2000.1073;
Schuster, S., Dandekar, T., & Fell, D. A. (1999) Detection of elementary flux modes in biochemical networks: A promising tool for pathway analysis and metabolic engineering. *Trends in Biotechnology*. Elsevier Ltd. doi:10.1016/S0167-7799(98)01290-6.

meaning. To connect the idea of a flux space direction with biochemical concepts, consider that a specific direction defines the relative sizes of all flux components. Different flux vectors along the same direction signifies differences in the total flux carried, while the proportions of fluxes in individual reactions remain the same.

For example, a conventional biochemical pathway involves only a small selection of reactions in the metabolic network, so would have all other flux components zero. This means geometrically that it corresponds to a direction located in a hyperplane that is orthogonal to all such flux axes. Moreover, inside this hyperplane, there will generally only be a single direction for which all the non-zero flux components have the correct mutual ratio's dictated by e.g. the stoichiometry of the participating reactions. So in this case there is a direct correspondence between this direction and a particular pathway. A metabolic state where this pathway carries a particular flux, will correspond to a vector along this direction with an appropriate length.

Generalizing this, a random direction in flux space might involve many or even all reactions of the network, and if it lies in the feasible region that is compatible with stoichiometry and other model constraints, might be considered as a metabolic "mode" composed of many participating biochemical pathways. Various points along this direction corresponds to metabolic states that all belong to the same mode, but carrying different overall fluxes. The set of model constraints may in some cases be compatible with only a single point along this direction, in other cases with a finite or even infinite range of flux points that lie along the direction.

The boundaries of the SS polytope are formed by a set of hyperplanes. These are mathematically specified by a **constraint matrix** and a **constraint values vector**. The coordinate origin is required to be inside or on the polytope boundary. Then each row of the constraint matrix is a unit vector defining a direction in flux space. The corresponding boundary hyperplane is orthogonal to this direction, and intersects it at a radius given by the corresponding entry in the values vector. With an interior origin, entries in the values vector are nonnegative.

Figure 1 illustrates the boundary representation for a triangle in 2D. The boundary hyperplane representation of a polytope is an equivalent alternative to specifying all the vertices, but has the major advantage that for a space of dimension N, the number of hyperplanes is generally only of the order of N (the minimum number required to get a closed polytope, is the (N+1) needed for a simplex). The disadvantage is that many geometrical quantities are much harder to calculate than for the vertex based specification.

The approach used by *SSKernel*, consistently uses the hyperplane representation and avoids ever calculating vertices. This is a major distinction from elementary mode based descriptions of the SS.

The part of a boundary hyperplane that is **feasible** (i.e., satisfies all constraint matrix constraints with a less or equal condition) is termed a **facet** of the SS polytope. Similarly, intersections of facets are themselves facets, at a **facet level** defined by the number of intersecting hyperplanes. Thus, for a polyhedron in 3 dimensions, its sides are the level 1 facets, edges are level 2 facets, and vertices level 3. Generally, facet levels range from 0 (the SS itself) to the dimension number N, and level N facets are the vertices.

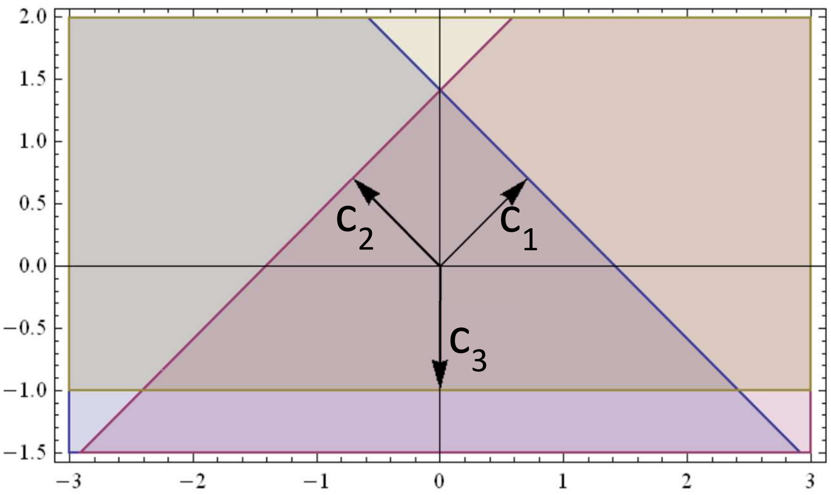

$$\begin{bmatrix} c_1 \\ c_2 \\ c_3 \end{bmatrix} \begin{pmatrix} x \\ y \end{pmatrix} = \begin{bmatrix} .707 & .707 \\ -.707 & .707 \\ 0.000 & -1.00 \end{bmatrix} . \begin{pmatrix} x \\ y \end{pmatrix} \leq \begin{pmatrix} 1 \\ 1 \\ 1 \end{pmatrix} \quad (1.1)$$

*Figure 1: 2D example of representing a polytope by boundary planes specified by a constraint matrix and values vector. The first row of the constraints matrix contains unit vector $c_1$ and the grey region in the figure is the set of points p = (x,y) for which $c_1.p \leq 1$, the corresponding entry in the values vector. The polytope, here a triangle, is the area where the three half-planes defined by Equation 1.1 overlap.*

Having removed artificial flux constraints the SS is almost always unbounded. In this case at least some of its facets will also be unbounded. Any direction along which the SS is unbounded, is called a **ray** direction. A ray that falls within a facet, defining the direction along which that facet is unbounded, is considered to be a **peripheral ray** of the SS and the higher the facet level, the “more peripheral” the ray would be.

As a counterpart to ray directions, which signify an infinite range of flux variation, it is commonly observed (e.g. by doing flux variability analysis (FVA) ) that many fluxes in a FBA model have a constant value throughout the solution space. These are designated as **fixed fluxes**. The *SSKernel* program detects these by a special algorithm that runs faster than FVA by several orders of magnitude, and isolating them at the start is a key step to reduce SSK dimensions and make the finding of rays and facets computationally tractable.

To this end, the calculation in the program is divided up in different stages further discussed below. Crucially, in an initial stage the fixed fluxes and an easily calculated class of rays (prismatic rays) are removed and stoichiometry constraints applied which results in a **reduced solution space** or RSS. The RSS dimension count is typically between 1% and 10% of the value for the full flux space and is determined without requiring heavy computing, making it a worthwhile intermediate step.

It is possible that all facets at all levels, except for a single vertex where they all intercept, are unbounded; in that case the SS shape is a **simple cone** and its SSK is trivially just the single vertex point forming the apex of the cone.

More commonly, the SS has both unbounded and **bounded facets** and is then designated as a **faceted cone**. A bounded facet is mostly referred to as an **FBF** (Feasible Bounded Facet) to emphasize that it needs to satisfy both feasibility and boundedness conditions. Typically, all facets at level 1 and other low levels remain unbounded, but FBF's appear at higher levels. Figure 2 illustrates the relationship between a faceted cone SS in 3D and its RSS and a prismatic ray vector.

The importance of bounded facets are that they express the ways in which the diversity of physical, biochemical and biological flux constraints interact with each other to limit the feasible flux values to finite ranges. That is in contrast to the artificial flux range constraints that express no other reality than the mere observation that fluxes cannot be infinite.

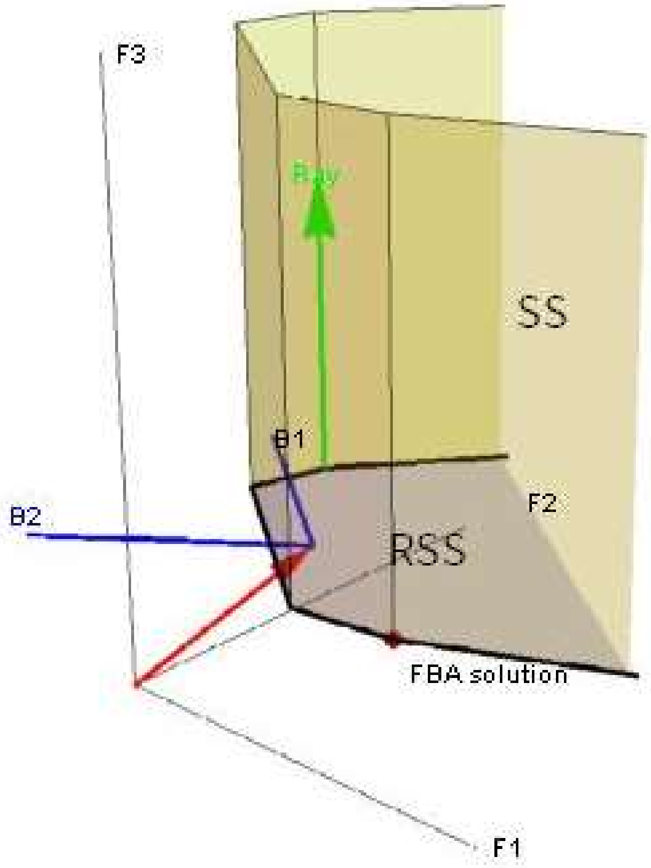

*Figure 2: Schematic representation of a faceted cone solution space (SS, light green) in a 3D space composed of flux vectors F1, F2 and F3. The FBA solution is a vertex of the SS. All level 1 facets (the sides) of the SS are unbounded, but some level 2 facets (the black edges at the bottom) are bounded. One ray direction (green arrow) which is a (level 2) prismatic ray is shown, and there are also conical rays as shown below in Figure 3 .*

*Projecting out the prismatic ray leaves a 2D reduced solution space (RSS, light blue). Its location and orientation is given by the coordinates of the RSS origin (red arrow) , and the RSS basis vectors (blue arrows B1 and B2), as vectors in the 3D embedding space.*

So it is plausible to distinguish between the part of the SS that is bounded by enforcing meaningful constraints, and the part that allows infinite fluxes due to the absence of such meaningful constraints. The key is to introduce additional **capping constraints** to perform this separation. The capping constraints are carefully constructed not to constrain finite fluxes any further than the model constraints already do. To achieve that, each capping constraint hyperplane is set at the minimal radius that ensures it does not intersect any FBF at more than a single point, i.e. tangentially. This is designated as its **capping radius**.

The resulting finite region, bounded by all FBF's that belong to the SS, and the capping hyperplanes, is the sought after **Solution Space Kernel** or SSK and is the central concept in this work. Figure 3 shows the SSK for the RSS described in Figure 2, as well as its associated rays and capping constraint.

It is emphasised that this is not meant as a truncation of the SS (as artificial flux limits do) but merely as a partitioning. Any feasible flux beyond the SSK can still be recovered by adding a suitable multiple of a ray vector to a flux that falls within the SSK. A **ray vector** is defined as a unit vector that defines a direction in flux space for which the allowable flux is not bounded by the model constraints.

For this reason the specification of the SSK is supplemented by giving a set of ray vectors. These ray vectors are not necessarily mutually orthogonal, but form a complete (possibly overcomplete) basis for a subspace designated as the ray space of the FBA model. The **ray space** is a subspace of the flux space that contains every ray of the SS, but not every flux vector in the ray space is necessarily a ray. In other words, every ray is a linear combination of the ray basis vectors, but not every linear combination is a ray. Conversely, if the linear combinations are restricted to **convex** combinations, every convex ray basis combination is a ray, but some rays may not be representable as convex combinations of the computed ray vectors. In order to minimize the range of rays that are missed by making convex combinations, without allowing the size of the basis set to increase much beyond the number of dimensions, the computational procedure used by *SSKernel* prioritizes maximally **peripheral rays** for the ray basis, by maximizing the level of the facet that contains each of them.

Two major (and interrelated) computational tasks performed by the *SSKernel* program is to find all FBF's and the ray basis (which is internally manipulated as part of a **ray matrix**). There are a few different types of rays, found by different algorithms, namely **prismatic** rays, **linealities**, **singleton** rays, and **conical** rays. In the first 3 cases the required capping is of a type called **coincidence capping**, which in fact reduces to a vector projection and makes a major contribution to reducing the dimensionality of the SSK. For conical rays, coincidence capping is also possible in some situations but otherwise **tangential capping** that does not reduce dimensions is applied.

It is obvious that the subfacets of an FBF are also feasible and bounded, whereas unbounded facets may have both bounded and unbounded subfacets. Figure 2 shows some examples of both cases.

A tree representation is used to capture such relationships. For a given FBF, the lowest level ancestor that is also an FBF, is denoted as the **base feasible bounded facet** or **BFBF**. It is enough to know just the set of all BFBF's, because all remaining FBF's are by definition subfacets of them.

Furthermore, there is always an unbounded facet that is an ancestor of all BFBF's. The level 0 facet (i.e., the polytope itself) is always such a common ancestor, as is the case in Figure 3. But in higher dimensions there are often closer ancestors located at much higher levels. The highest level such common ancestor is denoted as the **progenitor facet**. So as a rather trivial example, in Figure 2 the facet coloured light blue and marked "RSS" is the progenitor facet as it contains all BFBF's as subfacets.

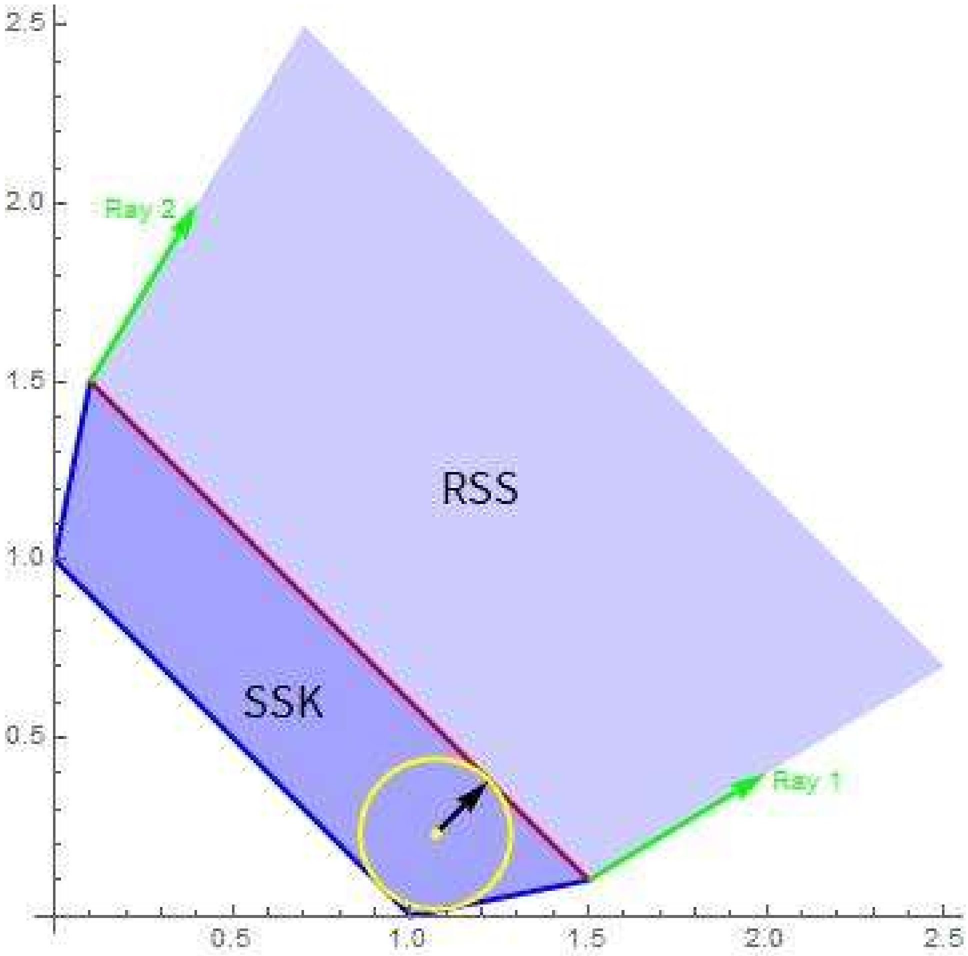

*Figure 3: Reducing the 2D RSS (light blue) to a bounded SSK (darker blue) by tangent capping. The two green arrows represent peripheral conical rays which are listed in the ray matrix. The 2D plane of the figure is the ray space and is spanned by the two ray vectors shown as green arrows. In this simple example they also happen to form a convex basis (i.e., all rays are convex combinations of Ray 1 and Ray 2). Note that if e.g. Ray 2 was replaced with a non-peripheral ray, such as the diagonal y = x in the figure, the resulting set would still be a complete basis, but not a convex basis any more.*

*There are three FBF's, the blue lines, and as they are not subfacets of any lower level facet (except the RSS, level 0) they are also the BFBF's. The red line signifies a capping constraint defined by its capping vector (black arrow). Generally, two capping vectors, one along each ray direction, would be used. But here, only a single capping constraint is used to simplify the illustration. The capping radius is set at the minimal value such that the capping line does not intersect any BFBF at more than a single point (i.e., it is tangent to the SSK). In so doing the two RSS constraints passing through the tangent points become redundant for the SSK and are eliminated from it.*

*The yellow circle is a maximal radius inscribed circle, and is not unique. Its centre at the yellow dot is well interior and so is suitable as the coordinate origin relative to which all constraint lines are specified. The inscribed radius happens to coincide with the capping radius in this particular example.*

The significance of the progenitor is that the multinomial tree search for BFBF's can be restricted to the subtree with the progenitor as the root node rather than the full SS polytope. And furthermore, the solution space can be coincidence capped to the progenitor (provided that there exists a ray that is orthogonal to the progenitor). This can produce a significant dimension reduction and eliminate all rays that are orthogonal to the progenitor facet.

To make use of these observations, a preliminary stage in finding BFBF's is to determine the progenitor facet, and then perform coincidence capping to it if possible, so that the full BFBF search can be more efficiently done in a lower dimensional space.

For each peripheral ray vector that remains in the progenitor facet, its direction is used to define a tangential capping constraint vector and the capping radius determined by an LP minimization. Constraints that become redundant as a result of capping are also identified and eliminated. Figure 3 illustrates this.

As the calculation progresses through various dimension reduction steps, it is important to shift the coordinate origin inside the reduced polytope after each step, and this is referred to as **centering**. LP methods that include the feasibility constraints can do this efficiently but often produce a point located on the polytope periphery, usually on a high level facet. A useful device to find a point that is truly interior is to find the **maximal inscribed hypersphere**. This is also done by formulating an appropriate LP problem. The centre of this sphere is well interior, but not unique and often not well centred in the sense of being at an equal distance (as far as possible) from all sides of the polytope. An example of that appears in Figure 3. An additional step called **center refining** is implemented to improve on that. It is based on shifting the centre to minimize the sum of pairwise percentage differences between the forwards and backwards radii to the periphery, over a number of suitably (see below) chosen sampling directions. There may be multiple equivalent refined centre points.

Exact formal specification of the final SSK polytope in the program output, is presented as a constraint matrix and values vector that is usually several orders of magnitude smaller than the stoichiometry matrix of the FBA model. To connect the low dimensional subspace in which the SSK constraints are formulated, to the physically meaningful flux space, a vector transformation is also supplied. This transformation consists of the position of the SSK coordinate origin in flux space, and a set of mutually orthogonal vectors in flux space that defines the basis in which the SSK constraint vectors are expressed (See Figure 2 ).
This information is encapsulated in a special data structure in the form of a nested list, written as in standard *Mathematica* notation with curly brackets to denote comma delimited sublists as:

$$\begin{aligned} kernelspace &= \{polytope, transformation\} \\ &= \{\{constraints, values\}, \{SSKorigin, SSKbasis\}\} \end{aligned} \tag{1.2}$$

Both the SSK origin and basis are chosen by *SSKernel* to have special significance.

The *origin* is chosen to be a refined centre of the closed, low dimensional SSK polytope. For a SSK of N dimensions, of the order of 2N sampling directions are chosen to do the refining, in a way that aims to distribute them uniformly over the (N-1)-dimensional angle space but also to include the endpoints of the main polytope chords. The set of approximately 4N **peripheral points** generated by this centralization procedure also serve to further characterize the SSK polytope.

The maximal chord of a polytope could in principle be calculated by a mixed integer linear programming procedure, but this is tractable only for single digit values of N. Instead, a special heuristic is used to select the longest chord among candidates from a number of randomized LP-based maximizations. To express the distinction, this is called a **main chord** and can be considered an approximation to the maximal chord. A second main chord is found similarly but constrained to be orthogonal to the first, and likewise subsequent chords until a full set of N mutually orthogonal main chords are known. The result is somewhat analogous to finding principal components for a set of multidimensional data points. But note that the main chords are found without reference to the coordinate origin and will not have a common intersection except if the SSK happens to have a highly symmetric geometry.

The SSK *basis* is then taken as the normalised vectors along the main chord directions, taken in descending order of chord lengths. Note that the SSK **diameters** along the SSK basis vector directions are generally considerably smaller than the main chords, because they by definition are constrained to all intersect at the SSK origin.

For large values of N, even calculating all the main chords can become too time consuming. In these cases only the first few are calculated by LP, and the rest estimated from diameters passing through the set of peripheral points.

The set of mutually orthogonal main chord lengths gives a broad indication of the geometric shape of the SSK. Dividing the longer length by the shorter, for each pair of chords, defines an **aspect ratio** of the SSK polytope as seen along a direction perpendicular to the plane that the pair defines. For a highly symmetric shape many (or all, for a hypersphere) aspect ratios may be equal. But in practice typical FBA models show a declining series of main chord lengths and some aspect ratios can become high, 2- or 3-digit numbers. For context, the aspect ratio associated with the thickness of a credit card is in the range between 50 and 100.

When the aspect ratio defined by a chord pair is larger than a chosen threshold, the direction of the short chord is designated as a **thin direction**. Such a direction represents a combination of fluxes that remains approximately constant throughout the SSK, in comparison with the range of variation found along other directions. This is an extension of the concept of a fixed single flux value that represents zero thickness along the relevant flux axis. To treat them similarly, *SSKernel* can optionally flatten the SSK to zero thickness along all thin directions, with a corresponding reduction in the SSK dimension count. The maximal flattening threshold can be set interactively by the *SSKernel* user.

In addition to the formal specification of the SSK by the SSKernel data structure of equation (1.2), a more intuitive depiction results as a byproduct of the centering procedure described above.

Firstly, the SSK origin defines a single **typical flux**. This is typical in a sense that generalises the concept of the median of a 1-dimensional set of data, to multiple

dimensions. By construction, there are (as closely as is allowed by the generally irregular shape of the SSK) equal ranges of feasible flux values to either side along any direction in the SSK subspace.
That is in contrast to the FBA flux, which (depending on the LP solver used) is mostly at an extreme such as a vertex of the polytope and so highly atypical.
The origin is merely *a* typical point: even in 2D, an asymmetric triangle has no uniquely defined centre point.

Secondly, the set of peripheral points that are delivered by centre refining, are by construction distributed more or less evenly in all directions around the central point. Each of them defines the maximal deviation from the central point allowed by SS constraints, along its particular direction. So the peripheral points define a broader **central region** of the SSK, and delineate a typical range of variation of fluxes. In the process they indicate the shape of the SSK, somewhat extending the level of detail represented by the main chords.. Also, they can serve as a **representative sample** of flux values that are compatible with the optimized objective and the model constraints, e.g. to calculate average values for flux-dependent quantities.

Another perspective is that since any convex combination of the peripheral point vectors is also feasible, they can be considered to form the vertices of a polytope that defines such a central feasible region, the **peripheral point polytope** or PPP. Of course the SSK is itself a polytope, but its vertices are unknown and there may be an unmanageable number of them if the SSK dimension N is large. By contrast, the PPP has only a number of vertices of the order of 4N. It could perhaps be used as a ”poor man's version” of elementary flux modes that underly this central typical region.

In order for this to be useful, one would need to gauge if the central region covers an appreciable fraction of the full SSK. A tricky point in such a comparison, is that the power law dependence of  hypervolume on dimensionality, means that e.g. even the inscribed hypersphere of a hypercube, only covers a negligible fraction of its volume. So it makes more sense to compare a measure of the radius of the PPP to the corresponding radius of the SSK.

For low dimensions, *SSKernel* makes a Monte Carlo estimate of this by direct sampling. In higher dimensions, it instead compares the average inscribed hypersphere radii of similarly oriented, regular simplexes that respectively encloses the PPP and the full SSK polytope. The values are averaged over multiple random orientations of the simplexes. The radius ratio usually comes out at between 60% and 90%, even where the power law causes the volume ratio to be become negligible in high dimensions.

An example that shows calculated periphery points and the enclosing simplexes is shown in Figure 4. Also, section 8.1 illustrates a number of SSK concepts for a simple toy network example.

The various steps outlined above require a considerable number of new algorithms, and many of those involve random sampling. In some cases a more exact algorithm such as tree traversal is used for smaller dimensions, but an approximate method based on sampling is needed to make the calculation tractable for large dimensions. One consequence is that some intermediate results, such as the number of BFBF's found, can vary somewhat in repeated runs, although e.g. in that case the actual capping radii derived from the BFBF search is found anecdotally to remain virtually unchanged.

Nevertheless, it is expedient to **validate** the final SSK independently of its calculation. This is achieved by a special LP-based **deconstruction** procedure that only uses the SSK data structure to decompose a given flux vector into its SSK and ray vector parts, and any discrepancy found is assigned as a residual vector.

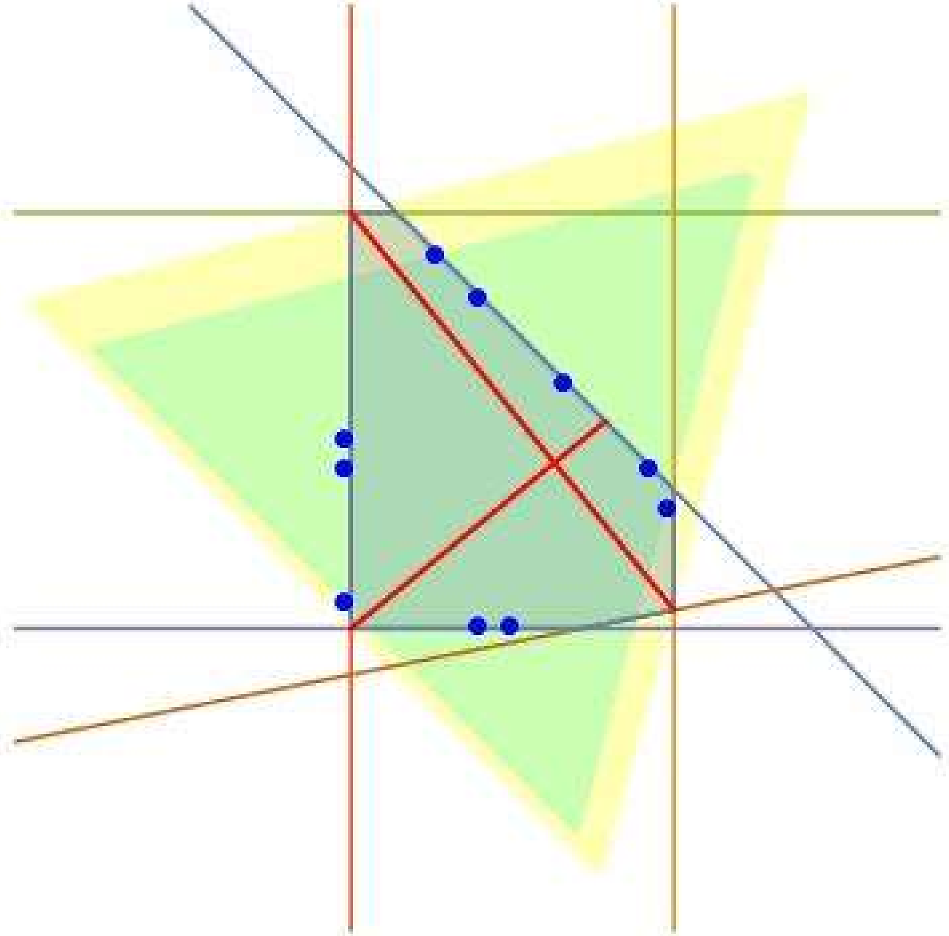

*Figure 4: 2D example of peripheral points, main chords and enclosing simplexes.*

*The coloured lines represent a set of constraint lines that define an irregular polygon shown as the shaded region.. The red lines are the calculated main chords, giving an aspect ratio of 1.56. The blue dots are a set of semi-random periphery points and define a central region of the polygon, the PPP.*

*The regular simplex for a 2D space is an equilateral triangle, The yellow and green equilateral triangles shown are respectively the calculated enclosing simplex of the polygon and of the PPP. The enclosing simplexes have the same randomly chosen orientation. In the SSKernel calculation, the mean radius of the inscribed circle of each triangle, taken over multiple random orientations, is taken as a measure of the spatial extent of the polygon and its PPP, in order to express the relative coverage of the polygon by the PPP. In this example, the radius ratio is 0.83 giving an area ratio of 0.7.*

A basic premise of the SSKernel project is that a residual-free deconstruction is possible for any feasible flux. So if a residual vector component remains, its norm indicates the extent to which the calculation has fulfilled the premise. This test as applied to periphery points, is used to control the flattening procedure. Almost invariably, the residual is zero or negligible for a SSK constructed without flattening, but has a noticeable value when flattening is applied and may indicate the need to flatten fewer thin directions.

In the final stage of the calculation, the validation test is applied to the two FBA solutions (with/without artificial flux bounds) that were calculated initially independently of the SSK. In this case, the accuracy of the deconstruction is assessed by noting by how much the deconstructed flux vector deviates from the original FBA vector in terms of direction and relative magnitude.

### 4 The User Interface and Workflow

The *SSKernel* software can be run in two distinct environments. The recommended version described here is to load and execute the notebook file *SSKernel Launcher.nb* in *Mathematica*. This provides by far the most user friendly, flexible and informative way to do a SSKernel analysis. It provides a fully interactive interface that gives dynamic feedback and allows user intervention accordingly.

The alternative command line implementation is discussed in section 6.

The user interface consists of a **control window** and a **status report window**.

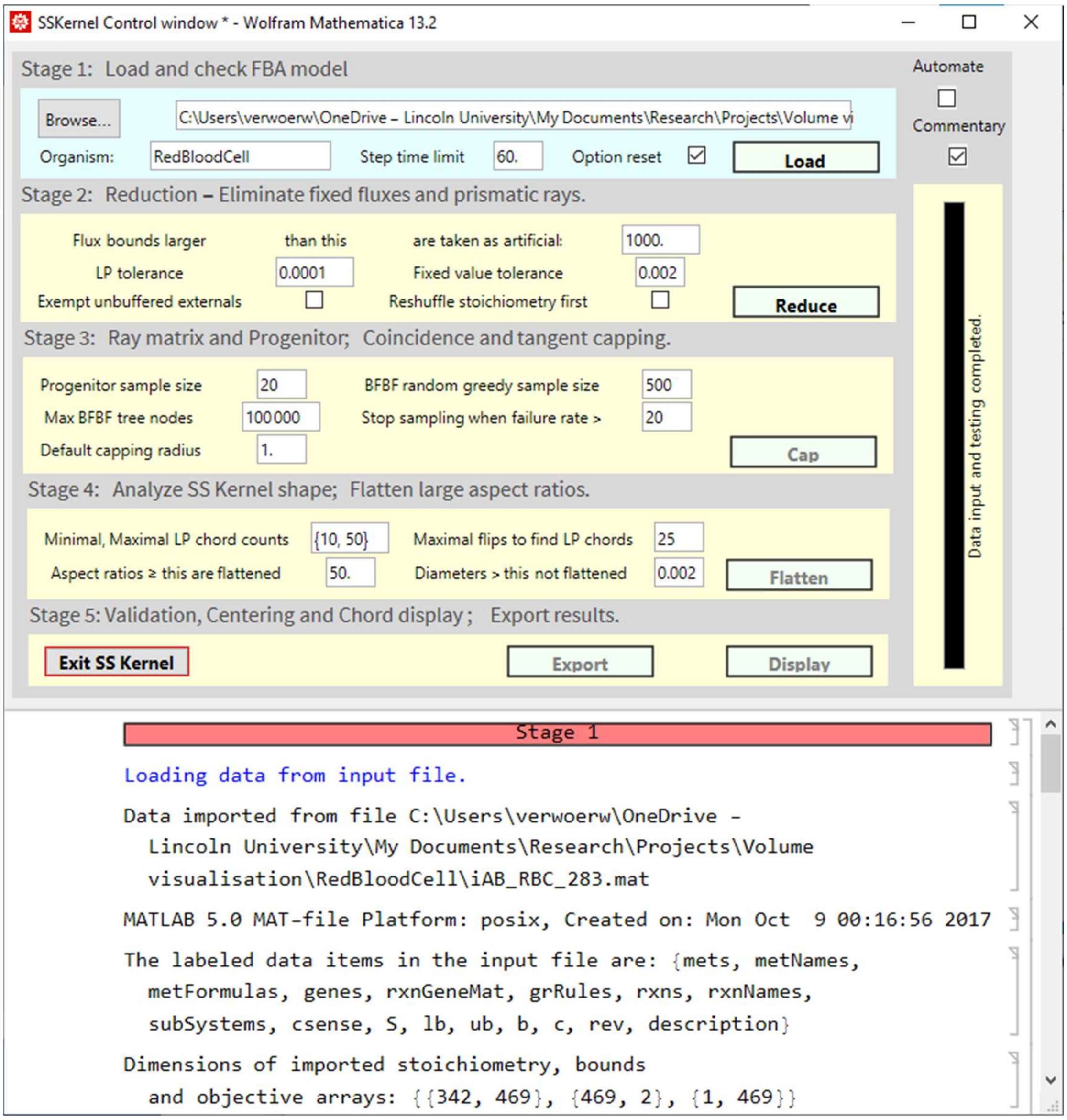

*Figure 5: Example of the SSKernel control window.*

The software is supplied as a .zip file that unpacks into a directory named SSKERNEL, containing various files and the Packages directory where most of the program code resides.

The program is launched by simply double clicking on the SSKernel Launcher.nb icon, or e.g. a shortcut for it installed on the desktop. This first loads *Mathematica*, and displays the notebook with a warning that it contains dynamic content. Also, with the default configuration of the *Mathematica* frontend it will present a dialog offering to evaluate the initialization cells in the notebook. If so, ignore the warning and just click “OK” to proceed with executing it automatically. This step can be skipped by changing the preference “InitializationCellWarning” to FALSE in the *Mathematica* front end.

In a first run, use the browse button in the control window to enter the TEST directory within SSKERNEL. It contains a single data file, iAB_RBC_283.mat. Selecting this the control window should appear as above (Figure 5) after loading an FBA model for the red blood cell.

The calculation is divided into 6 discrete stages, each of which is controlled by a distinct frame in the control window into which its relevant control parameters can be entered. Default values for the control parameters are read from a configuration file.

Normally, execution of each stage is initiated manually by clicking the rightmost button in each frame. The stages need to be executed in numerical sequence, but any stage can be executed repeatedly, with the same or new parameter values. This includes going back to a previous stage and repeating the calculation from that point onwards.

The tick box <u>Automate</u> at the top right of the control window also allows the user to choose performing stages 1-5 automatically.

An overview of the calculation status is given by the background colour of the stage frames:

Yellow – not yet executed

Green – currently executing

Cyan – executed successfully

Pink – unsuccessfully executed

More detailed feedback is supplied by the vertical progress bar on the right, which indicates the actual procedure being performed at any time and its expected time to completion.

Furthermore, below the control section is a scrollable “running commentary” window where text and graphics feedback on detailed intermediate results are presented. This is often useful in determining whether to proceed to the next stage and if not, what parameter changes are needed. The <u>Commentary</u> tick box at the top right allows this feedback to be restricted to essentials.

The current state of the calculation is summarised in a verbal report, also displayed on the screen, in the status report window (Figure 6) which for the RBC model appears as follows after loading:

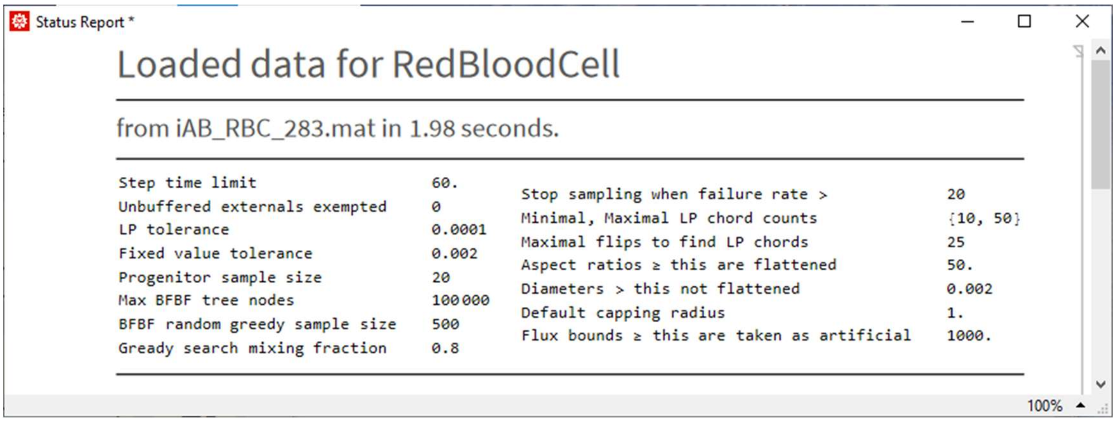

*Figure 6: Example of the SSKernel report window*

Additional entries are added in this window as the calculation progresses, and in addition values such as the heading, running time and control parameter values are dynamically updated.

The final output from the calculation is produced in the form of two output files: the verbal report displayed in the status window, and a data file that contains the SSK specification, a listing of rays, thin directions, periphery points, and the maximal inscribed hypersphere. Export of these two files has to be explicitly initiated by the user.

The program gives some guidance on how to proceed through the stages by greying out steps for which the necessary outputs from previous stages are not yet available. It attempts to intercept any occasional numerical failures and terminate a stage when this occurs. The user is alerted of this by the stage frame turning pink, and standard *Mathematica* error messages are displayed in the running commentary box. The most common culprit of such error is when one of the many LP calculations fails.

It can also happen that the calculation stalls, or the user simply becomes impatient! In this case the stage can be manually terminated by the standard *Mathematica* interrupt <Alt> <,> or abort intervention < Alt > + < . >. This may leave some variables being invalid, so it is advisable to restart from Stage 1 after a manual abort.

Many failures can be remedied by simply repeating the stage. Even without any change of parameter values, the random aspects usually ensure enough variation to the course of the calculation that a previously encountered problem case is avoided. Occasionally it may be necessary to repeat the previous stage to supply a different input to the current stage, or at worst to restart from stage 1.

When a new model is loaded, control parameters are initialized to the values set in the *Configuration.m* package file. Interactive control of the calculation is possible by adjusting the control parameters before each stage as described below. The variable names and default values are listed in Table 2 in section 6.

#### *4.1 Executing Mathematica commands manually*

The lower part of the Control window in which the running commentary appears, is in fact a normal *Mathematica* notebook, into which the user can manually enter and execute commands, delete unwanted cells, etc.

This may be useful particularly in between the execution of individual stages as detailed below, for example to inspect or modify variables in the program. All the variables listed and described in the *Configuration.m* file are accessible and can be displayed or further manipulated.

Displaying variables: For example, after loading a model in Stage 1, the names or ID's of reaction fluxes are not normally displayed, but can be listed by entering the command

```
Multicolumn[ReactionNames, 4]
```

or even better, to interpret flux numbers as associated with the stoichiometry matrix columns and used when specifying rays, thin directions etc., try

```
Multicolumn[Transpose[{Range@Length@ReactionNames,ReactionNames}], 4]
```

Similarly, *MetaboliteNames* can be listed if contained in the input data, even though they are not relevant to the SSKernel calculation. Further examples are found in the case study of section 8.1.

Add an external FBA solution vector: Model specifications usually specifies the objective to be optimized by FBA but not the FBA solution, i.e. a flux vector resulting from the optimization. If available e.g. from the literature or a preexisting FBA calculation using different LP solver software, a comparison may be of interest. SSKernel provides for that, for example in the deconstruction done in stage 5. If the input file is prepared in the *.m format described later, this can be included as *FBAvector*. However, the standard *.mat format does not provide for that. In that case, after stage 1 this can be manually added by entering the following command with applicable numerical values for the $f_i$ :

```
FBAvector = {f1, f2, ……fN};
feasiblepoints = {FBAvector};
```

Modify or add an objective: Another use of this facility would be to modify the objective of a model. The current objective vector is graphically displayed by the control panel, but can be inspected by executing the command

```
objectselector
```

To modify this, enter the desired form by the commands

```
objectselector={s1, s2, ……sN};
FBAdone=False;
```

and execute Stage 2 by clicking the Reduce button. The second line sets a flag to prevent the program from reusing the FBA results from the old objective.

In the case of a model with a trivial zero vector objective or even none defined, it may be necessary to introduce a new objective such as biomass maximization, by adding a corresponding biomass flux as a new column to the stoichiometry matrix and selecting this to be maximised. This column may for example contain the metabolite composition of biomass as obtained from literature.

The code to be executed in the control window before running stage 2, might in this case be

```
ModelName=ModelName<>"+Biomass";
ReactionNames=Join[ReactionNames,{"Biomass"}];
biomass={0,0,0,0,-0.8,-0.2,0};
bounds=Join[bounds,{{0.,999.99}}];
S=Join[S,Transpose@{biomass},2];
{rows,cols}=Dimensions[S];
objectselector=UnitVector[cols,cols];
FBAvector=Join[FBAvector,{0.}];
FBAdone=False;
```

Multiple columns could be added by just adding additional entries in the braces in the various Join statements in the code above and adjusting the objectselector as needed..

Accessing greyed out stages: In the course of a normal calculation, the buttons on the control panel to start subsequent stages remain greyed out until the previous stages have been completed successfully. This ensures that all required data is available. But occasionally, if a stage fails it might be possible to correct this manually or the failure may be non-critical. In such situations, it is possible to override the barrier by manually resetting the variable *available* which takes the form of a list of 6 True/False values. For each, the corresponding stage button is active when set to True. Caution is advised when using this possibility, but it can save the frustration of a stalled calculation.

For extensive manipulations, of which the addition of an objective might be an example, it may not be desired to clutter the Control window output (although, any superfluous cells can simply be deleted when finished with them). In that case an alternative is to open another *Mathematica* notebook, while SSKernel is still running, and perform the further analysis there.

As it is part of the same Mathematica session, all variables, commands and their results executed in the additional notebook are shared by the running control window.

Interrupting or aborting a calculation: For difficult models, the calculation may appear to stall occasionally and require user intervention. It is possible to interrupt the calculation by the button press <alt> comma. Then, in the ensuing dialog the option "Enter subsession" will suspend the SSKernel calculation and allow the user to enter and execute *Mathematica* commands in the control window as above. For example, the command `totalfacets` in an interrupted FBF search will show how many facets have been visited so far, and if excessive some of the control variables might be adjusted to force quicker completion of the random or tree search. Once done, the main calculation is resumed when the subsession is ended by the command `Return[]`. Finally, a running calculation can also be aborted by the standard Mathematica keypress press <alt> fullstop.

#### 4.2 Stage 1

The Browse button allows the user to select an input data file that defines the FBA metabolic model, in either the *Matlab*- oriented .mat format or as a *Mathematica* .m file. More details are given below. By default, it is assumed that the input file is located in a directory named after the biological species of the organism, but any other name can be entered in the Organism box.

To avoid excessive running times, some steps of secondary importance (such as validation tests during flattening, and reorientations of enclosing simplexes) are time limited and this is controlled by the Step time limit box.

Normally, when a new model is loaded, all control parameters are reset to the values read from the configuration file. Avoid this by unticking the Option reset box.

If the Automate Stages and Option Reset boxes are both ticked, adjustments to parameters are generally lost when loading a new model. To avoid that, it is usually best to run stage 1 manually, then make adjustments such as choosing the maximal flattening to be allowed, and only then tick the Automate box before running stage 2.

After loading the model, some consistency checks are performed and metabolite coefficients are graphically displayed for the objective if present.

A special section in the commentary is devoted to inspection of the stoichiometry matrix to identify whether sources and sinks are supplied by the model, particularly for external metabolites. The rationale for this is further discussed in section 9 below.

Also, histograms showing the distribution of input upper and lower bounds are displayed. Based on this the program proposes an appropriate value for the value of a bound to be considered artificial and hence to be eliminated for the SSK calculation. This value is shown as a vertical blue line on each histogram, if it falls within the histogram range. Upper bounds to the right, or lower bounds to the left of the blue line are considered artificial and will be eliminated in Stage 2. An example is shown in Figure 7 :

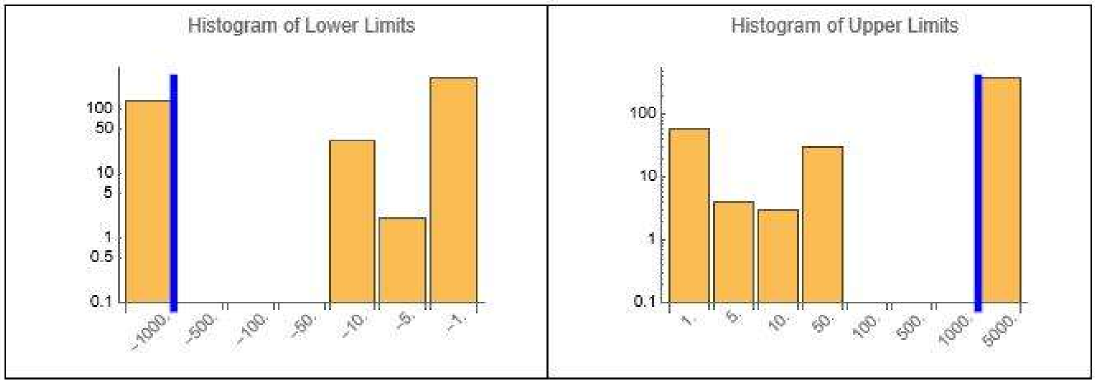

*Figure 7: Example of upper and lower bound histograms.*

The user should inspect the histograms and modify this choice as seems appropriate. In particular, in the case of models that have suffered multiple rounds of revision, some authors may for example use 1000 as the artificial bound value and others 100 000, resulting in multiple bands in the histogram. SSKernel is unlikely to pick this up, so an appropriate value may need to be manually entered in the lower box of stage 2.

#### 4.3 Stage 2

The first step of this stage is to perform an FBA calculation for the metabolic model as loaded, including artificial bounds on flux components. This is followed by a second FBA, where all artificial constraints are removed, and an unbounded LP objective optimization is performed. This serves as the basis for the fixed flux identification that follows, and that requires it to be done by using the Simplex LP algorithm. That can be considerably slower than the Interior Point algorithm used for the initial bounded FBA calculation for large models. Some control over this can be exerted by adjusting the LP tolerance box for which default values are set according to the model size.

For some large or troublesome models, increasing the LP tolerance to obtain convergence for the bounded FBA, can cause later LP calculations to fail. An example is the E-Coli K12 model IML1515, which has 2712 flux variables and it fails to converge at the default LP tolerance of 0.001. Resetting this to 0.1 does give convergence, but spoils the convergence of later stages. The best way to deal with this, is to enter a pair of tolerance values, separated by a comma, in the LP tolerance box instead of the normal single value. The first of these (*FBAtol*) is used for the bounded LP only, while the second (*LPtol*) is used for all subsequent LP's. An alternative is to simply allow the bounded LP to fail; as its result is only for validation, a failed value will simply be ignored and most of the rest of the calculations proceed as normal.

Artificial bounds are automatically identified during stage 1 based on a histogram of limit values as described above. If this was unsatisfactory, a value can be entered manually in the artificial bounds box before stage 2 is run.

Identification of fixed fluxes requires a numerical criterion for the range of variation of an individual flux over the solution space, that can be considered to be fixed. This value is normally chosen to be 5 – 10 times larger than the LP tolerance, to avoid picking up spurious variation caused by LP inaccuracies. However, it can be adjusted in the fixed value tolerance box, which contains the value of variable *fixtol*.

In addition to allowing for inaccuracies, choosing the fixed value tolerance can be a key lever to reduce large models to manageable dimensions. In stage 2, fixed fluxes as well as some classes of rays are removed, and the stoichiometry constraints applied to produce the RSS (reduced solution space). For medium sized models, the RSS dimension (i.e., its variable count) is usually well below 100. This allows subsequent calculations to be performed efficiently. Values around 150 are usually OK, but once it gets beyond 200 subsequent stages are likely to become intractable.

The best remedy for such a situation is usually to increase the fixed value tolerance until the RSS dimension decreases to an acceptable level. Even an apparently large fixed tolerance may be plausible if it remains small in comparison to the overall range of variation of the remaining variables. This can be judged retrospectively e.g. from the main chord lengths of the resulting SSK and its periphery points. *SSKernel* keeps a watch on this and alerts the user if the significance of the SSK result seems threatened by the choice made for the fixed value tolerance.

There is also an interplay between the value chosen for this tolerance and the maximal extent that is allowed to be flattened in Stage 4. As flattening amounts to taking the flux combination that corresponds to some thin direction of the SSK as a constant, it is plausible to use the same cutoff value there. *SSKernel* automatically enters the value used for Stage 2 also in the appropriate box for Stage 4, although the user can change this before executing the later stage.

The connection between thinning and the fixed tolerance also gives some justification for choosing a relatively large *fixtol* value in stage 2. There is no point in carrying along fluxes with minor variation ranges, only to be eliminated by flattening in stage 4 to avoid excessive aspect ratios. In that case it makes more sense to isolate such fluxes as fixed from the start, and so facilitate the computing intensive operations in stage 3 by reducing the dimensions of the space in which they are carried out.

A further option to influence the course of a calculation is offered by the Reshuffle first tick box, which is normally left blank. When ticked, Stage 2 is started by a random reordering of the rows and columns of the stoichiometry matrix, i.e. a random rearrangement of the metabolites and reactions.

In principle, the order in which reactions and metabolites are listed is arbitrary, so reordering them should make no difference. In practice, however, the numerical procedures can be sensitive to this ordering. For example, in the Simplex procedure for linear programming, the final optimized objective corresponds to a particular polytope vertex out of the many that share this objective value, and the reordering will most likely cause it to end up at a different vertex, i.e. with a different flux value, even though the objective value is the same. That can influence the course of the entire calculation. Anecdotally, significant differences are only observed for large models where the initial FBA calculation sometimes has trouble to converge.

In such cases, it is worth trying to tick the Reshuffle box and repeat Stage 2. Not only can this help convergence per se, but in these cases the number of fixed fluxes found also seems to be sensitive to reordering, presumably because the search starts from a different point in the solution space polytope. A larger number of fixed flux values can greatly improve the viability of the entire SSK calculation.

The Reshuffle option is also in general useful to investigate the robustness of the calculated numerical values for a particular model. absent

A final tick box labelled Exempt unbuffered externals offers a way to experiment with the effects of source or sink reactions that are ineffective or absent from the model. By default it is unticked and is usually best left that way.
Section 9 of this manual discusses the reasoning behind this facility and the cases where it might be useful.

#### *4.4 Stage 3*

Stage 3 starts by finding a set of rays to form a basis for the RSS ray space that is vectorially (over)complete (although not convex complete), and enters the vectors as well as their overlaps with constraint vectors, into a ray matrix. It uses the ray matrix to find a sample of FBF's, that are backtracked on the facet tree to discover their progenitor facet. The number of FBF's in the sample can be set in the Progenitor sample size box, but the default of 20 is usually adequate.

Coincidence capping to the progenitor facet is the next dimension reduction step, before a full search for BFBF's is attempted. Based on the ray matrix, *SSKernel* estimates the number of tree nodes that will need to be traversed for a comprehensive search. An upper limit for this can be set in the Max BFBF tree nodes box. As a guideline, around 100 nodes might be traversed per second.

If the estimate predicts that this limit is insufficient to find all BFBF's, a randomised greedy search for FBF's is made instead of the tree search. As this progresses, more and more of the FBF's that are found just backtracks to the set of basis facets (BFBF's) already found. This is counted as a failed attempt at finding a new BFBF. The search is abandoned when an allowed number of consecutive failed attempts is exceeded (with the counter being reset to zero at each success). This allowed number can be set in the failure rate box. To prevent this from continuing indefinitely, an overall maximal target for the number of BFBF's to be collected by random search is also set in the BFBF random greedy sample size box. In practice, the default value of 500 is seldom reached, and the capping radii subsequently found in actual trials turn out to be insensitive to this number.

The final adjustable parameter in stage 3 is the Default capping radius. This value only comes in play if the SSK is of the type *Simple Cone* i.e. there are no BFBF's and the minimal SSK is a single point at the apex of the cone. In this case, strictly speaking all capping radii would be zero, but instead an arbitrary default radius is used for capping to establish a finite SSK so its (cross-sectional) shape can be investigated.

Having achieved a bounded SSK, the final step in stage 3 is to find a maximal radius inscribed hypersphere and move the coordinate origin to its centre. This was illustrated in Figure 3.

The calculations performed in stage 3, particularly finding FBF's, can become very computationally intensive for large models. The determining factor is not so much the original model size, but the dimension count of the RSS. For more than 200 flux variables, the calculation can appear to come to a standstill. To alleviate this problem, SSKernel presents the user with three choices after the ray matrix has been calculated and if the RSS dimension is larger than the limiting value set in the configuration file.

The 1$^{st}$ choice is to abort stage 3 so that stage 2 can be repeated with an increased *fixtol* value and so reduce the RSS dimensions. This choice may compromise a clear distinction between fixed variables and those that are eventually eliminated if flattening is performed in stage 5.

The 2nd choice is to skip the rest of stage 3 entirely and simply use the default capping radius to perform tangent capping of all the ray directions that remain in the RSS. In effect the SSK is treated as a simple cone, and the potential dimension reduction from coincidence capping is sacrificed. Also, the extent of the RSS becomes somewhat arbitrary instead of being limited by the combined flux constraints as embodied in the bounded facets. Still, if a realistic value is used for the default capping radius that may be acceptable.
The 3rd choice is to continue stage 3, after drastically reducing the sample sizes so that only a few BFBF's are calculated. Reducing the progenitor sample size to a value between 1 and 3 may be enough for coincidence capping to reduce the dimensions sufficiently that the BFBF search for tangent capping can still be performed. But usually the latter sampling value also needs to be reduced from its default value of 500, to somewhere between 1 and 10. Both of these changes increase the risk that the capping radii may be too small, so that some bounded facets are intersected by capping hyperplanes, in which case the SSK gets truncated. Anecdotally, testing this for small models still gives quite a good approximation to the proper SSK, but this cannot be guaranteed.

#### 4.5 Stage 4

Stage 4 is concerned with calculating parameters that characterize the shape of the finite SSK, and using this to determine if flattening is appropriate. If so, the flattening is performed and the shape of the reduced SSK recalculated.

The start of this process is to calculate the main chords of the SSK. Preferably this is done using LP, but where this takes too long the remaining chords are approximated by diameters through the refined centre point. Centre refining is intertwined with chord calculation, as any known chord endpoints help to improve centre refining.

The user can set the minimum and maximum number of chords to be calculated by LP, in the Minimal, Maximal LP chord counts box. The format is as a bracketed list, e.g. {10,50}, including the curly brackets. *SSKernel* determines the actual number within these limits, based on the SSK dimensions. The LP calculation of main chords involves sampling over several SSK orientations. Reducing this sample size entered into the Maximal flips to find LP chords box speeds up the chord calculations, but can decrease the reliability.

Decisions by *SSKernel* whether to perform flattening, are made on the grounds of two adjustable parameters. First, a maximal acceptable aspect ratio for the SSK (*maxaspect*) is set in the Aspect ratio's larger than this are flattened box.

However, a value can also be set for the maximum thickness that is allowed to be flattened (*maxthin*), in the Diameters larger than this not flattened box. This value overrides the aspect ratio criterion, so setting a value of 0 prevents any flattening while a large value (even Infinity) in effect bases decisions purely on the aspect ratio.

Repetition of Stage 4 is often needed to obtain the SSK shape judged most appropriate by the user. The program gives feedback on the decision it made, in the form of a "Flattening Decision Plot" in the report window. Note that individual numeric chord lengths are interactively displayed in the report window, by floating the mouse pointer over the relevant coloured plot point.

For the example seen in Figure 8, the SSK has 31 dimensions. There are 12 mutually perpendicular directions in which the SSK has large chord lengths in the range 100 – 1000. Then there is a sharp drop to a set of values between 0.01 and 0.1. Purely based on flattening all aspect ratios larger than 50, would set the flattening threshold at the level indicated by the blue line, a value of roughly 3.

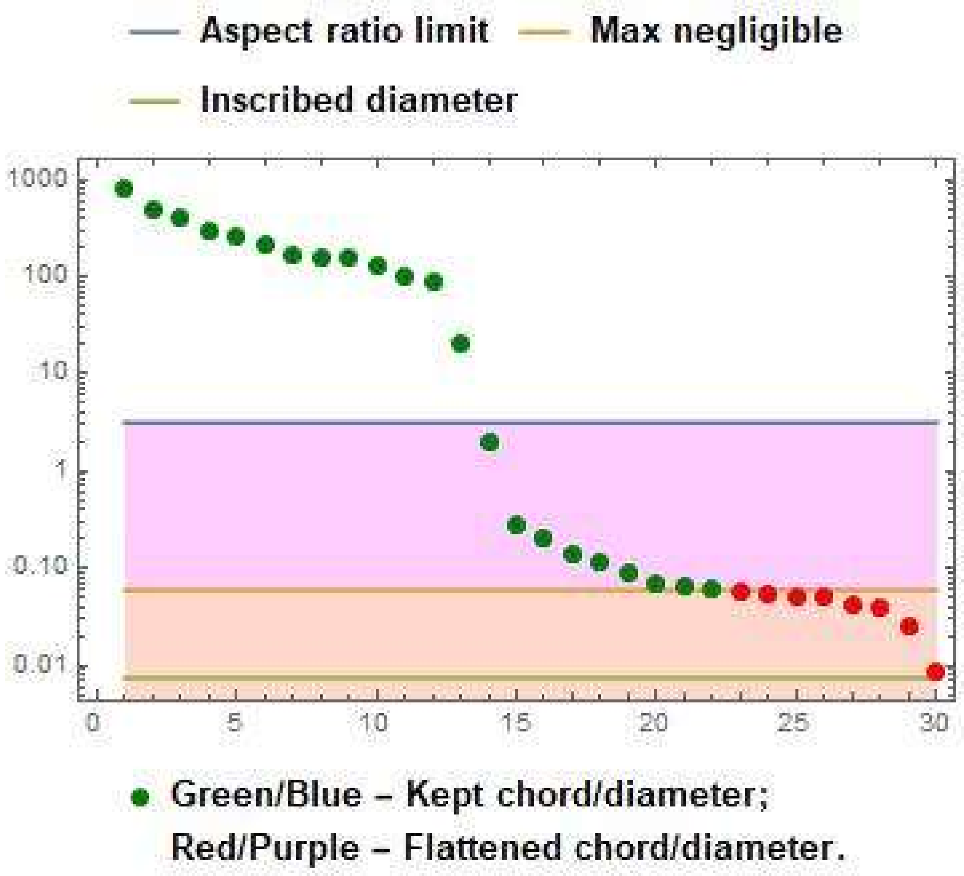

*Figure 8: Example of a Flattening Decision Plot ( for model iCCSA643_BMC ). Flux space directions with chord lengths in the pink area qualify for flattening because they represent aspect ratios larger than the preset limit; but only those in the brown area (red dots) were actually flattened because the maximum thickness allowed to be neglected was set at 0.06. The light green line indicates the smallest diameter of the SSK in any direction. The absence of blue and purple dots indicate that all chords were calculated by the more accurate LP method in this example.*

However, the result of the validation analysis given in the status report window estimates that this results in an average error of 13% for marginal flux values, i.e. those reached from points near the periphery of the SSK by addition of an appropriate ray. In the example, the user has decided that this is unacceptable and by setting *maxthin* in this case to 0.06 and repeating Stage 4, the estimated flattening error was reduced to 3.4% at the expense of increasing the maximal aspect ratio to nearly 10000. If the default *maxthin* value equal to the fixed value tolerance of 0.002 was used, there would be no flattening and the aspect ratio would be even more extreme.

User decisions on choosing a value for *maxthin*, and hence to control flattening can be based on a number of considerations:

1. Consistency with the criterion that was chosen to define fixed flux values, dictates that *maxthin* should not be smaller than *fixtol*.
2. No SSK chord can be shorter than the inscribed hypersphere diameter (before flattening). So choosing *maxthin* as zero or any value smaller than the inscribed diameter, will prevent any flattening.
3. After flattening, the inscribed diameter should normally increase to a value larger than *maxthin*. If this fails to happen (and which is flagged by a message in the status report), it indicates that there are still undiscovered thin directions that need flattening out (or that the user deliberately chose not to flatten out some known ones). The reason that thin diameters can remain undiscovered, is that because the direction of each main chord is constrained to be orthogonal to all previous ones, there is no guarantee that the last chord on the list will actually be along the thinnest direction. Simply repeating Stage 4 will often solve this.
4. At the other extreme, setting *maxthin* to the value Infinity allows flattening of all directions that give aspect ratios larger than *maxaspect*.
5. Intermediate choices can be made on the grounds of acceptable flattening error levels as in the example of Figure 8.
6. In addition, it may also be useful to inspect the final validation values delivered by stage 5. However, when these show significant deviations, it should be borne in mind that the FBA solutions are not very representative of the entire SSK and especially when they only carry a small flux, quite large relative errors may still only be small absolute flux deviations.

#### *4.6 Stage 5*

There are no adjustable parameters for stage 5. Its main purpose is to round off the calculation by presenting summary measures in the status report window that describe the SSK itself and the process that leads to it. Among these is a table listing the outcomes of the validation tests on FBA flux values, before and after flattening. Also, two graphs shown in Figure 9 are produced to give a visual overview of the calculated chords, aspect ratios and centring.

In Figure 9, the number of bars gives an immediate visual impression of the dimensionality of the SSK, since each represents a mutually orthogonal chord.
For a hypersphere all lengths will be equal; for a highly symmetric shape, such as a hypercube or simplex, at least half are usually the same length, and the rest are only shorter by a single digit factor. Bearing in mind the logarithmic scale, the example shown in Figure 9 is clearly far from such a regular shape and this is typical of most SSK polytopes.

The monotonous decrease of the bar length in the top plot of Figure 9 merely reflects the fact that chords have been ordered in this way. The figure only shows the chord lengths, but their directions in flux space are given by the SSK basis vectors explicitly listed in the data structure of equation (1.2) and that appears as the first item in the data output file. The basis vectors are ordered in the same way, so there is a one-to-one correspondence between each displayed chord length and its corresponding basis vector direction.

The mean aspect ratio value shown below the top plot in the figure is calculated as the average of the pairwise aspect ratios of chords, taken over all possible pair combinations. Also, the maximal aspect ratio, i.e. the ratio of the longest to the shortest chord, is shown.

A context for these values is provided by considering the values for highly regular shapes. In 2 or 3 dimensions, it is intuitively clear that the more facets a regular polytope has, the closer it approaches a hypersphere; for example, a regular hypercube (square or cube) is closer to a hypersphere (circle or sphere) than a regular simplex (triangle or tetrahedron). Typically, the number of facets for a SSK (i.e., its number of constraint hyperplanes) is around 1.5 – 1.8 N where N = number of dimensions. The minimum number to give a closed polytope is the (1+1/N) N for a simplex while a hypercube has 2 N facets. So a SSK (if it was regular in shape) could be considered to be intermediate between the simplex and hypercube cases in the trend towards full spherical symmetry.

*Table 1: Aspect ratios as function of dimension for regular polytopes*

| | Dimensions N | 2 | 3 | 15 | 50 | 100 |
|---|---|---|---|---|---|---|
| Mean aspect | Hypersphere | 1 | 1 | 1 | 1 | 1 |
| | Hypercube | 1.0 | 1.3 | 1.2 | 1.2 | 1.3 |
| | Simplex | 1.2 | 1.3 | 1.4 | 1.5 | 1.5 |
| Max aspect | Hypersphere | 1 | 1 | 1 | 1 | 1 |
| | Hypercube | 1.0 | 1.4 | 2.2 | 2.5 | 2.7 |
| | Simplex | 1.2 | 1.4 | 2.7 | 5 | 6.8 |

Table 1 shows aspect ratios calculated by *SSKernel* for these regular polytopes in different dimensional spaces. The mean aspect ratios in the table reflects the trend noted above and as the values show very little dependence on the dimensions, it serves as a plausible measure for isotropy and symmetry.

The maximal aspect ratio also reflects this trend, an in addition shows that even for these highly symmetric shapes there is a tendency for the difference between the largest and smallest orthogonal chord lengths to increase moderately with dimensions. This is simply because the length maximisation of each subsequent chord is more constrained by the requirement that it is orthogonal to all previous ones. The values in the table shows that this factor only accounts for single digit factor increases in the maximal aspect ratio over the range of dimensions relevant to SSK calculations.

By contrast, the example of Figure 9 shows a mean aspect ratio of 38 and a maximal value of 7593, indicating that its SSK polytope shape is far from being isotropic.

The lower plot in Figure 9 shows the radius to the periphery, along the forwards and backwards directions of each of the chords shown in the top figure. The plot is arranged in such a way that the longer of these is always shown above the axis. The total length of each coloured bar amounts to the diameter of the SSK along each chord direction. This is generally considerably smaller than the corresponding chord length, because it is by definition constrained to pass through the coordinate origin.

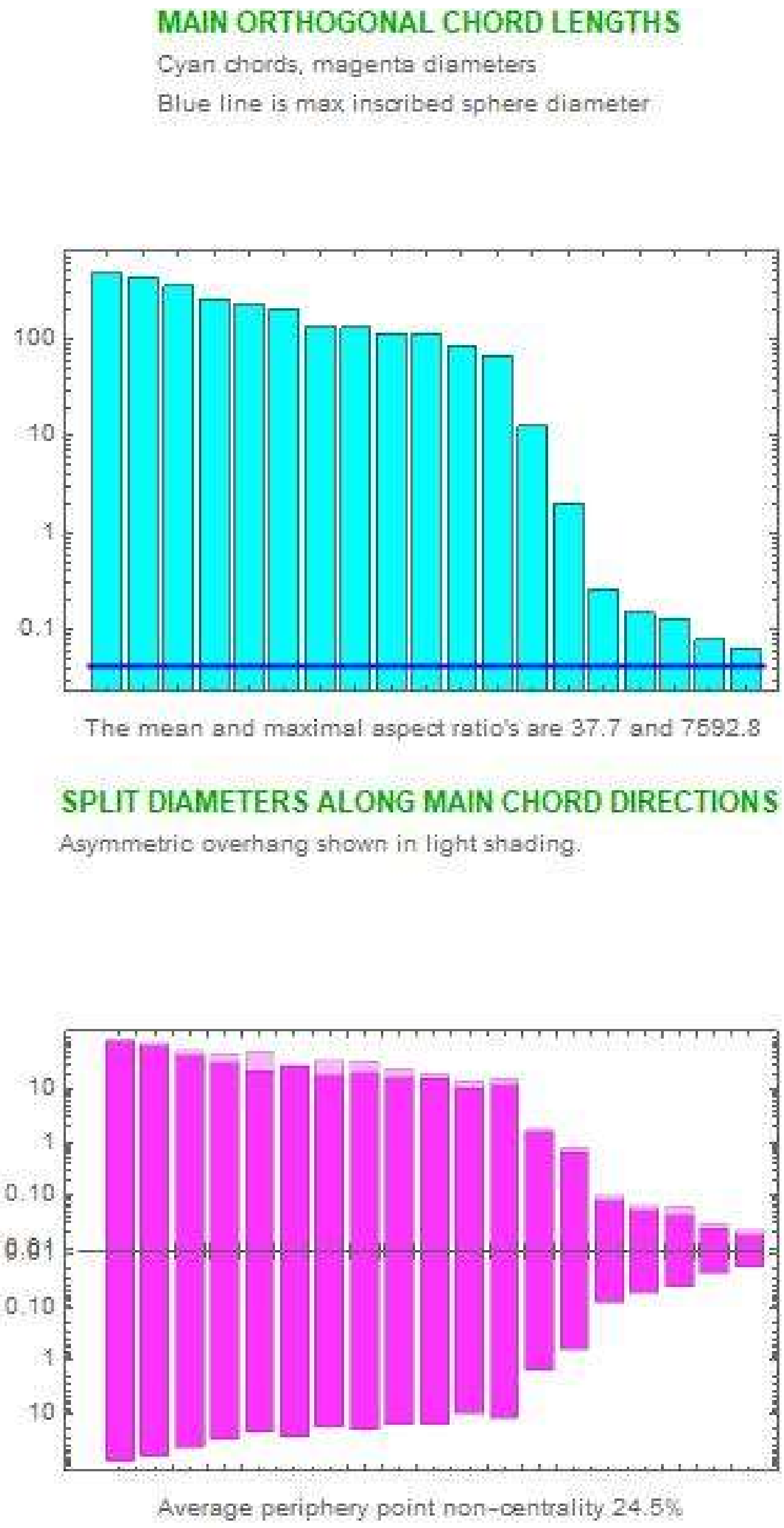

*Figure 9: SSK shape and geometry as expressed by chords and centring, as calculated for model iCCSA643_BMC.*

The magenta bars below the axis are symmetrically copied above the axis, to bring out the difference as a lighter shaded overhang. The fact that in the example shown, these overhangs are quite small, indicates firstly that the SSK is reasonably symmetric in each individual direction, and secondly that the centre refining process was quite successful in selecting a point that is centrally located, as the coordinate origin.

The non-centrality measure shown below the figure, indicates that the overhang is on average 24.5% of the corresponding diameter, taken over all the peripheral opposing point pairs (not just the chord directions shown in the figure). By way of comparison, this value works out as 0% for the hypercube in all dimensions, but ranging from 12% at N = 2 to 47% at N = 100 for the simplex. Consideration of the 2D and 3D cases makes it clear why perfect centring is possible for a hypercube but not for a simplex.

The final result produced by Stage 5, is the comparison of mean inscribed radii for the simplexes that respectively enclose the full SSK and the PPP (see Figure 4). For small N, a sampling estimate of the PPP coverage is also displayed in the status report window.

#### 4.7 Stage 6

The final stage optionally exports a PDF copy of the status report window and the detailed numerical specification of the SSK, ray vectors etc. as two output files. The files are stored in the same directory as the input file, and with generic names based on that of the input file. See section 7.2 for a full description. If either of these files already exists, the new file is saved with an allocated number.

### 5 Subsequent use of the SSK results.

The *SSKernel* program is intended as a tool for analysing and comparing metabolic models, not as providing the final outcome.

The "Report" output file documents the results of a particular run of the program, including the control options used. It shows the dimension reductions achieved in the various stages, and various characterisations of the SSK obtained such as its dimensionality, inscribed hypersphere, enclosing radius, aspect ratios and chord lengths. These values give a qualitative view of the size and shape of the SSK as a geometric object, in a way that can perhaps be usefully compared between different metabolic models.

However, for further quantitative processing, the detailed numerical data in the DIF output file is provided separately. To demonstrate how this data may be read and processed, an additional *Mathematica* notebook file *ExploreKernel.nb* is included in the SSKernel software.

As a first example, it displays fluxes that acquire fixed values or directions in the model as named reactions and flux values.

The key result, the SSK specification, is disassembled into its components and each of them displayed in graphical form or with reaction names where applicable. Complementary to this, ray directions and thin directions that were taken out are similarly displayed.

The relationship between the small dimensional SSK and the original full flux space (FFS) is explored by showing the periphery points in terms of both coordinate systems, in the process also demonstrating how the SSK specification is used to transform vectors between them. To achieve this, a few utility functions are provided as part of the *ExploreKernel.nb* notebook and their use demonstrated.

Next, the distribution of periphery points are explored by constructing a histogram of pair angles between them. Another use of periphery points to gauge the range of variation of individual named fluxes over the extent of the SSK, is given as a demonstration, as well as a small example of using them for averaging calculations.

Finally, it demonstrates how to explore the effects of bioengineering interventions, such as gene knockouts, on the metabolic space. The tools for this are taken from the *Resampling.m* Mathematica package that forms part of the SSKernel software, although not directly used by the interactive user interface. The main tool is the function *HyperSampler*, which resamples representative points from the intersection of the SS with any hyperplane that can be specified by a set of linear constraints.

It is hoped that these demonstrations can serve as a template that can be expanded for further calculations and analysis.

### 6 Command line implementation

For users who do not have access to a full *Mathematica* installation, a command line version can be run which only needs installation of the Wolfram Engine software, which can be obtained free of charge together with the WolframScript interpreter as set out in section 2.

This version uses exactly the same *Mathematica* code files in the Packages directory as the notebook version, but executes all stages in sequence without any possibility of user intervention. Adjustable parameters are in this case initialized in the configuration file, and can be set to desired values in the command line used to invoke the wolfram script file *SSKernelscript.wls*. Some of these variables may still be further adjusted by the program itself based on calculation results. The final values that were used in a specific run are listed in its Report output file.

*SSKernelscript.wls* is located in the subdirectory *CommandlineFiles* together with a batch file *SSKernelbatch.bat.* This subdirectory should be at the same directory level as the subdirectory *Packages* that contains the main SSKernel code as a series of *.m files. The code in the script and batch files is plain text that can be modified to suit any other file organisation.

Detailed explanations of how wolfram script files are used and run in various operating systems, are available at https://www.wolfram.com/engine/?source=nav . The examples here are based on running the script in the *Windows 10 PowerShell* environment and may need slight adaptation elsewhere.

The script file can be run with up to 2 arguments, each given as a string enclosed in double or single quotes:

- Without an argument, it will process the first .mat or .m input file that it finds in the subdirectory "TEST" located in the same directory as itself, and with default option values.
- If there is one argument, this will be taken as the file path of an input file to be processed. Whether this path specification is to be absolute or relative, depends on the values assigned to the variable *DataDirectory* in the file "Configuration.m" located in the "Packages" directory.
  With the default assignment `DataDirectory=""` a full (absolute) path to the data file needs to be given in the command line argument. To avoid the need for a tedious command line, any non-empty string value assigned to *DataDirectory*, is prepended in the script to the path string supplied in the argument.

  Examples of appropriate assignments in Configuration.m are:

  ```
  DataDirectory="C:/Users/John Smith/Documents/Data/"
  DataDirectory="C:\\Users\\John Smith\\Documents\\Data\\"
  ```

  Either forwards (Unix-style) or backwards slashes (Windows style) may be used, but in the latter case each backslash needs to be doubled because *Mathematica* uses this as an escape character in a quoted string.

With *Datadirectory* terminated by a slash, an appropriate command line argument in each case might be

```
"Clostridium/IHN637.mat"  or  "Clostridium\IHN637.mat"
```

with no need to double the backslash in the command line.

- If there are two arguments, the first is the input file as above and the second is a quoted, comma separated list of variable assignment statements, enclosed in curly brackets. Only changed control variables need to appear, in any order, and override the corresponding statement in the configuration file. E.g.:

```
"{verbose= False, Tolerances='0.001,0.0001', maxthin = 0.02 }"
```

Note the use of double and single quotes, shown for Windows. It may be necessary to swap these for other operating systems. The control variables that can optionally be adjusted in the command line are listed and described in the table below.

*Table 2: Adjustable control parameters*

| Name | Type | Default | Description |
|---|---|---|---|
| verbose | Logical | True | True means detailed progress feedback in console/control window. |
| loadreset | Logical | True | True means program may reset values according to input data |
| LPmethod | String | Automatic | Method used for initial bounded FBA: “Simplex” or “Interiorpoint”. |
| Tolerances | String | “0.0001” | Tolerance(s) for LP convergence |
| fixtol | Real | 0.002 | A flux varying less than this is taken as having a fixed value |
| artificial | Real | 100. | Flux upper limits equal or larger than this are considered artificial and set to Infinity |
| targetcount | Integer | 20 | The number of random FBF's for initial progenitor estimate |
| maxthin | Real | 0. | SSK thickness less than this may be flattened to zero |
| maxaspect | Real | 50. | Avoid a SSK aspect ratio larger than this, by flattening as allowed by maxthin |
| samplesize | Integer | 500 | The target number of BFBFs to be found, if using greedy random search. |
| greedyfails | Integer | 20 | Terminate sampling if this number of consecutive rejections indicate a small prospect to find any more BFBF's. |
| treesize | Integer | 200000 | The maximum estimated number of tree nodes to be visited by exhaustive search, otherwise use greedy search |
| chordmin | Integer | 10 | The minimal number of main chords found by LP. |

| | | | |
|---|---|---|---|
| chordmax | Integer | 50 | The maximal number of chords found by LP, any more are approximated from periphery points. |
| flipmax | Integer | 25 | The maximal number of times quadrants are flipped during main chord calculation. |
| DefaultCap | Real | 1. | Default capping radius is applied if SSK is a simple cone with no FBF's |
| timeconstraint | Real | 60. | Seconds allowed for calculating flattening errors and enclosing simplexes |

Note that, depending on the value of variable *loadreset*, the program may modify some parameter values during the course of the calculation. The values actually used are listed in the Report output file.

There are various ways to invoke *SSKernelscript.wls* and transmit its arguments.

a) In Windows Explorer, simply double click *SSKernelscript.wls* .
This passes no arguments and processes an input file it finds in the TEST directory. It is mainly useful for testing that the installation functions correctly.

b) In Windows Explorer, right click the .mat or .m data input file and using the "Open with" submenu in the context menu that appears, select the batch file *SSKernelbatch.bat*, (not the script!) in order to process it.
In some operating systems, it may be possible to simply drag-and-drop the input file on the batch file (or even the script file, if the default control parameters are to be used) but this facility appears not to be supported from Windows 11 onwards.
The control parameter argument passed in this case, is defined in the batch file and may be adapted as needed by first editing *SSKernelbatch.bat.* Also see comments in the batch file for hints on dealing with possible problems that may arise when running it on a network file system rather than locally.
This method is useful when processing several input files with the same set of control parameters.

c) Open Windows Powershell by a right click on the Windows Start button.
   a. Change directory by a command such as
```
CD "C:/Users/John Smith/Documents/SSKernel"
```
to the directory containing the commandline and SSKernel packages subdirectories; confirm this with the DIR command if needed.
   b. Enter a command line such as
```
.\CommandlineFiles\SSKernelscript.wls
"Clostridium/IHN637.mat"
  "{verbose=False, Tolerances='0.001', maxthin=0.02}"
```
This relies on the operating system associating *wolframscript* with the file extension .wls, as should happen during installation of the Wolfram Engine. A console window is opened where the control window text output as described in section 4 appears, including error messages.
This is the recommended way to run SSKernel using a command line.

d) If the file association fails, the script interpreter can be invoked explicitly by replacing the command line in method c)b by a line like
```
wolframscript -f CommandlineFiles\SSKernelscript.wls
"Clostridium/IHN637.mat"
"{verbose=False, Tolerances='0.001', maxthin=0.02}"
```
In this case unlike the previous methods, there is no dedicated window for the program text output, and it appears directly in the Powershell console window, which may be useful for diagnosing problems but is otherwise probably not optimal.

e) Finally, instead of using Powershell, the commands in methods 3 and 4 may also be executed in the Windows command interpreter, most easily accessed by typing “CMD” in the Windows search box. Again, the disadvantage is that output appears directly in the command console instead of opening a dedicated window.

### 7 Directories and files

The *SSkernel* files are publicly available under the GPL license from GitHub.com, at the repository "SolutionSpaceKernel", from where this manual was presumably obtained. The full package also contains various files with source code, etc. in a zipped archive.

The downloaded zip-file unpacks into a directory, e.g. "SSKernel_V1.2.3", which contains the main *Mathematica* notebook file *SSKernel Launcher.nb* that drives the application, a subdirectory "TEST" that contains one or more example data files, and the subdirectory "Packages" where most of the program code resides as well as the "CommandlineFiles" subdirectory. The only critical part of this is that the naming and positioning of the CommandlineFiles and Packages directories relative to the notebook should be maintained in order for *SSKernel* to find its code.

In the "Packages" directory, the file "Configuration.m" contains default initialisation data for setting choices on the control panel. These can be edited by the user with a text editor, using care not to disturb the formatting.

It is recommended that the input data file specifying the FBA model, is located in a directory named after the biological species of the organism it refers to. This is convenient because the parent directory name is taken by default as the organism name when reading data. But this is not critical, as the organism name can be edited in the control panel by the user. So generally, the model specification input file can be located anywhere in the file system because it is explicitly selected by the user.

Output files are stored in the same directory as the input file.

#### *7.1 Input Files*

*SSKernel* can read the FBA model specification file in one of three alternative formats, signified by the file name extensions *.mat* , *.sbml* or *.xml,* and *.m* or equivalently *.wl* (i.e., WolframLanguage).

The first format is the *Matlab*-oriented *.mat* format fully defined in the COBRA-toolbox V3.0, see Table 3 in:

**Creation and analysis of biochemical constraint-based models: the COBRA Toolbox v3.0**, Nature Protocols, volume 14, pages 639–702, 2019 doi.org/10.1038/s41596-018-0098-2.

Models in this format can for example be downloaded directly from the BIGG metabolic database .

SBML (Systems Biology Markup Language) files of two types can be read by the program. Most current metabolic models make use of the Flux Balance Constraints (FBC) extension of SBML Level 3. This should generally be read successfully. SSKernel can also read older SBML files that do not store the objective and flux bounds specifications in separate sections as in FBC, but instead incorporate these as parameters in each individual reaction specification. In this case it searches for the parameters with respective id's given as either UPPER_BOUND or UPPERBOUND, or lower case or mixed case versions of these, and similarly for the lower bound and for OBJECTIVE_COEFFICIENT. Any other naming conventions may need the program code for the function SBMLReader found in the text file DataHandler.m which is in the Packages directory of SSKernel, to be appropriately edited.

In case an SBML file is not read successfully, note that the COBRA toolbox contains functions to read various other commonly used file formats such as SBML and even spreadsheets, and can store them as *.mat* files that can then be used as *SSKernel* input files.

Where the same model is available in *.mat* and SBML formats (as for most BiGG models) it is usually found that the reactions and metabolites are arranged in a different order in the two files. As pointed out in the discussion of Stage 2, this can lead to somewhat different results for each; the difference is usually negligible for small or medium sized models, but for large models it may e.g. influence whether the initial FBA calculation can converge. In such cases it may be helpful to try both input formats to see which works best.

The 3rd alternative option is to create an input file as a *Mathematica .m* or *.wl* file. This is a standard text file that contains the numeric arrays as a list with the structure

Modelname
ReactionNames
MetaboliteNames
S
Svals
Bounds
Objectvector
FBAvector
maxmin

Each item starts on a new line.

*Modelname* is a string, which may include a formal ID and a desciption of the model. It has to appear first.

The remaining items are comma separated, nested lists in curly brackets and may appear in arbitrary order. Any additional list items are ignored.

*ReactionNames* is a simple list of strings, enclosed in 1 level of curly brackets.
Each string is taken as the name of the reaction flux of the corresponding column of the stoichiometry matrix.
The *ReactionNames* list is **optional**, if absent all reactions are allocated the name "NoName" by default.

Since the SSKernel calculation takes place in flux space, *MetaboliteNames* are not really relevant, but are optionally included in the input data for information purposes. If present, they should appear after *ReactionNames*.

*S* is the stoichiometry matrix. It is listed in a 2-level nested list, row by row, each row specifying coefficients for one constraint such as a metabolite or the objective vector.

*Svals* specifies both the values (i.e., right hand sides of the constraint equations) and which inequality type the constraint represents. It is a 2-level list, consisting of the pair {*value, type*}, where types -1, 0, 1 respectively mean a Less/equal, Equal, Greater/equal equation, for each corresponding constraint in *S*.
Specifying *Svals* is **optional**, since for a strict stoichiometry matrix all equations are of type " = 0", and this is taken as the default if *Svals* is absent. It is mainly provided to enable further constraints to be added as additional rows, to the matrix that defines the chemical reaction stoichiometry for all reactions in a model.

The *Bounds* matrix is a similar 2-level list, but has a row for each flux, consisting of the pair {*loflux, hiflux*} that specify minimum and maximum values for each flux. Where the model does not specify a limit, values -Infinity and Infinity (no quotes) can be used.

Note that the *Svals* and *Bound*s arrays are recognised by their dimensionality. So the number of rows for *Svals* needs to be the same as for *S*, i.e. the number of constraints. The number of rows for *Bounds* needs to be the same as the number of columns in *S*, i.e. the number of fluxes.

*Objectvector* is a single level list of coefficients for weighting flux contributions to the objective that is minimized/maximized by FBA. Typically it has only a single nonzero entry, which selects the column of S that defines e.g. the biomass growth flux. As currently implemented, up to 4 nonzero entries are accepted in order to define a compound objective. This limitation is necessary for the program to distinguish between the objective and the FBA vector that is also a numeric vector with the same number of components.

*FBAvector* can **optionally** be included to specify an externally calculated FBA solution that optimizes the objective. It is tested during input that the given vector does satisfy all models constraints, otherwise it is rejected and a zero vector assumed by default. If present, SSKernel checks that this is consistent with its internally calculated FBA solution.

Finally, a simple string, either "max" or "min" can be given to indicate whether the objective should be maximised or minimised. It is **optional**, and if absent minimization is chosen by default.

#### *7.2 Output files*

Output files are assigned generic names based on the input file name. For the input file "XYZ.mat" there will be two output files: "XYZ SSKernelReport.pdf" and "XYZ SSKernel.dif". The report file is just a copy of the Status Report window displayed during program execution.

The second file is in Data Interchange Format ( *.dif) and contains the numeric calculation results. It is a text file that can be inspected in any text editor. It consists of several sections. The first of these is a heading that describes the model and the date and time of execution. This is followed by a sequence of section headings to identify the data that follows, and then the numerical data. Headings are quoted strings enclosed by *Mathematica* comment brackets as in (* …… *). Data are given as comma separated, nested lists enclosed by curly brackets as in standard *Mathematica* notation, and also stored as quoted strings. The main structures are interspersed by metadata that belong to the DIF specification.

The section headings are as follows, to indicate the actual output data:

(* Kernel Space Data Structure *)
(* Periphery points in flux space *)
(* Fixed flux numbers and values *)
(* Thin directions in flux space *)"
(* Ray vectors in flux space *)
(* Maximal inscribed sphere radius and centre *)
(* Reversible reactions becoming fixed in direction *)
(* Names of metabolites, and reactions that form the flux space basis vectors *)
(* Reduced Solution Space Data Structure *)

The Mathematica notebook *ExploreKernel.nb* demonstrates how to read and use the numeric output in the DIF file. See section 5.

Note that the main chords and chord lengths, that characterize the SSK shape and are graphically displayed in the *XYZ Report.pdf* file, are not explicitly identified in the *XYZ SSKernel.dif* file. However, as chord endpoints are by construction included in the list of periphery points, chords can be readily extracted once the sequence numbers of relevant periphery points are known. A special utility function **ChordPairs** is provided in the *Centering.m* package to perform this task. Its use is demonstrated in ExploreKernel.nb.

### 8 Case Study: A Toy Network

The *inputfile.m* format is best illustrated by a toy model example, Figure 10:

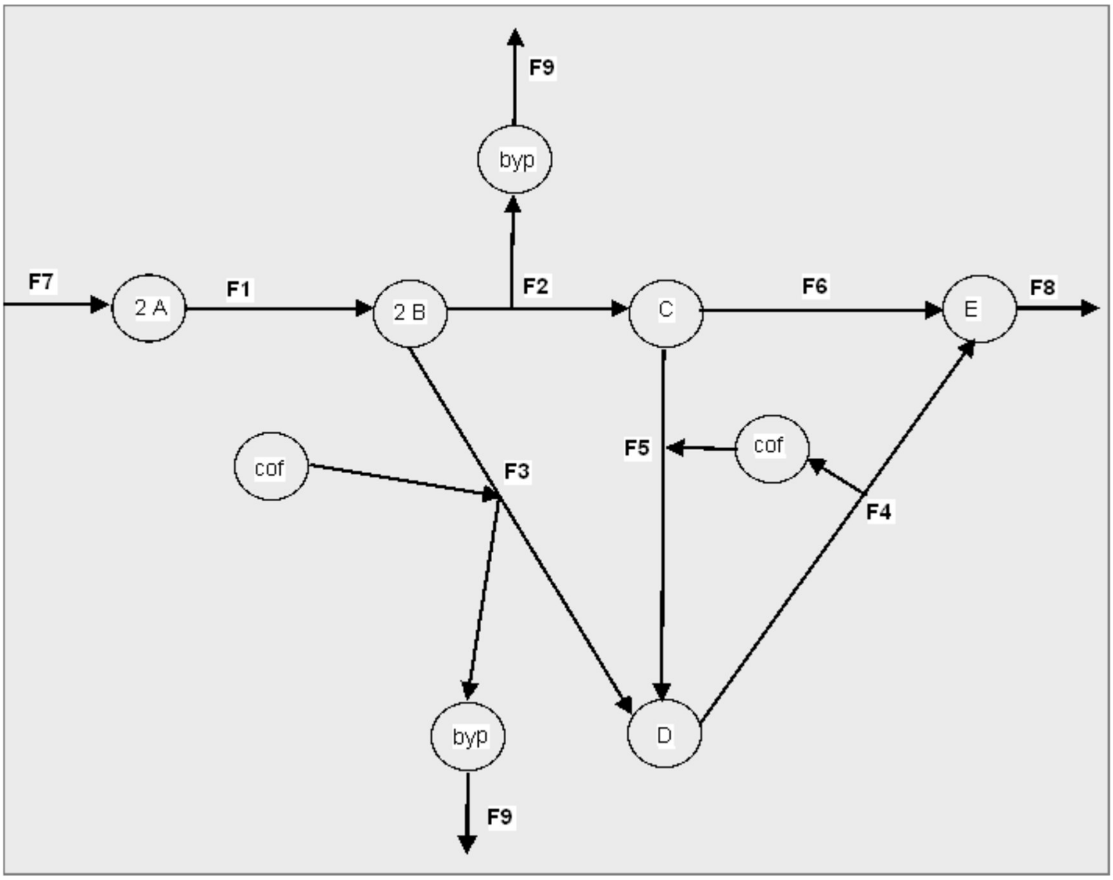

*Figure 10: A toy network, with 9 reaction fluxes F1, F2,... F9 and 7 metabolites A,B,C,D,E,byp and cof.*

This model is discussed in "*Systems Biology – Properties of Reconstructed Networks*" by B. O. Palsson, 2006, p. 204.
The .m input file for this network follows below. The reactions have been named according to flux numbers. The stoichiometry matrix is given by

| | $F1$ | $F2$ | $F3$ | $F4$ | $F5$ | $F6$ | $F7$ | $F8$ | $F9$ |
|---|---|---|---|---|---|---|---|---|---|
| $A$ | −1 | 0 | 0 | 0 | 0 | 0 | 1 | 0 | 0 |
| $B$ | 1 | −2 | −2 | 0 | 0 | 0 | 0 | 0 | 0 |
| $C$ | 0 | 1 | 0 | 0 | −1 | −1 | 0 | 0 | 0 |
| $D$ | 0 | 0 | 1 | −1 | 1 | 0 | 0 | 0 | 0 |
| $E$ | 0 | 0 | 0 | 1 | 0 | 1 | 0 | −1 | 0 |
| $byp$ | 0 | 1 | 1 | 0 | 0 | 0 | 0 | 0 | −1 |
| $cof$ | 0 | 0 | −1 | 1 | −1 | 0 | 0 | 0 | 0 |

(1.3)

which may be compared to the figure, noting the stoichiometry coefficients appearing in the nodes for A and B.

The input file for the model, but extended to include explicit bounds is the following.

```
"Toy network"
{"Reaction 1", "Reaction 2", "Reaction 3", "Reaction 4",
"Reaction 5", "Reaction 6", "Reaction 7", "Reaction 8",
"Reaction 9"}
{"A","B","C","D","E","byp","cof"}
{{-1, 0, 0, 0, 0, 0, 1, 0, 0}, {1, -2, -2, 0, 0, 0, 0, 0,
0}, {0, 1, 0, 0, -1, -1, 0, 0, 0}, {0, 0, 1, -1, 1, 0, 0,
0, 0}, {0, 0, 0, 1, 0, 1, 0, -1, 0}, {0, 1, 1, 0, 0, 0,
0, 0, -1}, {0, 0, -1, 1, -1, 0, 0, 0, 0}}
{{0., Infinity}, {0., Infinity}, {0., Infinity}, {0.,
Infinity}, {0, Infinity}, {0., Infinity}, {0., 3.0}, {0.,
Infinity}, {0., Infinity}}
{0, 0, 0, 0, 0, 0, 0, 0, 0}
{0., 0., 0., 0., 0., 0., 0., 0., 0.}
"max"
```

The objective as defined by the 3rd last line in the input, is a placeholder that sets a trivial zero vector objective. It allows nominal maximization to proceed, but in fact just calculates a flux value for the network, subject to the flux bounds set in the list appearing just before the objective.

For the "bare" network with no objective and no bounds, all nominal upper bounds would be set to "Infinity". This gives an infinite cone SS. In order to keep it finite, the stated model has limited the single input flux F7 to the range {0., 3.}. That facilitates displaying the SS shape as in the next section.

#### *8.1 SSK and FVA perspectives on solution spaces*

The toy model gives an amenable demonstration of some of the concepts encountered in the SSK analysis. It also nicely contrasts the SSKernel characterisation of the solutionspace, versus that given by flux variability analysis (FVA).

Starting with just the network model without an optimized objective, an FVA calculation shows that despite the lack of upper bounds on any flux except F7, the solution space is in fact bounded with finite, nonzero ranges for all 9 fluxes. So the FVA bounding box in this case is a cuboid in 9 dimensions.

To perform the FVA calculation, the following commands can be executed once an FBA objective has been calculated by running Stage 2 in the control panel:

```
FVAConstraints = Append[S,objectselector];
FVAlimits = Append[Svals,{objective,0}];
{lowerflux,upperflux} = Transpose[Map[Chop[{
   LinearProgramming[ #,FVAConstraints,FVAlimits,bounds].#,
   LinearProgramming[-#,FVAConstraints,FVAlimits,bounds].#}]&,
      IdentityMatrix[Length@objectselector]]];
TableForm[{lowerflux,upperflux},
TableHeadings->{{"Lower limit","Upper limit"},ReactionNames}]
```

In the SSK calculation, the fact that the solution space is bounded, circumvents the need for finding rays and capping hyperplanes. Hence the SSK is in this simple case identical with the solution space itself. Performing the vector transformations, constraint projections, centering and periphery detection on this space, agrees with FVA that there are no fixed fluxes. However it reveals in addition that the solution space has in fact just 3 dimensions and using the calculated SSK constraints, can hence be depicted as in Figure 11 (a) .

The SSK is a 3D shape, somewhat like a cheese wedge, embedded in the 9D flux space. The 3 coordinate axes in the figure do not correspond to individual flux space axes, but are constructed by the calculation to define the directions of the maximal orthogonal chords of the SSK volume (and shown as red arrows on the picture). The numerical values are obtained by entering the command **`ChordLengths`** in the control panel, and are 5.4, 3 and 1.95 respectively.

Detailed information such as the position of the SSK centre as well as the chord direction vectors, are contained in the vector transform part of the SSK data structure as described by equation (1.2). However, executing the command **`KernelSpace`** displays this as specified relative to the RSS, while it may be of more interest to see the centre location and chord directions in the full 9D flux space. This is available in the file "*Toy Network SSKernel.dif*" that is created by clicking the Export button to execute Stage 6.

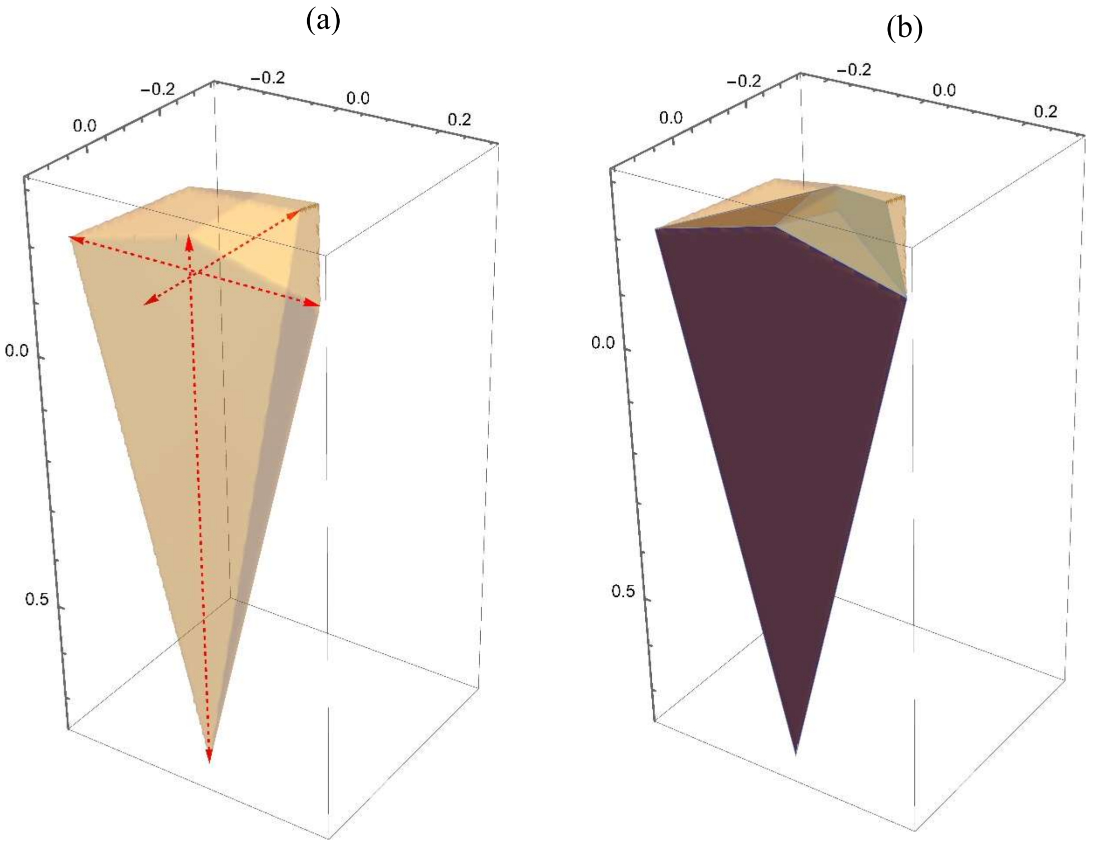

*Figure 11: (a) The bounded solution space (i.e. SSK) of the toy model without an FBA objective. The main chords are shown as red arrows. (b) The PPP superimposed on the SSK.*

However, the numerical parts of the export data are immediately available in the *ExportResults* variable as soon as Stage 5 has been completed. As described in section 7.2 below, the SSK datastructure is the first item in the exported results and according to equation (1.2) the second part of that is the vector transform. Hence the command to display the centre and chord directions, is simply

```
ExportResults[[1,2]]
```

Other exported data such as ray vectors or the inscribed sphere are similarly accessible by inspecting appropriate items in the *ExportResults* array.

As part of the centering and chord calculations, a collection of explicit peripheral points is generated. In Figure 11 (b) the Peripheral Point Polytope or PPP formed by using these as vertices (strictly, the convex hull of the set of points) is shown, superimposed on the SSK as defined by its set of constraint equations. It is seen that the PPP covers a large part though not all of the SSK interior.

The apparent contradiction between the fact that the SSK calculation shows that there are only 3 degrees of freedom while the FVA calculation finds all 9 fluxes to be variable, is accounted for by the fact that the three SSK basis vectors each involve all 9 flux axes. This point is further clarified in the example below where a biomass objective is added to the model.

But first, consider a minor elaboration of the toy model in which the FBA objective is taken to be maximisation of the sum of the two metabolic product fluxes F8 and F9. This is easily done by setting `objectselector = {0,0,0,0,0,0,0,1,1}` as detailed in section 4.1.

This optimisation is found to produce four fixed fluxes, namely F1, F7, F8 and F9. That leaves the FVA bounding box as a 5D cuboid. The SSK in turn reduces to a 2D quadrangle, in effect a cross section of the solution space wedge. There is again a marked dimension discrepancy between the FVA and SSK results. Both agree on the value range of individual fluxes, but the SSK gives far more exact geometrical information on the metabolic flux variation that is allowed by the stoichioimetry and other flux constraints.

A more realistic elaboration of the toy model is to assume that metabolic products have to conform to a known biomass composition. In this case, assume this to be 80% of metabolite *E* in Figure 10, and 20% of the byproduct *byp*. There are various ways that seem plausible to introduce an objective in order to maximize biomass production with this 4:1 proportion:

1. Set `objectselector = {0,0,0,0,0,0,0,0.8,0.2}` to maximize the appropriately weighted sum of output fluxes F8 and F9..
2. Replace the output fluxes F8 and F9 by a single output flux, `biomass = {0,0,0,0,-0.8,-0.2,0}` and maximize its flux.
3. Keep F8 and F9 but add the same biomass flux as an additional flux that is then maximized.

Option 1 is the easiest to implement but in fact gives exactly the same set of optimized fluxes as maximizing the simple sum as mentioned above, as does any other weighted sum with positive coefficients. Moreover, the maximized objective value does not actually give the optimized biomass flux value as it represents only 80% of F8 and 20% of F9, whereas the total biomass flux in this case consists of 100% of F8 and the appropriate fraction, in this case 25%, of F9 to make up the required 4:1 ratio.

Option 2 would force the output fluxes to be produced in 4:1 ratio. That may or may not be compatible with the mass ratio's determined by the other network constraints. In the present case, these constraints require metabolites E and byp to be produced in equal amounts and so if option 2 is implemented, the only solution is the trivial one where all fluxes are zero. Generally, option 2 might work in cases where the network mass balances leave sufficient flexibility, but that is hard to predict.

So only option 3 is in general a viable option. Note that (as for option 2) the coefficients for F8 and F9 have to be taken as negative because they form the inputs to a nominal biomass reaction. Creating this additional output channel for the biomass leaves the original F8 and F9 intact to account for any excess metabolite production that is required for mass balance. Its optimised flux indeed represents the maximal biomass that can be created when input flux F7 is limited. The required code for adding this modification as an additional biomass flux was already shown in section 4.1, page 19.

Out of the resulting 10 fluxes, a total of 7 become fixed by maximizing the biomass flux and leaves a 3D FVA bounding box. On the other hand, only a single degree of freedom remains, so that the solution space and hence also the SSK reduces to a single finite line in the 10D flux space. It turns out that it is fluxes F4, F5 and F6 that remain variable, and the subspace that they define can be sketched to show both the FVA bounding box and the SSK at the same time. This is done in Figure 12.

The figure illustrates how the discrepancy between the dimensions of the SSK and the FVA bounding box, that was remarked upon in discussing the previous versions of the toy network model, arises in this case. It is seen from the figure that the FVA bounding box gives a rather scant account of the actual solution space. It gives a general impression of where in the flux space it is located, and the extent of flux variation allowed. However, it gives virtually no indication of the shape, orientation or even dimensionality of the SS. It might be thought that sampling points from the bounding box could be used pin down these details. But the SS occupies only a negligible fraction of the bounding box interior, making the sampling approach unviable.

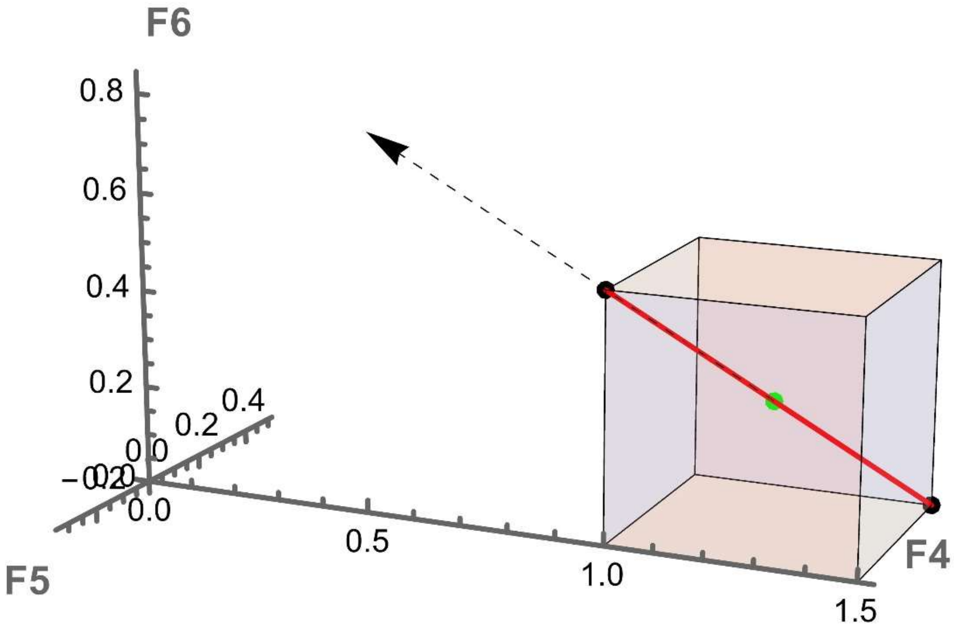

*Figure 12: SSKernel vs FVA bounding box. The coloured cube is the bounding box calculated by FVA for the the toy example network. The red line is the actual solutionspace (and also the SS kernel, since there are no rays).The SS centre and periphery points are shown as blue and green dots respectively. The dashed arrow is the unit vector that is delivered as the basis vector for the solution space, and defines its orientation in the 3D flux space formed by the variable fluxes F4, F5 and F6.*

The SSKernel calculation, by contrast, delivers a full and exact mathematical definition of the SS as a pair of 1D constraints relative to a specified coordinate origin and vector basis, as defined by equation (1.2). This specifies its shape as a line segment with a given orientation. Moreover, the explicit numerical specification of the SS centre point *SSKorigin* and orientation *SSKbasis* is directly readable from the DIF output file, as well as the numerical coordinates of the two periphery points. Also the report output shows graphically that the SS is one dimensional, and gives its single chord length and centering.
While these details are somewhat trivial in this particular case, already for the only slightly more challenging case illustrated in Figure 11 there is obviously far more information available in a similarly compact form, from the SSK description.

Some further examples of how the toy model can be manipulated to give explicit visualizations of bounded vs unbounded facets, various types of rays, and coincidence vs tangent capping, can be found in a forthcoming article [W. S. Verwoerd and L. Mao, *The FBA Solution Space Kernel – Introduction and Illustrative Examples*, bioRxiv 2024)].

### 9 Blocked reactions and external metabolites

A large part of the dimension reduction observed for the SSK of a typical metabolic model, is due to the elimination of many fixed fluxes. Most of these are in fact fixed to a value of zero – i.e., they represent blocked reactions. There are several ways that a reaction can become blocked, but a major cause is the absence of a source or sink reaction by means of which the network can exchange metabolites with the cellular environment.

Most models identify specific *external metabolites* that the network absorb from or deliver to the environment. For data imported from an SBML file, the cellular compartment is defined for every metabolite and is typically named "Extracellular" or similar for external metabolites. In the *.mat input there is no explicit provision for this, but a common convention is to append a qualifier such as "_e" or "_[e]" etc. to the names of external metabolites.

Conceptually, there is an infinite reservoir that buffers external metabolites so that despite being imported/exported, the metabolic network does not change their concentrations. This can be incorporated into the FBA calculation in one of two ways:

1) External metabolites are omitted from the list of metabolites for which the network maintains flux balance.
2) Explicit source or sink reactions are included in the model that either imports or exports each external metabolite (collectively called *exchange reactions*).

The SBML specification provides for the first method by assigning the attribute boundaryCondition = True to a metabolite. Deleting the row for such a metabolite from the stoichiometry matrix $S$ that defines the flux balance constraints $S.f = 0$, allows it to be freely imported or exported, but means that network reactions that involve this metabolite do not have a balanced stoichiometry any more.

The second buffering method, however, is preferred, and is followed by most models. Not only does this avoid disturbing the stoichiometry, but it also allows the direction and uptake or production rate to be set to realistic values by specifying flux bounds for the exchange reaction.

Examples of such explicit import and export reactions are seen in the toy network model discussed in section 8; there fluxes F7, F8 and F9 are unbalanced import/export fluxes. They are easily identified in the stoichiometry matrix of equation (1.3) by the fact that all non-zero entries in each corresponding column (usually just one) have the same arithmetic sign. This is in contrast to the main stoichiometrically balanced reactions which by definition have both inflows and outflows. Note that the exchange reactions are individually not stoichiometrically balanced, but their presence helps to ensure overall mass balance for the network as a whole.

The role they play is demonstrated, for example, by supposing that the last column of $S$, representing F9, is omitted. Then the second last row of $S$, representing metabolite *byp*, would contain only positive entries. Assuming all reactions to proceed in the forward direction as indicated on the network diagram, a nonzero flux value for either or both F2 and F3 would then violate the flux balance condition for *byp* that is contained in $S.f = 0$. In other words, in the absence of the sink reaction F9, reactions F2 and F3 become *blocked reactions* with fixed flux values of zero.

In general, inspection of just the stoichiometry matrix is not sufficient to identify such blocked reactions, because it does not contain information about reaction directions. It needs to be considered in conjunction with the flux bounds separately specified for each reaction in a metabolic model. In the above example, if the bounds indicated that either one of F2 and F3 runs in the reverse direction, or if either or both are bidirectional, it is in principle possible to satisfy the balance of *byp* by cancellation of their contributions to $S.f$ . Whether that can actually happen still depends on the joint effects of all other flux constraints on feasible values of F2 and F3, as determined by solving the full FBA problem. However, the point made here is that it is possible to determine by inspection, if the *structure* of a given stoichiometry matrix and flux bounds, a priori prohibits some flux components from achieving nonzero values. The term *blocked reaction* is for present purposes limited to these "structurally" prevented reactions, and the metabolite that requires this in order to satisfy the balance condition may be termed the *blocker*.

Anecdotally it is observed that in many current metabolic models, exchange reactions are missing for some of its external metabolites. But this does not necessarily make them blockers. It may still be possible to satisfy the balance condition for such a metabolite, in case it is involved in multiple stoichiometrically balanced reactions, some of which take up and others that produce the metabolite. The absence of a sink or source still restricts such a metabolite, because it requires that there is no net transfer to/from the environment, which is a more restrictive assumption than implied by its status as an external.

Moreover, surprisingly, in many models some of the exchange reactions that are provided by the model, are themselves actually blocked, due to the reaction directions implied by the flux bounds. As this makes them ineffective at the only purpose served by an exchange reaction, it seems most likely an unintended consequence.

In cases where the exchange reaction is absent or ineffective, there is a logical contradiction between specifying the metabolite as external, yet not allowing its free interchange with the environment. The reactions that consequently become blocked, result in a perhaps unjustified reduction in the dimensions of the SSK.

The best way to deal with this would be to revise the model by removing inconsistent flux bounds and adding missing exchange reactions. However, the task of the SSKernel program is to analyse the features of an existing model rather than to try to improve it. Such revisions would also require detailed knowledge of the applicable biochemistry, out of scope for automated correction.

As a compromise, the SSK implementation offers an option that reverts to the first way of representing the external metabolite reservoir: it allows exempting unbuffered external metabolites from the flux balance condition, by deleting their rows from the stoichiometry matrix. This may or may not result in a more realistic SSK, as discussed further below. At least, the comparison between the model with/without the exemptions allows an impression to be formed of whether the model is significantly compromised by defects in the buffering of external metabolites.

Internal metabolites might be assumed to be free of such difficulties, as flux balance is primarily designed to ensure mass conservation when metabolites are passed on from one metabolic network reaction to another. However, many models also provide exchange reactions for a few internal metabolites. One reason might be to avoid dealing with the details of currency metabolites such as ATP and NADH being exchanged between a great many individual reactions. Also, most models are presumably still incomplete, and may contain the origin but not the destination (or vice versa) of some obscure metabolites. Such a metabolite usually acts as a blocker, and large models often have many such. Providing an exchange reaction is a way to temporarily avoid this from happening until the model can be elaborated with the proper stoichiometric reaction or pathway.

In the SSK implementation, a summary in the form of number counts for internal exchange reactions and blockers are provided as part of the preliminary Source and Sink analysis of the stoichiometry matrix. However, these cases are taken as unavoidable parts of the model specification and no further action is taken on internal metabolites.

#### *9.1 How effective are exemptions of unbuffered externals?*

A common observation is that when exemptions are allowed, the number of fixed fluxes found by the SSK calculation decreases, and the SSK dimension count consequently increases. This is to be expected, as exemptions decrease the number of constraints. But the reduction is often quite large because blocking a single reaction usually blocks further reactions along a serial pathway.

The effects of this vary among different models. The best case, mostly observed in small and well curated models, is that there are no unbuffered externals and allowing exemptions makes no difference. Some examples have only a few exemptions and the rest of the SSK calculation proceeds without problems.

However, the more common case seems to be that there are 20 or more exemptions, leading to an even larger increase in SSK dimensions, and consequent problems. The removal of constraints can mean that the objective maximization becomes unbounded. Even if not, the FBA LP may have difficulty converging. Either of these cases can stop the SSK calculation in its tracks, although the latter can sometimes be alleviated by relaxing the LP tolerance.

Successful execution of the FBA may still leave the RSS with dimensions that exceeds the 150 – 200 limit for practical SSK calculation. The only remedy for this is to increase the fixed flux tolerance *fixtol*, as described elsewhere, but this may complicate the comparison of SSK's with/without exemptions.

Another problem case sometimes encountered, is a model that omits exchange reactions altogether. Perhaps this is meant to preserve the purity of the model by only including stoichiometrically balanced reactions. But usually this will yield an infeasible LP problem, or at best give only the trivial zero flux solution. All external metabolites are unbuffered for such a model, and exempting them is likely to make the LP problem unbounded so is no solution. The only viable solution in such a case is to elaborate the model by adding exchange reactions with realistic flux bounds for at least some of the exchange reactions.

To summarise, the examples suggest that choosing the exemptions option in the software implementation is helpful to explore the effects of unbuffered external metabolites on the SSK, for models where there is only a limited number of them.

However, perhaps the majority of models seem to have too many unbuffered externals to make this viable. In such cases a revision of the model, to remove inconsistent flux bounds and/or supplying missing exchange reactions with appropriate bounds, is the best plan of action.

### 10 Bioengineering and Sampling Solution Space Subregions

The SSKernel analysis gives a large simplification of the description of an FBA solution space, to a relatively low-dimensional bounded kernel supplemented by a set of ray vectors that account for its unbounded aspects. The set of chords and periphery points it produces as byproducts further helps to indicate the overall shape of the kernel.

In principle such insights should clearly be helpful for understanding and manipulating the metabolic state of the cell, for purposes such as bioengineering. In practice, however, it may not be obvious how best to exploit this knowledge.

The SSKernel software includes a *Mathematica* package, Resampling.m, that contains some tools to assist in this regard. This package is not explicitly used by the interactive user interface that terminates in the creation of the summary report and numerical output files. However, the ExploreKernel.nb notebook demonstrates how these tools can be applied afterwards to process the stored kernel space and ray data. If preferred, the tools can also be used in the control window after a SSKernel run as described below.

In fact, the resampling procedure runs autonomously once the hyperplanes to be sampled have been specified. This can be done in the notebook environment as done in ExploreKernel.nb, but alternatively a separate data file constructed as in the file *Template Hyperspec.wl* supplied in the CommandlineFiles subdirectory can be used. In this case the entire resampling analysis can be done in command line mode. For this purpose, the two files *Resamplebatch.bat* and *Resamplescript.wls* are also supplied in CommandlineFiles and are run in the same way as explained in section 6. The details are described in a later section.

To explain the rationale for resampling, it is easiest to start from a simple concrete example: suppose we wish to enhance production of a desired target metabolite *m* by knocking out a single gene. The knockout has the effect of fixing the flux $f_i$ of the corresponding reaction to a zero value. In terms of the solution space, it means that only its intersection with the flux space hyperplane defined by $f_i = 0$ remains feasible. So the question is, what effect does blocking reaction *i* have on the achievable values at which *m* can be exported from the metabolic network? This can be answered, at least approximately, by sampling representative fluxes on the hyperplane intersection mentioned above.

To perform this crucial task, a special function **HyperSampler** is provided in Resampling.m. This starts from the known set of points on the kernel periphery. By construction those are distributed to sample the periphery fairly uniformly, but the points will generally not fall on the desired hyperplane. Two distinct strategies are used to resample representative points that do fall on the hyperplane. Interpolating between the known points guarantees that the new points remain feasible, because of the convexity of the solution space, but only generates points on the intersection of the hyperplane with the PPP or periphery point polytope. To generate new points on the hyperplane intersection with the solution space outside of the PPP, extrapolation is done by including ray vectors that extend from the PPP. In this case it is necessary to enforce the FBA constraints on the extrapolation to ensure that the new point remains feasible. The set of resampled points generated by both interpolation and extrapolation is usually somewhat larger than the original periphery points set, and special computational strategies are employed to ensure that they are approximately uniformly distributed over the hyperplane intersection.

To answer the question about achievable values for the target metabolite *m*, the range of values shown by the resampled points gives an approximate indication, but the restricted point count limits the accuracy. This is addressed by explicitly calculating two more points that respectively minimizes and maximizes $f_m$, over the extent of the intersection, and adding these to the new sample. This gives an accurate range characterization. However, even if the range of $f_m$ values are the same for the complete SS and its $f_i = 0$ intersection, there may be more a more subtle shift in typical values. To get a handle on this, the mean value of $f_m$ taken over the augmented sample is also calculated. That gives an overall indication of the effects on the target flux, of a gene knockout that restricts the solution space to the $f_i = 0$ hyperplane.

Other interventions, such as restricting the nutrient concentration so that an uptake flux is fixed to a non-zero fixed value, is modelled similarly by resampling from the corresponding hyperplane intersection with a different fixed $f_{i\ value.}$

A more significant extension of the method is to consider cases where $f_i$ is restricted to a subrange of the values it traverses in the full SS, e.g. if conditions are chosen that makes an otherwise reversible reaction irreversible. Suppose for example that $f_i \geq a$ where *a* is a fixed numerical value. This translates to resampling from a subregion formed by using the $f_i = a$ hyperplane to partition the SS. Just like the full SS, this subregion is again a polytope, which may or may not be bounded. If not, it will have its own rayspace, which may coincide or be a subset of the SS rayspace. The resampling is again done by combining interpolation and extrapolation, while enforcing the additional $f_i \geq a$ constraint. But in addition, the ray space has to be separated into the rays used for extrapolation and those that remain valid rays in the subregion.

In fact, it is also true of the simpler case of the $f_i = 0$ hyperplane that it can be bounded or unbounded and so the remarks above about ray space partitioning applies also to the single knockout intervention.

Quite a number of scenarios can arise, and the *HyperSampler* function is designed to deal with all of them. In fact, it can even handle the more general case where interventions affect multiple fluxes, and the target flux may also be a combination of reaction fluxes, not just a single one. With multiple fluxes the constraint for each is set individually, so collectively may involve any combination of fixed values for some and range restrictions for others.

Hence quite apart from its use for investigating a particular bioengineering intervention, more generally *HyperSampler* allows considerable flexibility to explore different subregions of the solution space as long as these can be specified by a combination of linear flux constraints.

Returning to the simplest case of a single knockout, a plausible bioengineering question may be the more general one: Which single gene knockout will be most effective to enhance production of the target metabolite? The Resampling.m package contains a second function, named **TargetVariation** to investigate this. Given a list of all plausible single knockouts as input, it calls *HyperSampler* for each, and presents a sorted table of the resulting mean value and flux range of the target flux for each case.

The table shows at a glance which knockouts make the metabolism unviable, which remain feasible but leave the target unaffected, and the extent to which the rest affect the target flux, including any cases in which the target flux becomes fixed. Usually most of the latter involve the target flux being fixed to zero, so eliminated, and may be of particular interest if the bioengineering goal is to remove a particular metabolite.

#### 10.1 Interactive Notebook Implementation

The following examples demonstrate the use of these two functions to interactively analyze the RBC red blood cell model, that is supplied as a test case (see Section 4 and Figure 6). For routine use, it may be that the commandline implementation described in the next section is found more convenient.

From the SSK analysis, the RBC model has a sink reaction that exports water, namely EX_h2o_e and this has flux values that range from -6.3 to 4.6 (i.e., both import and export of water are feasible).

Suppose that we want a sample of metabolic flux states for which the RBC is in equilibrium, not exchanging water with its environment. Noting that EX_h20_e appears as column 37 of the stoichiometry matrix, this is represented by the flux space hyperplane where component 37 of the flux vector is zero.

The following code confirms the reaction identification and resamples the hyperplane. The first argument of *HyperSampler* gives the list of column numbers (here just a single one) and the second the corresponding list of constant values for each.

```
ReactionNames[[37]]
{hpoints,hrays}=HyperSampler[{37},{0.},PeriPoints,rays,ReducedSS];
```

Here, the arrays *PeriPoints*, *rays* and *ReducedSS* are assumed to have been assigned values when importing the *.dif output file from a completed SSKernel run for the model. If the analysis is done in the control window directly after the run, the arrays first need to be assigned their values by the instruction set

```
KernelSpace=ExportResults[[1]];
PeriPoints=ExportResults[[2]];
fixvals=ExportResults[[3]];
rays=ExportResults[[5]];
{MetaboliteNames,ReactionNames}=ExportResults[[8]];
ReducedSS=ExportResults[[9]];
```

As an example of extracting information from the resampled hyperplane points in the array *hpoints*, they can be used to investigate which of the fluxes that vary over the

SS, are significantly changed by fixing the net water transport to zero. The following code plots the differences, and lookup the names of those changed by more than 0.1 :

```
fluxdim=Length@KernelOrigin;
varfluxes = Complement[Range[fluxdim],fixvals[[All,1]]];
difs = Mean[PeriPoints[[All,varfluxes]]-
Mean[hpoints[[All,varfluxes]]];
BarChart[difs]
ReactionNames[[Flatten@Position[difs,_?(Abs[#]>0.1&)]]]
```

This produces the output shown in Figure 13. Many of the variable fluxes do change,

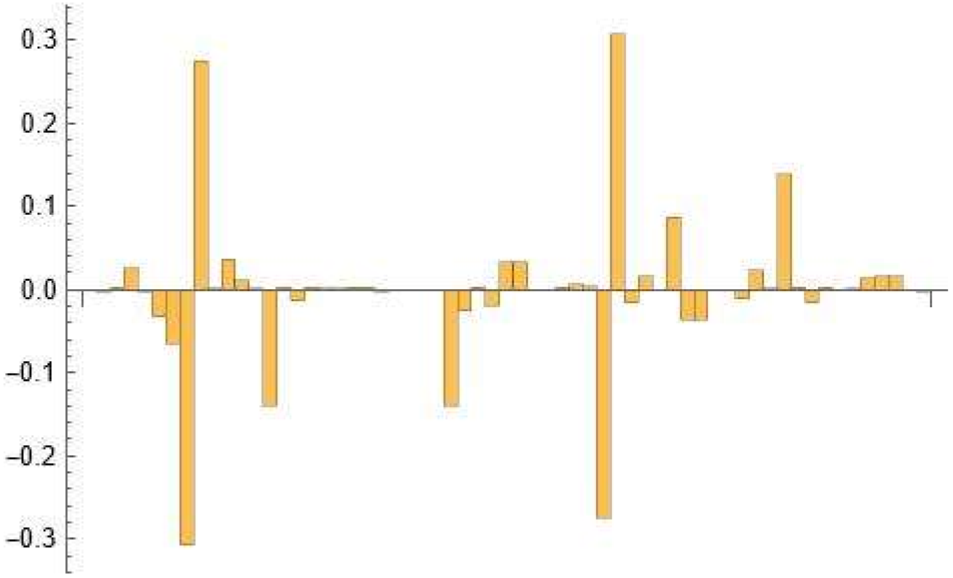

{"EX_ade_e", "EX_adn_e", "EX_bilglcur_e", "EX_fum_e", "EX_h2o2_e", "EX_hco3_e", "EX_man_e"}

*Figure 13: Differences between flux mean values taken over the $H_2O$ zero flux hyperplane, compared to the full solution space, for all variable fluxes. The reaction names of those that changed most are explicitly listed below the bar chart.*

and the ones most affected by restricting water exchange to zero are listed explicitly below the figure. The analysis could be repeated for the case where water exchange is restricted to be unidirectional, by just changing the call to *HyperSampler* to the form

```
{hpoints,hrays}=HyperSampler[{37},{{0.,1}},PeriPoints,rays,ReducedSS]
```

Here, the flux value restriction is specified by the pair expression {0., 1} where the first number gives the value and the second is -1, 0, or 1 to denote ≤ , = or ≥ respectively.

Moving on to an example of applying *TargetVariation* to the RBC model, note that since as the biological function of the red blood cell is to transport oxygen, a natural choice for the target is oxygen export. So we address the question, what would be the effect of knocking out individual genes for enzymes that control fluxes, on the oxygen export flux?

The first step is to install the target into the FBA model, which is performed by the utility function **TargetInsert** supplied by the Resampling package. This function allows the target to be specified in one of three ways:

a) A text string contained in the metabolite name.
b) The column number of a metabolite flux present in the stoichiometry matrix.
c) An explicit target flux as a vector in the flux space.

Depending on the specification, the target flux may or may not already occur in the model. If not, *TargetInsert* will add it as an extra column in the stoichiometry matrix *S* and in this case the SSKernel analysis needs to be repeated before proceeding. However, if it already occurs inserting a duplicate flux would introduce spurious flux variability, so *TargetInsert* checks for this and where possible simply identifies an existing column, else alerting the user to the need to rerun SSKernel using the augmented *S*.

For our example, the insertion step could be performed by the instruction

```
targetcol = TargetInsert["O2"]
```

It turns out that this is ambiguous since the model also contains CO2 and H2O2 as metabolites. *TargetInsert* recognizes this, so leaves the model unchanged and presents both the full names and any existing exchange reaction column numbers for each in an alert box, so the user can decide which to apply. For the RBC model, it identifies column 60 to correspond to oxygen export so a repeat call needs to made such as

```
targetcol = TargetInsert[60]
```

As this is a preexisting column, we can directly proceed to invoking *TargetVariation*. As input, this needs to be given a set of hyperplanes; for each it calls *Hypersampler* and collects relevant statistics from the resulting sample points, and sorts these by the mean value of the target flux. *TargetVariation* prints this as a formatted table as its final output while also returning all sample points it found for further processing.

For single knockouts, the zero value hyperplanes for all fluxes in the model could be specified as the set of cases for *TargetVariation* to inspect. However, explicitly fixing fluxes that already have fixed values over the SS would be either redundant or infeasible in the case the SS fixes them to a nonzero value. So, such knockouts should be confined to fluxes that were determined by the SSK analysis to remain variable. For the RBC model, that reduces the number of hyperplanes in the input list to 61 out of the total of 469 model fluxes. For this small model that is quite tractable, but for larger models biochemical knowledge may need to be employed to narrow down the set of interventions that are probed.

The following code performs the calculation as described:

```
(* Collect variable flux numbers *)
cols=Length@ReducedSS[[2,1]];
nonfixes=Complement[Range[cols],fixvals[[All,1]]];

(* Set up the list of hyperplanes to be sampled, as each
variable flux in turn fixed to zero*)
hyperlist = Partition[nonfixes, 1];
valuelist = ConstantArray[{0}, Length@hyperlist];

(* Sample each hyperplane and collect data on target flux *)
samples = TargetVariation[hyperlist, valuelist, PeriPoints,
rays,ReducedSS, targetcol, False];
```

The output appears as shown in Figure 14. For context, the mean value and range of the target flux (oxygen export) for the undisturbed SS is shown at the top of the table. At the bottom of the table, a number of knockouts appear that produce no samples, i.e. they render the model infeasible. Above them are some that produce a fixed target flux (since the upper and lower range limits are the same). Most of the rest leave the target flux range unchanged relative to the SS at 0 to 5, although the mean value is increased in somewhat fewer than half of them.

The most effective intervention, though, seems to be knocking out reactions H2Ot or EX_h2o_e, since in either case this will force an export flux of at least 2.32 units. At the same time the upper flux limit is reduced, so that the mean value is not greatly different from the original SS .

The **Formula** function supplied in the Resampling package gives a convenient way to inspect the chemical reaction represented by a particular flux; e.g. in this case the instruction

```
Formula["H2Ot"]
Formula["EX_h2o_e"]
```

produces the output

H2Ot: h2o_e → h2o_c
EX_h2o_e: h2o_e →

The *Formula* function accepts either a character string contained in a reaction name, or a number or list of numbers that specify the column position of the reaction(s) in the stoichiometry matrix.

This result illustrates how the SSK analysis can be used to interrogate a metabolic model to identify promising bioengineering avenues.

Further investigation is obviously needed to establish whether such interventions are practical, and the considerations may require biochemical expertise and extend beyond the limits of the model.

Even within its limits though, more information could still be extracted to help with this. For example, the question may arise how other fluxes would be affected if e.g. EX_h2o_e is knocked out.

Note that after running the code above, the actual sample points that were generated during the course of the *TargetVariation* run are available in a nested list form as the calculated variable *samples*. Each element of *samples* consists of the sample points collected for the corresponding hyperplane in the sequence as listed and numbered in the table.

For example, `samples[[20]]` would give sample fluxes resulting from the EX_h2o_e knockout. Any specific flux component in these gives an indication of how much it was affected, although the actual range of values may be wider than encountered in these samples. If that is important, the full range could be discovered by repeating the analysis using that particular flux as the target.

| Trial No | ASSIGNED Fluxes | Values | SAMPLE Count | Target: EX_o2_e Mean | Lower | Upper |
|---|---|---|---|---|---|---|
| 1 | SS Kernel | -- | 64 | 2.61 | 0 | 5. |
| 2 | {EX_ade_e} | {0} | 64 | 3.25 | 0 | 5. |
| 3 | {ME2} | {0} | 4 | 3.19 | 0 | 5. |
| 4 | {ADEt} | {0} | 64 | 3.13 | 0 | 5. |
| 5 | {PUNP1} | {0} | 64 | 3.13 | 0 | 5. |
| 6 | {FUMtr} | {0} | 64 | 3.02 | 0 | 5. |
| 7 | {FUM} | {0} | 64 | 3.02 | 0 | 5. |
| 8 | {EX_fum_e} | {0} | 64 | 3.01 | 0 | 5. |
| 9 | {UREAt} | {0} | 10 | 2.77 | 0 | 5. |
| 10 | {PTRCtex2} | {0} | 10 | 2.77 | 0 | 5. |
| 11 | {ORNDC} | {0} | 10 | 2.77 | 0 | 5. |
| 12 | {ARGt5r} | {0} | 10 | 2.77 | 0 | 5. |
| 13 | {ARGN} | {0} | 10 | 2.77 | 0 | 5. |
| 14 | {EX_urea_e} | {0} | 10 | 2.77 | 0 | 5. |
| 15 | {EX_ptrc_e} | {0} | 10 | 2.77 | 0 | 5. |
| 16 | {EX_arg__L_e} | {0} | 10 | 2.77 | 0 | 5. |
| 17 | {SBTR} | {0} | 14 | 2.76 | 0 | 5. |
| 18 | {SBTD_D2} | {0} | 14 | 2.76 | 0 | 5. |
| 19 | {H2Ot} | {0} | 64 | 2.75 | 2.32 | 3.14 |
| 20 | {EX_h2o_e} | {0} | 64 | 2.75 | 2.32 | 3.14 |
| 21 | {GTHPi} | {0} | 13 | 2.7 | 0 | 5. |
| 22 | {MALt} | {0} | 64 | 2.69 | 0 | 5. |
| 23 | {EX_mal__L_e} | {0} | 64 | 2.69 | 0 | 5. |
| 24 | {PUNP5} | {0} | 17 | 2.68 | 0 | 5. |
| 25 | {NH4t3r} | {0} | 17 | 2.68 | 0 | 5. |
| 26 | {HYXNt} | {0} | 17 | 2.68 | 0 | 5. |
| 27 | {ADA} | {0} | 17 | 2.68 | 0 | 5. |
| 28 | {EX_nh4_e} | {0} | 17 | 2.68 | 0 | 5. |
| 29 | {EX_hxan_e} | {0} | 17 | 2.68 | 0 | 5. |
| 30 | {GTHOr} | {0} | 8 | 2.66 | 0 | 5. |
| 31 | {LNLCCPT2rbc} | {0} | 64 | 2.61 | 0 | 5. |
| 32 | {LNLCCPT1} | {0} | 64 | 2.61 | 0 | 5. |
| 33 | {GALUi} | {0} | 64 | 2.61 | 0 | 5. |
| 34 | {GALT} | {0} | 64 | 2.61 | 0 | 5. |
| 35 | {C181CPT2rbc} | {0} | 64 | 2.61 | 0 | 5. |
| 36 | {C181CPT1} | {0} | 64 | 2.61 | 0 | 5. |
| 37 | {C160CPT2rbc} | {0} | 64 | 2.61 | 0 | 5. |
| 38 | {C160CPT1} | {0} | 64 | 2.61 | 0 | 5. |
| 39 | {GTHDH} | {0} | 20 | 2.59 | 0 | 5. |
| 40 | {DHAAt1r} | {0} | 20 | 2.59 | 0 | 5. |
| 41 | {ASCBt} | {0} | 20 | 2.59 | 0 | 5. |
| 42 | {EX_dhdascb_e} | {0} | 20 | 2.59 | 0 | 5. |
| 43 | {EX_ascb__L_e} | {0} | 20 | 2.59 | 0 | 5. |
| 44 | {DM_nadh} | {0} | 10 | 2.35 | 0 | 5. |
| 45 | {PYRt2} | {0} | 5 | 2.21 | 0 | 5. |
| 46 | {EX_pyr_e} | {0} | 5 | 2.21 | 0 | 5. |
| 47 | {O2t} | {0} | 2 | 0 | 0 | 0 |
| 48 | {H2O2t} | {0} | 2 | 0 | 0 | 0 |
| 49 | {CAT} | {0} | 2 | 0 | 0 | 0 |
| 50 | {Target: EX_o2_e} | {0} | 2 | 0 | 0 | 0 |
| 51 | {EX_h2o2_e} | {0} | 2 | 0 | 0 | 0 |
| 52 | {UGLT} | {0} | 0 | -- | -- | -- |
| 53 | {PGI} | {0} | 0 | -- | -- | -- |
| 54 | {LDH_L} | {0} | 0 | -- | -- | -- |
| 55 | {L_LACt2r} | {0} | 0 | -- | -- | -- |
| 56 | {Ht} | {0} | 0 | -- | -- | -- |
| 57 | {HEX7} | {0} | 0 | -- | -- | -- |
| 58 | {HEX1} | {0} | 0 | -- | -- | -- |
| 59 | {CO2t} | {0} | 0 | -- | -- | -- |
| 60 | {EX_lac__L_e} | {0} | 0 | -- | -- | -- |
| 61 | {EX_h_e} | {0} | 0 | -- | -- | -- |
| 62 | {EX_co2_e} | {0} | 0 | -- | -- | -- |

*Figure 14 Variation of the target flux " EX_o2_e " in response to individual flux knockouts for the RBC model.*

In the usual case that the RSS is unbounded, the target flux on a hyperplane or subregion that is sampled may or may not be bounded. If it is not, that will show up in the TargetVariation output table as an Infinity symbol for one of the flux limits. Also, the corresponding set of sample points will contain a couple of points with an Infinite value for the target flux component. Such sample points many need to be filtered out before using them for further numerical calculations. The majority of sample points are by construction located on the finite boundaries of the RSS/hyperplane intersection, although there may also be some points in its interior (such as inherited periphery points located on the SSK capping hyperplane). Further interior points can always be generated by convex combinations of the sample points.

Note that any sample points "at infinity" do not define the directions in which RSS/hyperplane intersection is infinite. To discover that, it is necessary to call *HyperSampler* which returns both sample points and a ray basis for the intersection.

Finally, note that the same method can be used to investigate far more elaborate interventions than single gene knockouts; all cases discussed in connection with *HyperSampler* can be used. The only change necessary is to modify the *hyperlist* and *valuelist* specifications in the code above so that each element in these lists conforms to the requirements of *HyperSampler*.
As an example, here is a hypothetical specification that could be used to generate a target variation table:

```
hyperlist={{20},{249},{20,22,27}};
valuelist={{-5.3},{{0.35,0}},{{-5.3,1},-0.05,{-0.2,-1}}};
```

Three different hyperplanes are listed in this specification. On the first, flux no 20 is kept fixed to the non-zero value -5.3. The second specification shows an alternative way to fix flux 249 to the value 0.35, by explicitly invoking an equality constraint. The third case demonstrates a subregion, specified by a mixture of inequalities and equalities: flux 20 is kept above -5.3, flux 22 is kept fixed at the value -0.05, and flux 27 remains below -0.2.

#### 10.2 Commandline Implementation

To do the resampling analysis in a commandline environment, the first step is to create a data file that specifies the target and hyperplanes to be sampled. A template file for this is supplied as Template Hyperspec.wl in the CommandlineFiles subdirectory. This file is to be copied to the directory where the model data file is located, and appropriately renamed as e.g. "iAB_RBC_283 HyperSpec.wl" for the red blood cell model discussed above. The following content of this file produces the output shown in Figure 14 :

```
target=60;
targetname="O2 export";
targetproduced=True;
(* Set up the list of hyperplanes to be sampled, as each
variable flux is in turn fixed to zero*)
cols = Length@ReducedSS[[2, 1]];
nonfixes = Complement[Range[cols], fixvals[[All, 1]]];
hyperlist = Partition[nonfixes, 1];
valuelist = ConstantArray[{0}, Length@hyperlist] ;
```

The goal of resampling is to analyse the solution space, and thus its primary input is the *XXX SSKernel.dif* file that contains the SSK and ray vector descriptions and is generated by the full SSKernel analysis. It also needs two secondary input files: the original model specification, e.g. *iAB_RBC_283.mat* from which it needs to read the stoichiometry matrix, and *iAB_RBC_283 HyperSpec.wl* as described.

Assuming that the SSKernel run has been completed, the simplest way to do the resampling analysis is to right click on *iAB_RBC_283 SSKernel.dif* and use the Open With menu to open it with the batch file *Resamplingbatch.bat.* This invokes the corresponding script file, that automatically finds and loads all 3 input files and calls the *TargetInsert* function.

In the simplest case, as for the iAB_RBC_283 example, the target flux is already present in the model and the user only needs to decide whether the *HyperSampler* function or *TargetVariation* needs to be invoked. In the latter case the output table is presented, and the actual sample points can optionally be stored in a separate file. In the former case, any of the hyperplanes specified in the HyperSpec file can be chosen to be sampled, and both the sample points and the rays in this hyperplane are optionally stored in a file.

It is slightly more complicated in cases where a new flux has to be created for the target metabolite. The resampling script performs this automatically, but then prompts the user that since the stoichiometry matrix has been modified, a new run of SSKernel is required. The input for this is stored as a new file, e.g. *iAB_RBC_283 target.m*. Then, SSKernel is run on this input file either using the interactive interface, or by right-clicking on the …target.m file and opening it with *SSKernelbatch.bat*. Once that is done, a new .dif file such as *iAB_RBC_283 target SSKernel.dif* has been created in the data directory. Then this file is in turn opened with the *Resamplingbatch.bat* file to make the final resampling analysis.

An advantage of running the resampling in commandline mode is that the sequence of operations is controlled by the script file and the user merely has to follow the prompts that are presented.
