## Supplementary material for "The FBA Solution Space Kernel – Introduction and Illustrative Examples": Input data and workflow

### Toy Model Input Data

The input data for the toy model defined in Figure 1 of the main article, is as follows:

Toy network

{"Reaction 1", "Reaction 2", "Reaction 3", "Reaction 4", "Reaction 5", "Reaction 6", "Reaction 7", "Reaction 8", "Reaction 9"}

{{-1, 0, 0, 0, 0, 0, 1, 0, 0}, {1, -2, -2, 0, 0, 0, 0, 0, 0}, {0, 1, 0, 0, -1, -1, 0, 0, 0}, {0, 0, 1, -1, 1, 0, 0, 0, 0}, {0, 0, 0, 1, 0, 1, 0, -1, 0}, {0, 1, 1, 0, 0, 0, 0, 0, -1}, {0, 0, -1, 1, -1, 0, 0, 0, 0}}

{{0, 0}, {0, 0}, {0, 0}, {0, 0}, {0, 0}, {0, 0}, {0, 0}}

{{0., Infinity}, {0., Infinity}, {0., Infinity}, {0., Infinity}, {0, Infinity}, {0., Infinity}, {0., Infinity}, {0., Infinity}, {0., Infinity}}

{0, 0, 0, 0, 0, 0, 0, 0, 0}

{0., 0., 0., 0., 0., 0., 0., 0., 0.}

{"A","B","C","D","E","byp","cof"}

max

This should be saved as a simple text file, such as "Scenario1.m" and used as input to the interactive SSKernel Mathematica software package to generate the kernel and ray specifications plotted in Figures 2 – 6.

The file extension *.m signals to SSKernel that the input data is in the Mathematica native code syntax, rather than the more involved *.mat files that are commonly used for realistic metabolic models.

The data above defines the case labeled as Scenario 2 in the article. Each of the other scenarios requires small data modifications as described in the text. The resulting data sets are as follows.

#### Scenario 1

F7 is restricted to an upper limit of 3.0 and F2 kept to a fixed value 0.5; biomass flux F10 added.

Toy network, FVA comparison

{"Reaction 1", "Reaction 2", "Reaction 3", "Reaction 4", "Reaction 5", "Reaction 6", "Reaction 7", "Reaction 8", "Reaction 9","Biomass"}

{{-1, 0, 0, 0, 0, 0, 1, 0, 0, 0}, {1, -2, -2, 0, 0, 0, 0, 0, 0, 0}, {0, 1, 0, 0, -1, -1, 0, 0, 0, 0}, {0, 0, 1, -1, 1, 0, 0, 0, 0, 0}, {0, 0, 0, 1, 0, 1, 0, -1, 0., -0.8}, {0, 1, 1, 0, 0, 0, 0, 0, -1, -0.2}, {0, 0, -1, 1, -1, 0, 0, 0, 0, 0}}

{{0, 0}, {0, 0}, {0, 0}, {0, 0}, {0, 0}, {0, 0}, {0, 0}}

{{0., Infinity}, {0.5, 0.5}, {0., Infinity}, {0., Infinity}, {0, Infinity}, {0., Infinity}, {0, 3.0}, {0., Infinity}, {0., Infinity}, {0., Infinity}}

{0, 0, 0, 0, 0, 0, 0, 0, 0, 1}

{0., 0., 0., 0., 0., 0., 0., 0., 0., 0.}

{"A","B","C","D","E","byp","cof"}

max

#### Scenario 3

F7 is restricted to an upper limit of 3.0:

Toy network, uptake limit

{"Reaction 1", "Reaction 2", "Reaction 3", "Reaction 4", "Reaction 5", "Reaction 6", "Reaction 7", "Reaction 8", "Reaction 9"}

{{-1, 0, 0, 0, 0, 0, 1, 0, 0}, {1, -2, -2, 0, 0, 0, 0, 0, 0}, {0, 1, 0, 0, -1, -1, 0, 0, 0}, {0, 0, 1, -1, 1, 0, 0, 0, 0}, {0, 0, 0, 1, 0, 1, 0, -1, 0}, {0, 1, 1, 0, 0, 0, 0, 0, -1}, {0, 0, -1, 1, -1, 0, 0, 0, 0}}

{{0, 0}, {0, 0}, {0, 0}, {0, 0}, {0, 0}, {0, 0}, {0, 0}}

{{0., Infinity}, {0., Infinity}, {0., Infinity}, {0., Infinity}, {0, Infinity}, {0., Infinity}, {0., 3.0}, {0., Infinity}, {0., Infinity}}

{0, 0, 0, 0, 0, 0, 0, 0, 0}

{0., 0., 0., 0., 0., 0., 0., 0., 0.}

{"A","B","C","D","E","byp","cof"}

max

#### Scenario 4

In addition, add a further export reaction labelled "Biomass" as flux F10 that combines metabolites E and byp in a 4:1 ratio. Define this as the objective to be maximized.

Toy network, maximize biomass

{"Reaction 1", "Reaction 2", "Reaction 3", "Reaction 4", "Reaction 5", "Reaction 6", "Reaction 7", "Reaction 8", "Reaction 9","Biomass"}

{{-1, 0, 0, 0, 0, 0, 1, 0, 0, 0}, {1, -2, -2, 0, 0, 0, 0, 0, 0, 0}, {0, 1, 0, 0, -1, -1, 0, 0, 0, 0}, {0, 0, 1, -1, 1, 0, 0, 0, 0, 0}, {0, 0, 0, 1, 0, 1, 0, -1, 0., -0.8}, {0, 1, 1, 0, 0, 0, 0, 0, -1, -0.2}, {0, 0, -1, 1, -1, 0, 0, 0, 0, 0}}

{{0, 0}, {0, 0}, {0, 0}, {0, 0}, {0, 0}, {0, 0}, {0, 0}}

{{0., Infinity}, {0., Infinity}, {0., Infinity}, {0., Infinity}, {0, Infinity}, {0., Infinity}, {0, 3.0}, {0., Infinity}, {0., Infinity}, {0., Infinity}}

{0, 0, 0, 0, 0, 0, 0, 0, 0, 1}

{0., 0., 0., 0., 0., 0., 0., 0., 0., 0.}

{"A","B","C","D","E","byp","cof"}

max

#### Scenario 5

Again the Scenario 4 model with a biomass objective is used, but now F2 is given a 0.5 lower limit and F3 is made reversible by removing its lower limit.

Toy network, biomass and single ray

{"Reaction 1", "Reaction 2", "Reaction 3", "Reaction 4", "Reaction 5", "Reaction 6", "Reaction 7", "Reaction 8", "Reaction 9","Biomass"}

{{-1, 0, 0, 0, 0, 0, 1, 0, 0, 0}, {1, -2, -2, 0, 0, 0, 0, 0, 0, 0}, {0, 1, 0, 0, -1, -1, 0, 0, 0, 0}, {0, 0, 1, -1, 1, 0, 0, 0, 0, 0}, {0, 0, 0, 1, 0, 1, 0, -1, 0., -0.8}, {0, 1, 1, 0, 0, 0, 0, 0, -1, -0.2}, {0, 0, -1, 1, -1, 0, 0, 0, 0, 0}}

{{0, 0}, {0, 0}, {0, 0}, {0, 0}, {0, 0}, {0, 0}, {0, 0}}

{{0., Infinity}, {0.5, Infinity}, {-Infinity, Infinity}, {0., Infinity}, {0, Infinity}, {0., Infinity}, {0, 3.0}, {0., Infinity}, {0., Infinity}, {0., Infinity}}

{0, 0, 0, 0, 0, 0, 0, 0, 0, 1}

{0., 0., 0., 0., 0., 0., 0., 0., 0., 0.}

{"A","B","C","D","E","byp","cof"}

max

#### Scenario 6

This replicates scenario 5, but with reaction 6 now also made reversible. The data specification is

Toy network, biomass and 2 rays

{"Reaction 1", "Reaction 2", "Reaction 3", "Reaction 4", "Reaction 5", "Reaction 6", "Reaction 7", "Reaction 8", "Reaction 9","Biomass"}

{{-1, 0, 0, 0, 0, 0, 1, 0, 0, 0}, {1, -2, -2, 0, 0, 0, 0, 0, 0, 0}, {0, 1, 0, 0, -1, -1, 0, 0, 0, 0}, {0, 0, 1, -1, 1, 0, 0, 0, 0, 0}, {0, 0, 0, 1, 0, 1, 0, -1, 0., -0.8}, {0, 1, 1, 0, 0, 0, 0, 0, -1, -0.2}, {0, 0, -1, 1, -1, 0, 0, 0, 0, 0}}

{{0, 0}, {0, 0}, {0, 0}, {0, 0}, {0, 0}, {0, 0}, {0, 0}}

{{0., Infinity}, {0.5, Infinity}, {-Infinity, Infinity}, {0., Infinity}, {0, Infinity}, {-Infinity, Infinity}, {0, 3.0}, {0., Infinity}, {0., Infinity}, {0., Infinity}}

{0, 0, 0, 0, 0, 0, 0, 0, 0, 1}

{0., 0., 0., 0., 0., 0., 0., 0., 0., 0.}

{"A","B","C","D","E","byp","cof"}

max

#### Toy model with target metabolite cof

This is the case used in section 3.2.1 of the article to demonstrate the use of resampling. The SSKernel input file is the one used for scenario 5, but with the stoichiometry matrix modified to allow excess production of the chosen target metabolite cof as follows:

Toy network, cof production

{"Reaction 1", "Reaction 2", "Reaction 3", "Reaction 4", "Reaction 5", "Reaction 6", "Reaction 7", "Reaction 8", "Reaction 9", "Biomass", "Export cof"}

{"A", "B", "C", "D", "E", "byp", "cof"}

{{-1, 0, 0, 0, 0, 0, 1, 0, 0, 0, 0.}, {1, -2, -2, 0, 0, 0, 0, 0, 0, 0, 0.}, {0, 1, 0, 0, -1, -1, 0, 0, 0, 0, 0.}, {0, 0, 1, -1, 1, 0, 0, 0, 0, 0, 0.}, {0, 0, 0, 1, 0, 1, 0, -1, 0., -0.8, 0.}, {0, 1, 1, 0, 0, 0, 0, 0, -1, -0.2, 0.}, {0, 0, -2, 2, -1, 0, 0, 0, 0, 0, -1.}}

{{0, 0}, {0, 0}, {0, 0}, {0, 0}, {0, 0}, {0, 0}, {0, 0}}

{{0., Infinity}, {0.5, Infinity}, {-Infinity, Infinity}, {0., Infinity}, {0, Infinity}, {0., Infinity}, {0, 3.}, {0., Infinity}, {0., Infinity}, {0., Infinity}, {0., Infinity}}

{0, 0, 0, 0, 0, 0, 0, 0, 0, 1, 0}

{0., 0., 0., 0., 0., 0., 0., 0., 0., 0.}

max

### Workflow

To reproduce the figures and tables in the article text, is a 2-stage process: first the kernel specification is generated by the SSKernel software (Verwoerd, 2024) (steps 1 – 4 below) and then this result is plotted or analysed by resampling.

**Computing environment.** Implemented in [Wolfram Language](https://www.wolfram.com/language/), the full utilization of the SSKernel software (Verwoerd, 2024) including its interactive control interface, does require installation of the proprietary *Mathematica* system (Wolfram_Research & Inc., 2024), but will hence run on all the operating systems for which *Mathematica* is available. Work is in progress to produce a stand-alone version of the software and make this available in due course as part of the GitHub release.

However, for reproducing the results of the main article a command line implementation is perfectly adequate and for that only the installation of the [Wolfram Engine](https://www.wolfram.com/engine/) is required. This can be downloaded for free, though subject to use restrictions that do allow testing applications. It is also available for online use via a Wolfram cloud server.

Assuming that the either of these requirements are met, the workflow reduces to the following steps:

1. Download the SSKernel software from Zenodo.
2. Copy and paste the required data from Section 2 above into a data file *ScenarioX.m* located in a convenient working directory.
3. Locate *ScenarioX.m* in Windows Explorer, right click to bring up the context menu, then use the "Open with" submenu to open the data file with the batch file *SSKernelbatch.bat* located in the subdirectory *CommandlineFiles*.
4. This will open a standard command window, load and parse the data. Follow the prompt to allow the calculation to proceed. At completion two output files are created in the working directory, a PDF file containing a summary report and the file *ScenarioX SSKernel.dif* that contains the kernel and RSS specifications and other supporting data for further analysis.
5. Open the *ToyPlots.nb* file (included as a separate supplementary file) with the *Mathematica* software system.

Use the Browse button it displays, to identify the applicable *.dif file and then execute the data loading cell in the notebook. Click the dropdown arrow for the appropriate section and execute the section to generate the desired figure in the article.

If full access to *Mathematica* is not available, see the notes below on alternatives.

6. Similarly, the *.dif file resulting from the data in section 2.6, is used as input for the resampling calculation discussed in the main article, Section 3.2.

Table 1 in that section is produced from the *.dif file produced for the data listed in Section 2.6. Open this file similarly as described in step 3 above, but with the different batch file *Resampling.bat* also found in the *CommandlineFiles* subdirectory.

7. Table 2 in the article is generated similarly from the RBC model supplied with the SSKernel software, in the *TEST* subdirectory.

**Note on plotting.** The plots are most easily produced as described using *ToyPlots.nb* in the *Mathematica* notebook environment.

However, any graphical software capable of rendering inequality specified polygons and polyhedra should be able to display the 2D and 3D SSK volumes in an equivalent way. The inequality coefficients are listed as matrices and vectors in the *.dif file. The SSK data is structured as in Equation above. Even just line plots of linear equations will suffice for the 2D polygons.

Note that these files are in the spreadsheet-oriented Data Interchange Format, and are simple text files. If not readable by the chosen graphics package, it can simply be opened in a text editor. There are explanatory headings, some text tags used by DIF readers, and the actual data are simply comma separated lists enclosed in curly brackets. Each such list denotes a vector. Matrices are represented as lists of row vectors, in a second nested level of curly brackets.

Plotting of periphery points, chords and rays require minor data adjustments carried out in the *ToyPlots.nb* notebook by some utility functions it contains. To make their coding accessible outside of Mathematica, the notebook is also supplied as a PDF document in a separate supplementary file.
